# Efficient clustering of large molecular libraries

**DOI:** 10.1101/2024.08.10.607459

**Authors:** Kenneth López Pérez, Vicky Jung, Lexin Chen, Kate Huddleston, Ramón Alain Miranda-Quintana

**Affiliations:** Department of Chemistry & Quantum Theory Project, University of Florida, Gainesville, Florida 32611

**Keywords:** clustering, similarity, chemical diversity, chemical space

## Abstract

The widespread use of Machine Learning (ML) techniques in chemical applications has come with the pressing need to analyze extremely large molecular libraries. In particular, clustering remains one of the most common tools to dissect the chemical space. Unfortunately, most current approaches present unfavorable time and memory scaling, which makes them unsuitable to handle million- and billion-sized sets. Here, we propose to bypass these problems with a time- and memory-efficient clustering algorithm, BitBIRCH. This method uses a tree structure similar to the one found in the Balanced Iterative Reducing and Clustering using Hierarchies (BIRCH) algorithm to ensure O(*N*) time scaling. BitBIRCH leverages the instant similarity (iSIM) formalism to process binary fingerprints, allowing the use of Tanimoto similarity, and reducing memory requirements. Our tests show that BitBIRCH is already > 1,000 times faster than standard implementations of the Taylor-Butina clustering for libraries with 1,500,000 molecules. BitBIRCH increases efficiency without compromising the quality of the resulting clusters. We explore strategies to handle large sets, which we applied in the clustering of one billion molecules under 5 hours using a parallel/iterative BitBIRCH approximation.

## INTRODUCTION

Clustering is an unsupervised machine learning (ML) technique that organizes unlabeled data into groups, or clusters, of related points. (1–3) This can be critical to obtain insights into the structure and organization of the data (3), which could also be used to aid the development of supervised ML models(4, 5). This is essential in developing more accurate predictive models.(4) In chemical applications, during the data generation and curation process, clustering can assist in identifying regions of chemical space that are lacking within the dataset.(6, 7) By ensuring that a representative range of chemical space is covered in the training and validation datasets, one can reduce biases and ensure that the model performs well on diverse data, broadening generalizability and enhancing the robustness of the model.(8)

In drug design, the importance of clustering is highlighted when considering the “molecular similarity principle”, which states that structurally similar molecules will often share similar chemical properties or biological activities.(9–11) By identifying similar compounds, clustering can help predict the behavior and properties of existing and new structures, expediting virtual screening.(12) The preponderant molecular representations in drug design are binary fingerprints(13), and Tanimoto similarity(14, 15) is widely used in cheminformatics tasks(16–19), including clustering.(20, 21) Fingerprints encode molecular information using bitstrings (arrays of *on* and *off* bits)(13), this simplicity is particularly attractive in the exploration of large chemical spaces, but also leads to robust results in Structure-Activity Relationship (SAR) studies.(22–24) There are multiple ways to generate fingerprints from a molecular graph or structure (MACCS(25), ECFP(26), RDKit(27), MAP(28), etc.), so it is important to emphasize that the results discussed below apply to any binary encoding. From this type of representation, it is easy to calculate the Tanimoto similarity(10, 14) between any two molecules *A* and *B*, T(*A*, *B*), as:

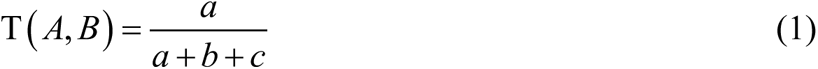

where *a* indicates the number of common *on* bits between *A* and *B*’s bitstrings, and *b* + *c* counts the *on*/*off* mismatches between them. While there are many possibilities to quantify the similarity between molecules, Tanimoto is the de facto option in the cheminformatics community.(18)

Pharmaceutical applications typically use one of a handful of clustering algorithms, with spectral(29, 30), hierarchical(31, 32), and Jarvis-Patrick(33, 34) clustering as common choices. However, arguably the most popular option is the Taylor-Butina clustering. All of these methods rely on the construction of the similarity matrix between the molecules, and then use some notion of neighborhood/locality to group the points.(20) Taylor-Butina does this in perhaps the simplest and most intuitive way, requiring a single parameter: a similarity threshold used to count the number of neighbors of each potential cluster centroid.(20, 35) At every iteration, the molecule with most neighbors is identified as the best potential centroid and, together with all its available neighbors, they form a cluster. This procedure is repeated until exhausting all molecules, or until no molecules remain that are closer than the pre-specified threshold. This results in clusters that are easily interpretable, as they contain molecules that are closely related to the corresponding centroid.(20) However, the fact that we need to compute the similarity matrix means that all of these methods scale as O(*N*^2^) in both time and memory(36), with this scaling potentially preventing their application to ever-increasing sectors of chemical space.

These time and memory bottlenecks are not unique to chemical applications of clustering, so it is not strange that methods with more favorable scaling have proliferated. One particularly attractive alternative is the Balanced Iterative Reducing and Clustering using Hierarchies (BIRCH) algorithm.(37) BIRCH solves the memory problems by using a Clustering Feature (CF) to encapsulate the cluster information in a compact way that still allows computing critical indicators, like the cluster centroids, radius, and diameter.(37) The time scaling is then improved using a CF-tree data structure, that allows to efficiently distribute the molecules into their corresponding clusters: the “leaves” of the tree.(37) (There has been a renewed interest in the cheminformatics/drug-design communities in using tree structures to accelerate similarity searches, especially in conjunction with low-dimensional molecular representations.(38)) These are very attractive features, however, they cannot be immediately transferred to drug-design applications for two main reasons. First, BIRCH was originally conceived for continuous inputs, which is in stark contrast with the discreet *on*/*off* character of the molecular fingerprint components. Moreover, the stored information in the CF limits BIRCH to the Euclidean distance, thus preventing the use of the Tanimoto similarity, or any other cheminformatic similarity index.

In this contribution we propose a novel clustering algorithm with the ease of interpretation of Taylor-Butina, but leveraging the advantages of the CF and CF-tree structures, resulting in more attractive memory and time scaling. Given the close relation with BIRCH, we termed our method: BitBIRCH. As discussed below, we can adapt the tree structure from the CF-tree without many changes (although BitBIRCH uses a more efficient criterion to split the tree nodes). The biggest challenge comes from the cluster representation, since we need an alternative to the CF that allows computing all the cluster’s properties from the collection of bitstrings. The solution to this comes from our work on quantifying the chemical diversity of large datasets by comparing an arbitrary number of molecules at the same time. We started applying this idea with extended similarity(39, 40) indices, which were successfully applied to dissecting epigenetic libraries(41), chemical space sampling(42), activity cliffs(43), and even in the study of Molecular Dynamics simulations.(44, 45) Recently, we expanded on this framework with the introduction of the instant similarity (iSIM) formalism.(46) iSIM shows how the average of the pairwise comparisons over a library can be calculated from two simple ingredients: the number of points in the set and a cumulative vector obtained after adding the fingerprints column wise. This is precisely what is needed to replace the CF by a Bit Feature (BF) that encapsulates the cluster’s information, and to be able to use the Tanimoto similarity in a BIRCH-like context. In the next sections, we discuss how iSIM can lead to a cluster’s centroid, radius, and diameter, and how to include information in the BF that allows to cluster a large set in a truly online way. Our estimates show that already for only 1,500,000 molecules BitBIRCH is > 1,000 times faster than the RDKit implementation of Taylor-Butina. We also compare these methods using several clustering quality metrics, which show that BitBIRCH’s improved time and memory efficiency do not diminish the final clustering results, with our algorithm outperforming Taylor-Butina in multiple instances. Finally, we discuss alternative formulations of BitBIRCH capable of handling billion-sized sets in just a few hours.

### BitBIRCH, Instant Similarity, and Taylor-Butina

BitBIRCH and BIRCH are based on two key features: a) the use of a reduced, vector-like, representation to encode the information about each cluster, b) a (CF-)tree structure to traverse and store the data. The latter is similar between both algorithms, with the most salient difference being the way to assign sub-clusters after splitting a node (details about this and the full pseudo-code are discussed in the SI, section S1). As far as the simplified cluster representation, BIRCH uses the CF, while for BitBIRCH we propose the Bit Feature (BF). Let 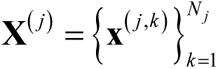 be the *j*^th^ cluster, containing *N*_j_ elements. Each of these elements, **x***^( j ,k )^* , is a *q*-dimensional vector, that is: 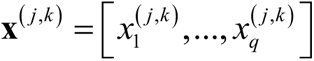. From now onwards, bold lowercase symbols will always represent vectors with *q* components/features. (In all the numerical results discussed below, unless otherwise explicitly mentioned, we worked with fingerprints with *q* = 2,048.) Then, the CF(20) of the *j*^th^ cluster, CF_j_, has three elements:

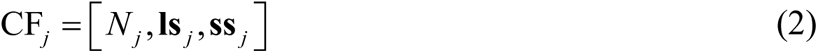

**ls**_j_ and **ss**_j_ are the linear sum and sum of squares, respectively, of the elements of each column of the matrix formed by the **x***^(j ,k)^*:

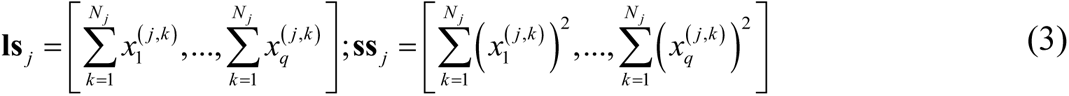

This is all that is needed to calculate either the radius or diameter of each cluster if the separation between the points is measured using the Euclidean distance. However, it is clear that for binary inputs (like molecular fingerprints) a new strategy is needed. First, we want to quantify similarity in different ways. Second, for binary data **ls**_j_ = **ss**_j_, meaning that they carry exactly the same information, and storing both vectors is unnecessary, independently of the chosen metric.

The BF_j_, on the other hand, is represented with a four-component structure:

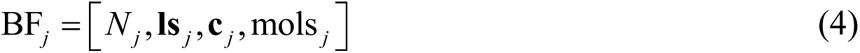

*N*_j_ and **ls**_j_ have the same meaning as in the CF_j_, and as we will show below, they are what is needed to calculate the radius/diameter of the clusters and perform the clustering. However, in order to add extra functionality, we decided to include two other elements in the cluster representation, mols_j_ and **c**_j_. First, mols_j_ is a list containing the indices of the molecules in the *j*^th^ cluster. Keeping track of the cluster assignments makes it possible to save the state of the clustering, while we expect to read new data. This is particularly attractive when all the molecules are not available at the same time, be it a matter of minutes/hours (e.g., after iterations of a generative pipeline) or even months/years (e.g., while expanding a library like ChEMBL(47) or ZINC(48)). In these cases, a BitBIRCH instance could be saved to disk, giving the possibility to update the clustering whenever needed.

The other new inclusion in the BF_j_ is the cluster centroid, **c**_j_. Since cluster membership is determined by comparing the new molecules being inserted in the tree with the cluster centroids, having the latter pre-calculated is more efficient than having to calculate them on-demand. Moreover, since we do not need the **ss**_j_, we can store the **c**_j_ at no extra memory cost, compared to the CF_j_. The centroid can be easily calculated from *N*_j_ and **ls**_j_ if we remember that, by definition, the center of a cluster is the point that, on average, is the closest to all the elements in the set. For instance, from the Tanimoto formula (Eq. (1)), we see that for every bit position, we should maximize the number of *on*-*on* coincidences, while minimizing the number of *on*-*off* mismatches. Thus, from the **ls**_j_ we can determine **c**_j_ as:

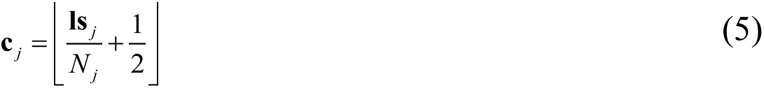

Where └*x*┘ is the floor function.

All that is left now is showing that *N*_j_ and **ls**_j_ are enough to calculate the diameter and radius of the cluster. The diameter of a cluster, *D*_j_, is defined as the average of all the pairwise inter-point separations:

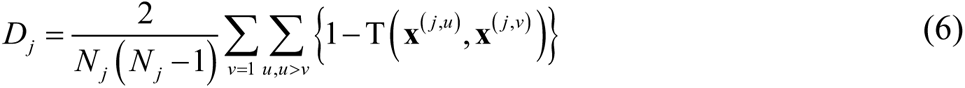

The key insight is that we can use the recently introduced instant similarity (iSIM)(46) to calculate the instant Tanimoto value of the set, *i*T _(_**X**^(^ *^j^*^)^ _)_, which is the average of the Tanimoto values over the cluster. In short, each element of **ls**_j_ contains information about the number of *on* bits in a column, so 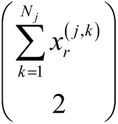 corresponds to all the possible *on*-*on* matches, while 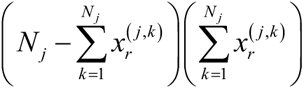 is the number of *on*-*off* mismatches in the *r*^th^ column, respectively. We can then write:

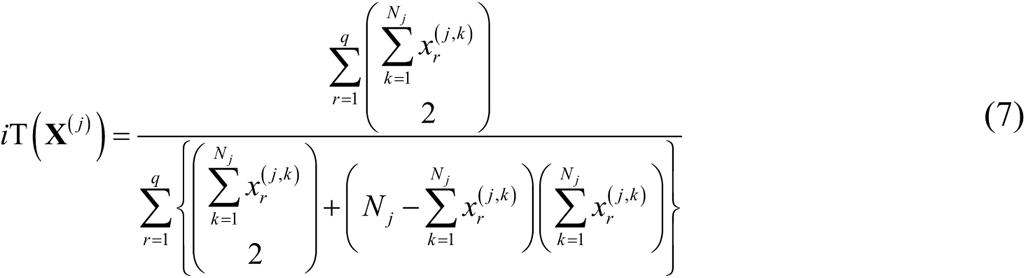

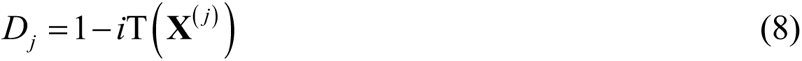

A similar argument can be made for the radius of the cluster, *R*_j_, defined as the average separation between all the points in the cluster and its centroid:

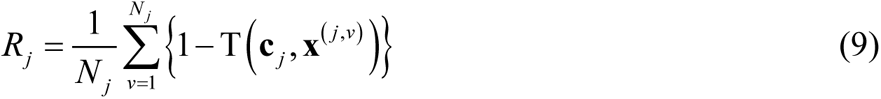

However, we can use iSIM to rewrite this as (see section S1 of the SI for a detailed derivation):

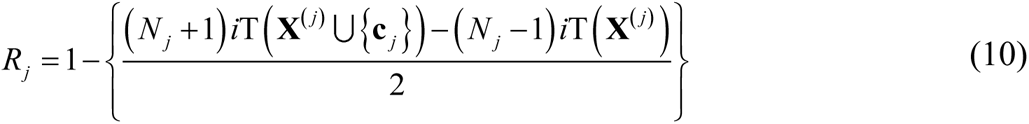

In other words, *D*_j_ and *R*_j_ can be calculated with just the number of elements and linear sum of a cluster thanks to iSIM. In the remaining of this contribution, we will discuss BitBIRCH based on *R*_j_, since this is the closest to the sphere-exclusion algorithm. Also, for simplicity, instead of referring to a radial threshold, we will be referring to a similarity threshold, calculated as 1− *R_j_*.

As shown in Fig. 1A, BitBIRCH retains the attractive O(*N*) scaling of BIRCH, due to the combination of the BF and tree structure, in stark contrast with the already-mentioned O(*N*^2^) time (also shown in Fig. 1A) and memory requirements of the Taylor-Butina algorithm. The Taylor-Butina implementation in RDKit(27) took more than 8 hours to cluster 450,000 molecules (each with 2,048 features) in the University of Florida’s HiPerGator cluster, and we were unable to allocate enough memory to cluster more than 500,000 molecules, since it required more than 4 TB of RAM. On the other hand, BitBIRCH was able to cluster 450,000 molecules in 2.2 minutes, and easily handled one million molecules in ∼ 5 minutes. Despite the great difference in performance, BitBIRCH and Taylor-Butina are in close agreement over the global structure of the data. Thus, the valuable insights and intuition built upon the sphere-exclusion method are also available through the BitBIRCH approach, but a much lower computational cost. In Fig. 1B we show how the number of clusters found by each method follows the same trends over diverse conditions. We considered clusters that had more than a minimum number of elements (min_size = 1, 2, 3, 5, and 10), in order focus on denser subsets. While the general agreement in the number of clusters is very good for all the considered libraries (see also SI, section S3), Taylor-Butina and BitBIRCH are particularly close when more singleton-like clusters are removed (min_size = 10). This global similarity is also present at the local level, if we compare set representatives from the top 10 most populated clusters found by both algorithms. Fig. 1C presents a heat-map with the Tanimoto comparisons between the medoids found by Taylor-Butina and BitBIRCH (for details on how to find the medoids, check the SI, section S1). Overall, there is a close relation between the “core” molecules identified by these methods, especially for the denser clusters. As discussed in the SI, this trend persists for the other ChEMBL subsets considered in this work. Moreover, in the SI we also discuss other tests showing the agreement between these two methods, including the Jaccard-Tanimoto set comparison between the most populated clusters, and the analysis of these trends over multiple coincidence thresholds.

**Figure 1:**
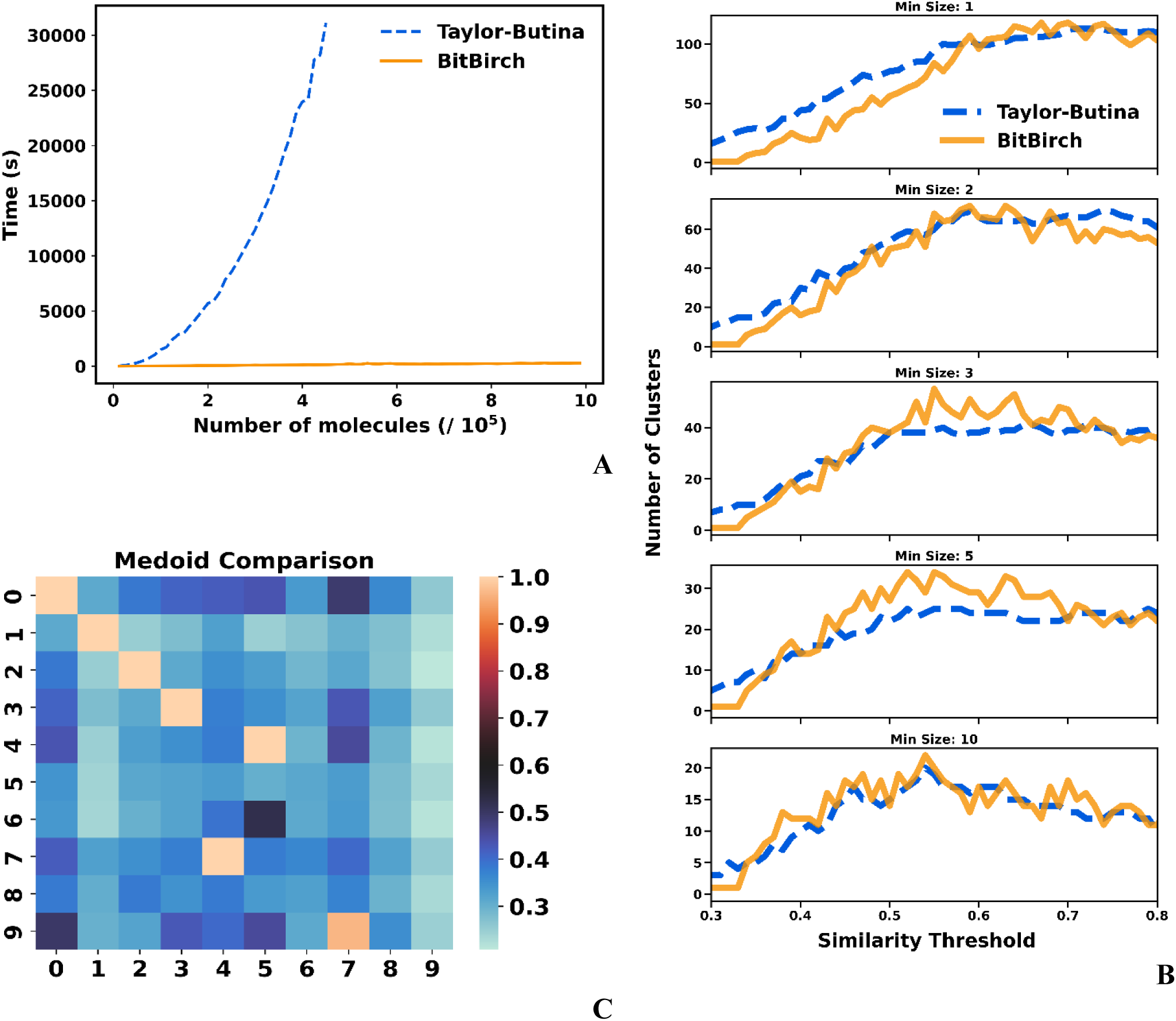
**A**: Time needed to cluster subsets of the Pubchem library up to one million molecules with the Taylor-Butina (RDKit implementation) and BitBIRCH algorithms; **B**, **C**: Analysis of the ChEMBL 262_Ki [19] library; **B**: Numbers of clusters with more than min_size (= 1, 2, 3, 5, 10) elements found by the Taylor-Butina (dashed blue line) and BitBIRCH (continuous orange line) algorithms for different similarity thresholds; **C**: Comparison of the medoids’ similarity of the top 10 most populated clusters (similarity threshold = 0.65, min_size = 10).

### Clustering Performance

Having established that the BitBIRCH and Taylor-Butina algorithms offer similar views on the nature of the data, we now compare the quality of the resulting clusterings. We used three main internal cluster validation measures(49): the Calinski-Harabasz (CHI)(50), Davies-Bouldin (DBI)(51), and Dunn (DI)(52) indices. The CHI depends on the ratio of between- and within-cluster dispersions, with bigger values indicating a better clustering.(50) This index can be re-formulated with Tanimoto as a similarity measure:

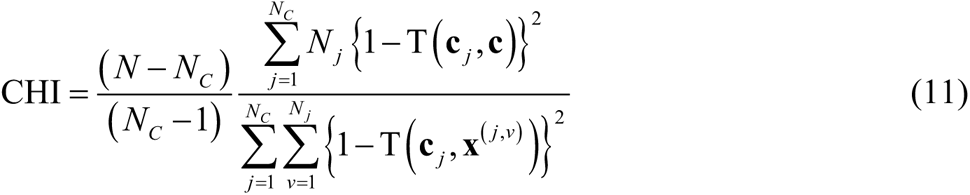

where *N* is the total number of molecules, *N*_C_ is the number of clusters, and **c** is the centroid of all the data. It is important to note that, with the help of iSIM, CHI can be estimated in O(*N*). DBI values are usually easier to interpret, since they are bounded in the [0, 1] interval, with lower values corresponding to tighter and more separated clusters.(51) However, calculating this index demands computing all the pairwise similarities between the cluster representatives. This is problematic, since for most practical values of the similarity threshold the number of clusters is proportional to the number of molecules (*N*_C_ ∼ O(*N*)), or in other words, the DBI calculation will roughly scale as O(*N*^2^). Even if we include the DBI analysis as a way to benchmark the performance of BitBIRCH, this index is not a viable option for ultra-large libraries; calculating it is more demanding than actually performing the clustering. The DI also suffers from this problem, since it demands calculating the separation between all pairs of clusters. Still, we report DI values for all the 30 ChEMBL subsets here and in the SI, keeping in mind that higher DI values indicate more well-separated clusters (more details about the expressions used to calculate the DBI and DI are included in the SI, section S4).

Figure 2A-C shows the mentioned quality clustering metrics for the ChEMBL 233 library. The CHI values in Fig. 2A indicate that BitBIRCH outperforms Taylor-Butina for lower similarity thresholds (< 0.6), with both methods giving clusters of the same quality until slightly over the 0.7 threshold, a range that covers most practical applications. This observation is supported when a Wilcoxon signed-rank test is done, the CHIs for the thirty ChEMBL libraries are significantly higher for BitBIRCH at low thresholds (*p* < 0.05) and there is no statistically significant difference at thresholds in the [0.5, 0.7] range, depending on the minimum size of the considered clusters (see Figs. S4.31 and S4.32 in the SI). The shaded areas in Figs. 2A-C indicate the standard deviation of the corresponding index values calculated after removing clusters with more than 1, 2, 3, 5, and 10 molecules. Interestingly, the BitBIRCH results are consistently more robust to the removal of outliers and singleton-like clusters than those of Taylor-Butina, which is once again more prevalent for looser thresholds. The DBI analysis (Fig. 2B) is similar to the CHI, now with a region of BitBIRCH that gives markedly better results until a 0.65 threshold, and then essentially equivalent results to Taylor-Butina around the 0.75 region. This observation for ChEMBL_233 also applies to all the 30 studied libraries, with Wilcoxon tests showing that there is no statistical difference at threshold values in [0.5, 0.7], while at low thresholds, the DBI score is significantly lower for BitBIRCH (see section S4 in the SI). The DI results are even more promising (Fig. 2C), with BitBIRCH consistently outperforming Taylor-Butina almost up to a 0.8 threshold. Overall, the DI is statistically higher for BitBIRCH in the [0.3, 0.8] range (SI, section S4). It is reassuring to see that these different metrics agree on the relative performance of both algorithms, with the CHI and DI plots, in particular, presenting very similar features. Essentially, for the range of similarity thresholds that are usually explored in drug design(53) and ML applications, BitBIRCH performs better or, at worst, equal to Taylor-Butina.

**Figure 2:**
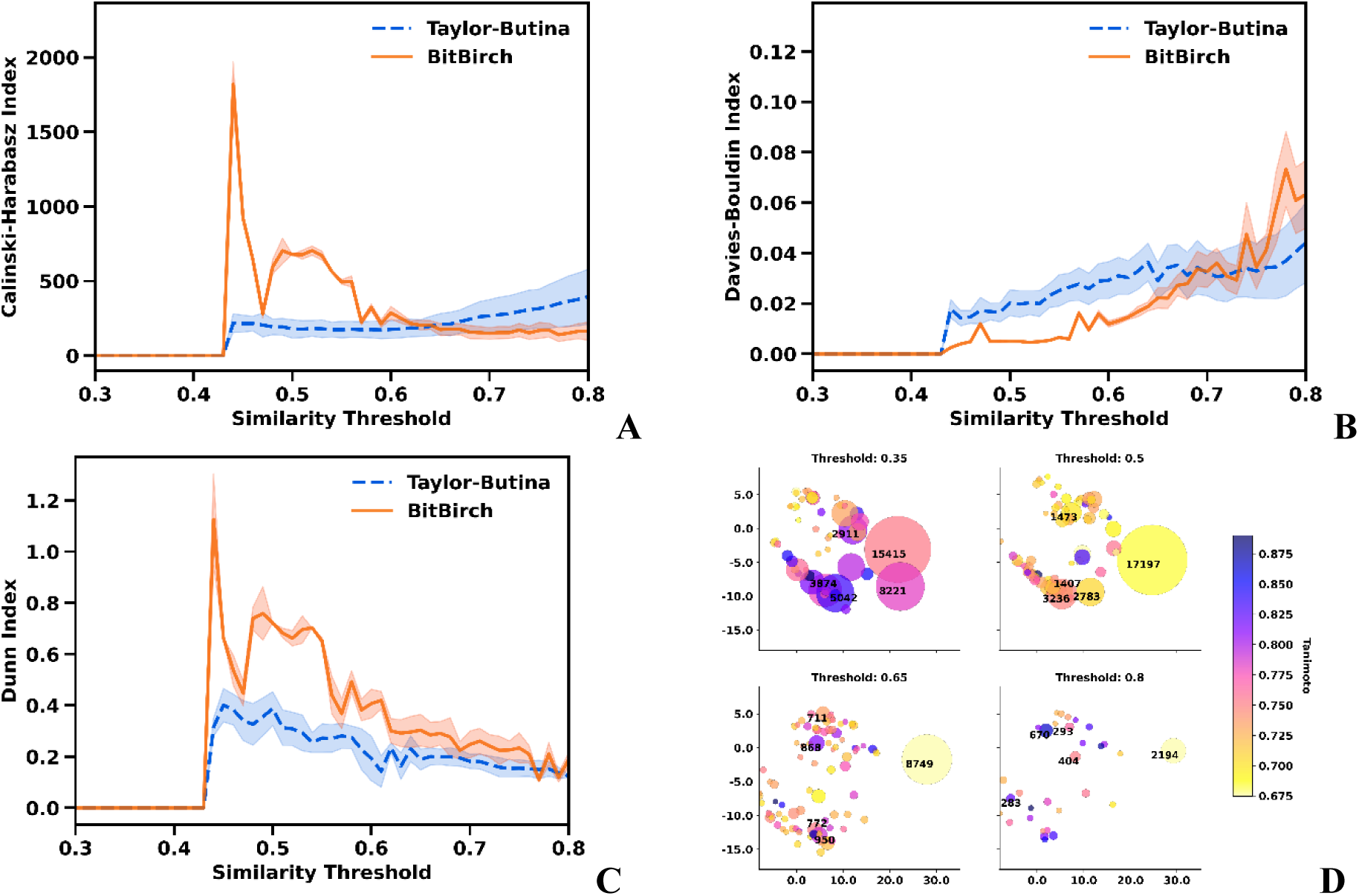
Comparison of Taylor-Butina (blue dashed line) and BitBIRCH (orange continuous line) clustering results for the ChEMBL 233_Ki [9] library. Dark lines and shaded regions indicate the average and standard deviation over min_size = 1, 2, 3, 5, 10 values, respectively. **A**: Calinski-Harabasz, **B**: Davies-Bouldin, and **C**: Dunn indices. **D**: Projection into the first two principal components of the BitBIRCH clustering of the ChEMBL33 natural products library with similarity thresholds = 0.35, 0.5, 0.65, 0.8. Colors indicate the iT values for the clusters, and the populations of the five largest clusters are explicitly indicated.

As a final test on the potential variability of BitBIRCH, we studied how the distribution of the clusters in chemical space changes under different conditions. (For this, we used a PCA projection into the first two principal components to conveniently visualize the cluster results.) For the natural products of the ChEMBL33(47) library (Fig. 2D) we observe that from a relatively loose threshold (0.35) to a much stricter one (0.8), the relative positions of the most populated clusters remain sensibly constant. This is especially clear for the 0.5-0.8 range, where not only the position of the clusters, but also the average similarities of the cluster members (calculated with iSIM, see color scale in Fig. 2D) is largely preserved. This indicates that BitBIRCH results are quite robust to changes in the similarity threshold. At least at a general level, we do not have to carefully fine-tune this parameter, especially within the [0.5, 0.8] interval.

### Tackling Ultra-Large Libraries

As shown above, the standard BitBIRCH implementation can easily handle millions of molecules, but now we explore strategies to handle much larger sets. We consider two possibilities: fingerprint folding and an iterative/parallel scheme.

Fingerprint folding is a well-known “trick” in the cheminformatics/similarity-search community.(54–56) A fold uses a pre-specified rule to combine the *on* and *off* bits of the two halves of a fingerprint, leading to a new binary representation with half the size of the original input. This procedure can be repeated, resulting in *n*-folded fingerprints, which directly translates to lower runtime and memory costs.(55, 56) Initially, it seems like a drastic simplification, however, it is well-documented that folding (up to 4-folding) still tends to preserve the ranking between the comparisons and to retrieve the same neighbors (particularly, for similarity thresholds > 0.5).(57)

Here, we explored one-, two-, and three-folding taking the 2048-length fingerprints and converting them to 1024-, 512-, and 256-bits vectors using a XOR logic operation, respectively.(54) First, we notice that the new methods follow the same trends in the number of clusters as the BitBIRCH algorithm (Fig. 3A). As expected, this agreement is better for the 1- and 2-folded fingerprints. However, the 3-folded approximation has significantly higher CHI values overall (Fig. 3B, see SI S6). The DBI for the folded fingerprints (Fig. 3C) shows a similar behavior as in the BitBIRCH-parallel case, with no single method being the clear-cut best option for all the studied libraries. In contrast with the DI (Fig. 3D), the original BitBIRCH outperforms all the folded clusters. With a Wilcoxon one-sided test, we confirm that the original algorithm has significantly lower DBIs and significantly higher DIs (SI, section S6), which suggests that folding the fingerprints can hurt the final clustering quality.

**Figure 3:**
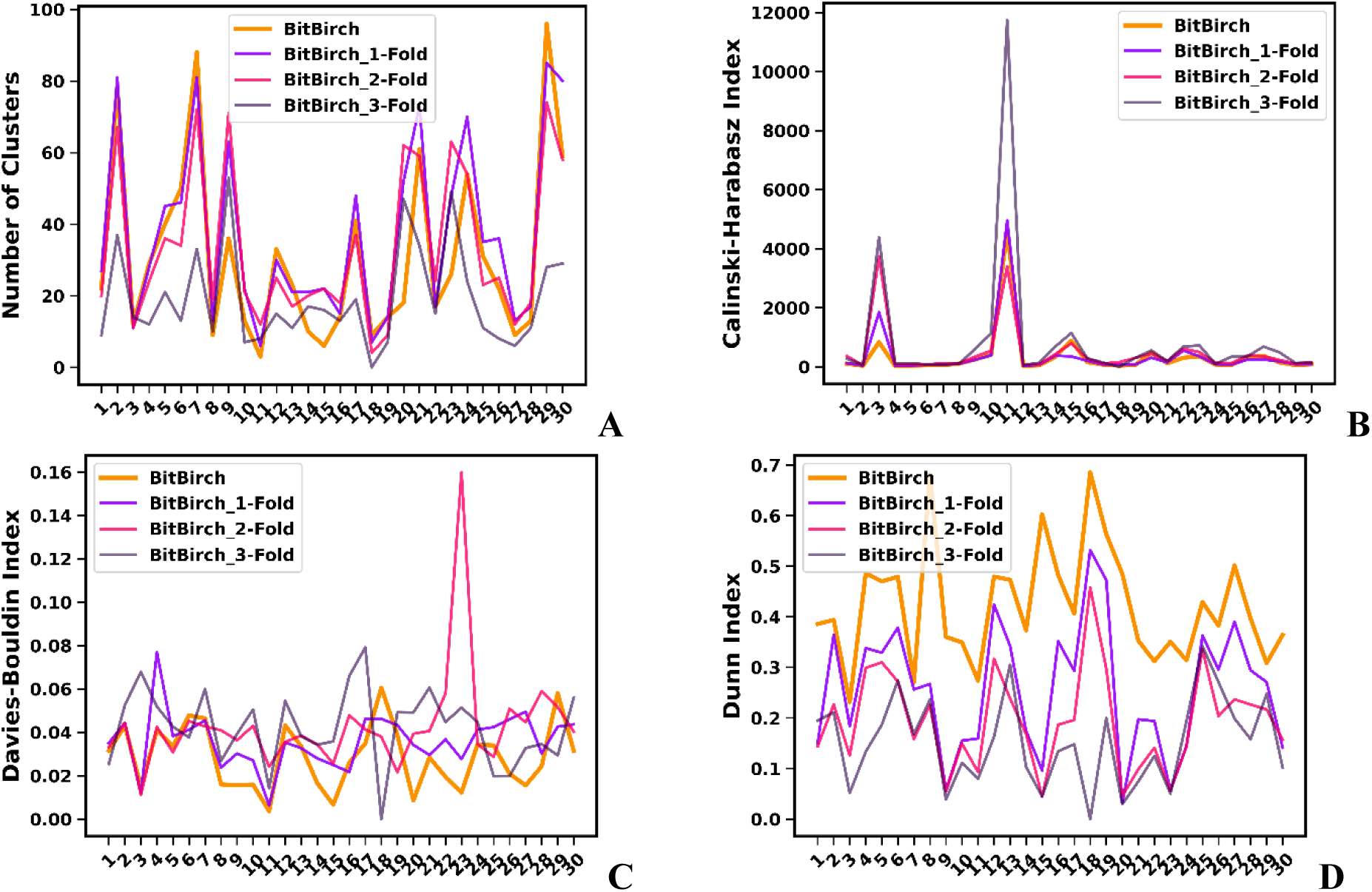
Comparison of the performance of the original (continuous orange line) and folded (dashed green line) BitBIRCH algorithms over 30 ChEMBL subsets. **A**: Number of clusters; **B**: Calinski-Harabasz, **C**: Davies-Bouldin, and **D**: Dunn indices.

The parallel method is inspired by the (optional) refinement step at the end of traditional BIRCH implementations(37) and clustering ensembles.(58) The basic idea is that the cluster centroids could be used as representatives of their corresponding sets. Centroids are used as input in a second clustering step. So, the BitBIRCH-parallel recipe is quite simple: 1-Separate the data into non-overlapping splits (we usually consider 10 splits, as discussed below); 2-Use BitBIRCH to cluster each of these splits; 3-Collect the cluster centroids and cluster them with BitBIRCH. This parallel/iterative approach is an approximation to just clustering all the data in one go, but the question remains: how much does the quality of the clustering suffer after this iterative procedure? First, Fig. 4A shows that for the 30 ChEMBL subsets considered here, the final number of clusters found by the standard and the parallel BitBIRCH methods (with 10 initial splits) is quite consistent, which reflects the ability of the simpler method to capture the same global structure as the exact BitBIRCH. In the SI we also present a detailed comparison between the local structure of the clusters found by these algorithms (SI, section S5). The CHI (Fig. 4B) and DBI (Fig. 4C) analysis reveal a promising pattern: in general, both methods have very similar performance, with the parallel approach being even slightly favored in some cases, as reported in other ensemble-like approaches.(59, 60) Even more reassuring, according to the DI (Fig. 4D), in most cases BitBIRCH-parallel finds better-separated and more compact clusters. The statistical analysis supports these observations, with the corresponding CHI and DBI values being statistically equivalent (*p* = 0.16 and *p* = 0.45, respectively), while the parallel DIs are significantly higher than BitBIRCH (*p* < 0.05) in the respective Wilcoxon’s tests. (Details on SI, S5)

**Figure 4:**
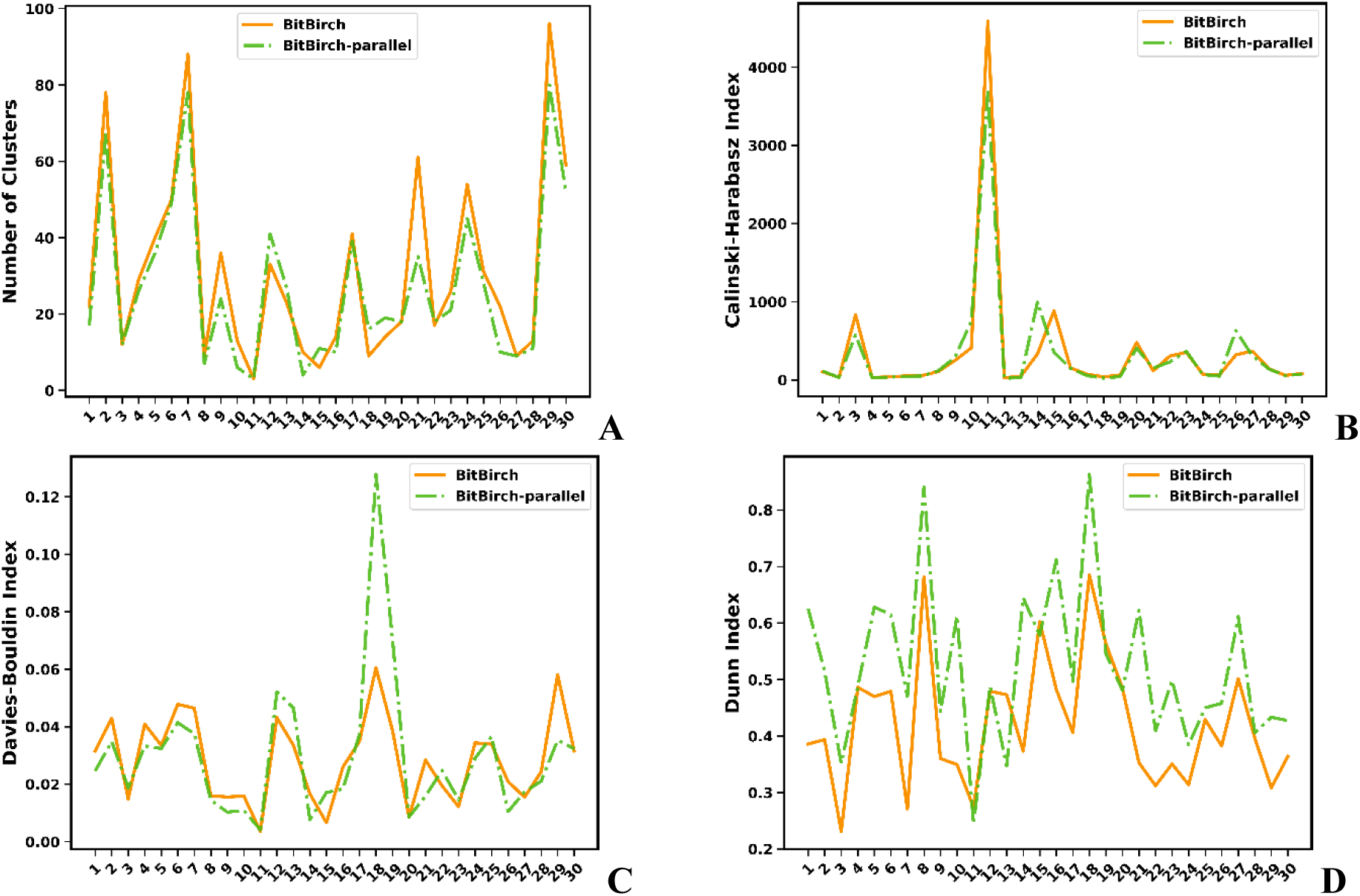
Comparison of the performance of the original (continuous orange line) and parallel (dashed green line) BitBIRCH algorithms (with 10 initial splits) over the 30 ChEMBL subsets. **A**: Number of clusters; **B**: Calinski-Harabasz, **C**: Davies-Bouldin, and **D**: Dunn indices.

Since by all the considered metrics the iterative/parallel approximation does not diminish the quality of the final results, we chose it to analyze a library with one billion molecules. As noted above, 4 TB were insufficient to cluster 450,000 molecules with Taylor-Butina. But even ignoring the memory issues, the trend in Fig. 1A ( *O* (Taylor-Butina) = 2*10^−7^ *N* ^2^ ) suggests that it will take ∼ 6,342 years to cluster one billion molecules. Using the BitBIRCH algorithm without any modifications ( *O* (BitBIRCH) = 2.94*10^−4^ *N* ) this estimate improves to only 3.4 days, which is already a practical time, but we wanted to test how much one could improve upon this result with the parallel BitBIRCH implementation. The one billion molecules were selected from the ZINC22(48) library and clustered using the parallel strategy using 1,000 splits (1,000,000 molecules each). While the average time to cluster a split was 4.34 min, to compute the total time we will consider the slowest one, 18.76 min, since it is the limiting step before clustering the centroids. On average, each split clustering yielded 26,653 centroids (lowest: 5,430; highest: 84,649). Then, the final clustering took 242 min between the centroid aggregation and clustering, resulting in a total of 2,081,662 clusters with more than 10 molecules. The total time to obtain the final clustering assignations of the billion molecules was 4.35 hours; a much more attractive time than the unmanageable Taylor-Butina estimate, and even the days required for this task using the unmodified BitBIRCH. In Fig. 5A we present a summary of the cluster population distribution for the 10,000 largest clusters, indicating a strong preference for smaller “pockets” of molecules. The distribution of the larger clusters (Fig. 5B) shows a relatively uneven location of the more populated centers (darker and bigger circles in Fig. 5B), which dominate the boundaries of the two-dimensional PCA projection. The asymmetric distribution of the denser clusters is also showcased by the “voids” observed in the central region of this chemical sub-space.

**Figure 5:**
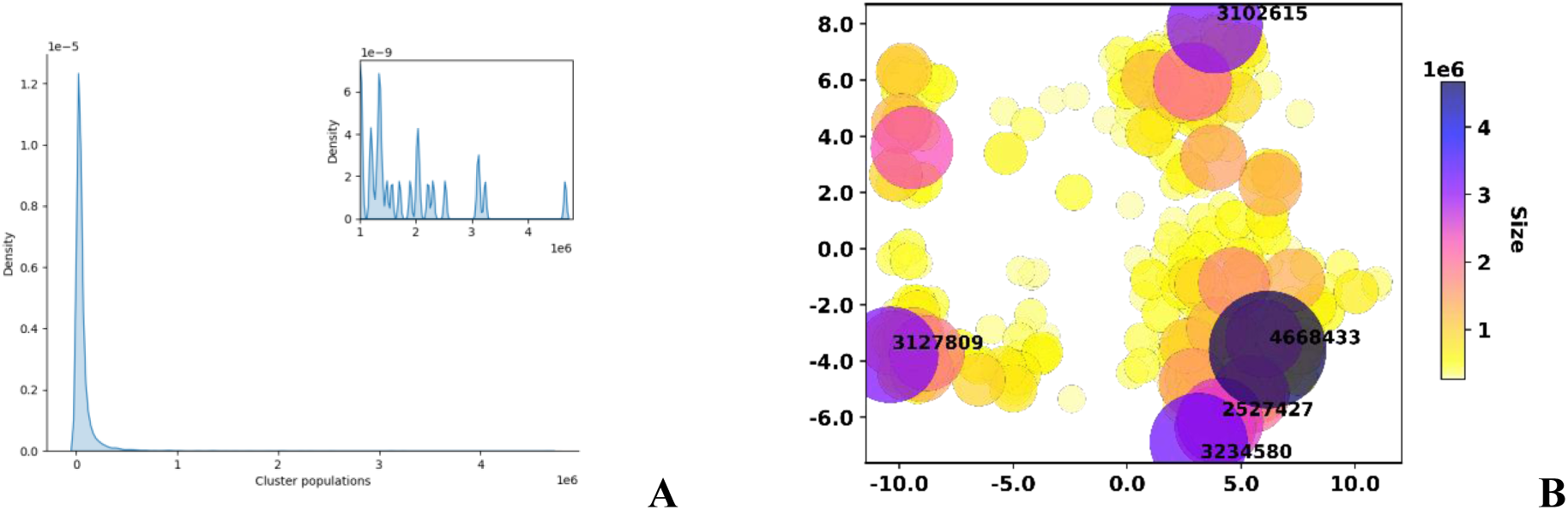
**A**: Kernel density estimation of the populations of the top 10,000 most populated clusters from 1 billion molecules of ZINC22. The zoomed image has the details of the density estimation for the clusters with > 1,000,000 molecules. **B**: Projection into the first two principal components of the 378 clusters with at least one million molecules. Circle color and size correspond to the number of molecules in each cluster. The populations of the top five largest clusters are explicitly indicated.

## Conclusions

We have developed an alternative to the BIRCH and Taylor-Butina algorithms, BitBIRCH, capable of handling binary data with the similarity metrics that are used in drug-design and ML applications targeting small molecules. Key components of BitBIRCH include the tree structure derived from the BIRCH method, as well as the ability to use the BF to encode the cluster’s information. The tree is critical to ensure the O(*N*) time scaling, by reducing the total number of comparisons required to allocate each new molecule to their corresponding cluster. The BF, on the other hand, is what allows to reduce the memory usage, by providing a convenient proxy to the cluster’s properties required to assign new molecules or create new sub-divisions. This is possible thanks to the iSIM formalism since having the chance to calculate the average of the pairwise Tanimoto comparisons is what opens the door to the cluster’s radius and diameter. As noted before, while we focused on Tanimoto similarity, iSIM can be used to calculate the average pairwise with other similarity indices; BitBIRCH can trivially incorporate other criteria, like Russel-Rao and Sokal-Michener. Likewise, this methodology is applicable to any type of fingerprints.

The comparison of BitBIRCH with the RDKit implementation of Taylor-Butina showcases the differences in efficiency between these methods. We saw that 4 TB of memory were insufficient for Taylor-Butina to cluster more than 450,000 molecules, while BitBIRCH was able to easily handle millions. Moreover, RDKit took more than 8 hours to cluster those 450,000 molecules, with BitBIRCH requiring ∼ 2.2 minutes to complete the same task. If we follow the trends in Fig. 1A, we see that for 1,500,000 molecules, BitBIRCH is > 1,000 times faster than Taylor-Butina. Compare this to other linear clustering algorithms, like linclust, that only got to a 1,000X speedup over their previous alternatives for billions of points, thus showcasing how much faster BitBIRCH is compared to Taylor-Butina. One of the key characteristics of BitBIRCH is that the increase in time and memory efficiency do not diminish the clustering quality compared to the O(*N*^2^) alternatives. First, we saw that BitBIRCH and Taylor-Butina partition the chemical space in similar ways (total number of clusters, etc.). Moreover, a more detailed analysis over 30 subsets of the ChEMBL library showed that BitBIRCH outperforms Taylor-Butina for lower similarity thresholds according to the CHI and DBI metrics, with both methods being statistically indistinguishable in the [0.5, 0.7] range. The DI, on the other hand, favors BitBIRCH up to a 0.8 threshold. So, for all the clustering metrics considered, BitBIRCH performs better or, at worst, with the same quality as Taylor-Butina. BitBIRCH also proved to be more robust to changes in the computational conditions (similarity threshold, minimum number of molecules in a cluster).

Finally, we explored alternatives to tackle billion-sized libraries in even more efficient ways. The fingerprint folding approach compromised the clustering results, with even one-fold, leading to worse DI values than the original BitBIRCH method. We found a more promising route with the parallel/iterative clustering strategy, it separates the data into different splits, which are then clustered and whose resulting centroids are then clustered again. This led to performances remarkably close to the unmodified BitBIRCH algorithm, while considerably boosting the efficiency of the method. We tested this with 1 billion molecules from the ZINC22 library, which we were able to cluster in 4.35 hours (in stark contrast with the estimated 6,342 years it would take the RDKit Taylor-Butina implementation to complete this task).

## Materials and Methods

The timing comparisons between Taylor-Butina and BitBirch were done on the first 1 million molecules from the PubChem 2005 library. For clustering performance calculations, 30 ChEMBL target-oriented subsets curated by van Tilborg et al were used (details on SI, section S2)(61). For the PCA clustering visualization changes with similarity threshold, the natural products from ChEMBL33(47) (n = 64,087) were used. For the billion molecules clustering, random tranches from the ZINC22 2D(48) database (https://cartblanche.docking.org/tranches/2d) were taken until 1 billion was reached. (see SI, section S7). In all cases, 2048-bit RDKit(27) fingerprints were generated from SMILES, corrupted SMILES were discarded from the databases. The statistical comparison between the clustering performance metrics for the methods was done using SciPy’s(62) Wilcoxon test(63), alternative hypothesis evaluated depended on the case. All the calculations (except the statistical analysis and plotting) were run in University of Florida’s HiperGator supercomputer. The BitBIRCH code is available at: https://github.com/mqcomplab/bitbirch

## Acknowledgements

KLP, LC, KH, and RAMQ thank the National Institute of General Medical Sciences of the National Institutes of Health for support under award number R35GM150620. VJ thanks the UF AI Scholars program for a fellowship.

