## Supplementary Information for "Efficient clustering of large molecular libraries"

### S1: BitBIRCH Algorithm and Complementary Similarity

Here we present the pseudo-code for the key parts of the BitBIRCH algorithm. Underlined sections indicate some of the key differences with respect to the traditional BIRCH method.

```
BitBIRCH(library)
  initialize root
    for molecule in library:
      subcluster = create_subcluster(1, molecule, molecule, molecule_index)
      split = root.insert_bf_subcluster(root, subcluster)

      if split
        split_node(root)
        update root
```

**Figure S1.1**: Main BitBIRCH algorithm.

```
create_subcluster(nmols, linear_sum, centroid, indices)
  subcluster.number_of_molecules = nmols
  subcluster.linear_sum = linear_sum
  subcluster.centroid = centroid
  subcluster.indices = indices
```

**Figure S1.2**: Create subcluster method.

```
split_node(node)
    initialize subcluster1 and subcluster2
    initialize new_node1 and new_node2

    subcluster1.child = new_node1
    subcluster2.child = new_node2

    extreme1, extreme2 = find_separated_molecules(node.centroids)

    for subcluster in node.subclusters
        if similarity(subcluster.centroid, extreme1) < similarity(subcluster.centroid, extreme2)
            append subcluster to new_node1
        else
            append subcluster to new_node2
```

**Figure S1.3**: Split node algorithm.

In the calculations discussed here and in the main text, the **similarity** function used was the pairwise Tanimoto index (Eq. (1) in the manuscript).

```
insert_bf_subcluster(node, subcluster)
  ind = argmax(similarity(node.centroids, subcluster.centroid))
  closest_sub = node.subclusters[ind]

  if closest_sub.child not NULL
     split = closest_sub.child.insert_bf_subcluster(node, subcluster)
     if not split
        merge(closest_sub, subcluster)
        return False
     else
        split_node(closest_sub.child)
        if number_of_subclusters > branching_factor
           return True
        return False

  else
     if jt_radius(closest_sub, subcluster) < threshold
        merge(closest_sub, subcluster)
        return False
     else if  length(node.subclusters) < branching factor
        return False
     else
        return True
```

**Figure S1.4**: Insert subcluster algorithm.

In all the calculations, we used a default value of 50 for the branching_factor.

```
merge(cluster1, cluster2)
   subcluster.number_of_molecules = cluster1.number_of_molecules +
       cluster2.number_of_molecules
   subcluster.linear_sum = cluster1.linear_sum + cluster2.linear_sum
   subcluster.indices = cluster1.indices + cluster2.indices
```

**Figure S1.5**: Merge clusters algorithm.

An important feature of the iSIM formalism, is the ability to easily rank the molecules in a set depending on how central-like or outlier-like they are. This is done through the concept of complementary similarity which, for a given molecule, is just the iSIM value of the set after that molecule has been removed. It is clear that when we remove outliers, the iSIM of the remaining molecules will increase. Likewise, when we remove a molecule that is central to the set, the final iSIM will decrease. We use this insight to identify the medoid of the set, as the molecule with the lowest complementary similarity value, which can clearly be done in O($N$).

As mentioned in the manuscript, the medoid can serve as a representative of the cluster (with the key difference between medoid and centroid being that the former is required to be an element of the set, while the former does not have to be a real molecule). However, in BitBIRCH we also used the medoid in a new way to speed up the **max_separation** function. In BIRCH, if one tries to insert a subcluster in a node that already has branching_factor subclusters, the node must be split. This is done by taking the branching_factor centroids, finding the two most separated ones, and then assigning the remaining centroids to whichever of these two maximally separated points they are closest to. This demands calculating all the pairwise distances between the centroids, which scales as O(branching_factor$^2$). In BitBIRCH, we take a different route, in that we do not aim to find the absolutely most separated centroids, but we only need to find two centroids that are guaranteed to be very separated, compared to the other centroids in the node. The recipe for this is very simple: 1- Find the medoid among the branching_factor centroids, 2- Find the centroid that is furthest away from the medoid (label this as molecule_1), 3- Find the molecule that is the furthest away from molecule_1 (label this as molecule_2). Then, molecule_1 and molecule_2 will serve as the two pivot points around which the node will be partitioned. This procedure scales as O(branching_factor).

Finally, for a detailed derivation of Eq. (10) in the main text:

By definition:

$$\begin{aligned} R_j &= \frac{1}{N_j}\sum_{v=1}^{N_j}\left\{1-\mathrm{T}\left(\mathbf{c}_j,\mathbf{x}^{(j,v)}\right)\right\} \\ &= 1-\frac{1}{N_j}\sum_{v=1}^{N_j}\mathrm{T}\left(\mathbf{c}_j,\mathbf{x}^{(j,v)}\right) \end{aligned} \tag{1}$$

$$iT\left(\mathbf{X}^{(j)}\right) = \frac{\sum_{v=1}\sum_{u,u>v} T\left(\mathbf{x}^{(j,u)}, \mathbf{x}^{(j,v)}\right)}{\binom{N_j}{2}}$$

$$iT\left(\mathbf{X}^{(j)} \cup \{\mathbf{c}_j\}\right) = \frac{\sum_{v=1}\sum_{u>v} T\left(\mathbf{x}^{(j,u)}, \mathbf{x}^{(j,v)}\right) + \sum_{v=1}^{N_j} T\left(\mathbf{c}_j, \mathbf{x}^{(j,v)}\right)}{\binom{N_j+1}{2}} \tag{2}$$

$$\begin{aligned} R_j &= 1 - \frac{\binom{N_j+1}{2} iT\left(\mathbf{X}^{(j)} \cup \{\mathbf{c}_j\}\right) - \binom{N_j}{2} iT\left(\mathbf{X}^{(j)}\right)}{N_j} \\ &= 1 - \left\{ \frac{\left(N_j+1\right) iT\left(\mathbf{X}^{(j)} \cup \{\mathbf{c}_j\}\right) - \left(N_j - 1\right) iT\left(\mathbf{X}^{(j)}\right)}{2} \right\} \end{aligned} \tag{3}$$

**S2: Details of the ChEMBL Subsets**

The 30 ChEMBL subsets can be found here: https://github.com/molML/MoleculeACE/tree/main/MoleculeACE/Data/benchmark_data/old.

| Library name | Number of molecules | Code |
|---|---|---|
| CHEMBL218_EC50 | 1031 | [1] |
| CHEMBL264_Ki | 2862 | [2] |
| CHEMBL2971_Ki | 976 | [3] |
| CHEMBL238_Ki | 1052 | [4] |
| CHEMBL228_Ki | 1704 | [5] |
| CHEMBL219_Ki | 1859 | [6] |
| CHEMBL214_Ki | 3317 | [7] |
| CHEMBL2047_EC50 | 631 | [8] |
| CHEMBL233_Ki | 3142 | [9] |
| CHEMBL4005_Ki | 960 | [10] |
| CHEMBL2835_Ki | 615 | [11] |
| CHEMBL287_Ki | 1328 | [12] |
| CHEMBL231_Ki | 973 | [13] |
| CHEMBL2034_Ki | 750 | [14] |
| CHEMBL237_EC50 | 955 | [15] |
| CHEMBL4616_EC50 | 682 | [16] |
| CHEMBL239_EC50 | 1721 | [17] |
| CHEMBL4203_Ki | 731 | [18] |
| CHEMBL262_Ki | 856 | [19] |
| CHEMBL236_Ki | 2598 | [20] |

| | | |
|---|---|---|
| CHEMBL244_Ki | 3097 | [21] |
| CHEMBL2147_Ki | 1456 | [22] |
| CHEMBL237_Ki | 2602 | [23] |
| CHEMBL235_EC50 | 2349 | [24] |
| CHEMBL3979_EC50 | 1125 | [25] |
| CHEMBL4792_Ki | 1471 | [26] |
| CHEMBL1862_Ki | 794 | [27] |
| CHEMBL1871_Ki | 659 | [28] |
| CHEMBL234_Ki | 3657 | [29] |
| CHEMBL204_Ki | 2754 | [30] |

**Table S2.1**: Library name, number of molecules, and numerical code for the 30 ChEMBL subsets.

### S3: Local Clustering Analysis of BitBIRCH and Taylor-Butina

In the main text we presented a type of analysis aimed to understand the relation between the local structure of the clusters obtained with BitBIRCH and with Taylor-Butina: comparing the medoids of the most populated clusters. Here we present the results of a similar analysis, but that takes into account all the elements of the top clusters and not just a single representative. To perform this comparison, we use the Jaccard-Tanimoto, JT, set similarity index. That is, for two sets A and B, we have:

$$\mathrm{JT}\left(A,B\right)=\frac{\left|A\cap B\right|}{\left|A\cup B\right|} \tag{4}$$

In this section we show the Medoid comparison and the Set comparison for all the 30 ChEMBL subsets. In general, we see that even in the cases where the BitBIRCH and Taylor-Butina final clusters do not have a perfect agreement, they still manage to largely identify the same regions in the denser regions of chemical space.

We also complement these analyses with another metric, in order to make it more quantitative. Notice that, in an ideal case, both the Medoid and the Set results would give an identity matrix (e.g., every BitBIRCH cluster perfectly matching every Taylor-Butina counterpart). To measure this, we introduce a Score, *s*, that measures how similar are the Medoid and Set results to the identity matrix of the same rank. In short, for a matrix *M*, *s* is given by:

$$s(M)=\frac{1}{2}\left\{n(M)+n\left(M^{T}\right)\right\}$$

$$n(M)=\frac{\sum_{\substack{r=0\\ r\rightarrow rows}}^{\dim(M)-1}\left(1-r\frac{0.9}{\dim(M)-1}\right)\begin{cases}\max\left(M[r]\right)+1, \text{ if } r=\arg\max M[r]\\ \max\left(M[r]\right)+1-\left|\arg\max M[r]-r\right|*\frac{0.9}{\dim(M)-1-r}, \text{if } r\leq\frac{\dim(M)-1}{2}\\ \max\left(M[r]\right)+1-\left|\arg\max M[r]-r\right|*\frac{0.9}{r}, \text{if } r>\frac{\dim(M)-1}{2}\end{cases}}{1.1*\dim(M)}$$

(5)

*s* is bounded in the [0, 1] interval, with a value of 1 indicating perfect agreement with the identity matrix, so higher scores indicate a better agreement between Taylor-Butina and BitBIRCH. As shown below, both the Medoid and Set scores are very robust with respect to changes in the similarity threshold, which showcases the stability of the BitBIRCH results.

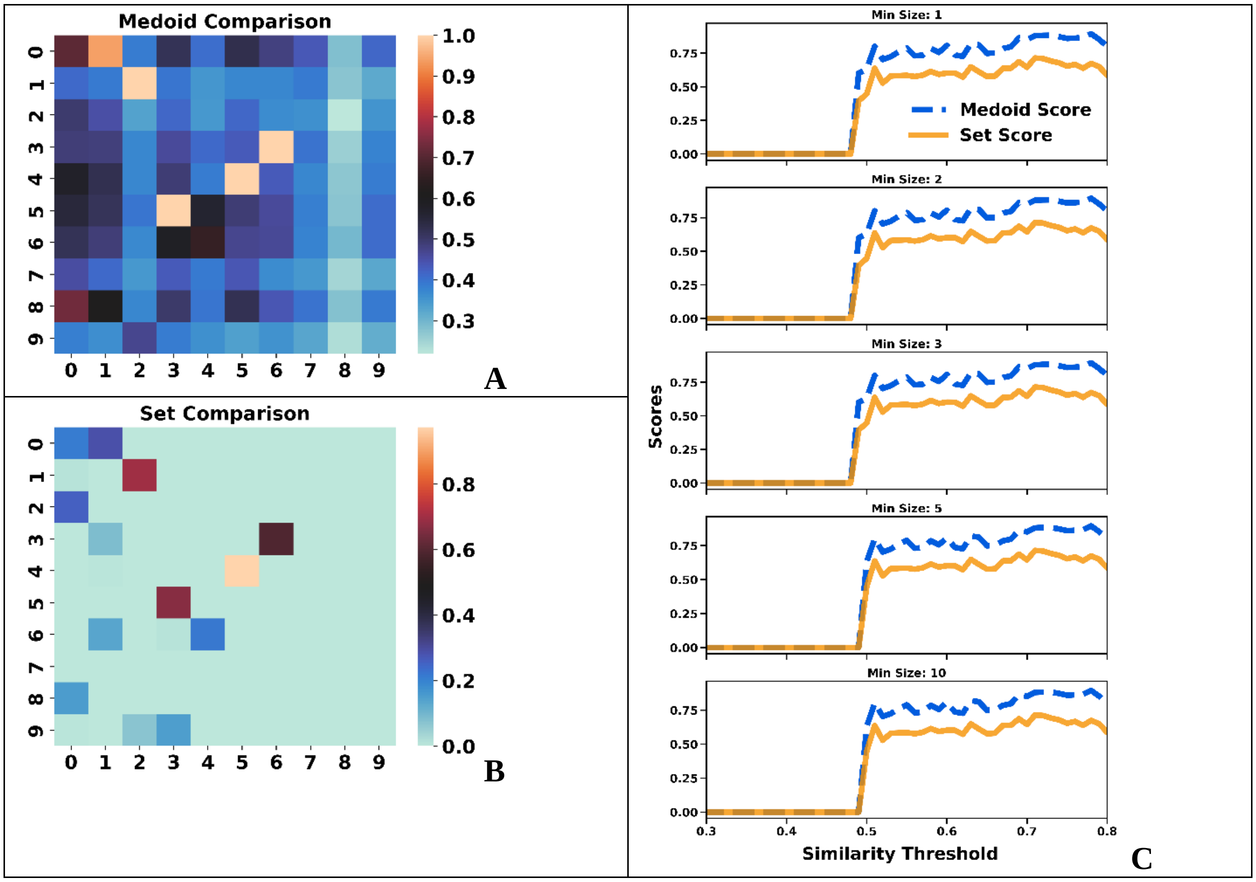

**Figure S3.1**: Comparison of the **A**: medoids, **B**: sets for the top populated clusters of the [1] ChEMBL subset (similarity threshold = 0.65, min_size = 10). **C**: Medoid and Set scores.

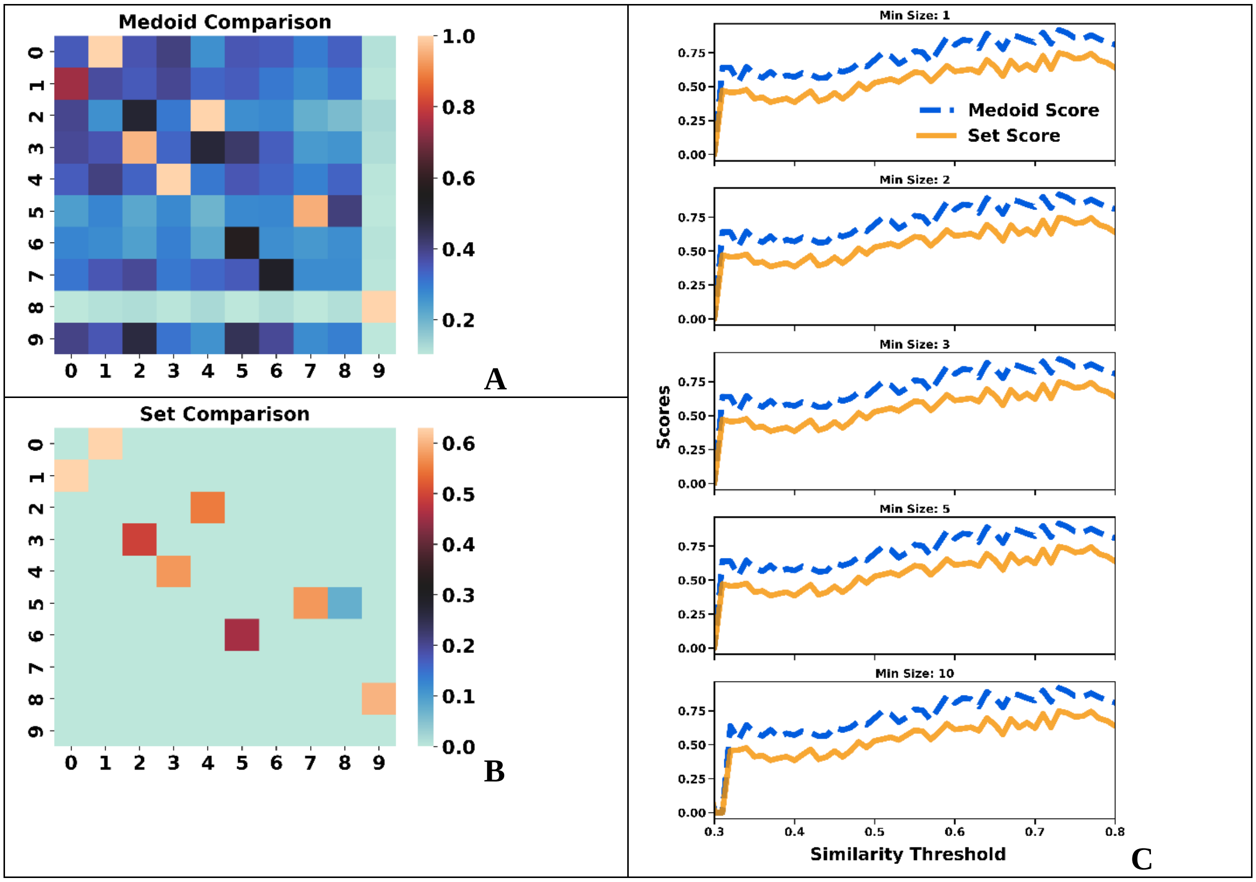


**Figure S3.2**: Comparison of the **A**: medoids, **B**: sets for the top populated clusters of the [2] ChEMBL subset (similarity threshold = 0.65, min_size = 10). **C**: Medoid and Set scores.

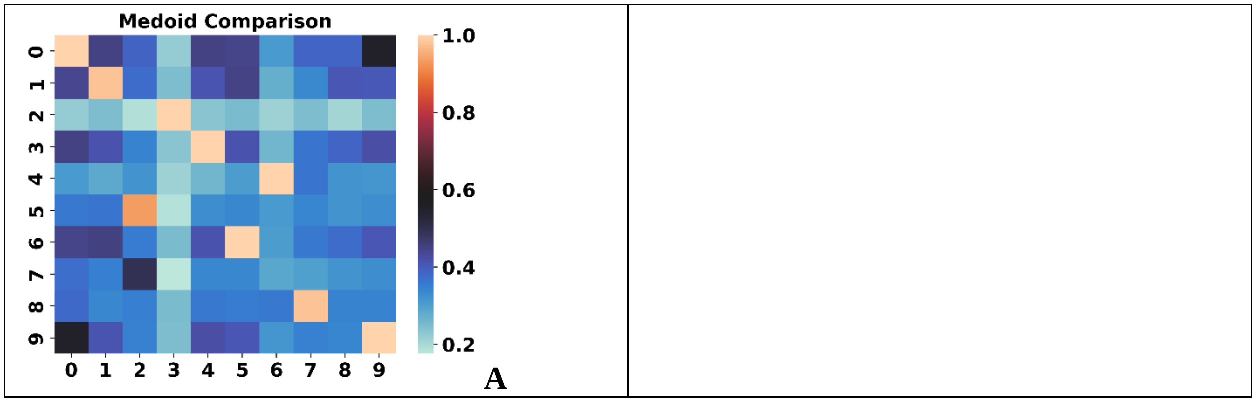

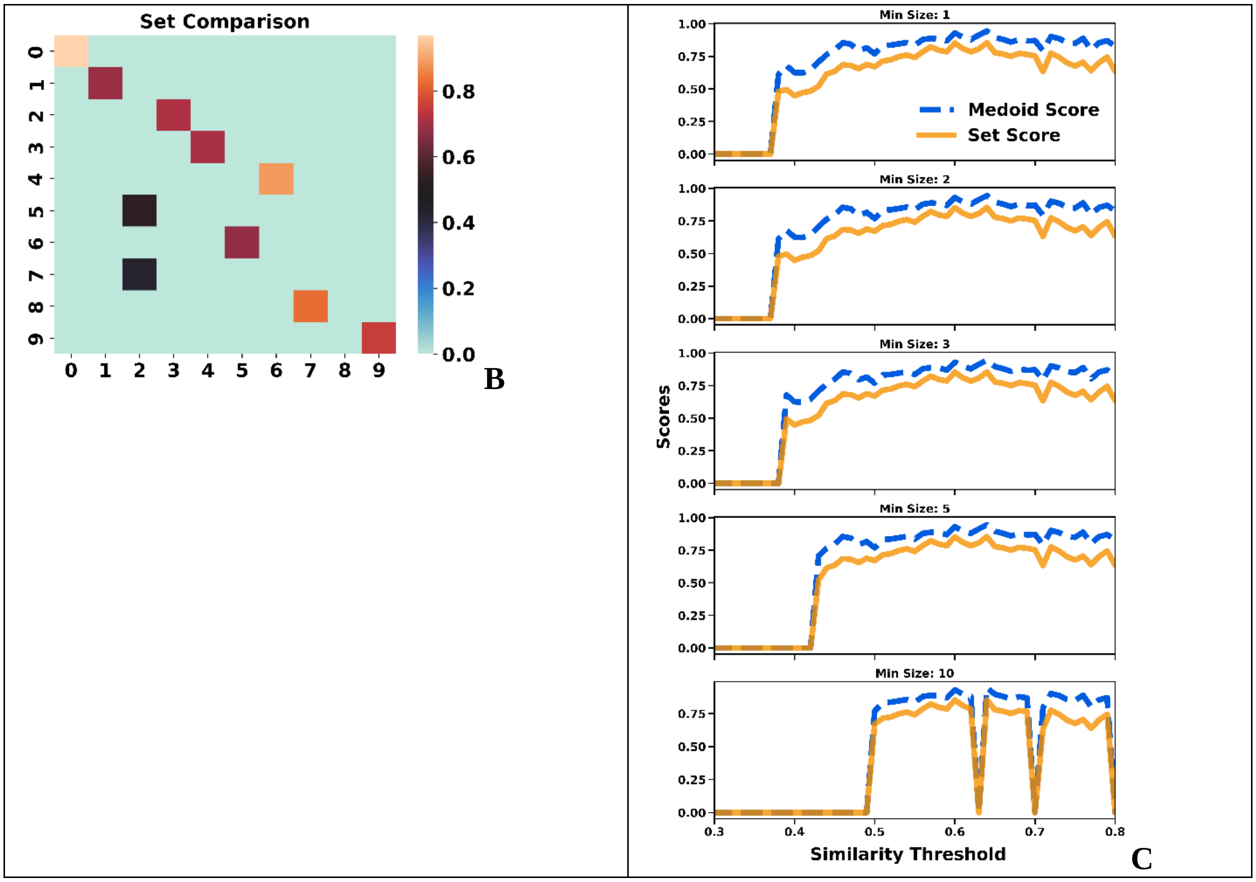


**Figure S3.3**: Comparison of the **A**: medoids, **B**: sets for the top populated clusters of the [3] ChEMBL subset (similarity threshold = 0.65, min_size = 10). **C**: Medoid and Set scores.

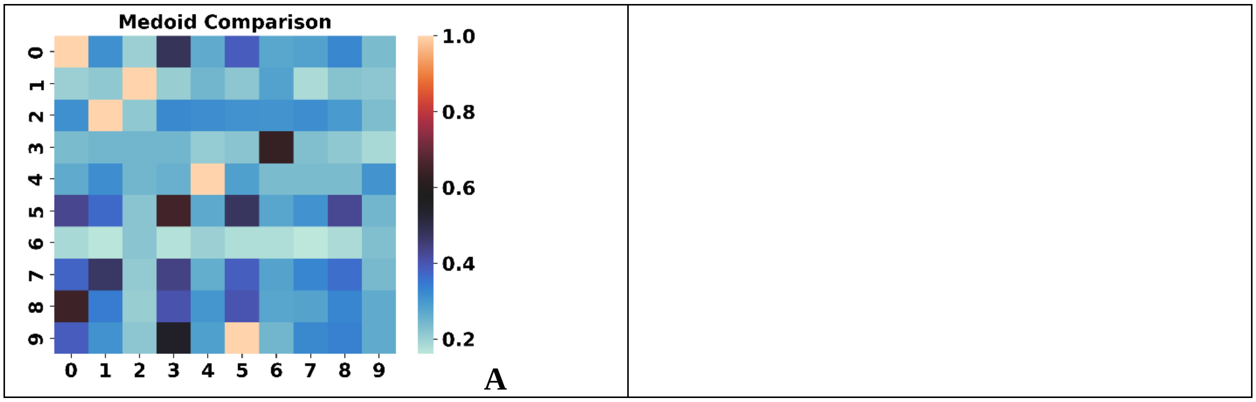

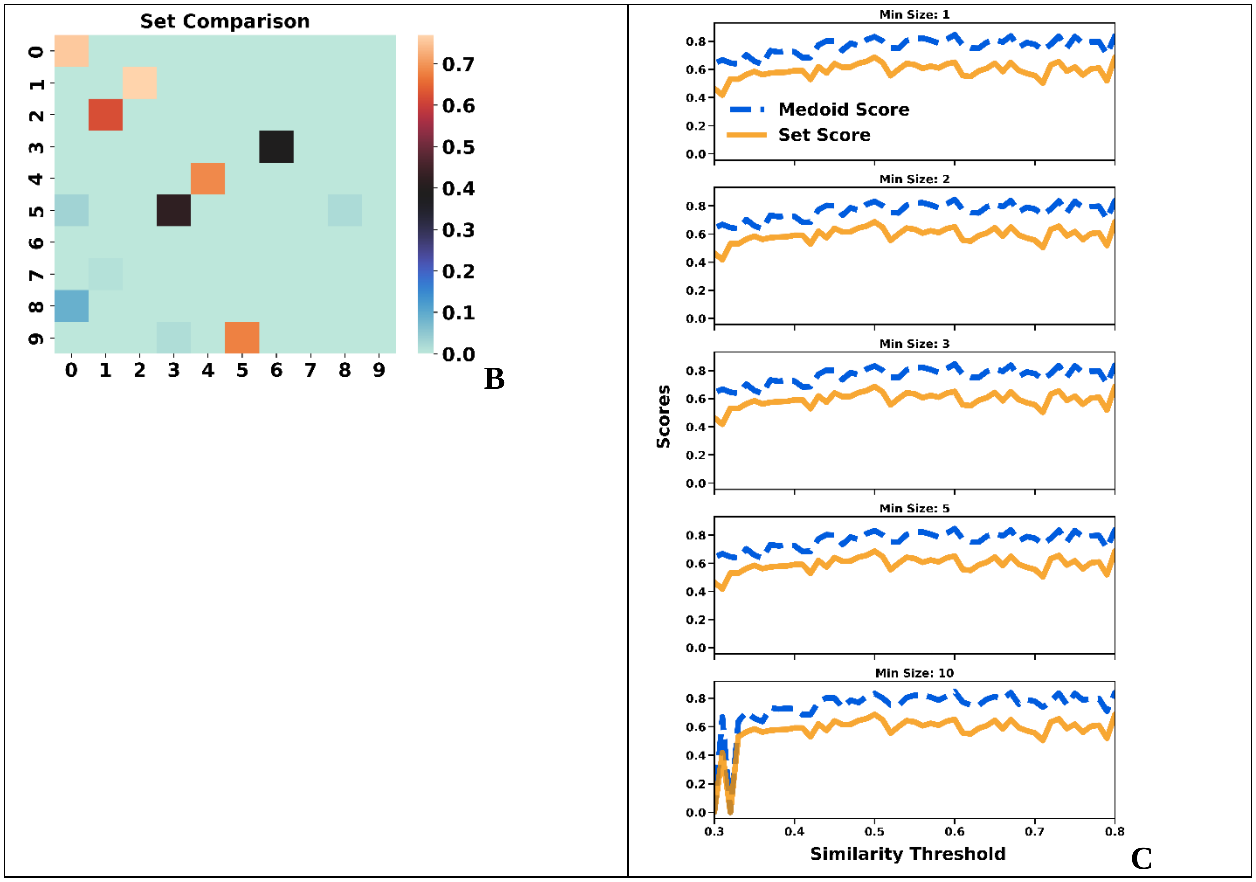


**Figure S3.4**: Comparison of the **A**: medoids, **B**: sets for the top populated clusters of the [4] ChEMBL subset (similarity threshold = 0.65, min_size = 10). **C**: Medoid and Set scores.

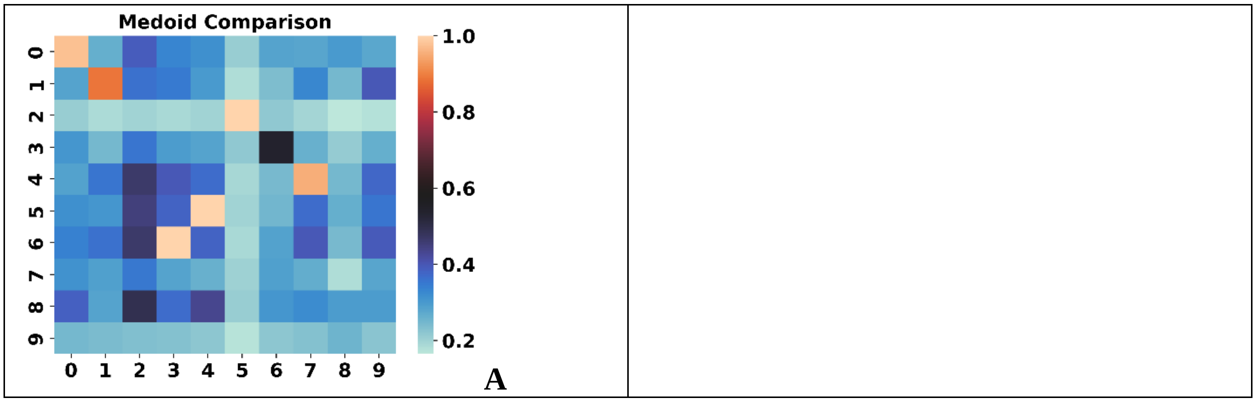

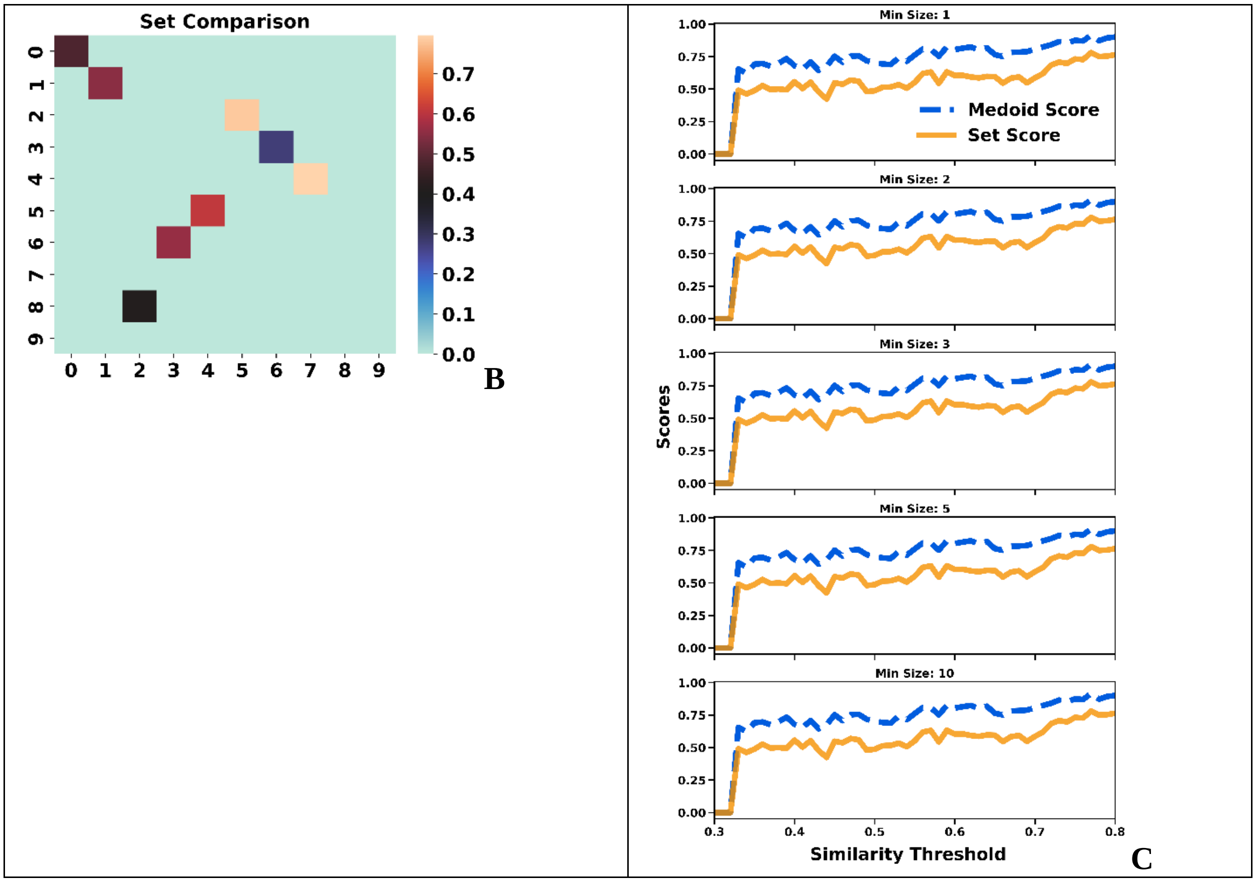


**Figure S3.5**: Comparison of the **A**: medoids, **B**: sets for the top populated clusters of the [5] ChEMBL subset (similarity threshold = 0.65, min_size = 10). **C**: Medoid and Set scores.

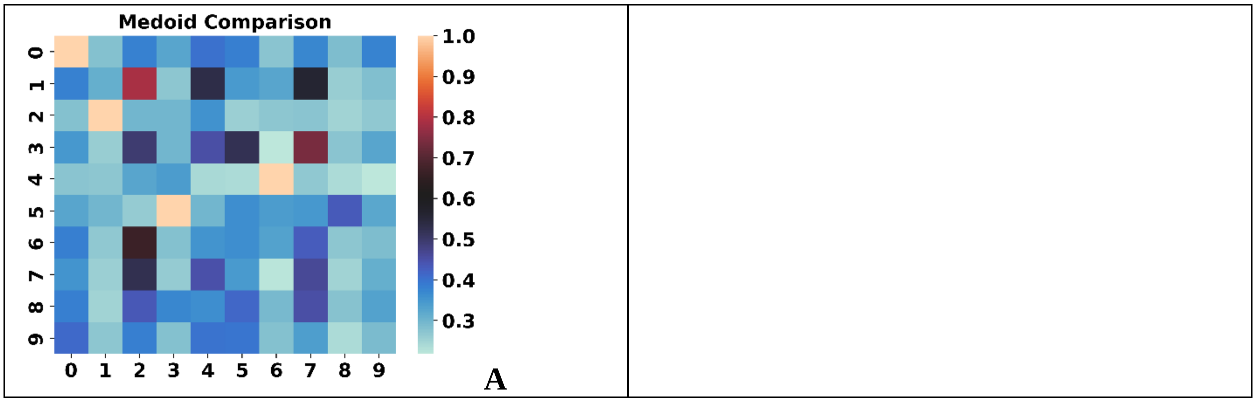

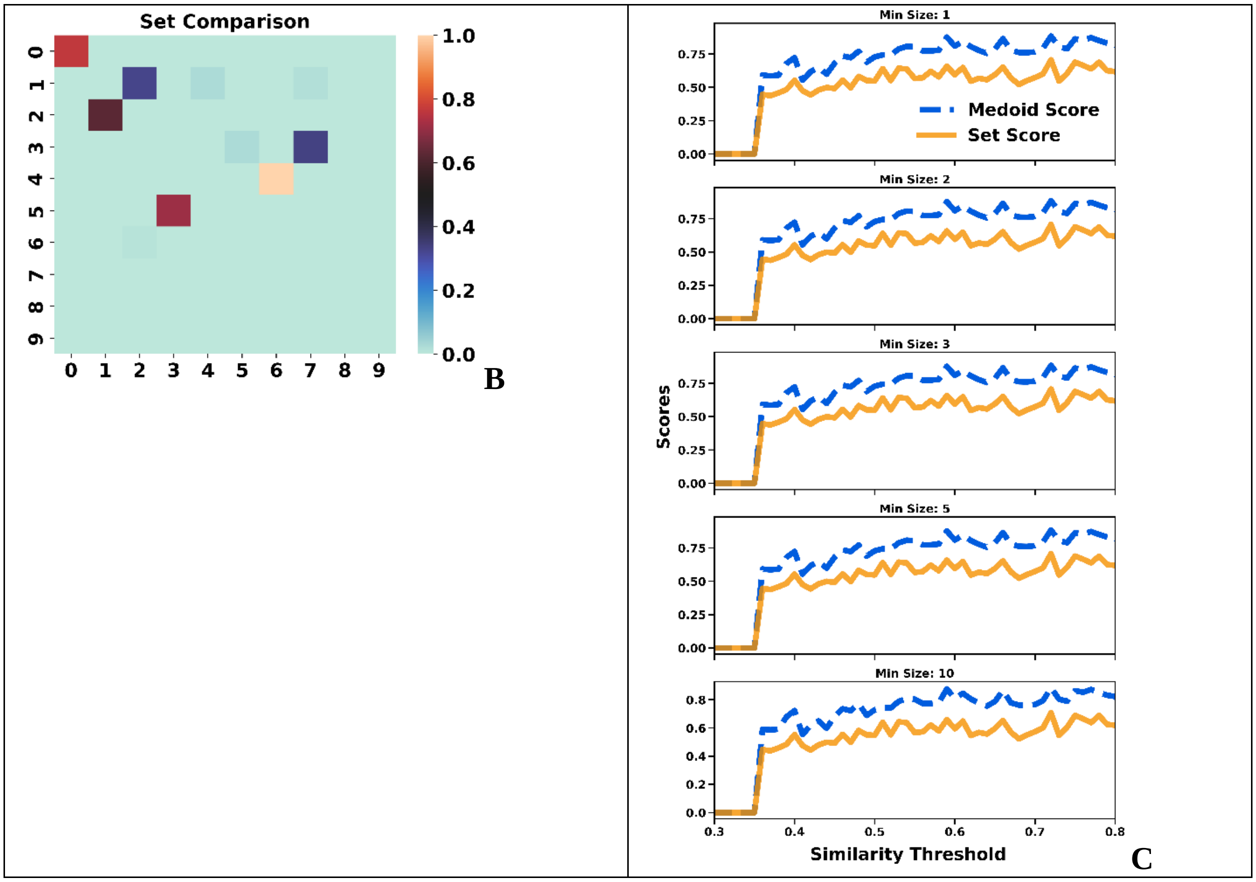


**Figure S3.6**: Comparison of the **A**: medoids, **B**: sets for the top populated clusters of the [6] ChEMBL subset (similarity threshold = 0.65, min_size = 10). **C**: Medoid and Set scores.

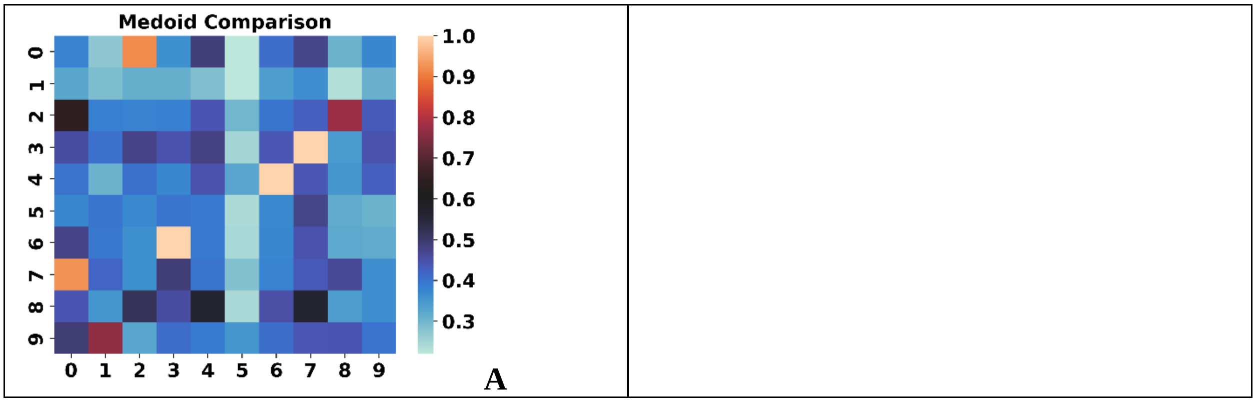

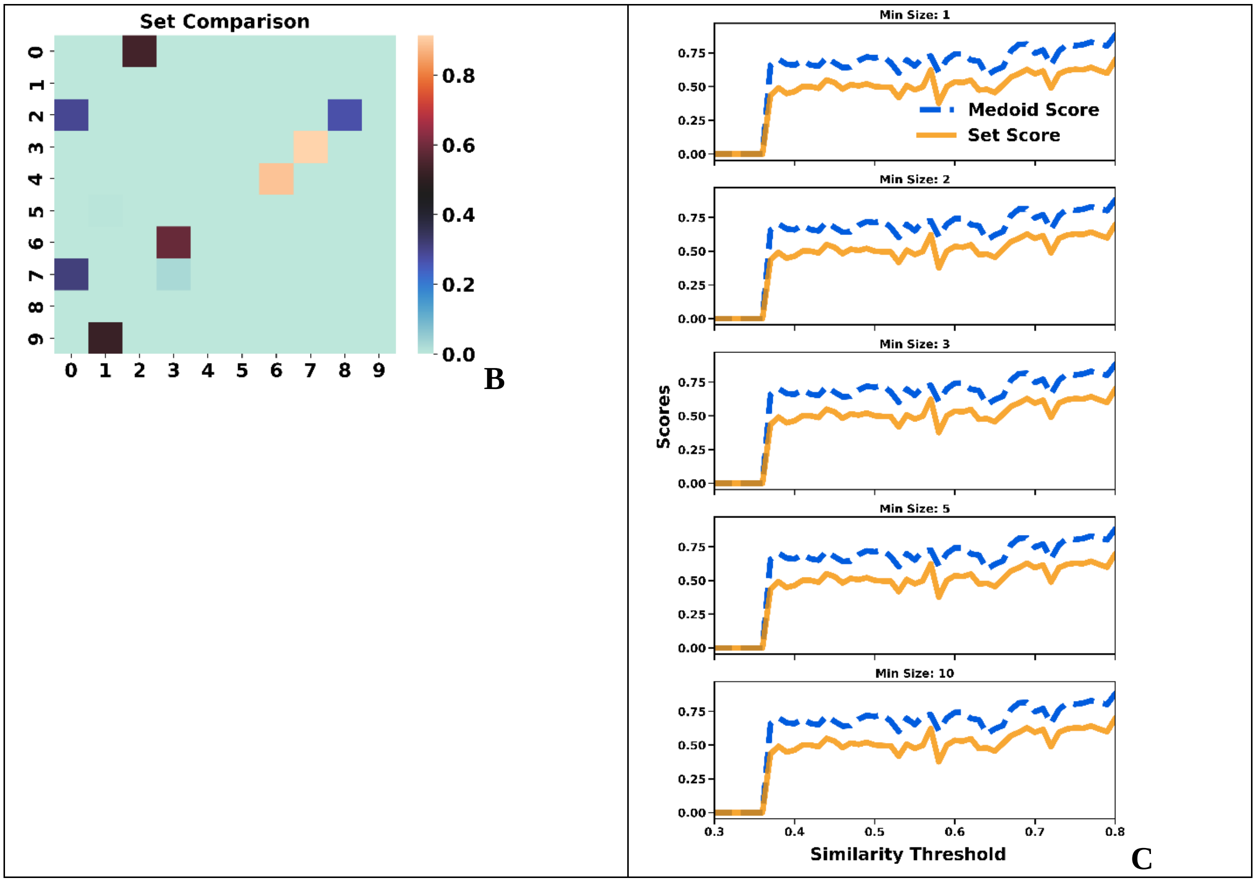


**Figure S3.7**: Comparison of the **A**: medoids, **B**: sets for the top populated clusters of the [7] ChEMBL subset (similarity threshold = 0.65, min_size = 10). **C**: Medoid and Set scores.

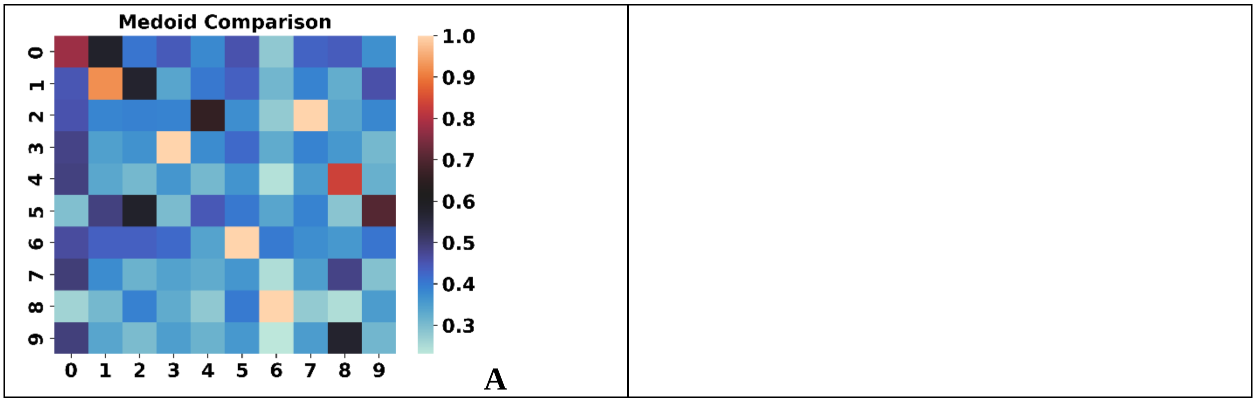

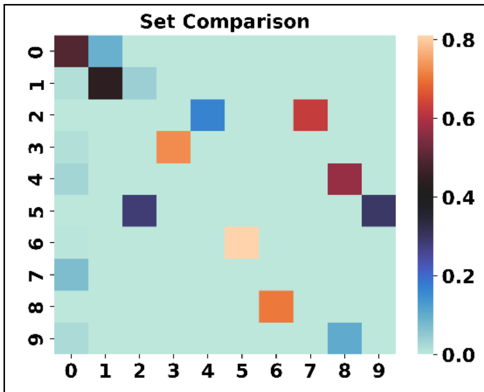


B

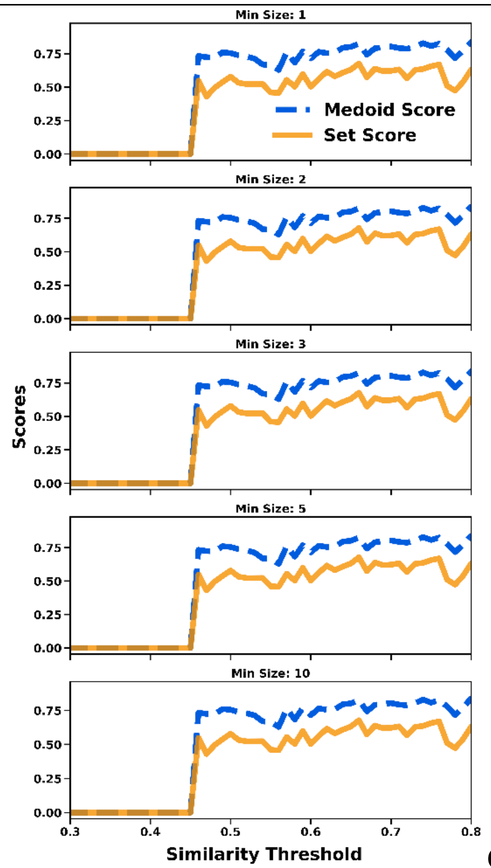


C

**Figure S3.8**: Comparison of the **A**: medoids, **B**: sets for the top populated clusters of the [9] ChEMBL subset (similarity threshold = 0.65, min_size = 10). **C**: Medoid and Set scores.

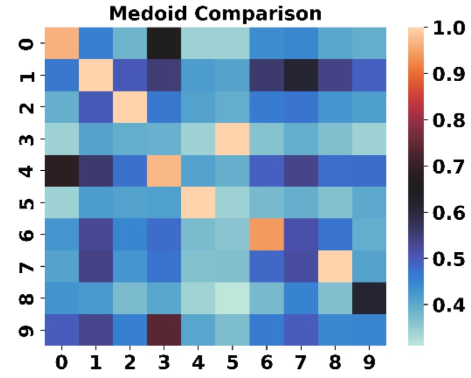


A

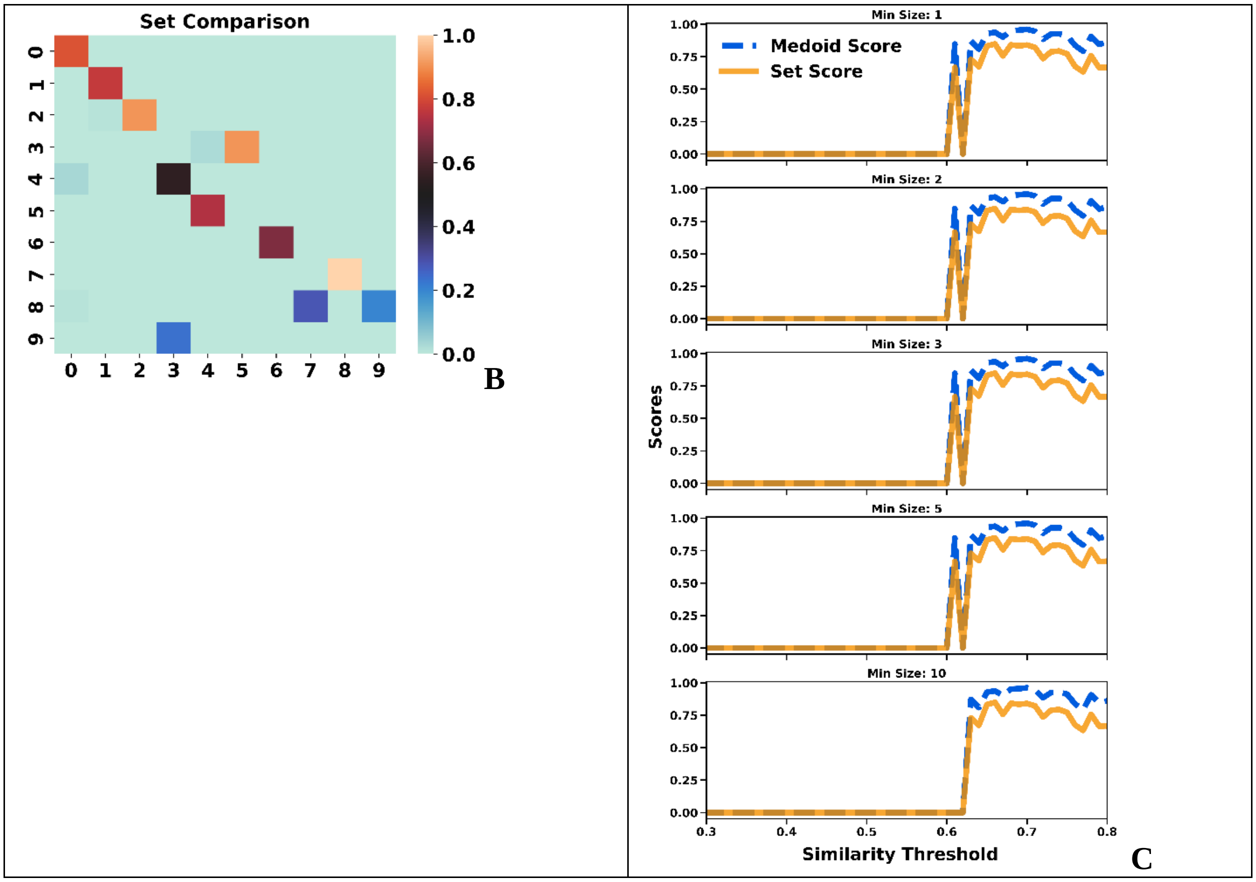


**Figure S3.9**: Comparison of the **A**: medoids, **B**: sets for the top populated clusters of the [10] ChEMBL subset (similarity threshold = 0.65, min_size = 10). **C**: Medoid and Set scores.

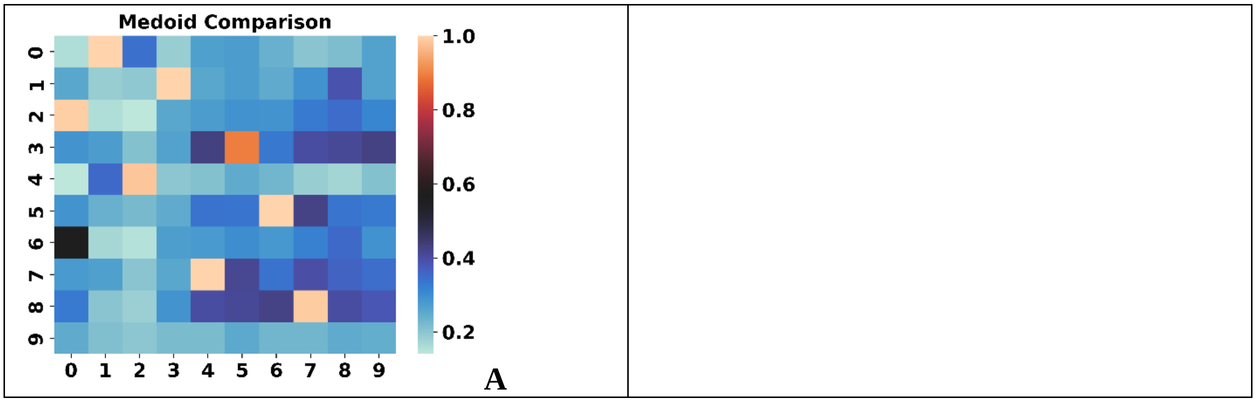

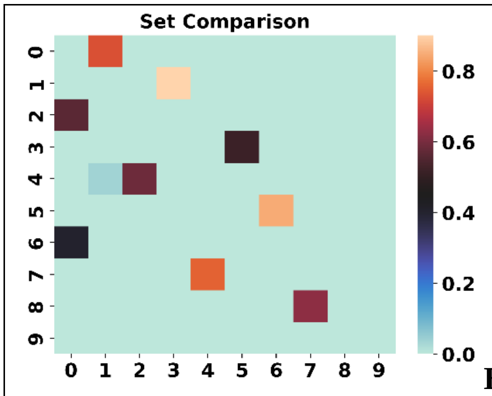


B

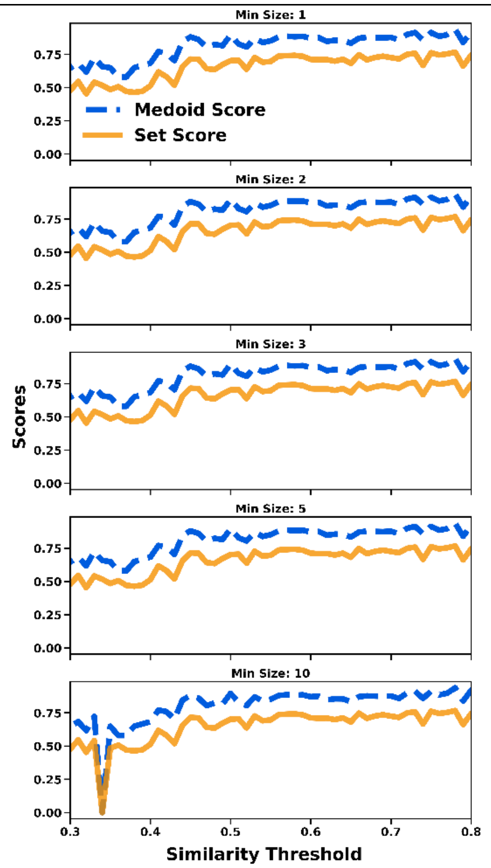


**Figure S3.10**: Comparison of the **A**: medoids, **B**: sets for the top populated clusters of the [12] ChEMBL subset (similarity threshold = 0.65, min_size = 10). **C**: Medoid and Set scores.

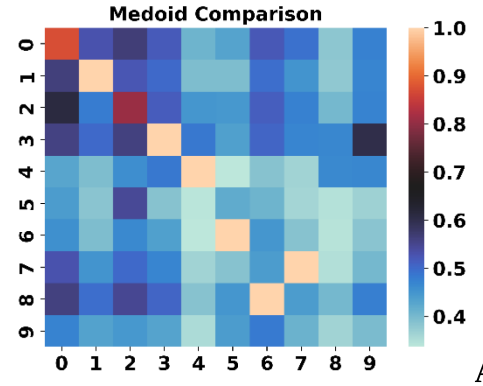


A

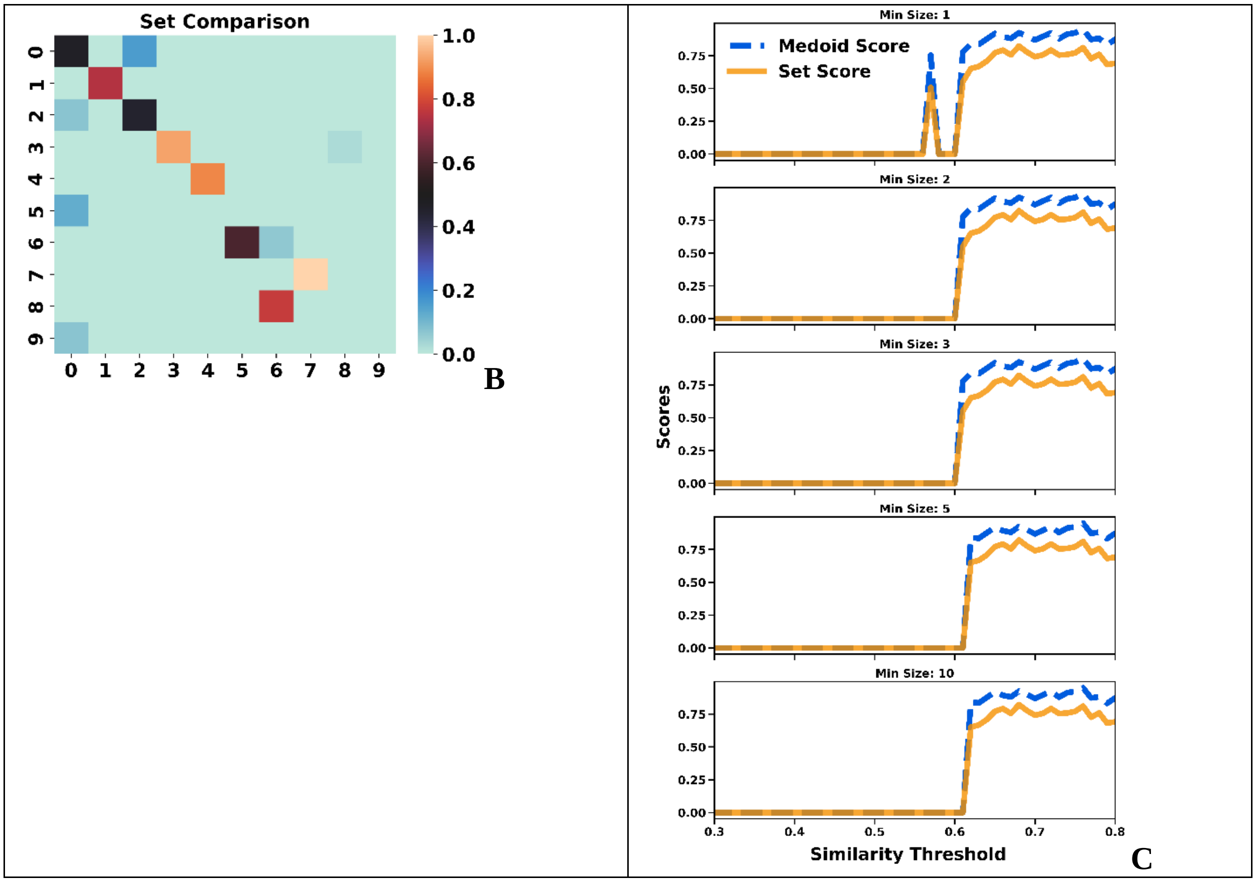


**Figure S3.11**: Comparison of the **A**: medoids, **B**: sets for the top populated clusters of the [14] ChEMBL subset (similarity threshold = 0.65, min_size = 10). **C**: Medoid and Set scores.

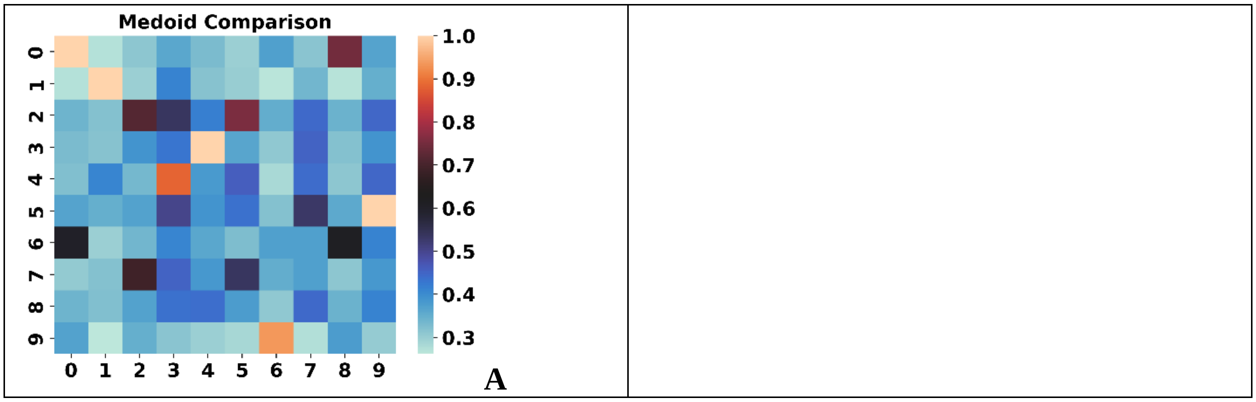

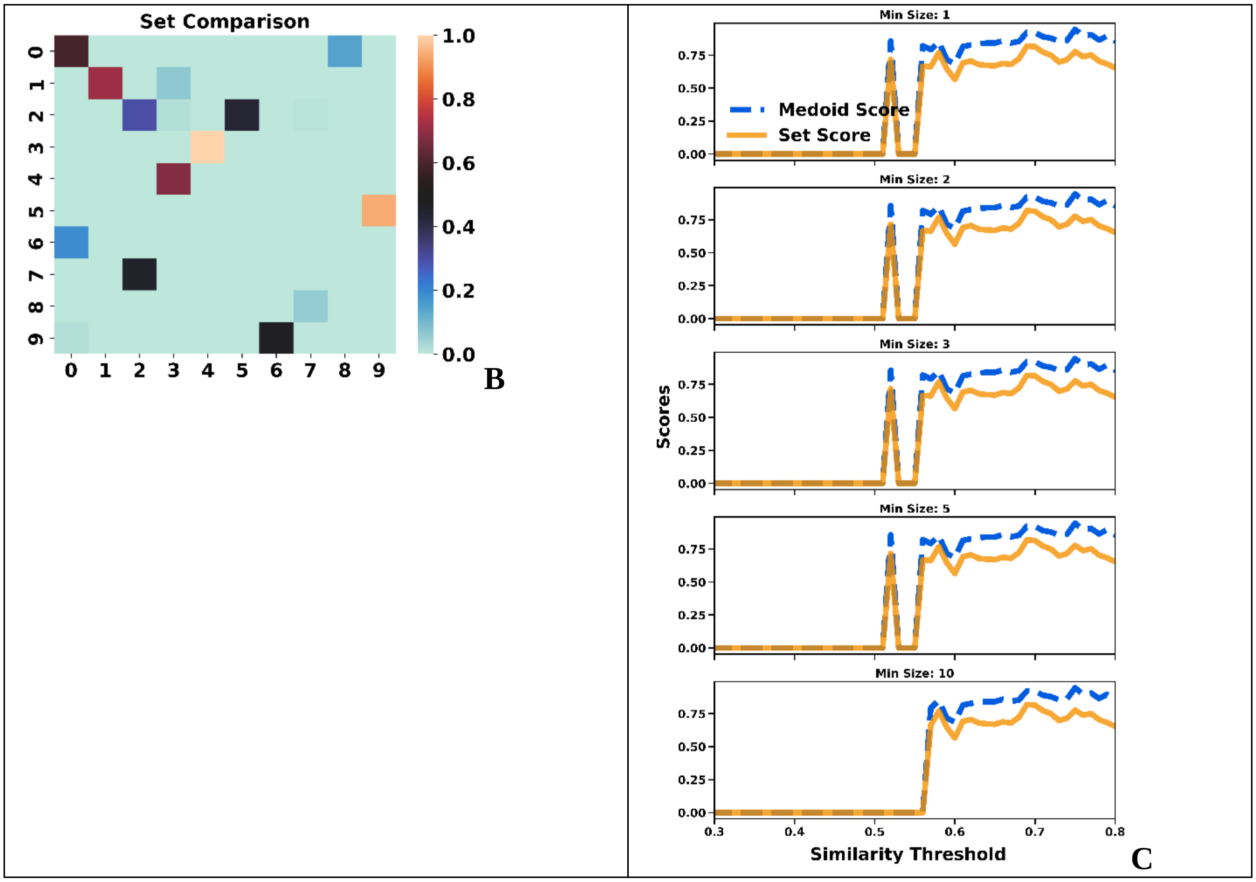


**Figure S3.12**: Comparison of the **A**: medoids, **B**: sets for the top populated clusters of the [16] ChEMBL subset (similarity threshold = 0.65, min_size = 10). **C**: Medoid and Set scores.

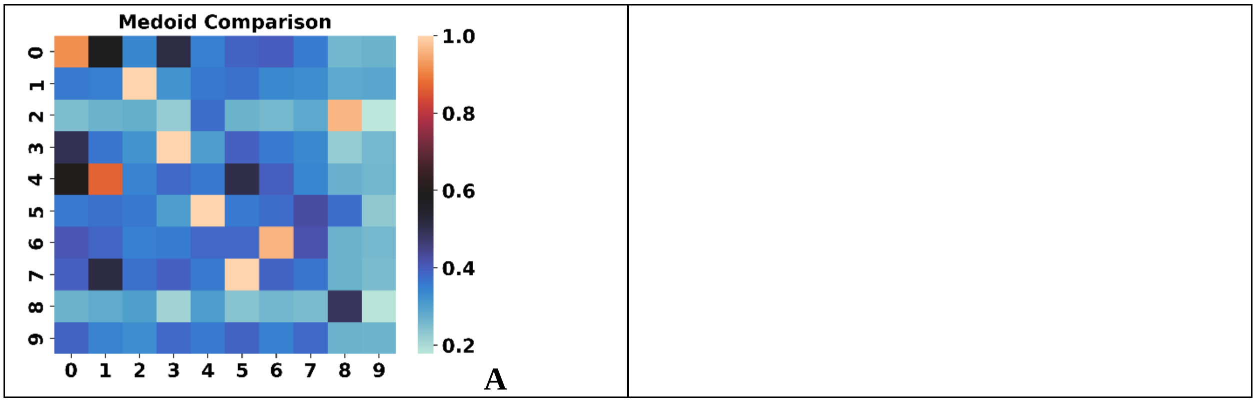

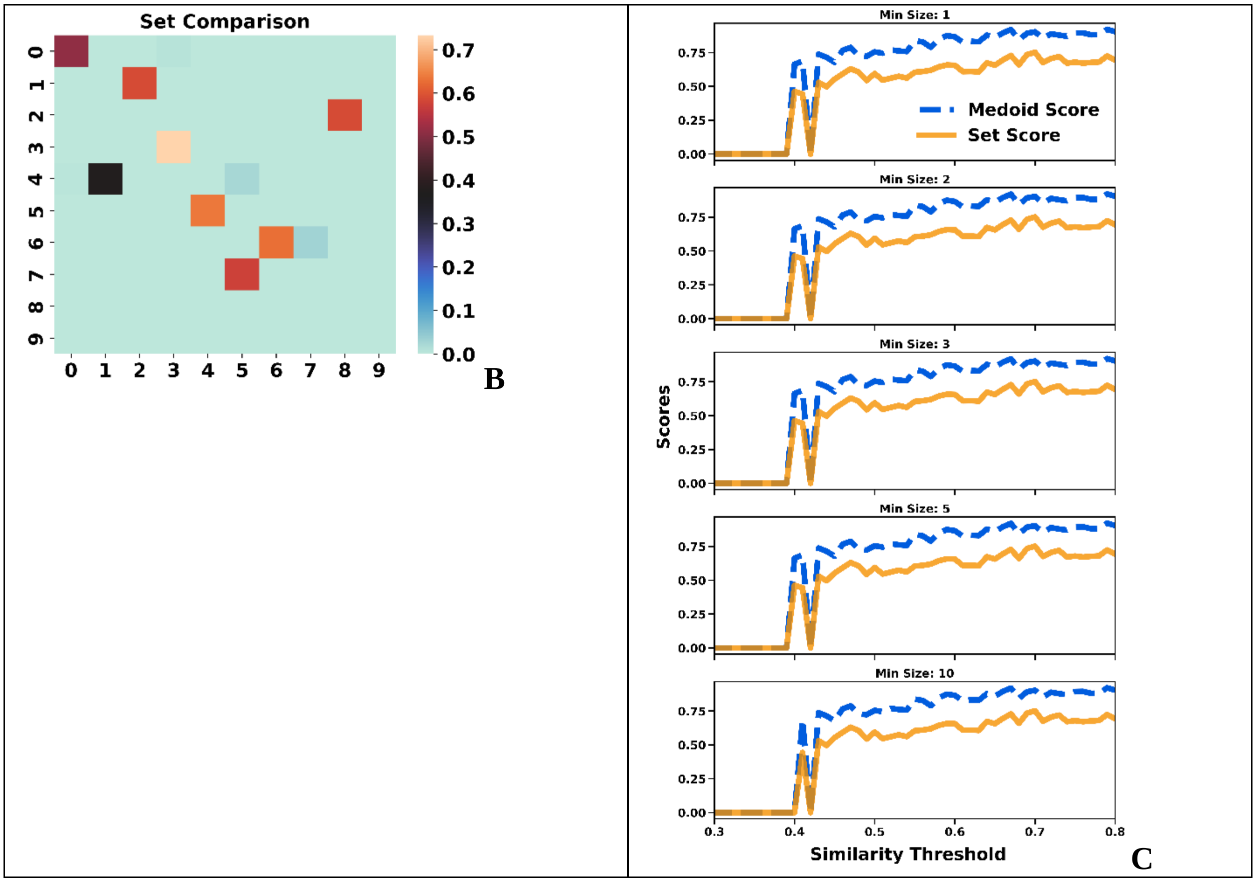


**Figure S3.13**: Comparison of the **A**: medoids, **B**: sets for the top populated clusters of the [17] ChEMBL subset (similarity threshold = 0.65, min_size = 10). **C**: Medoid and Set scores.

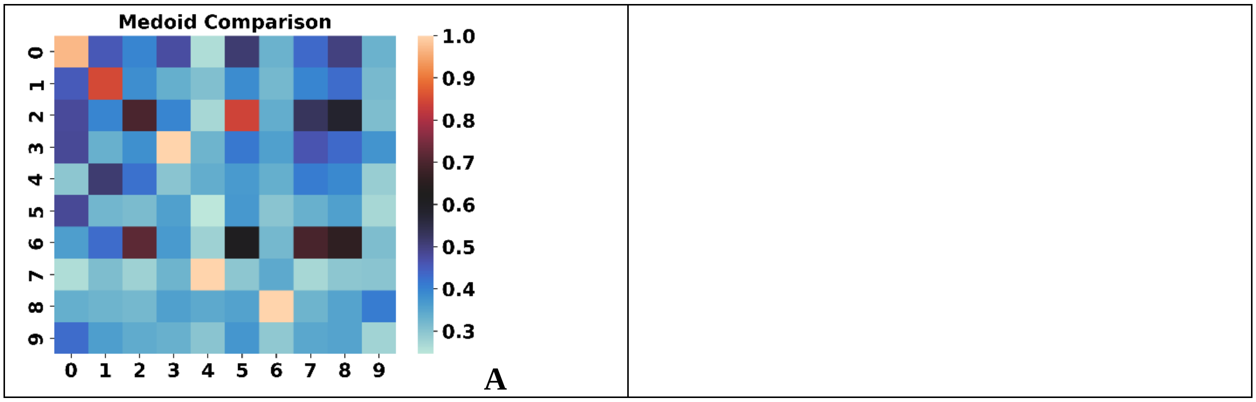

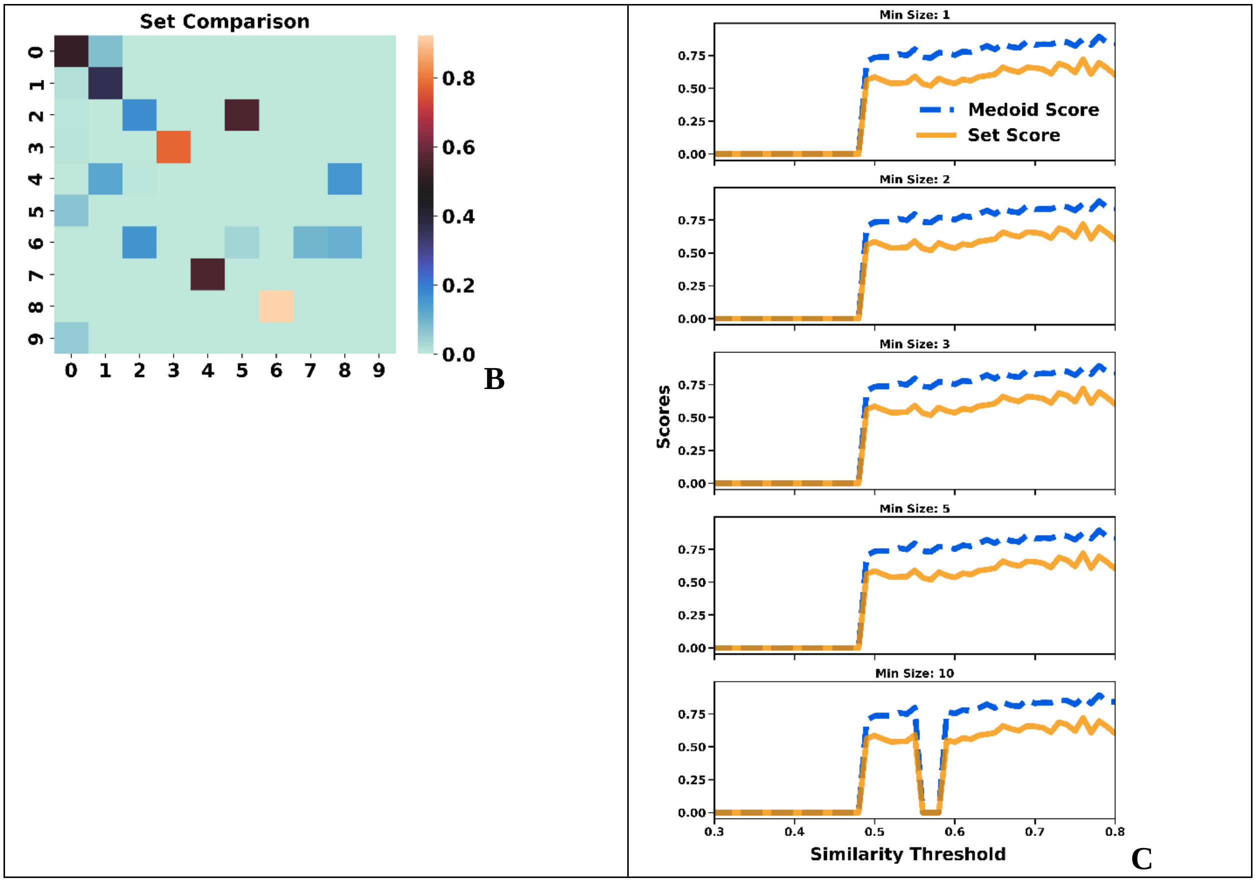


**Figure S3.14**: Comparison of the **A**: medoids, **B**: sets for the top populated clusters of the [20] ChEMBL subset (similarity threshold = 0.65, min_size = 10). **C**: Medoid and Set scores.

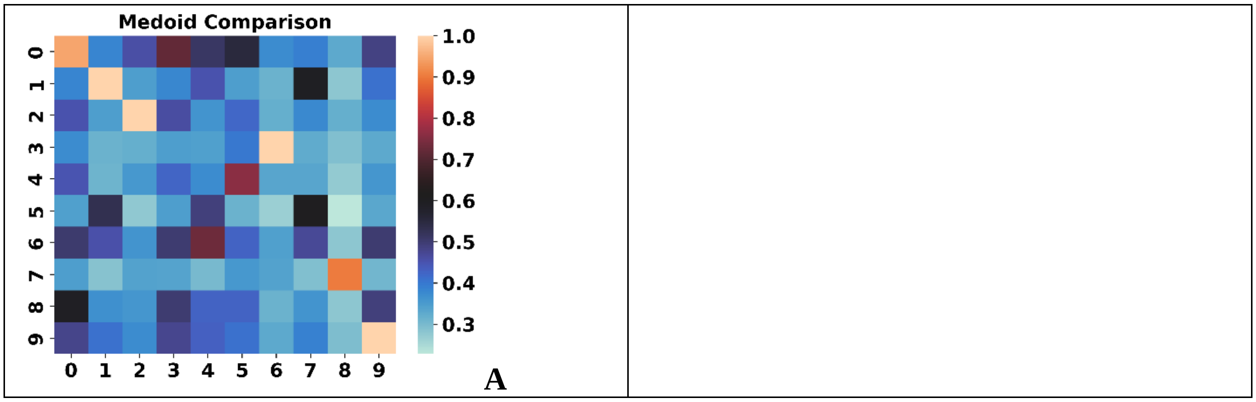

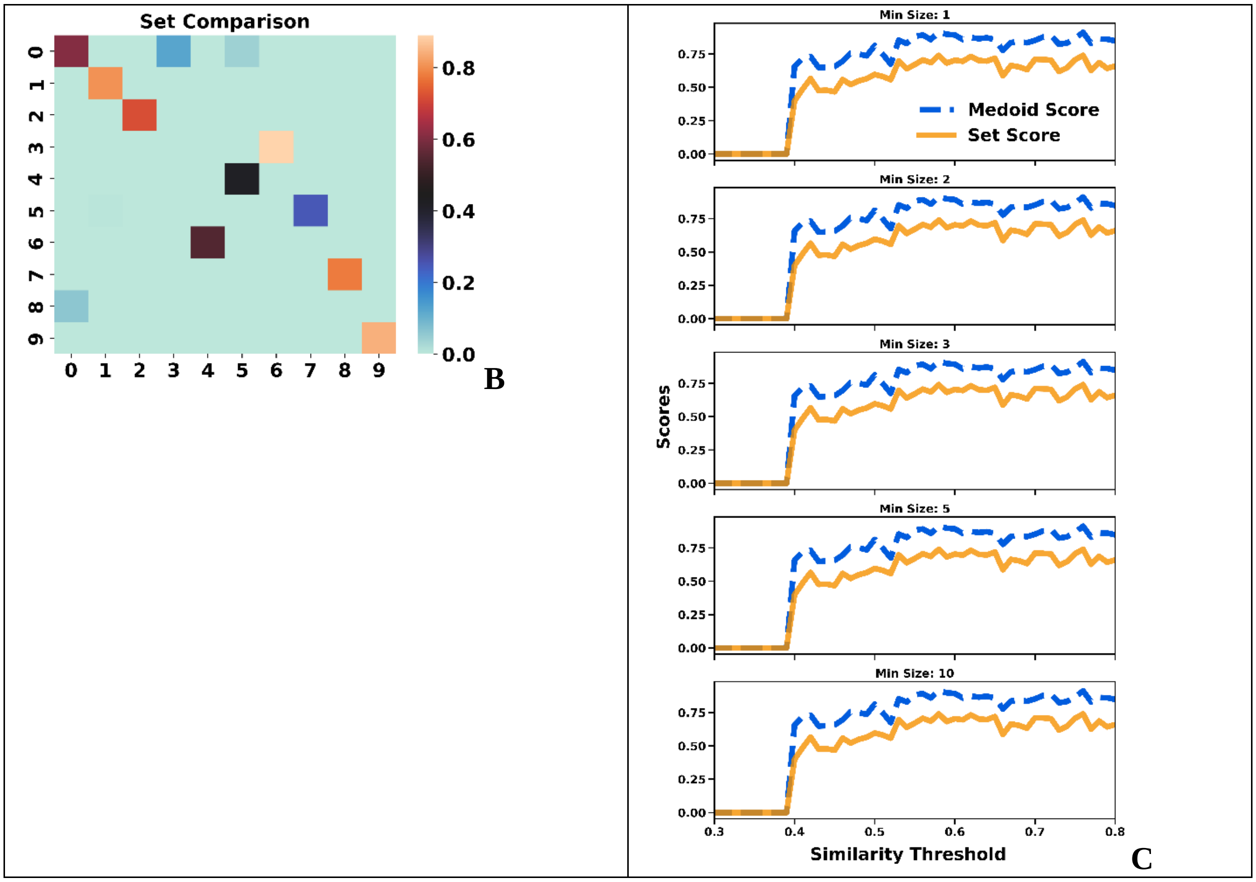


**Figure S3.15**: Comparison of the **A**: medoids, **B**: sets for the top populated clusters of the [21] ChEMBL subset (similarity threshold = 0.65, min_size = 10). **C**: Medoid and Set scores.

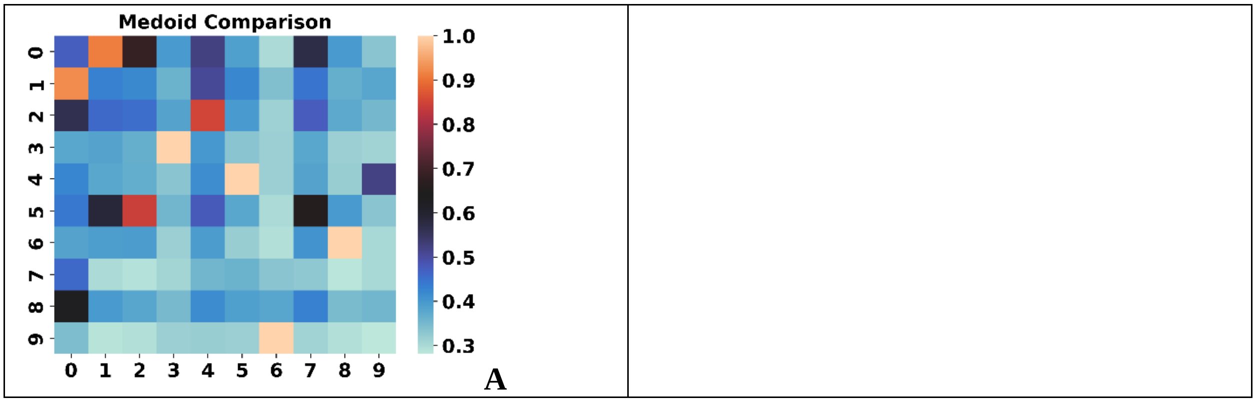

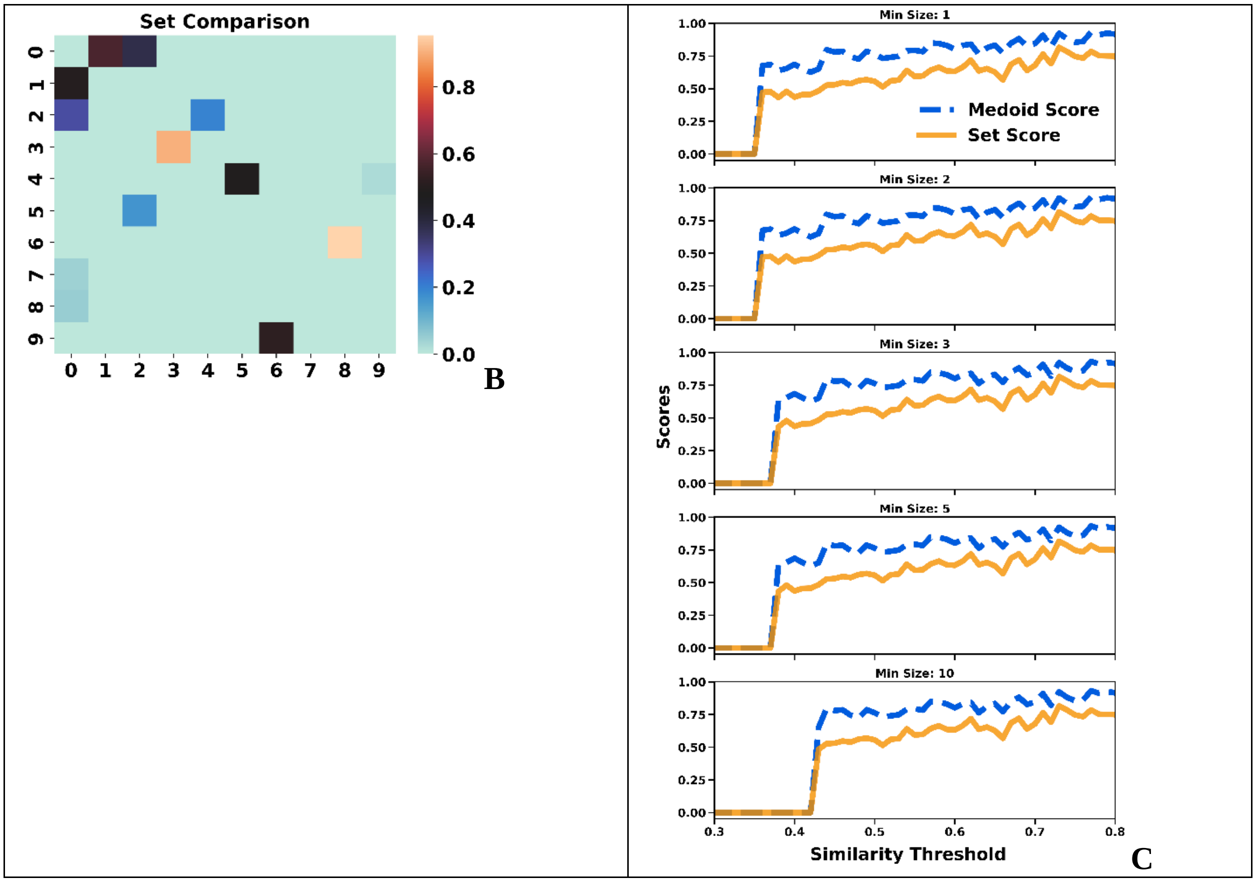


**Figure S3.16**: Comparison of the **A**: medoids, **B**: sets for the top populated clusters of the [22] ChEMBL subset (similarity threshold = 0.65, min_size = 10). **C**: Medoid and Set scores.

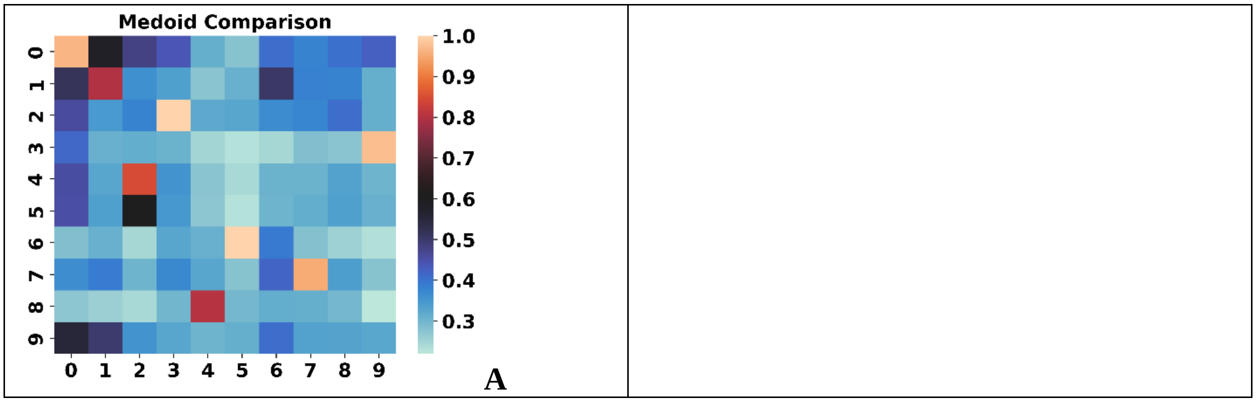

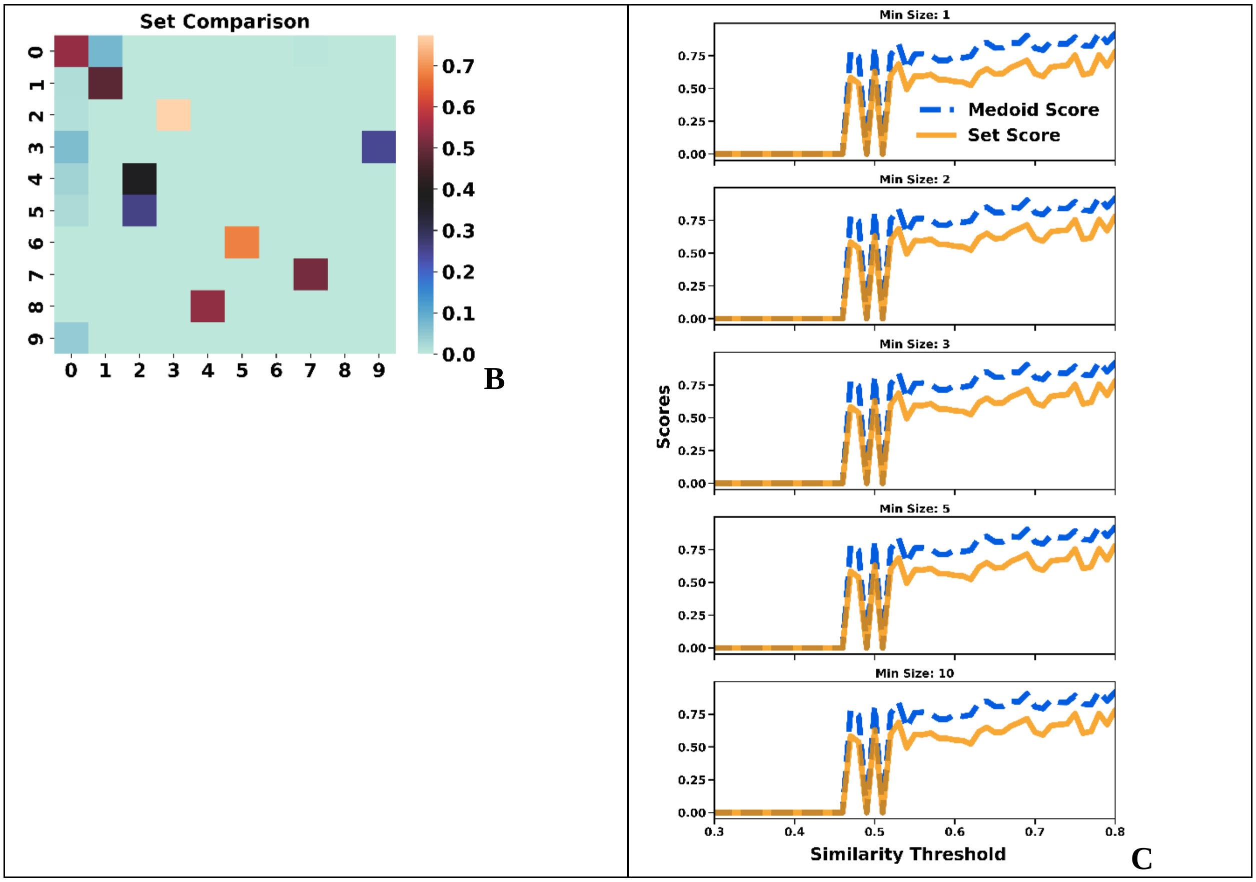


**Figure S3.17**: Comparison of the **A**: medoids, **B**: sets for the top populated clusters of the [23] ChEMBL subset (similarity threshold = 0.65, min_size = 10). **C**: Medoid and Set scores.

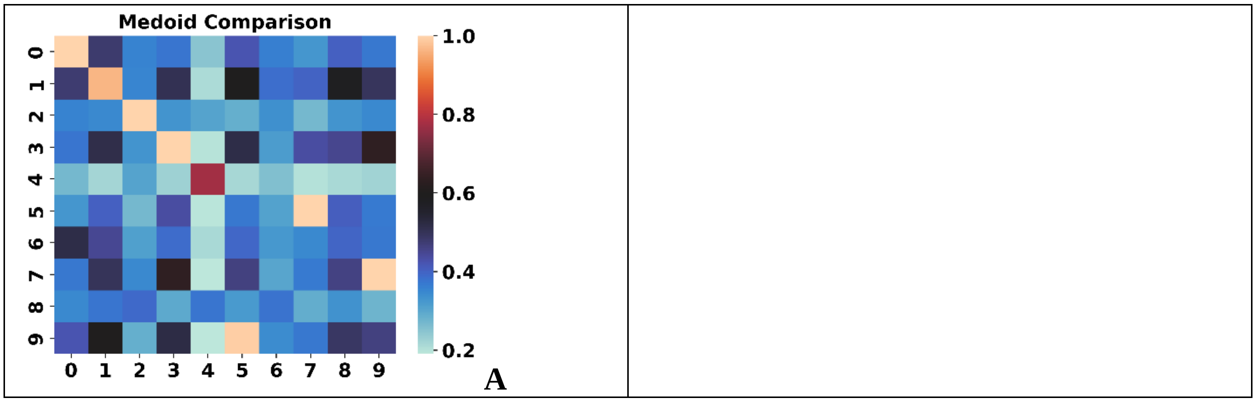

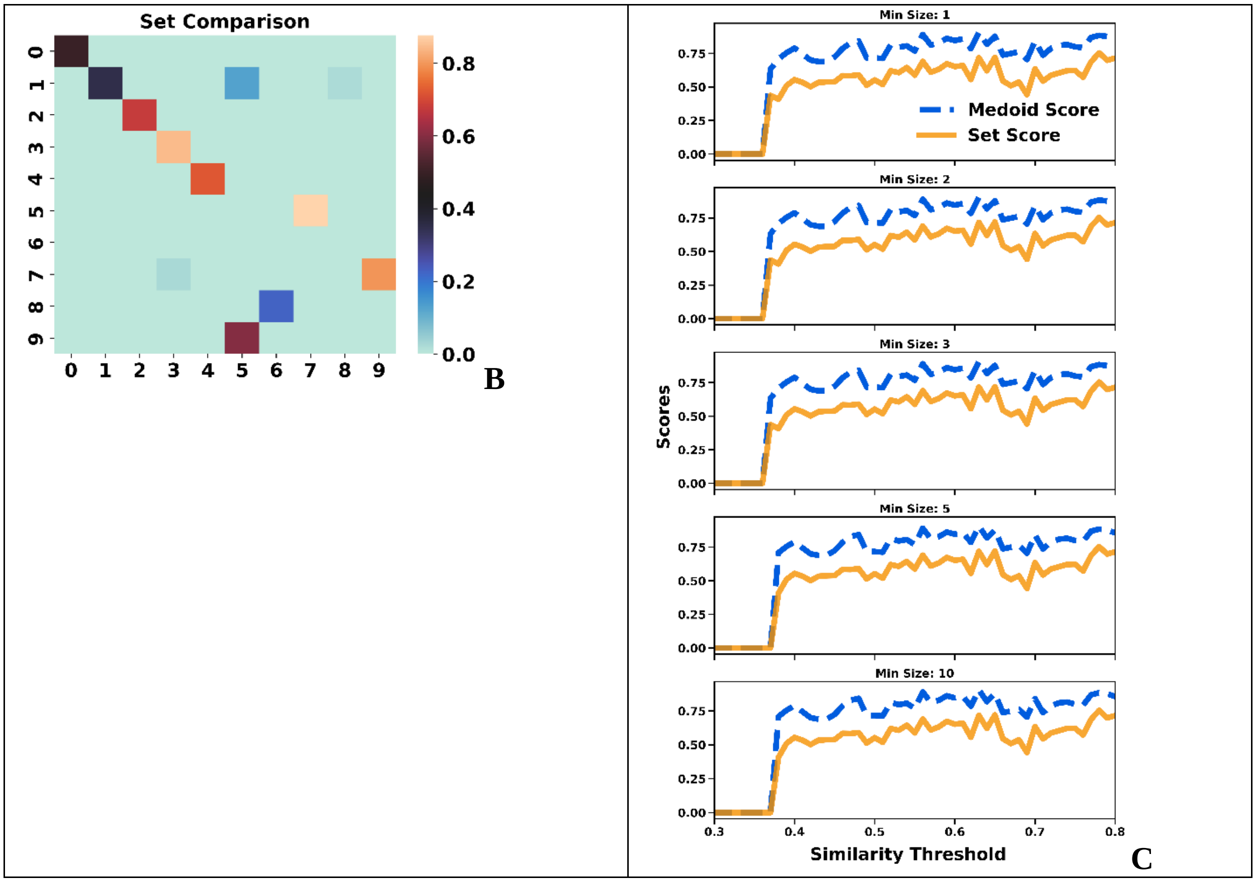


**Figure S3.18**: Comparison of the **A**: medoids, **B**: sets for the top populated clusters of the [24] ChEMBL subset (similarity threshold = 0.65, min_size = 10). **C**: Medoid and Set scores.

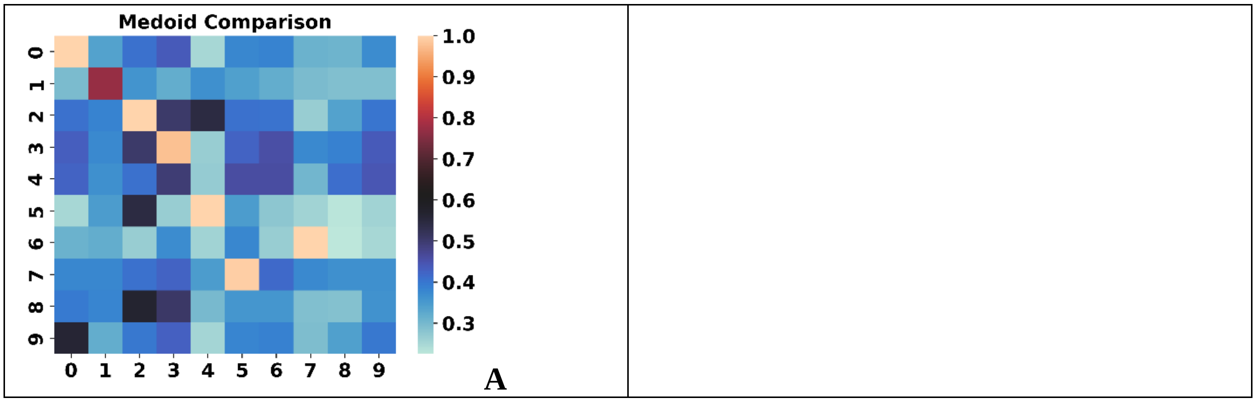

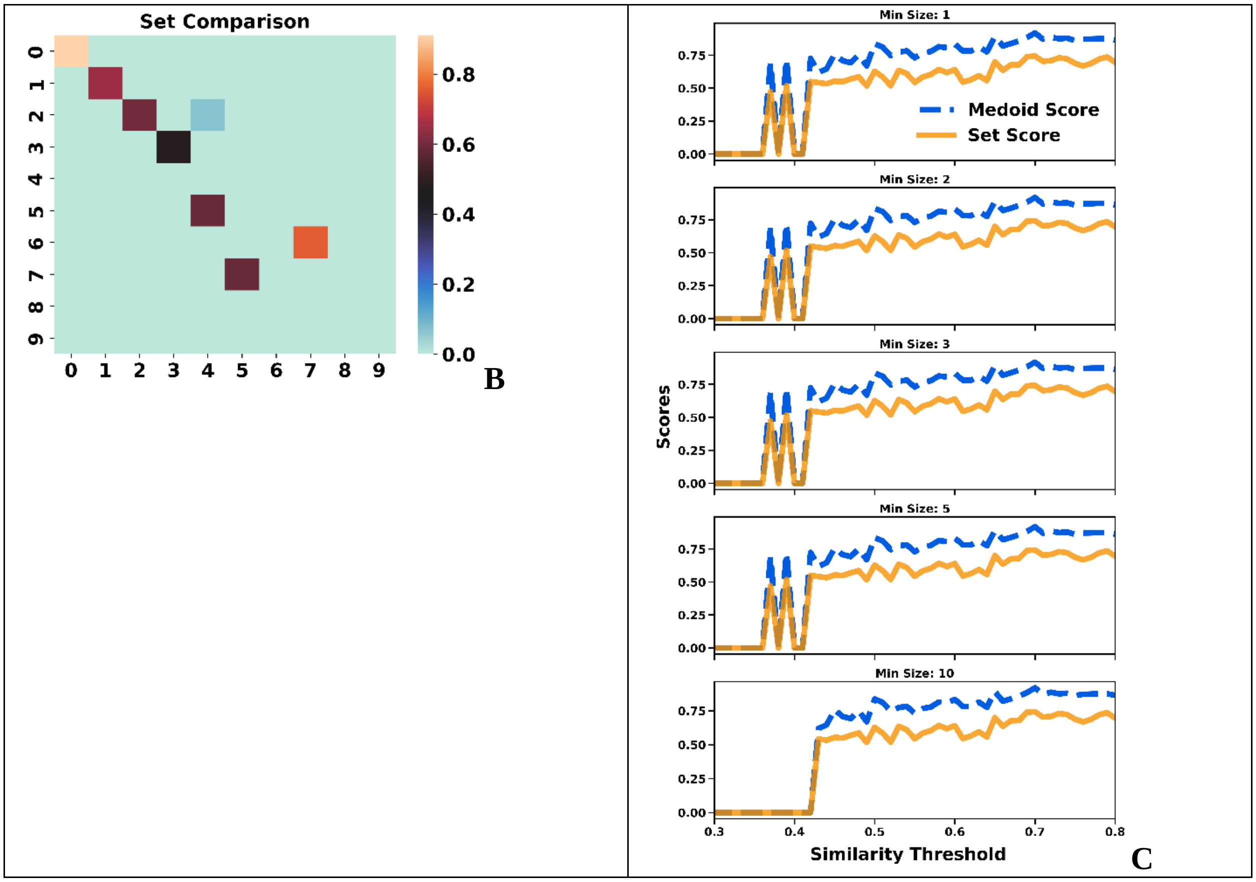


**Figure S3.19**: Comparison of the **A**: medoids, **B**: sets for the top populated clusters of the [25] ChEMBL subset (similarity threshold = 0.65, min_size = 10). **C**: Medoid and Set scores.

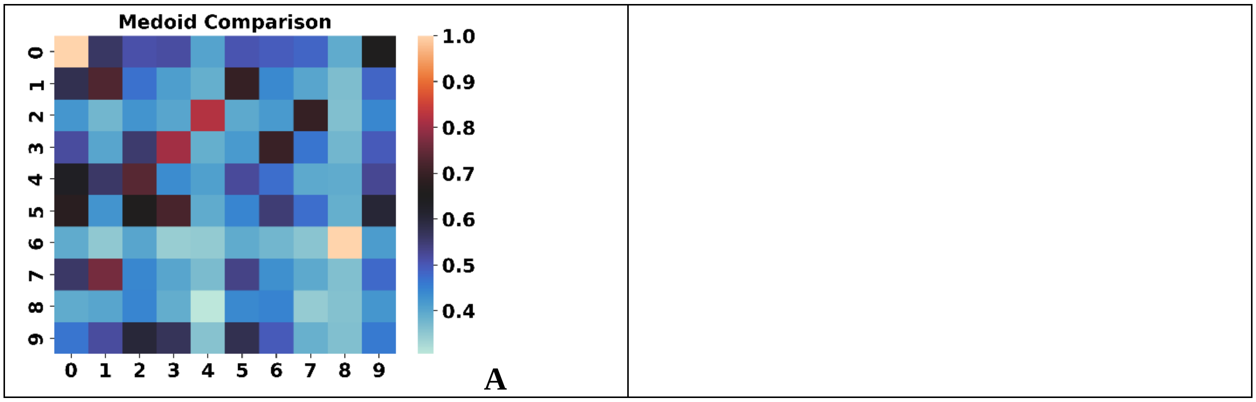

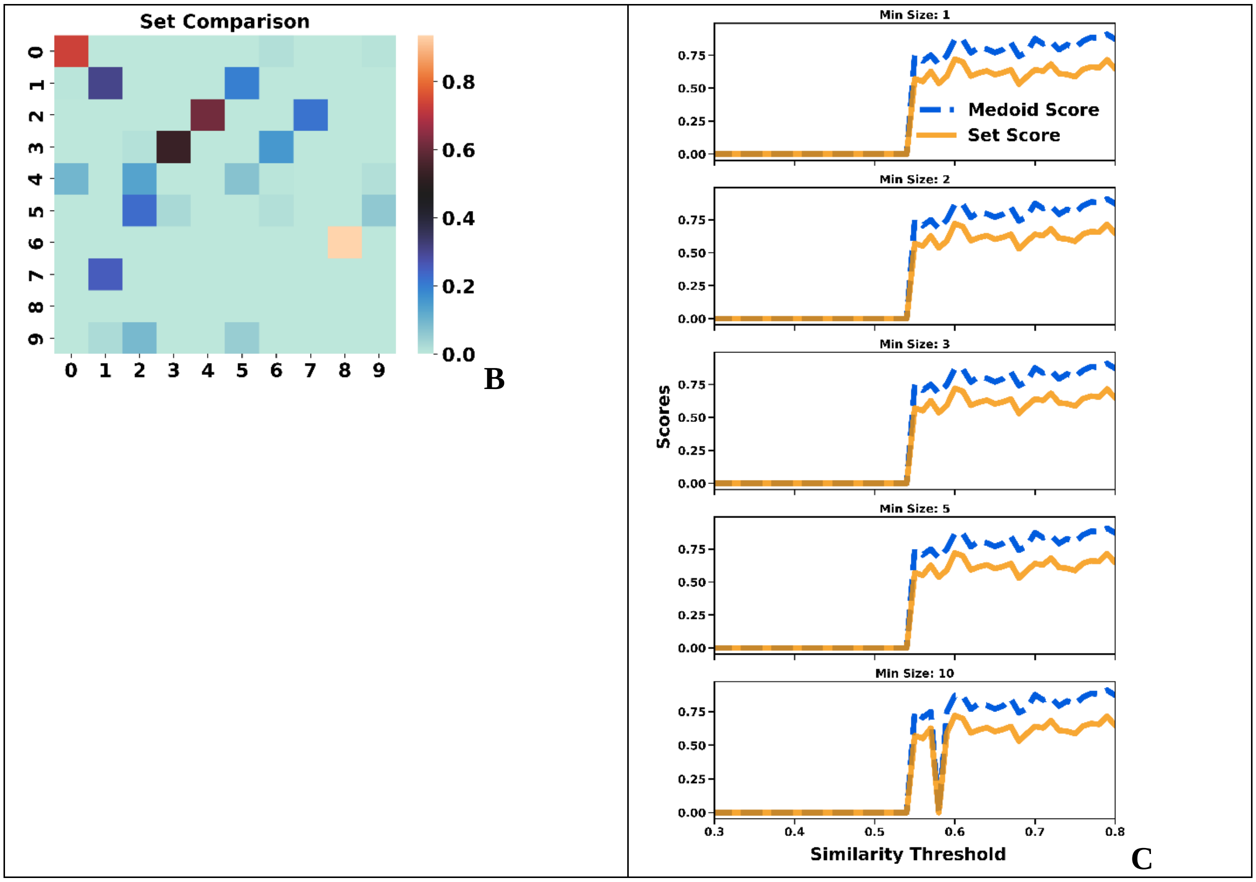


**Figure S3.20**: Comparison of the **A**: medoids, **B**: sets for the top populated clusters of the [26] ChEMBL subset (similarity threshold = 0.65, min_size = 10). **C**: Medoid and Set scores.

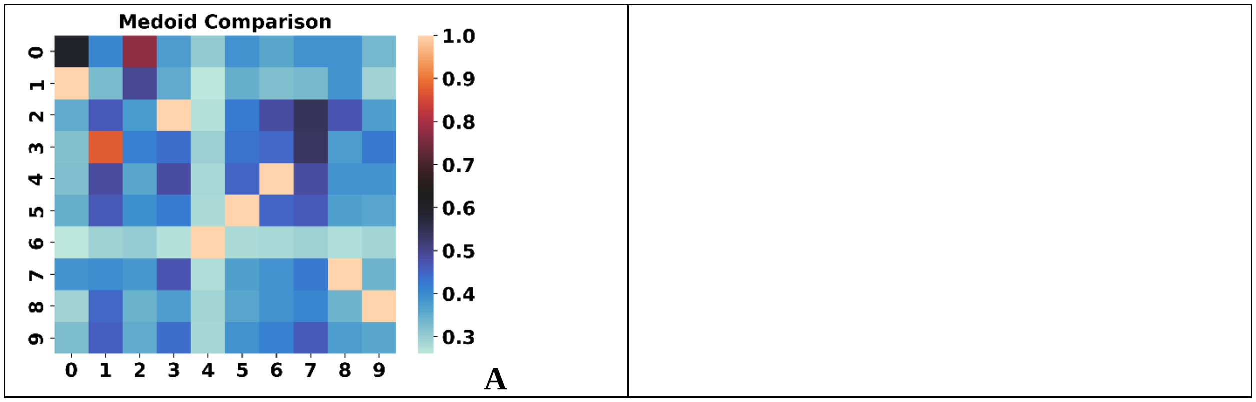

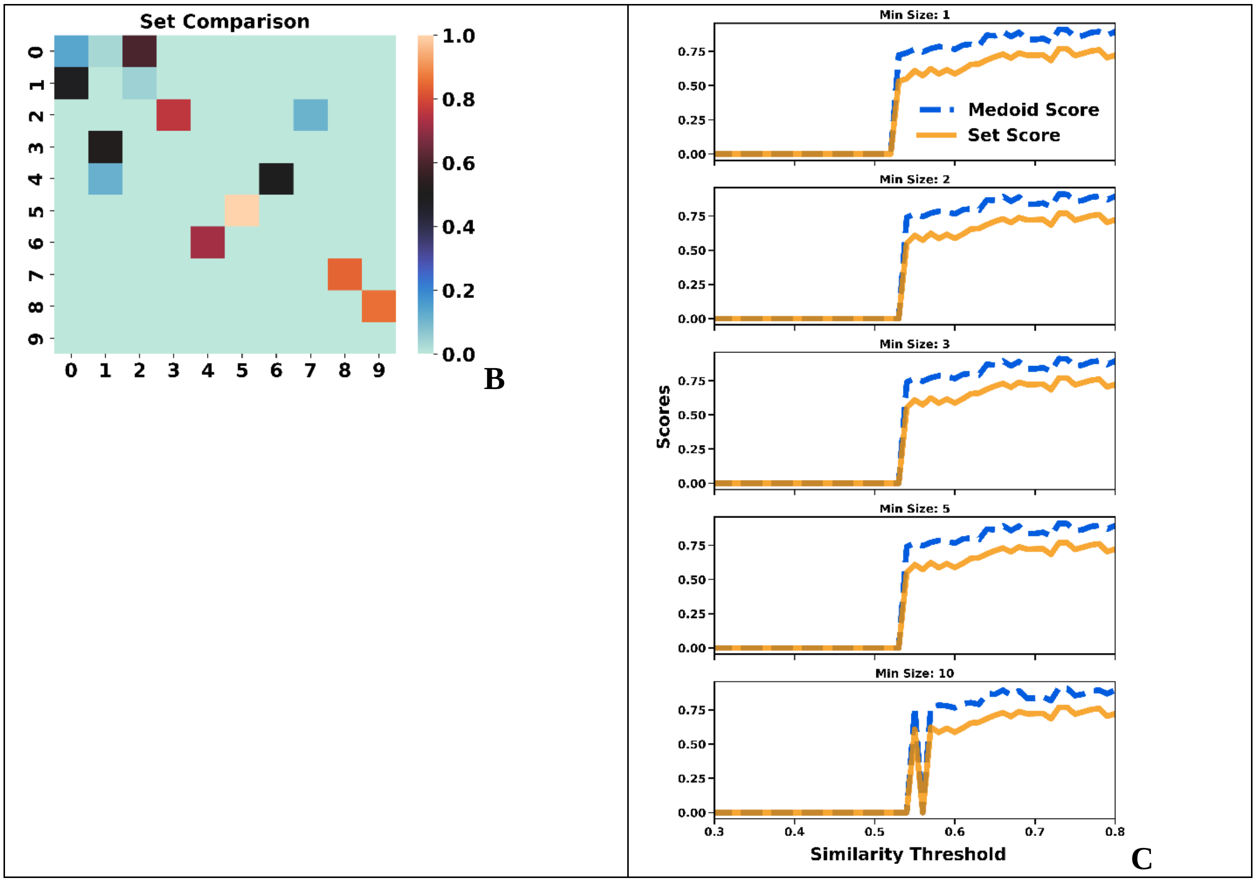


**Figure S3.21**: Comparison of the **A**: medoids, **B**: sets for the top populated clusters of the [28] ChEMBL subset (similarity threshold = 0.65, min_size = 10). **C**: Medoid and Set scores.

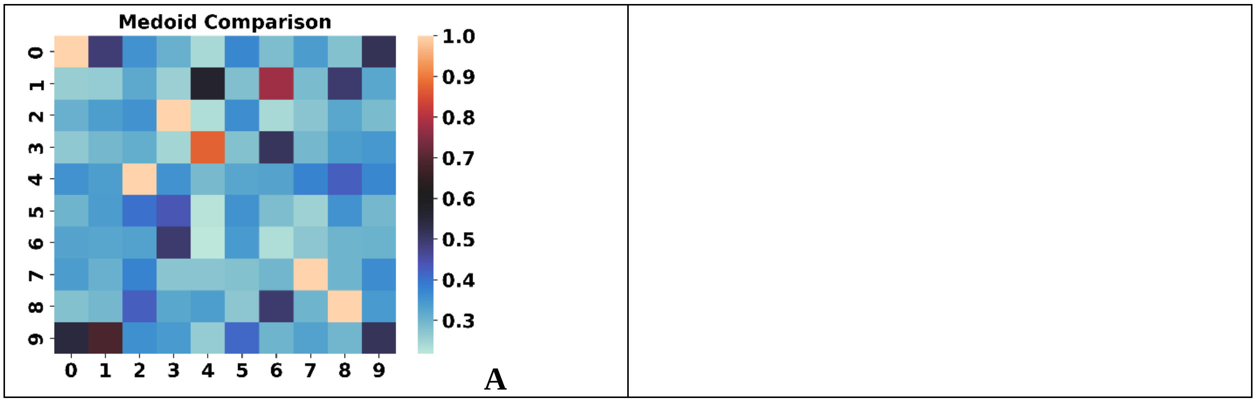

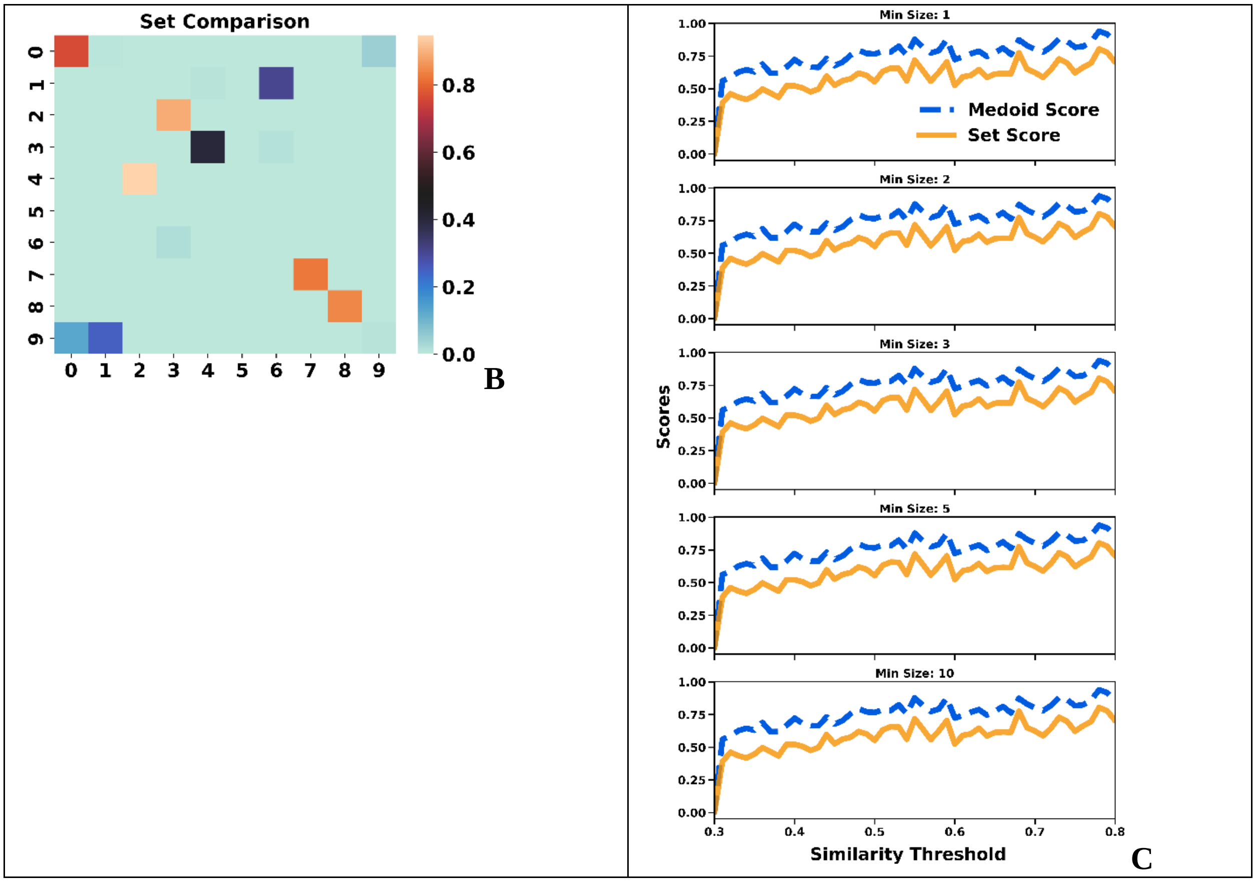


**Figure S3.22**: Comparison of the **A**: medoids, **B**: sets for the top populated clusters of the [29] ChEMBL subset (similarity threshold = 0.65, min_size = 10). **C**: Medoid and Set scores.

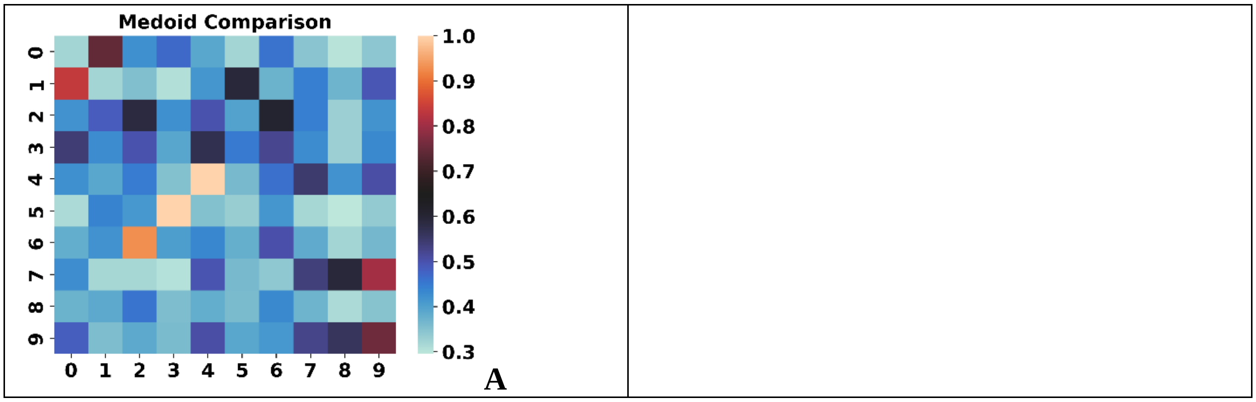

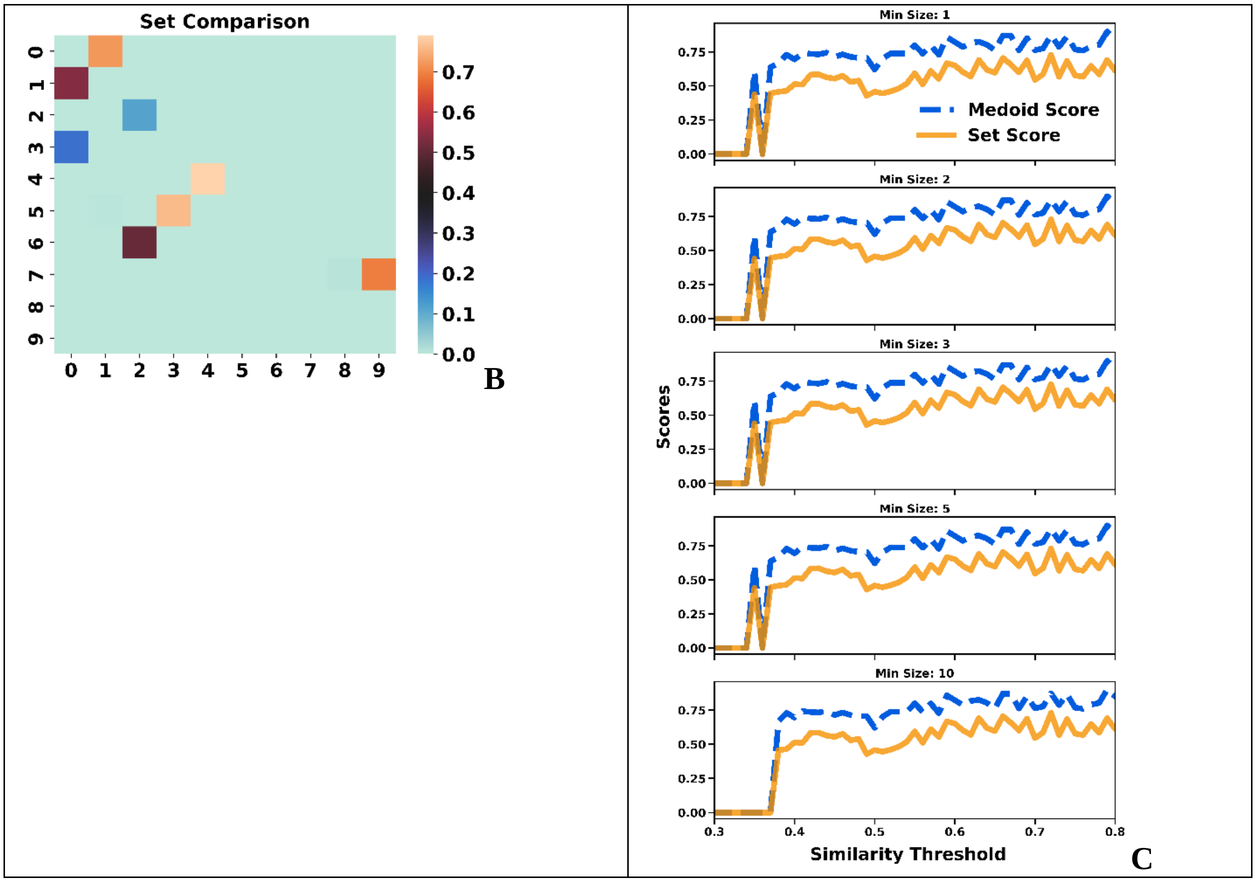


**Figure S3.23**: Comparison of the **A**: medoids, **B**: sets for the top populated clusters of the [30] ChEMBL subset (similarity threshold = 0.65, min_size = 10). **C**: Medoid and Set scores.

### S4: Clustering Performance Analysis BitBIRCH vs Taylor-Butina

The formula for the DBI used in the text is:

$$\mathrm{DBI}=\frac{1}{N_C}\sum_{j=1}^{N_C}\max_{j\neq i}\left\{\frac{\frac{1}{N_i}\sum_{v=1}^{N_i}\mathrm{T}\left(\mathbf{x}^{(i,v)},\mathbf{c}_i\right)+\frac{1}{N_j}\sum_{v=1}^{N_j}\mathrm{T}\left(\mathbf{x}^{(j,v)},\mathbf{c}_j\right)}{1-\mathrm{T}\left(\mathbf{c}_j,\mathbf{c}_i\right)}\right\} \tag{6}$$

The formula for the DI used in the text is:

$$\mathrm{DI}=\frac{\min\left\{1-i\mathrm{T}\left(\mathbf{X}^{(j)}\cup\mathbf{X}^{(i)}\right)\right\}}{\max\left\{i\mathrm{T}\left(\mathbf{X}^{(i)}\right)\right\}} \tag{7}$$

The full analysis of the quality of clustering for the 30 ChEMBL subsets is presented below.

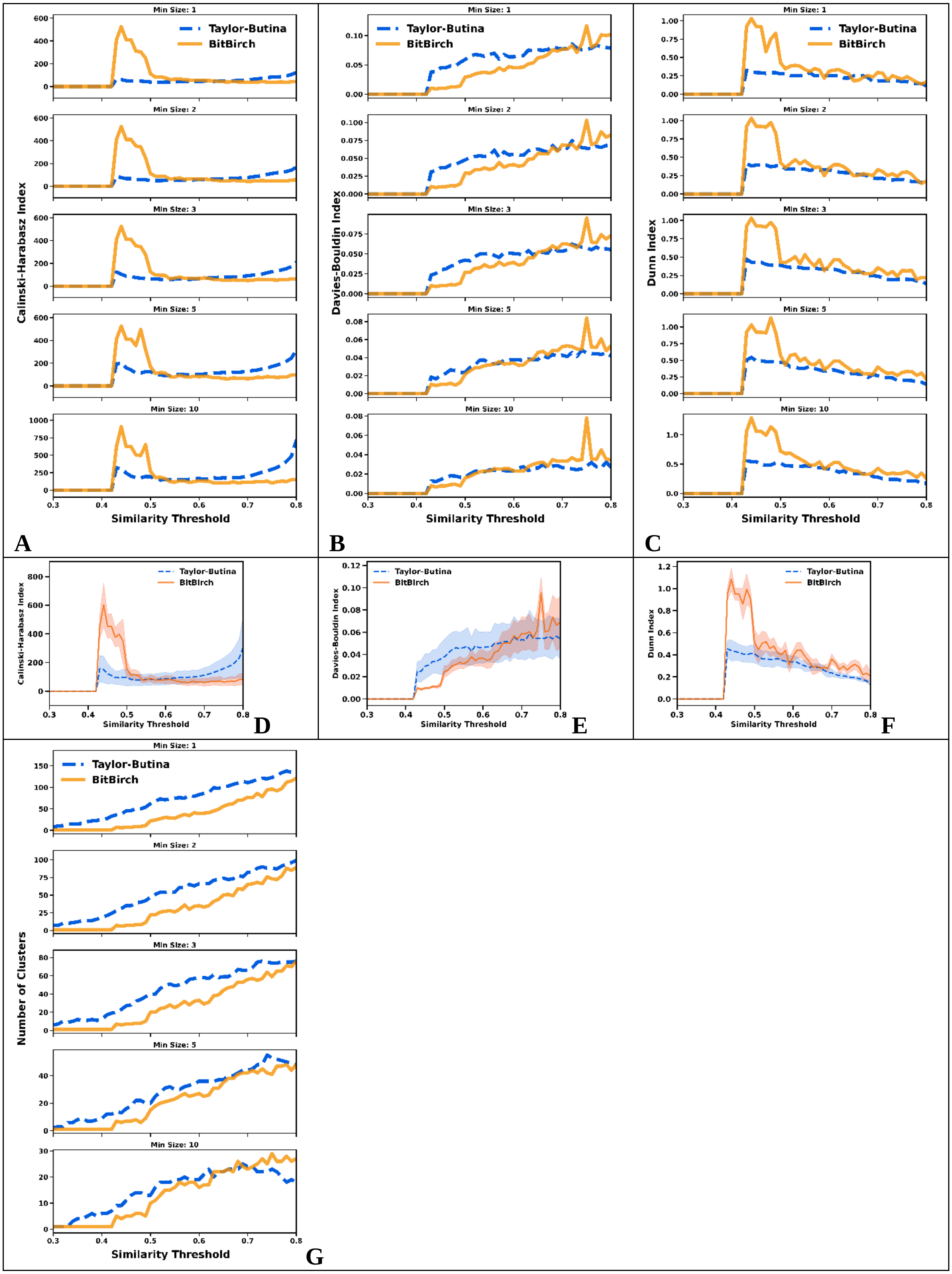

Min Size: 1
Min Size: 2
Min Size: 3
Min Size: 5
Min Size: 10
Taylor-Butina
BitBirch
Calinski-Harabasz Index
Davies-Bouldin Index
Dunn Index
Number of Clusters
Similarity Threshold
A
B
C
D
E
F
G

**Figure S4.1**: **A**: CHI, **B**: DBI, **C**: DI analysis for min_size = 1, 2, 3, 5, 10 for the [1] ChEMBL subset. Average **D**: CHI, **E**: DBI, **F**: DI values over the min_size variable. **G**: Number of clusters. BitBIRCH (orange continuous line), Taylor-Butina (blue dashed line).

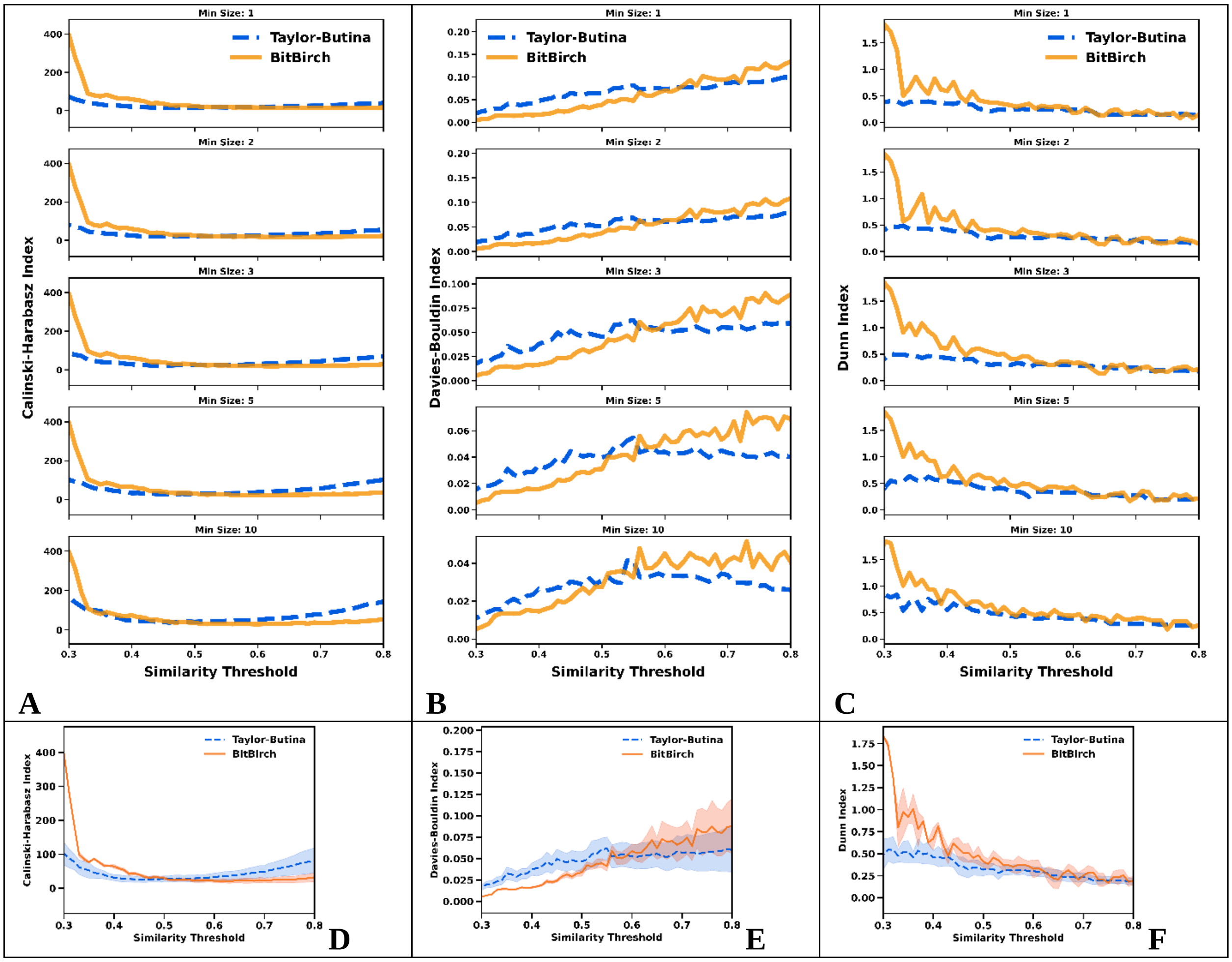

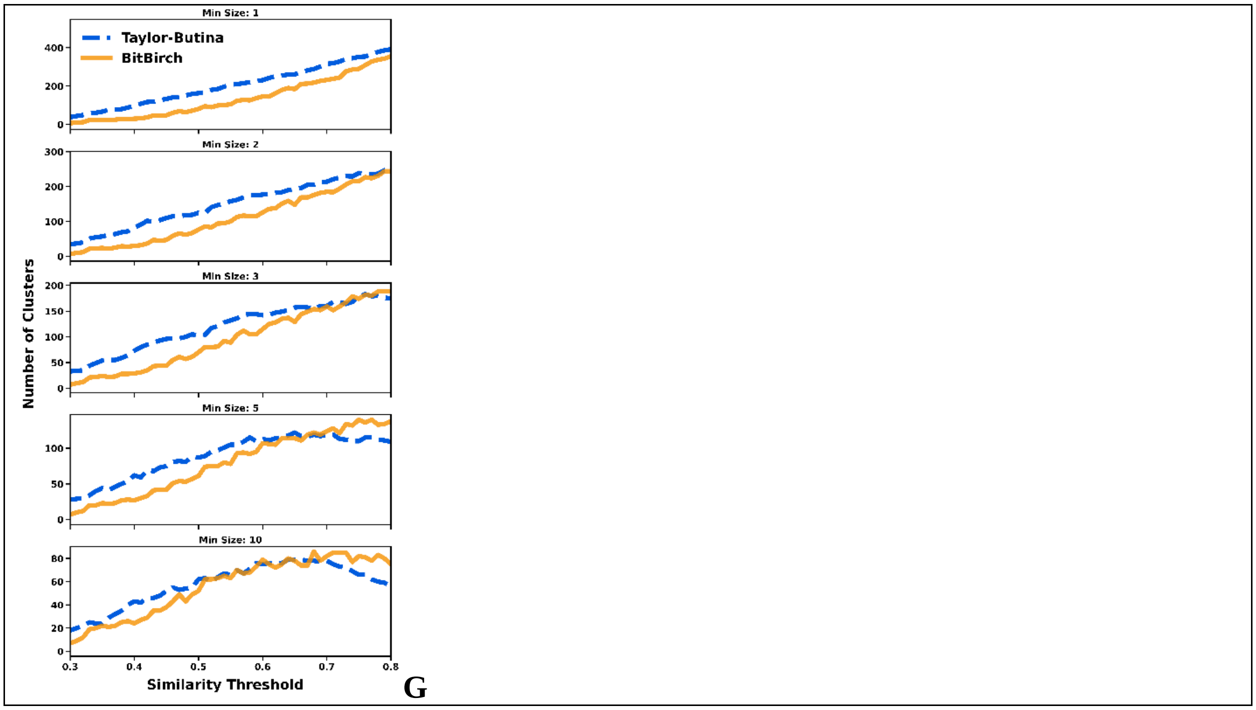


**Figure S4.2**: **A**: CHI, **B**: DBI, **C**: DI analysis for min_size = 1, 2, 3, 5, 10 for the [2] ChEMBL subset. Average **D**: CHI, **E**: DBI, **F**: DI values over the min_size variable. **G**: Number of clusters. BitBIRCH (orange continuous line), Taylor-Butina (blue dashed line).

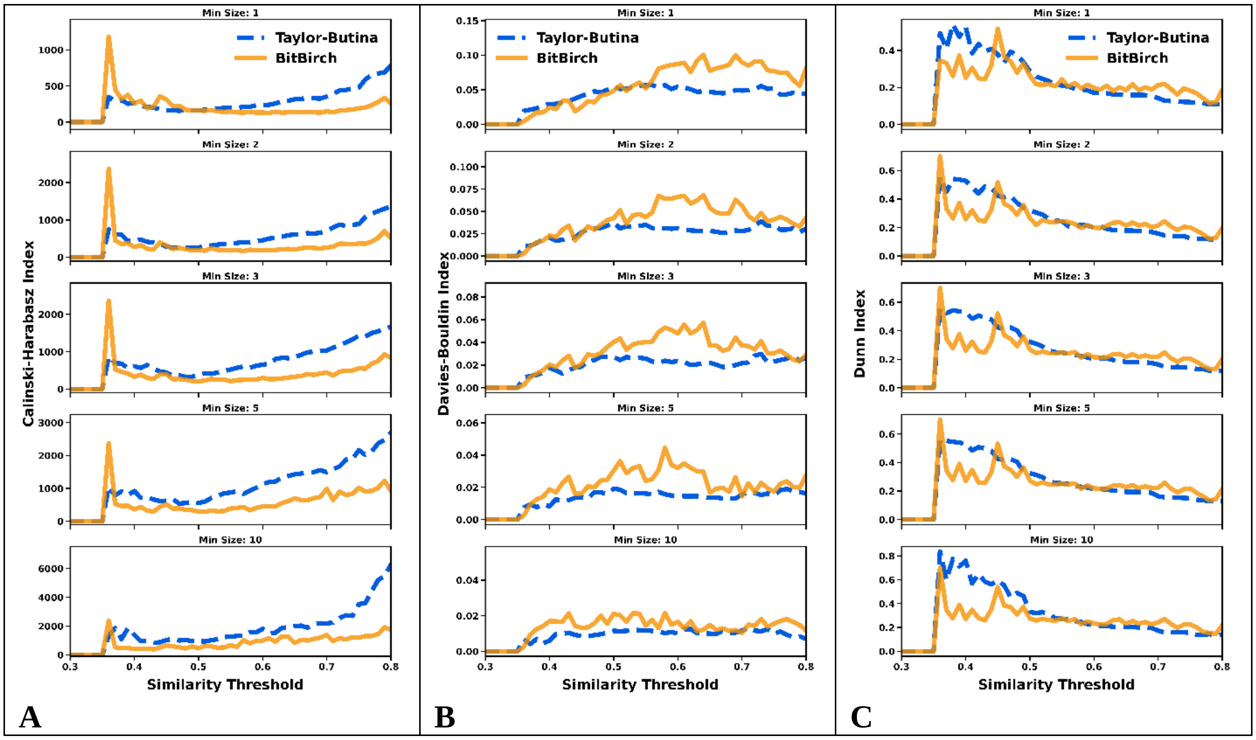

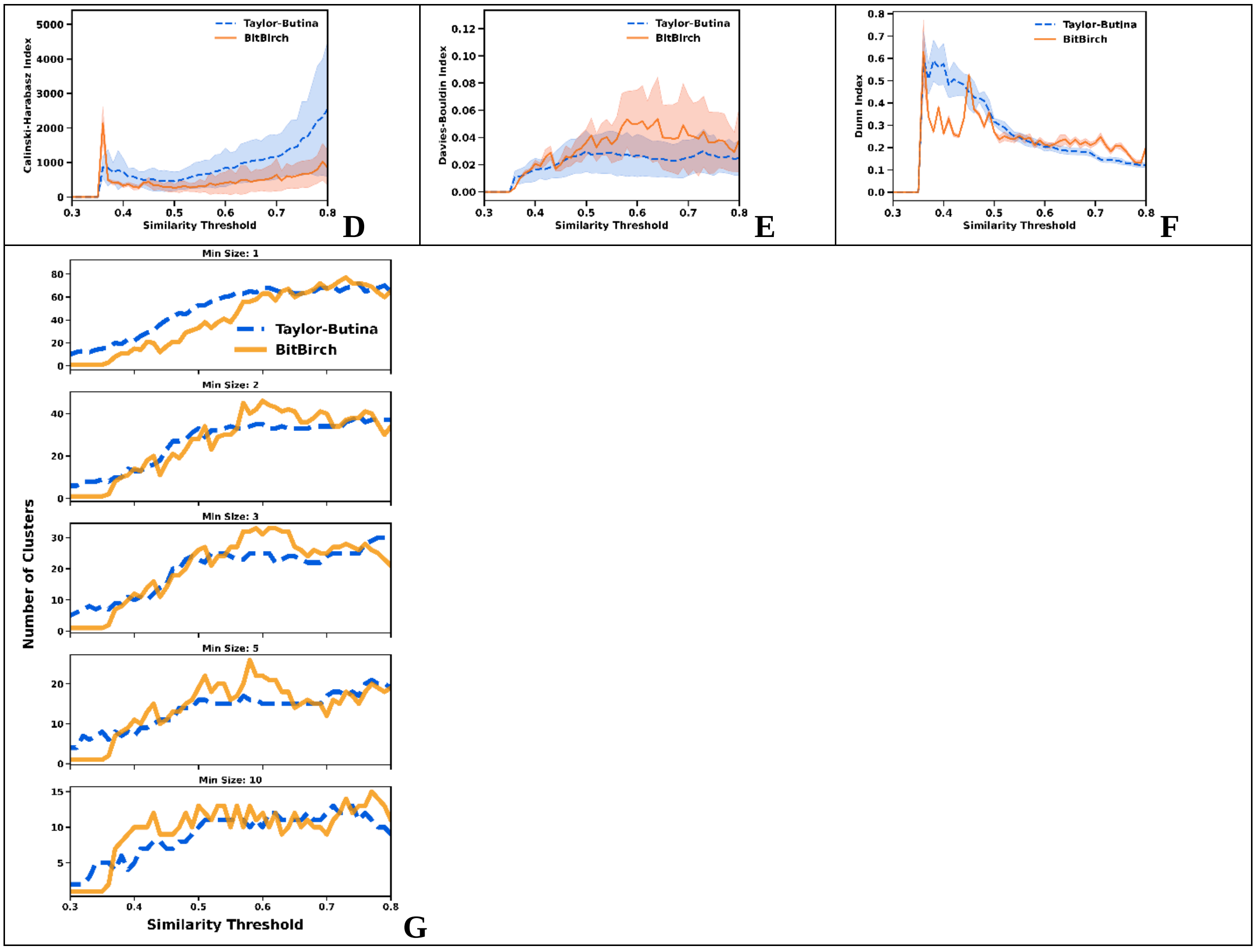


**Figure S4.3**: **A**: CHI, **B**: DBI, **C**: DI analysis for min_size = 1, 2, 3, 5, 10 for the [3] ChEMBL subset. Average **D**: CHI, **E**: DBI, **F**: DI values over the min_size variable. **G**: Number of clusters. BitBIRCH (orange continuous line), Taylor-Butina (blue dashed line).

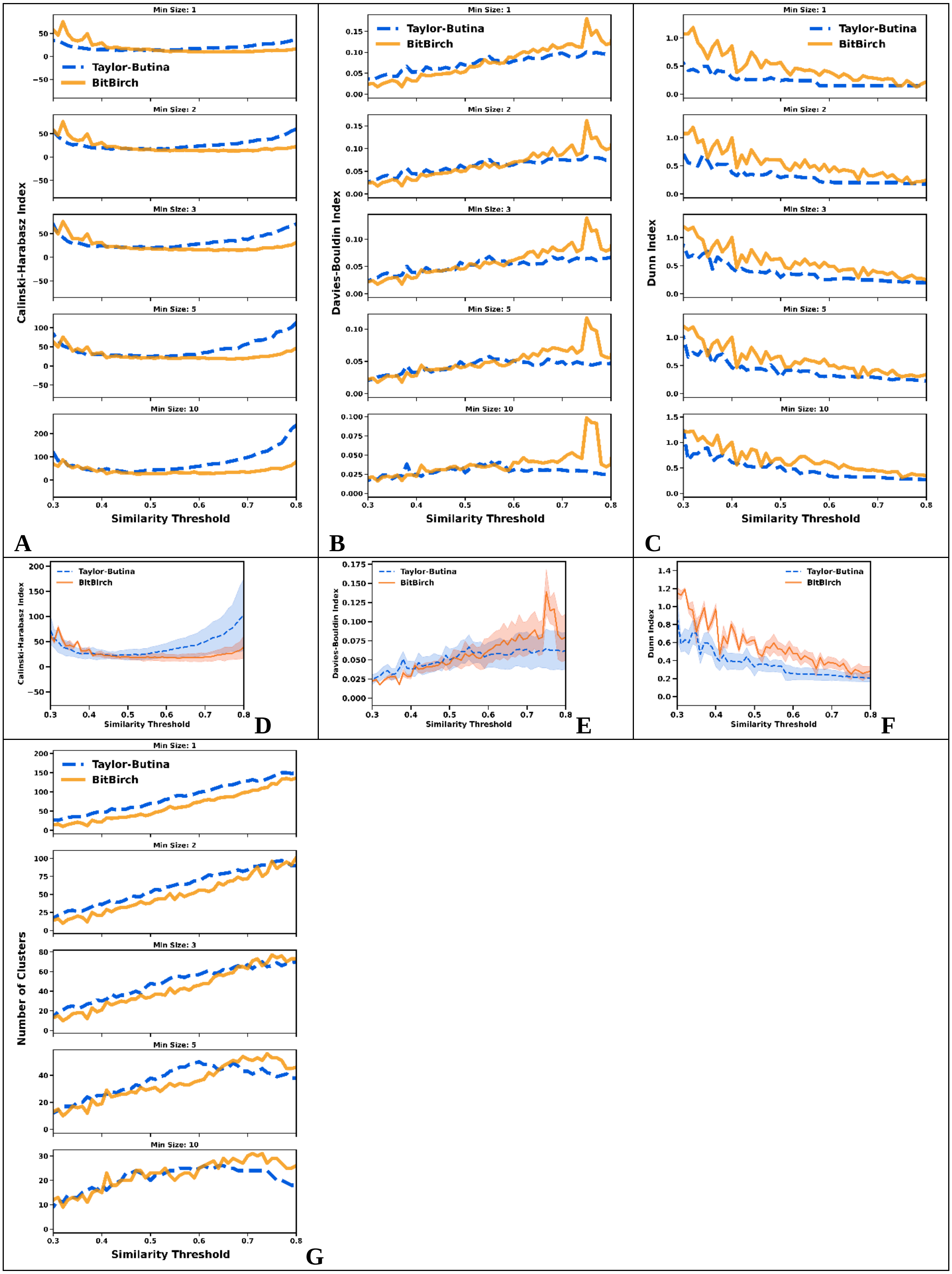

Min Size: 1
Min Size: 2
Min Size: 3
Min Size: 5
Min Size: 10
Taylor-Butina
BitBirch
Calinski-Harabasz Index
Davies-Bouldin Index
Dunn Index
Number of Clusters
Similarity Threshold
A
B
C
D
E
F
G

**Figure S4.4**: **A**: CHI, **B**: DBI, **C**: DI analysis for min_size = 1, 2, 3, 5, 10 for the [4] ChEMBL subset. Average **D**: CHI, **E**: DBI, **F**: DI values over the min_size variable. **G**: Number of clusters. BitBIRCH (orange continuous line), Taylor-Butina (blue dashed line).

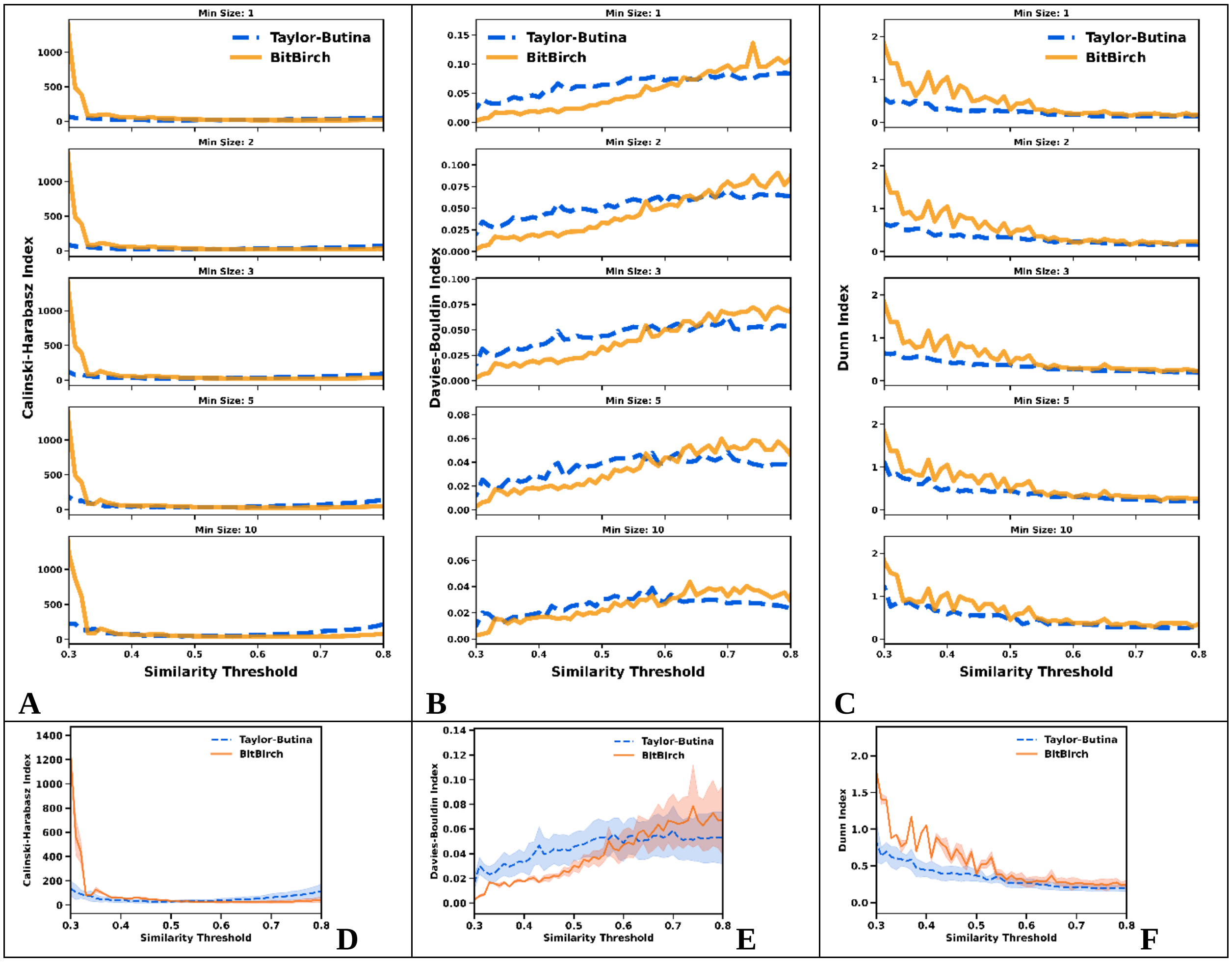

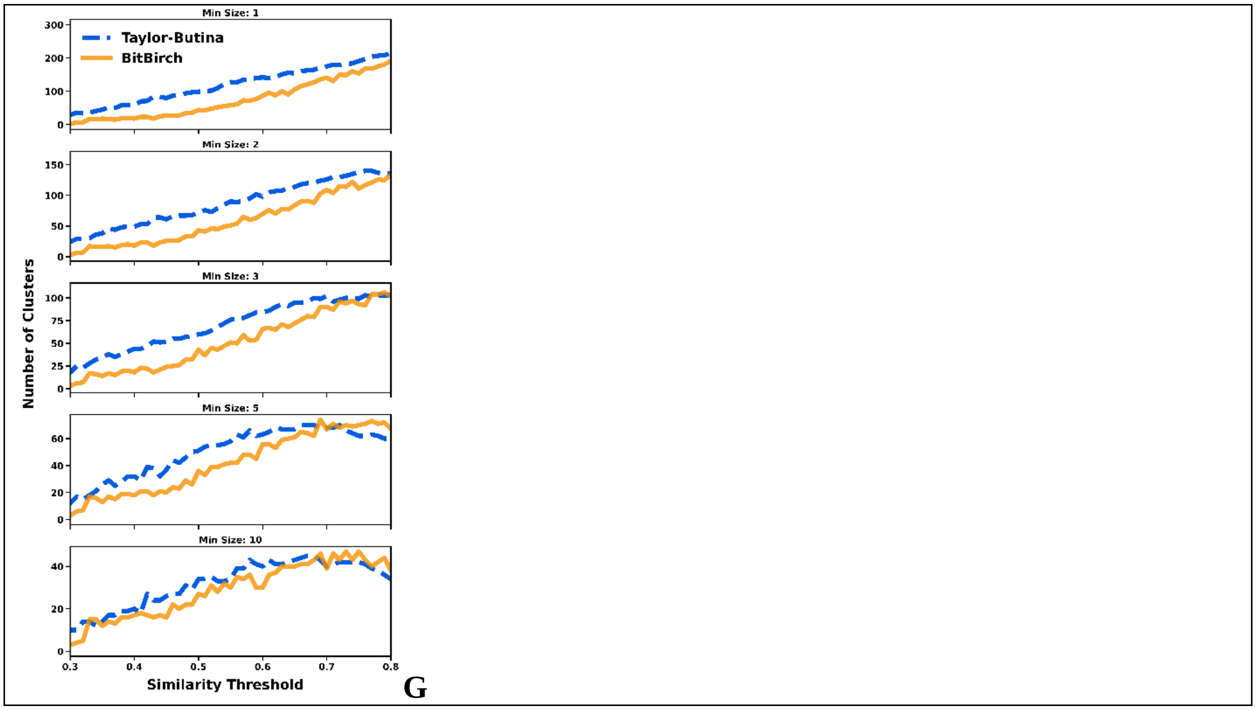


**Figure S4.5**: **A**: CHI, **B**: DBI, **C**: DI analysis for min_size = 1, 2, 3, 5, 10 for the [5] ChEMBL subset. Average **D**: CHI, **E**: DBI, **F**: DI values over the min_size variable. **G**: Number of clusters. BitBIRCH (orange continuous line), Taylor-Butina (blue dashed line).

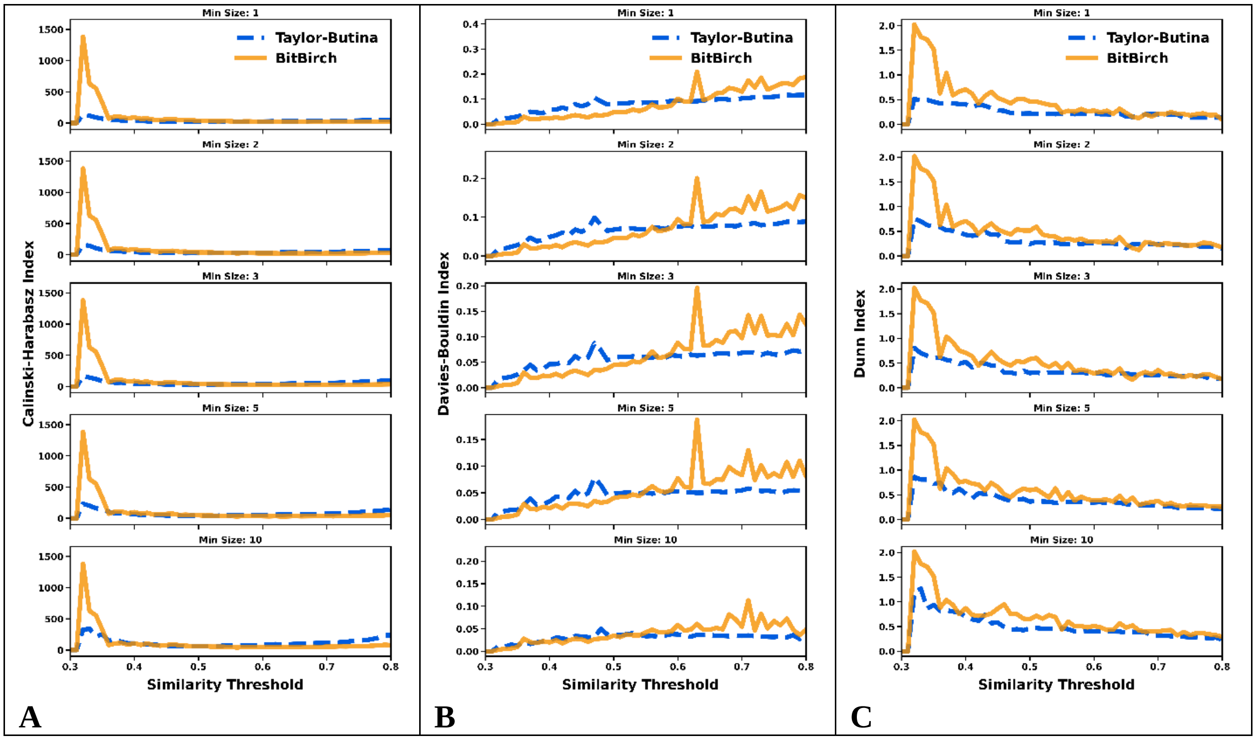

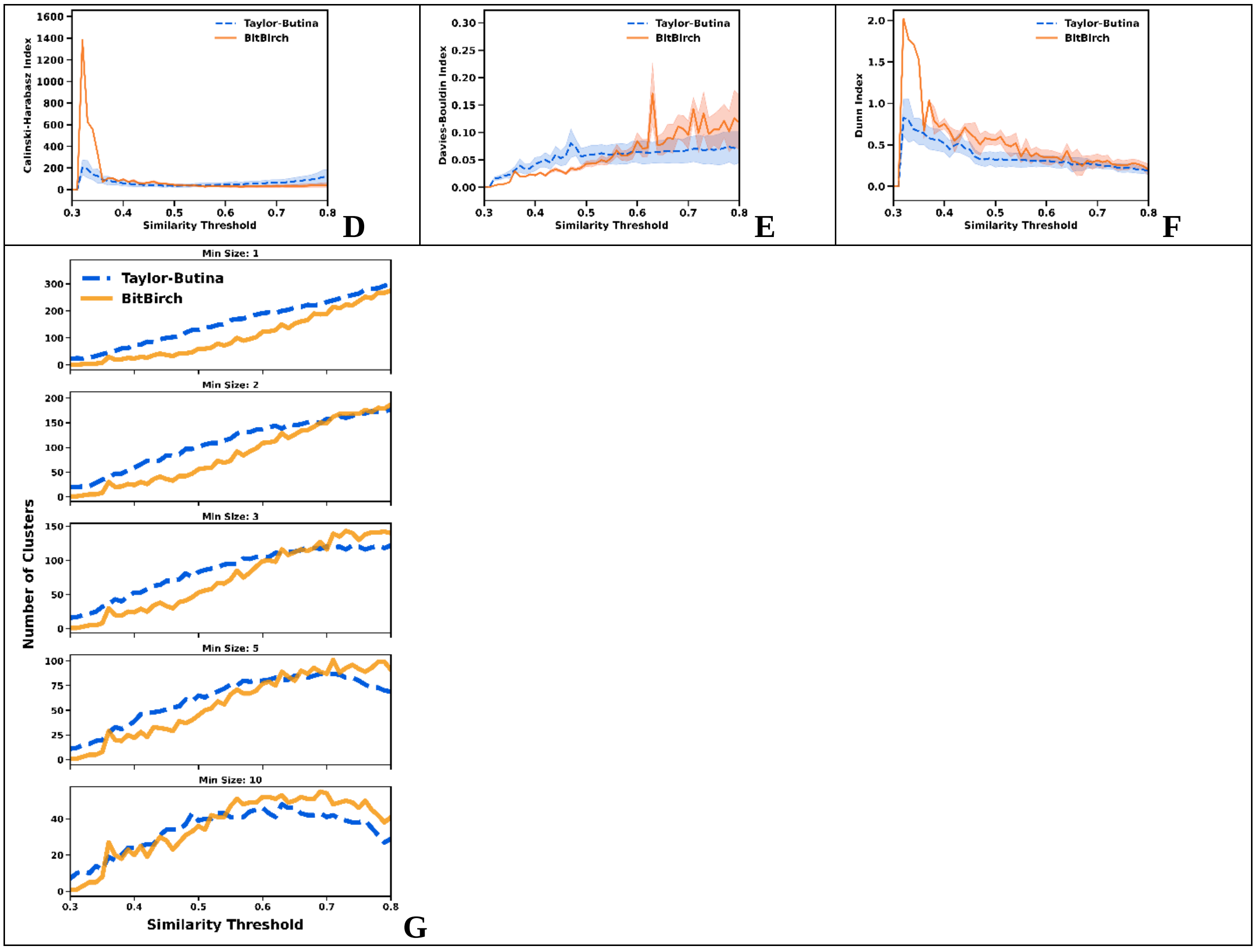


**Figure S4.6**: **A**: CHI, **B**: DBI, **C**: DI analysis for min_size = 1, 2, 3, 5, 10 for the [6] ChEMBL subset. Average **D**: CHI, **E**: DBI, **F**: DI values over the min_size variable. **G**: Number of clusters. BitBIRCH (orange continuous line), Taylor-Butina (blue dashed line).

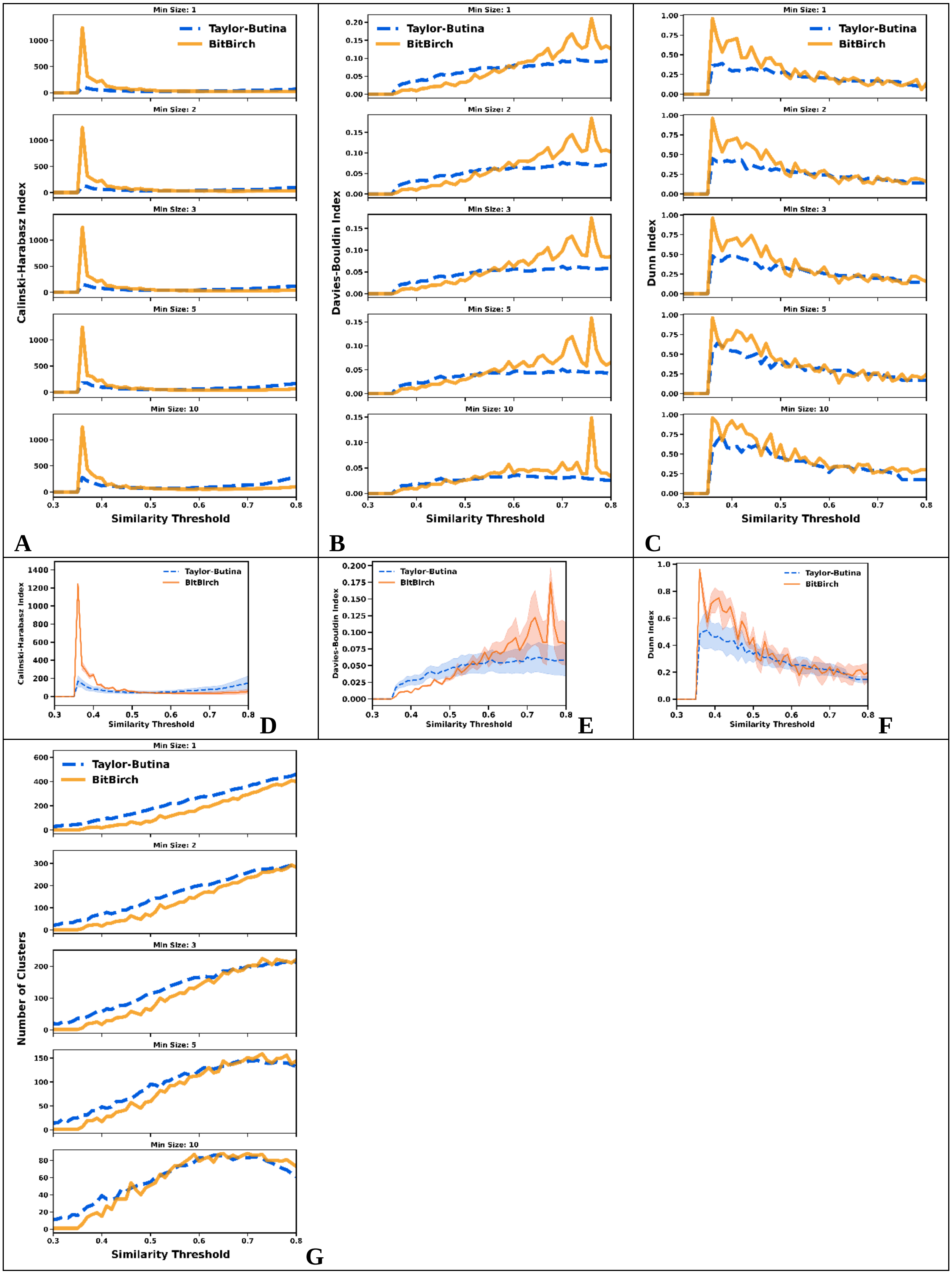

Min Size: 1
Min Size: 2
Min Size: 3
Min Size: 5
Min Size: 10
Taylor-Butina
BitBirch
Calinski-Harabasz Index
Davies-Bouldin Index
Dunn Index
Number of Clusters
Similarity Threshold
A
B
C
D
E
F
G

**Figure S4.7**: **A**: CHI, **B**: DBI, **C**: DI analysis for min_size = 1, 2, 3, 5, 10 for the [7] ChEMBL subset. Average **D**: CHI, **E**: DBI, **F**: DI values over the min_size variable. **G**: Number of clusters. BitBIRCH (orange continuous line), Taylor-Butina (blue dashed line).

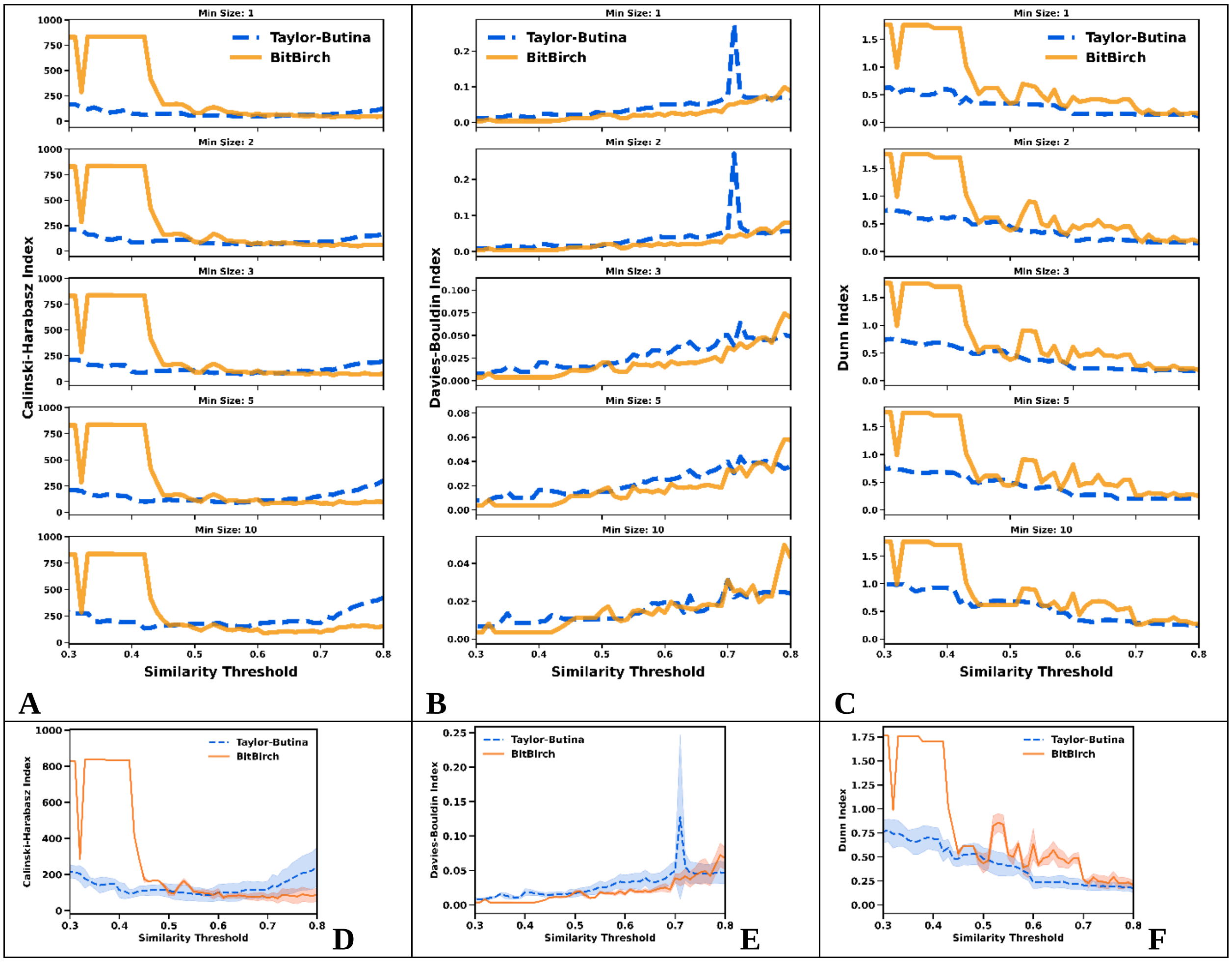

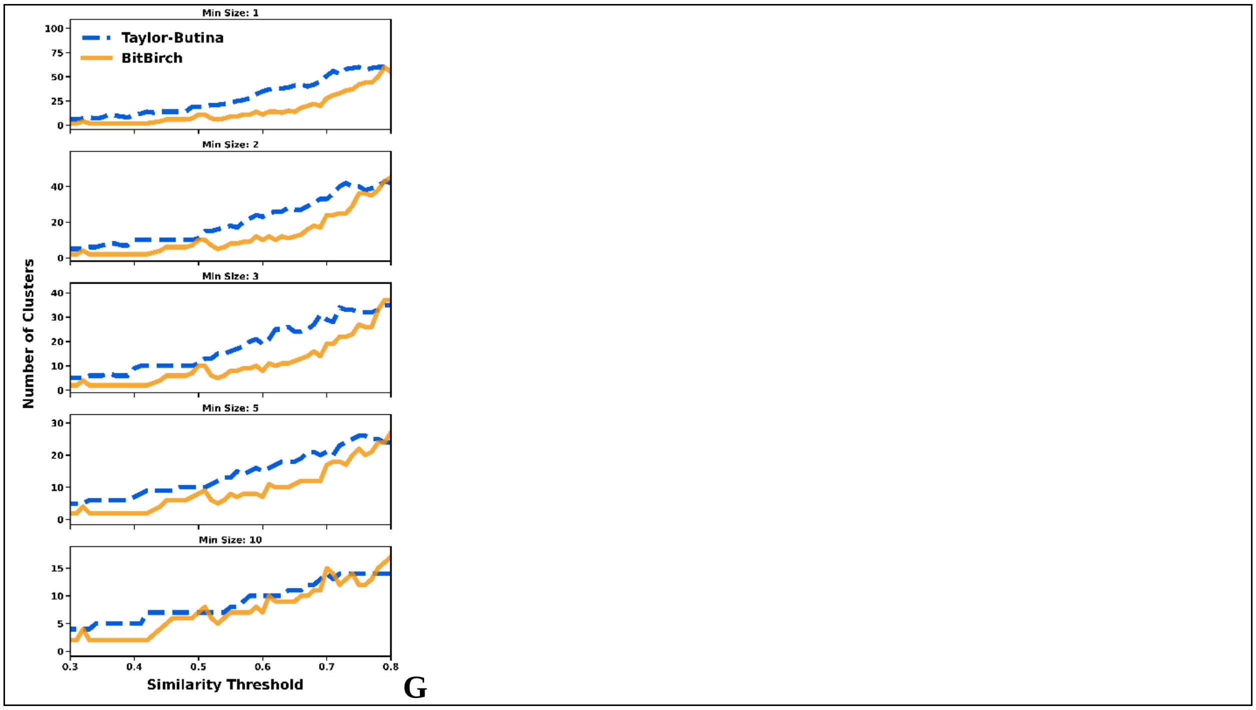


**Figure S4.8**: **A**: CHI, **B**: DBI, **C**: DI analysis for min_size = 1, 2, 3, 5, 10 for the [8] ChEMBL subset. Average **D**: CHI, **E**: DBI, **F**: DI values over the min_size variable. **G**: Number of clusters. BitBIRCH (orange continuous line), Taylor-Butina (blue dashed line).

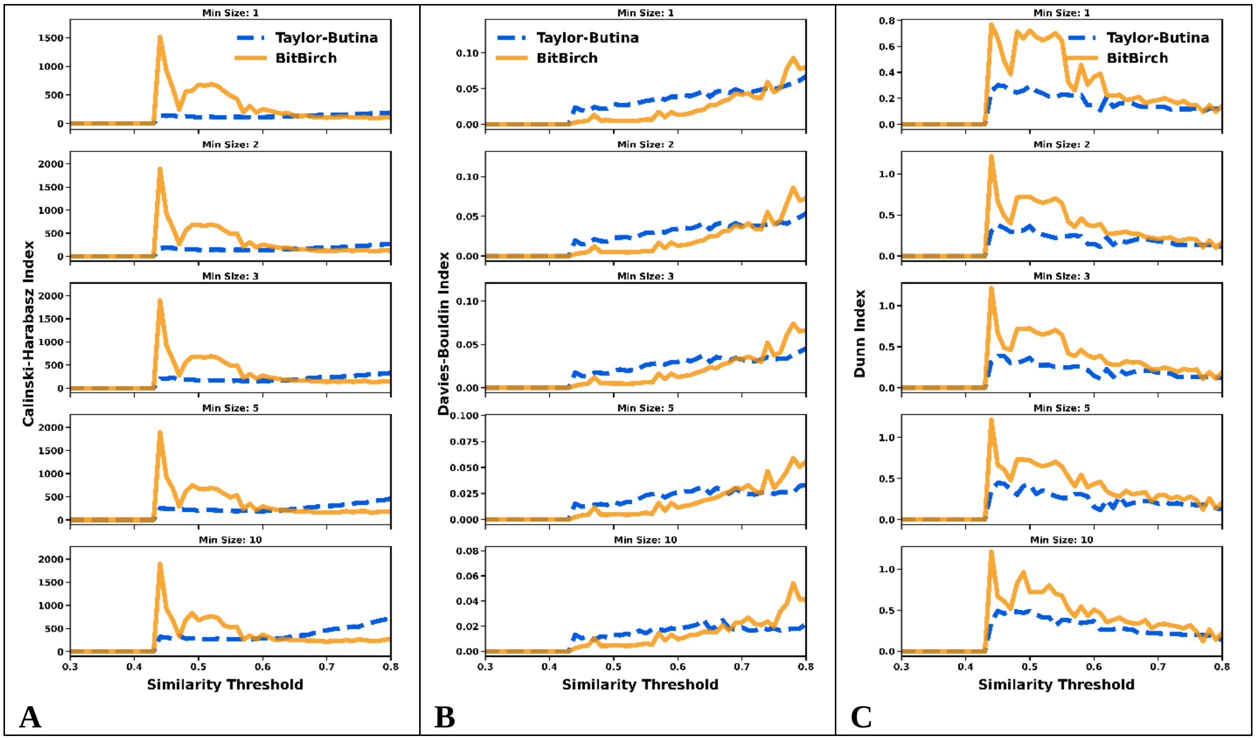

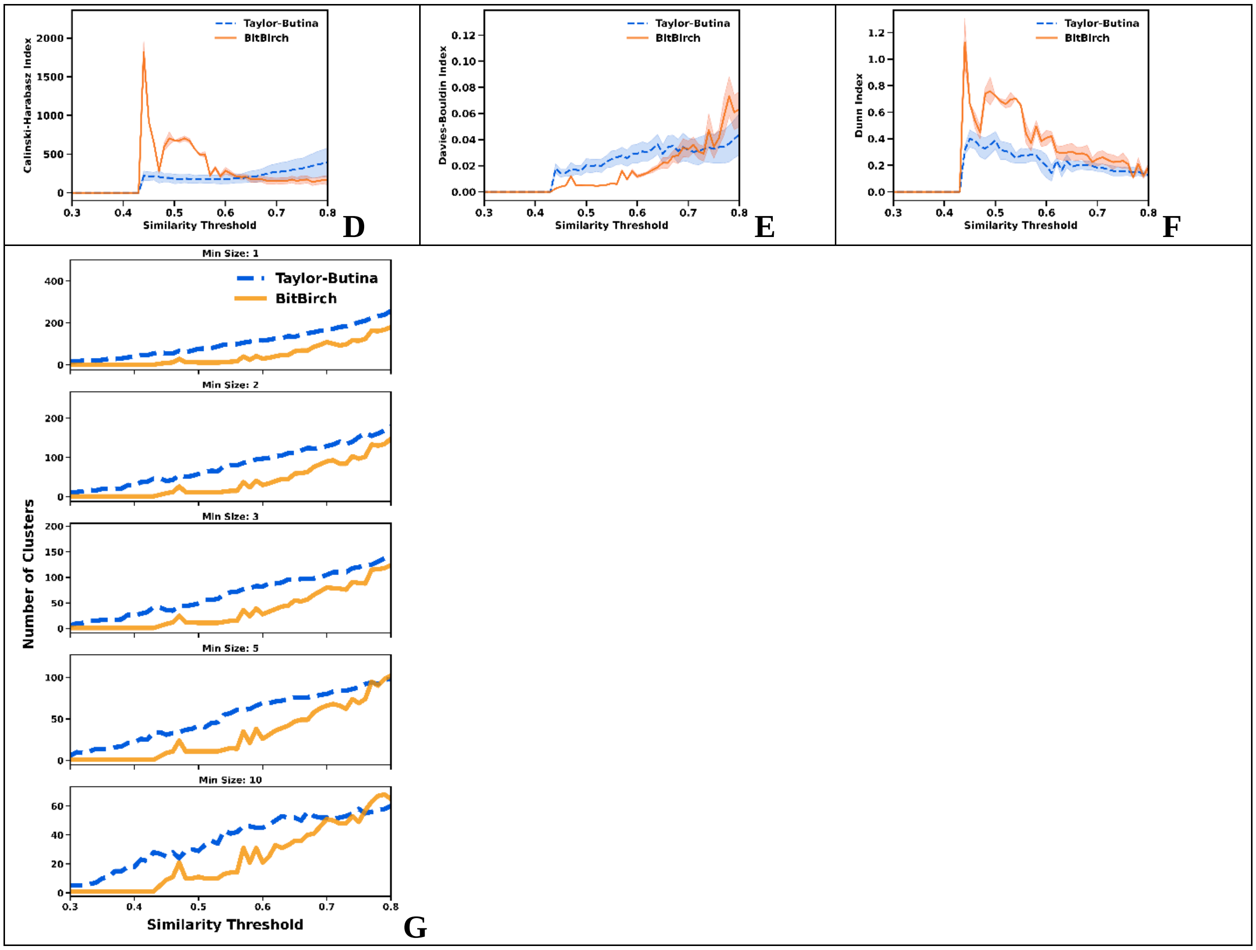


**Figure S4.9**: **A**: CHI, **B**: DBI, **C**: DI analysis for min_size = 1, 2, 3, 5, 10 for the [9] ChEMBL subset. Average **D**: CHI, **E**: DBI, **F**: DI values over the min_size variable. **G**: Number of clusters. BitBIRCH (orange continuous line), Taylor-Butina (blue dashed line).

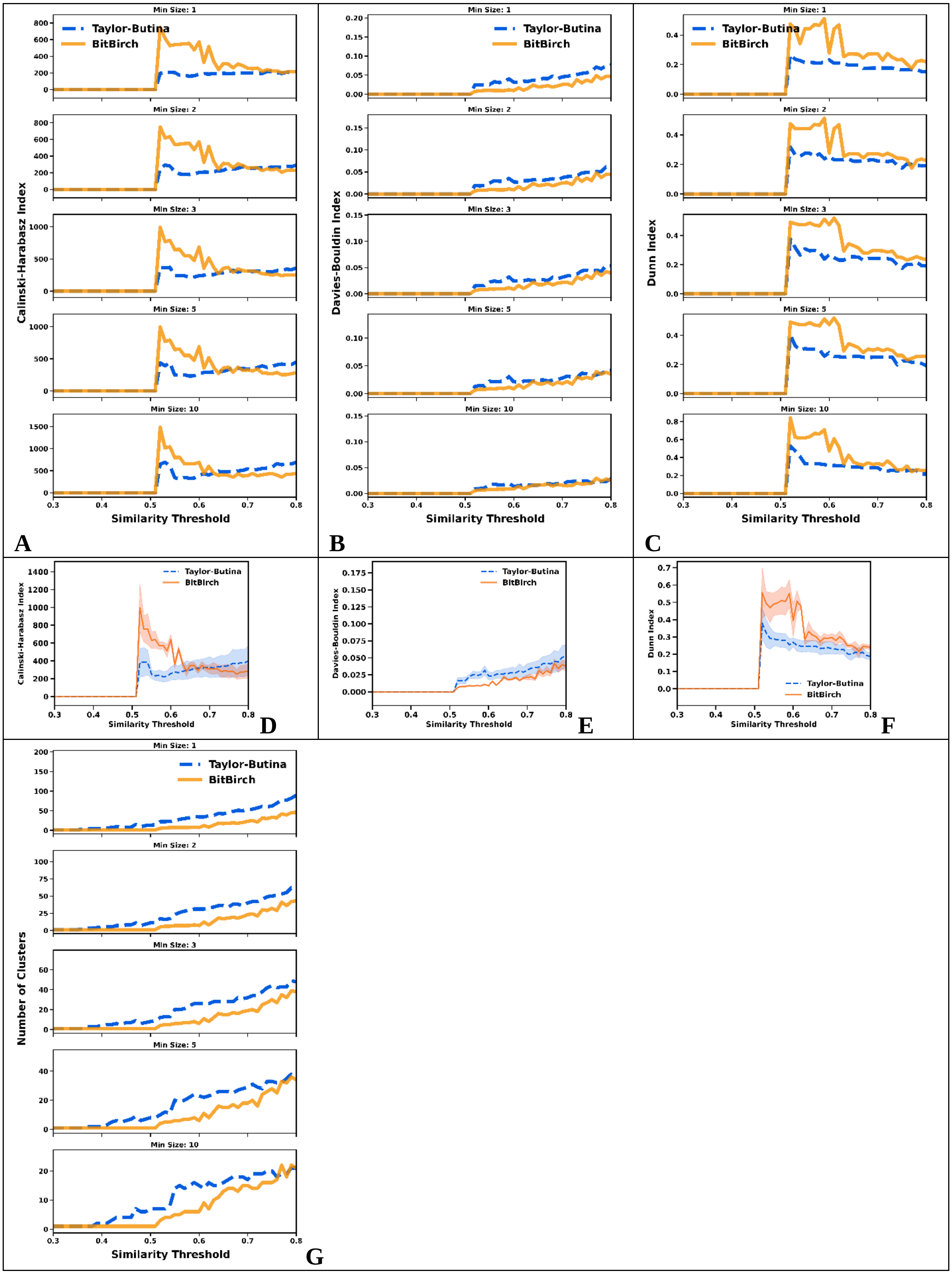
Min Size: 1
Min Size: 2
Min Size: 3
Min Size: 5
Min Size: 10
Taylor-Butina
BitBirch
Calinski-Harabasz Index
Davies-Bouldin Index
Dunn Index
Number of Clusters
Similarity Threshold
A
B
C
D
E
F
G

**Figure S4.10**: **A**: CHI, **B**: DBI, **C**: DI analysis for min_size = 1, 2, 3, 5, 10 for the [10] ChEMBL subset. Average **D**: CHI, **E**: DBI, **F**: DI values over the min_size variable. **G**: Number of clusters. BitBIRCH (orange continuous line), Taylor-Butina (blue dashed line).

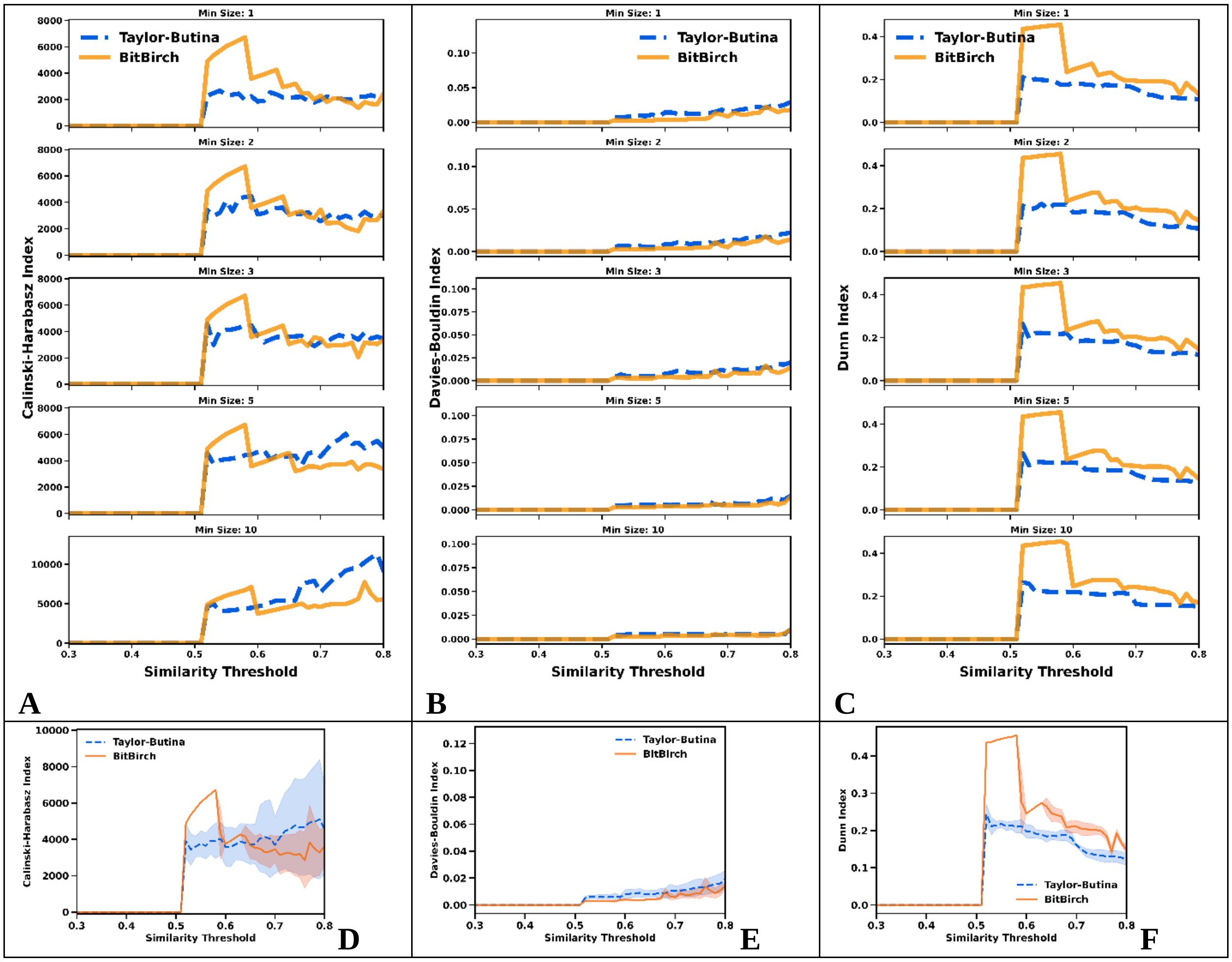

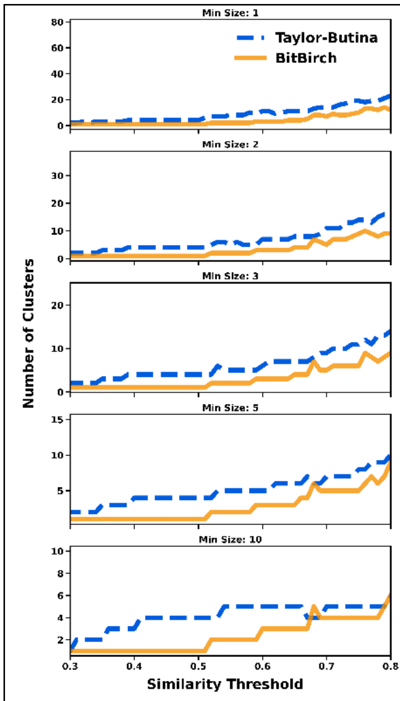


**G**

**Figure S4.11**: **A**: CHI, **B**: DBI, **C**: DI analysis for min_size = 1, 2, 3, 5, 10 for the [11] ChEMBL subset. Average **D**: CHI, **E**: DBI, **F**: DI values over the min_size variable. **G**: Number of clusters. BitBIRCH (orange continuous line), Taylor-Butina (blue dashed line).

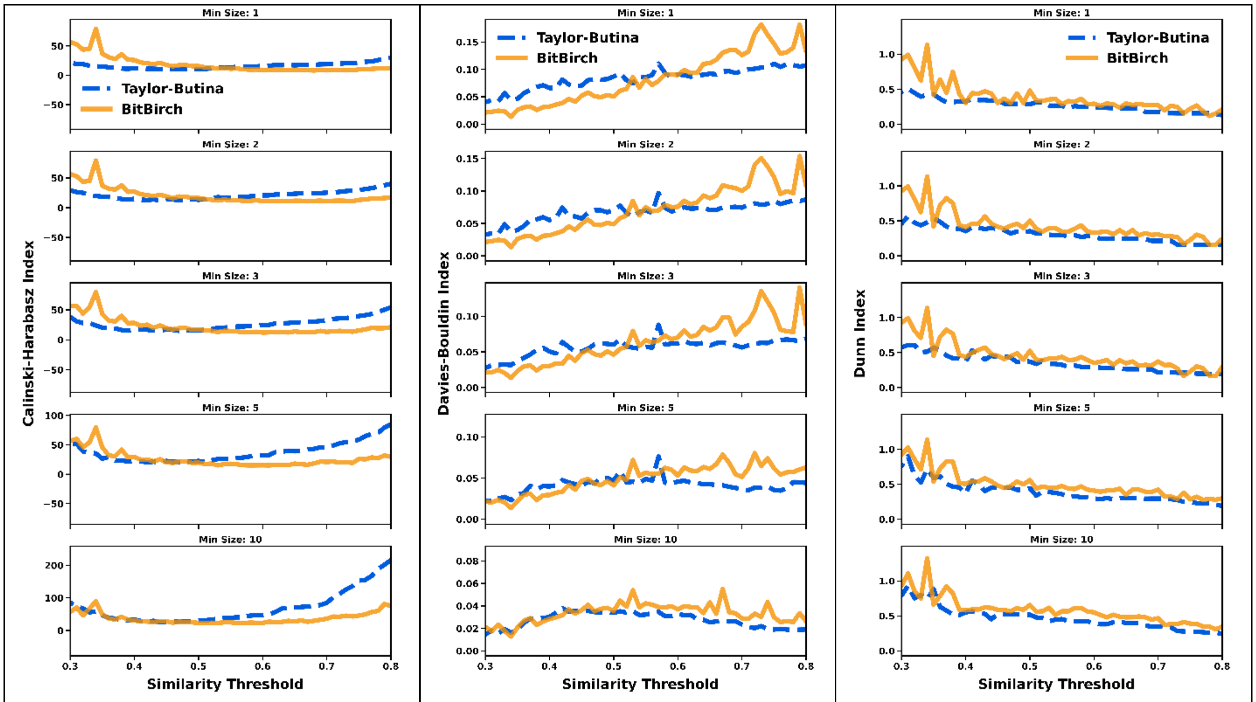


**A** **B** **C**

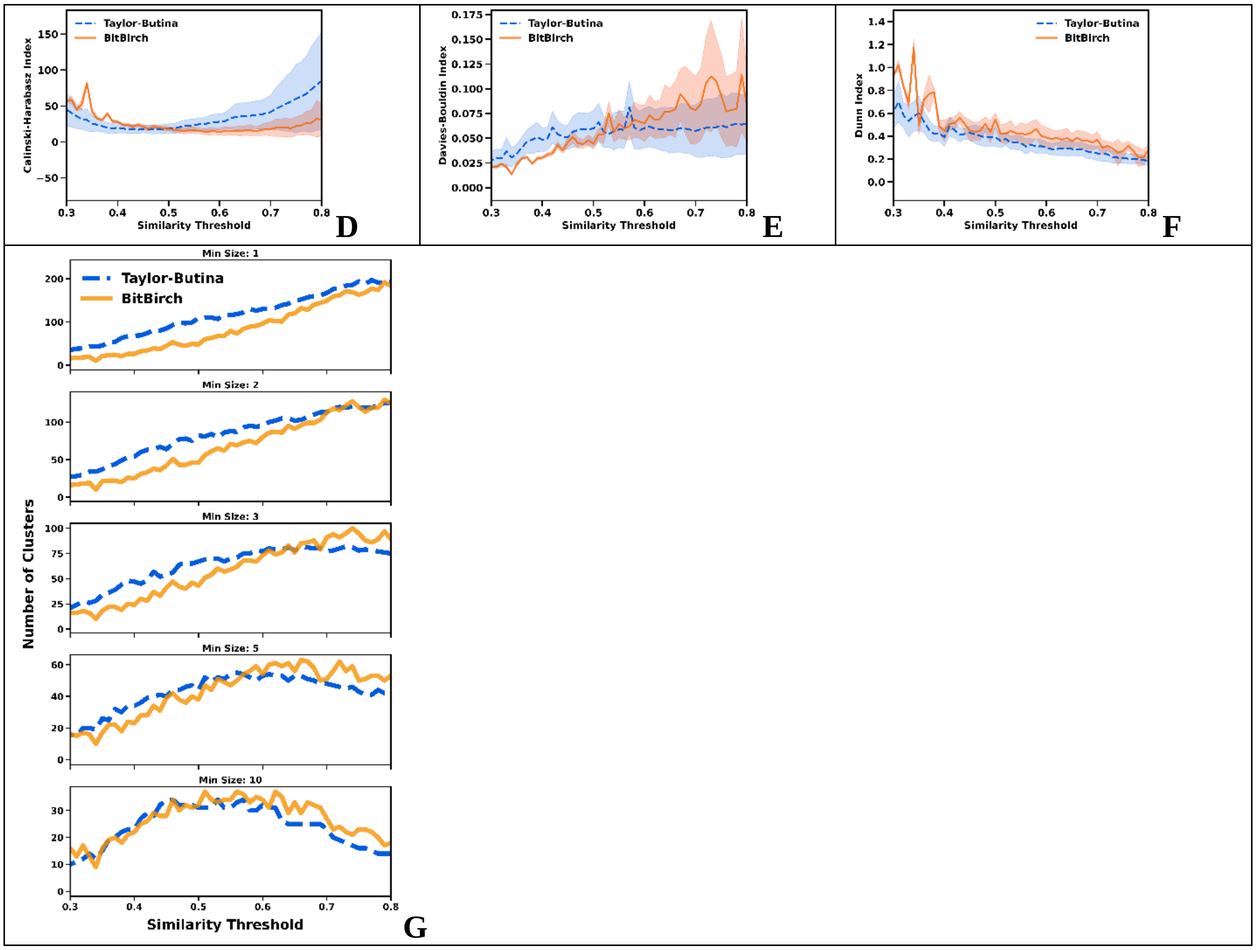


**Figure S4.12**: **A**: CHI, **B**: DBI, **C**: DI analysis for min_size = 1, 2, 3, 5, 10 for the [12] ChEMBL subset. Average **D**: CHI, **E**: DBI, **F**: DI values over the min_size variable. **G**: Number of clusters. BitBIRCH (orange continuous line), Taylor-Butina (blue dashed line).

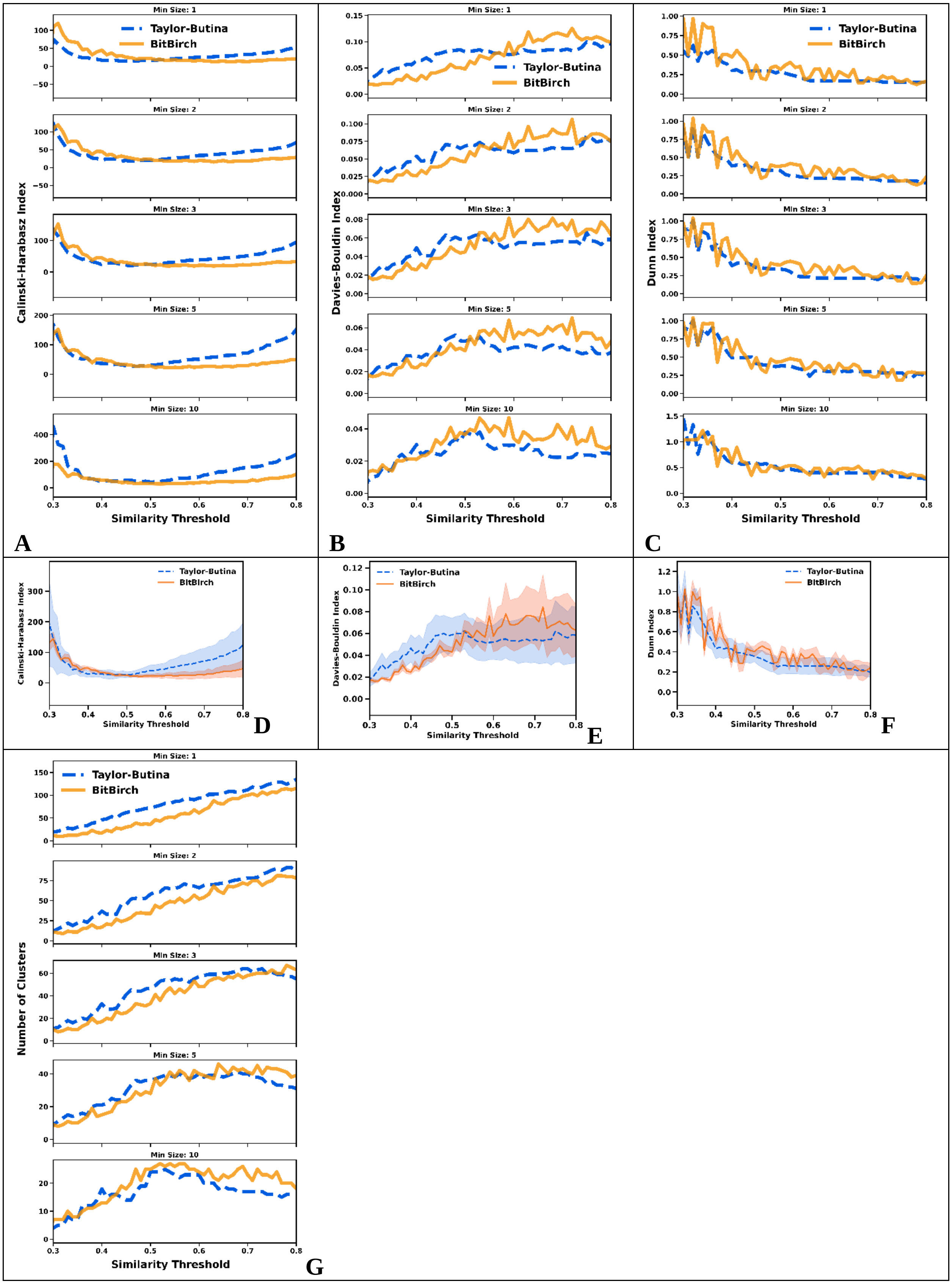
Min Size: 1
Min Size: 2
Min Size: 3
Min Size: 5
Min Size: 10
Taylor-Butina
BitBirch
Calinski-Harabasz Index
Davies-Bouldin Index
Dunn Index
Number of Clusters
Similarity Threshold
A
B
C
D
E
F
G

**Figure S4.13**: **A**: CHI, **B**: DBI, **C**: DI analysis for min_size = 1, 2, 3, 5, 10 for the [13] ChEMBL subset. Average **D**: CHI, **E**: DBI, **F**: DI values over the min_size variable. **G**: Number of clusters. BitBIRCH (orange continuous line), Taylor-Butina (blue dashed line).

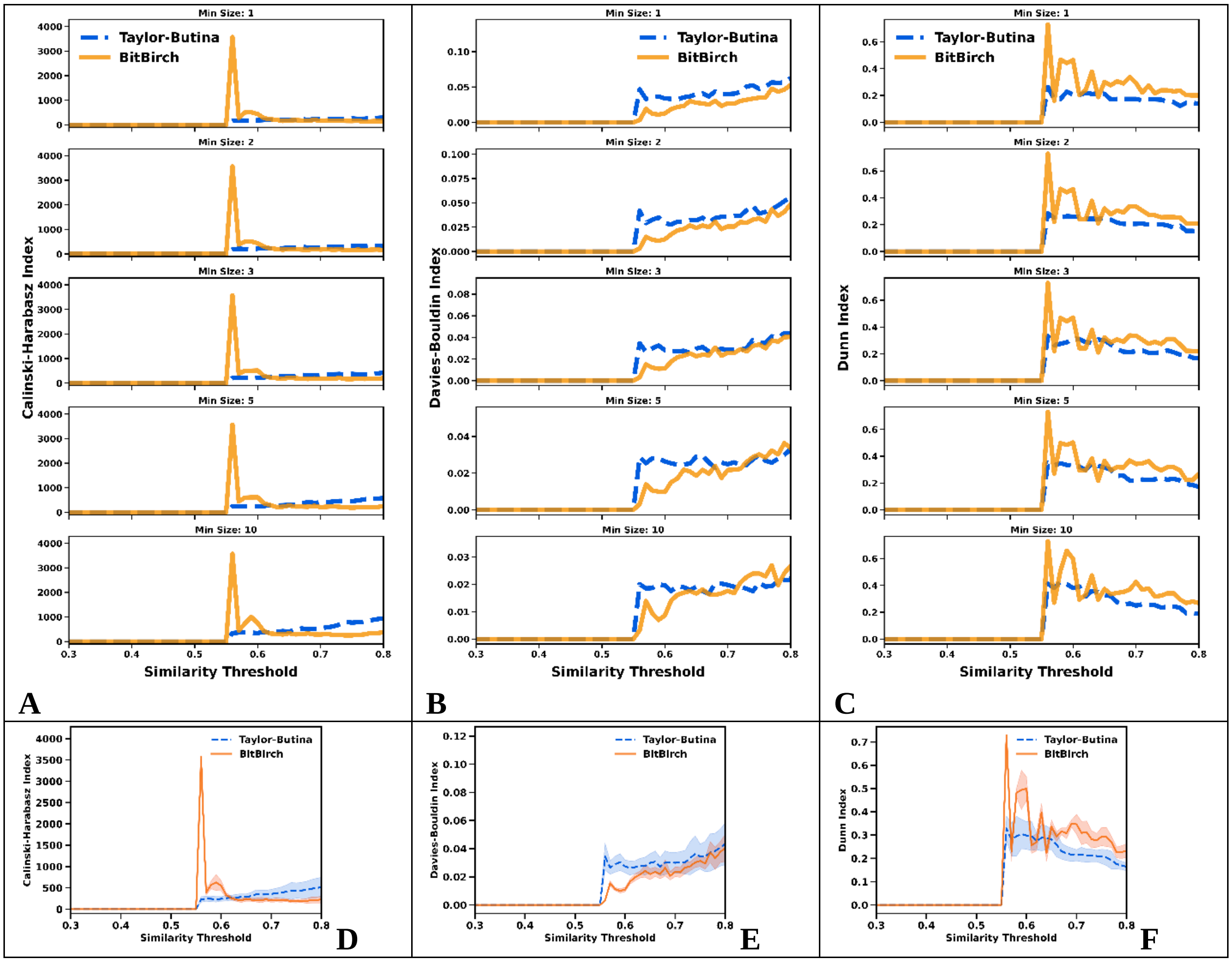

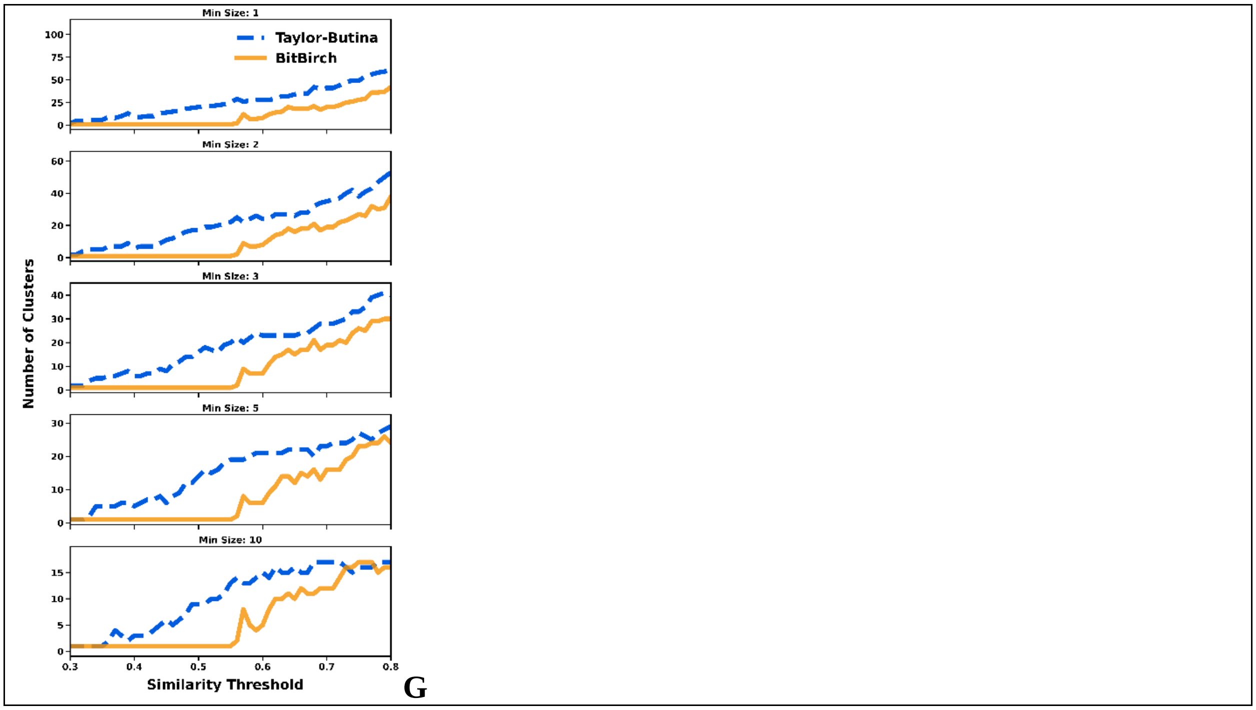


**Figure S4.14**: **A**: CHI, **B**: DBI, **C**: DI analysis for min_size = 1, 2, 3, 5, 10 for the [14] ChEMBL subset. Average **D**: CHI, **E**: DBI, **F**: DI values over the min_size variable. **G**: Number of clusters. BitBIRCH (orange continuous line), Taylor-Butina (blue dashed line).

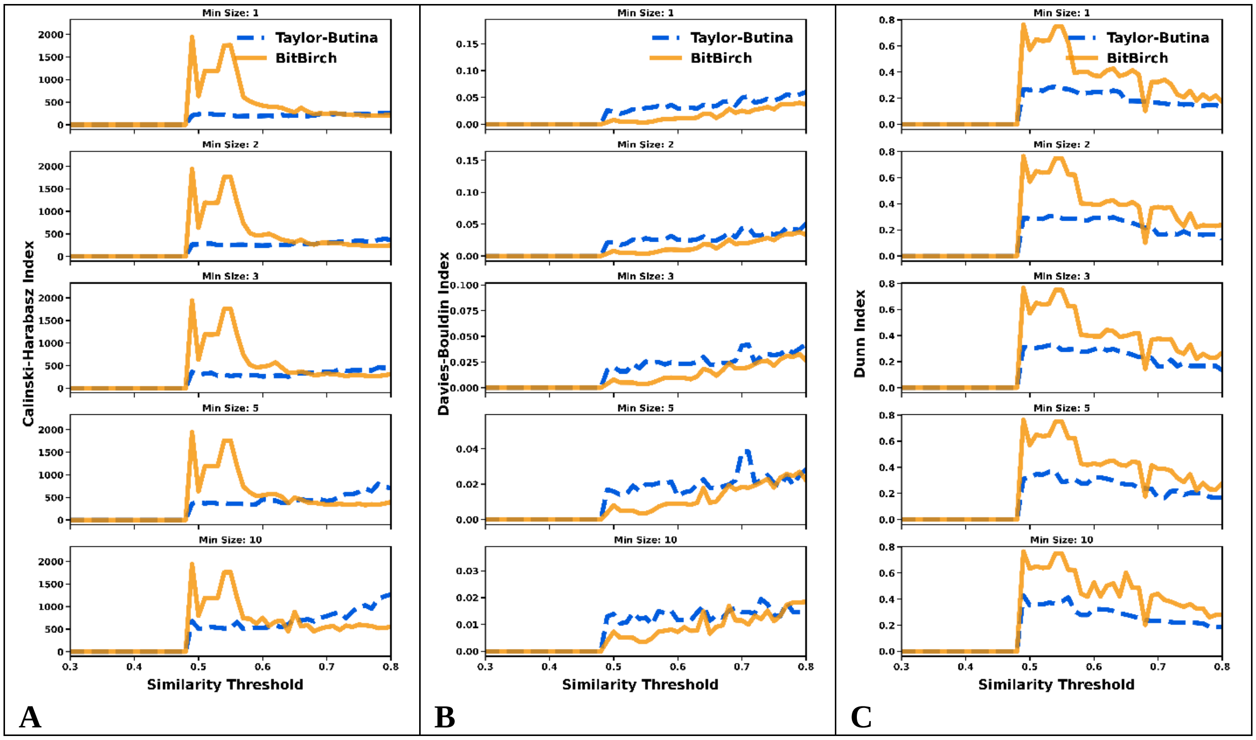

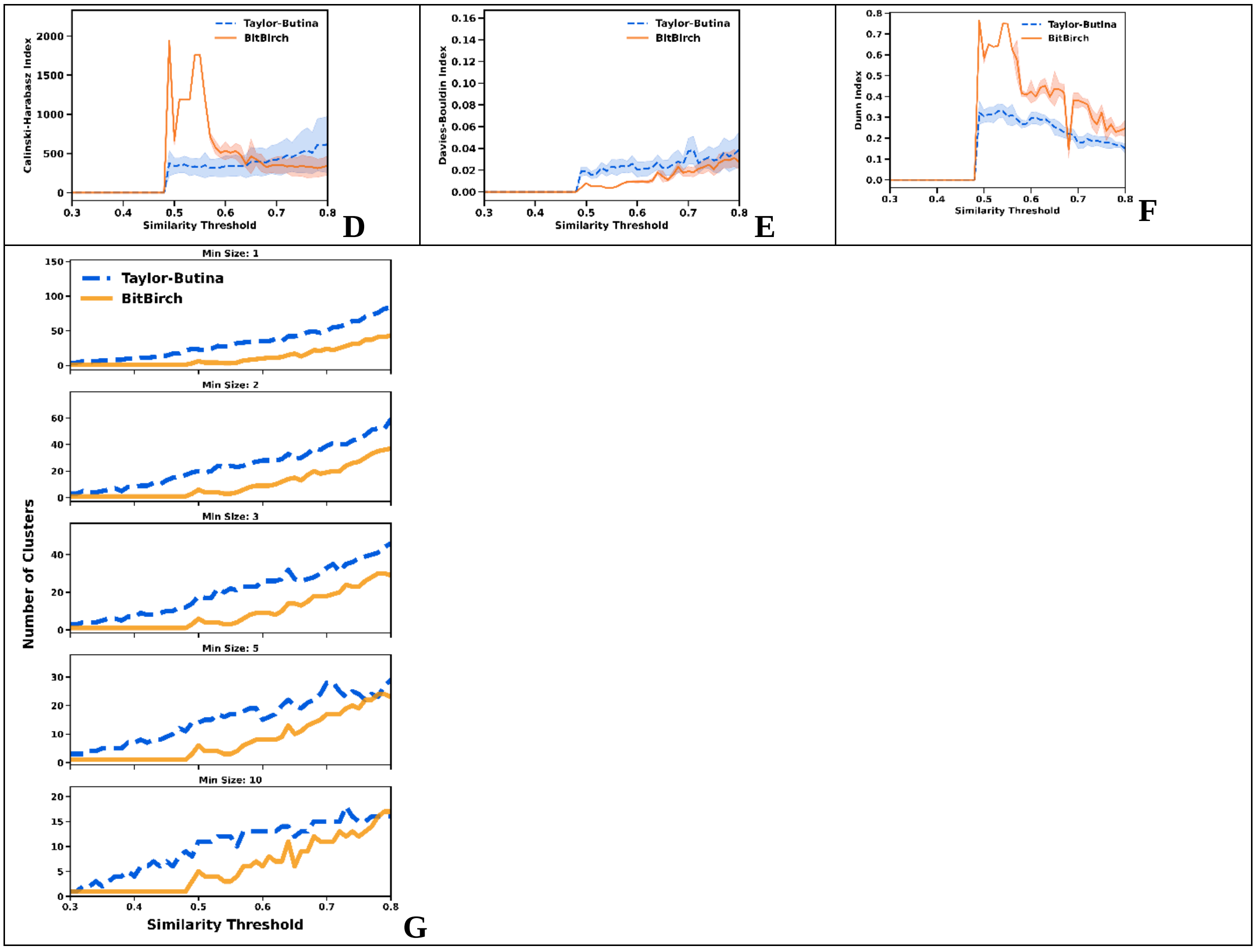


**Figure S4.15**: **A**: CHI, **B**: DBI, **C**: DI analysis for min_size = 1, 2, 3, 5, 10 for the [15] ChEMBL subset. Average **D**: CHI, **E**: DBI, **F**: DI values over the min_size variable. **G**: Number of clusters. BitBIRCH (orange continuous line), Taylor-Butina (blue dashed line).

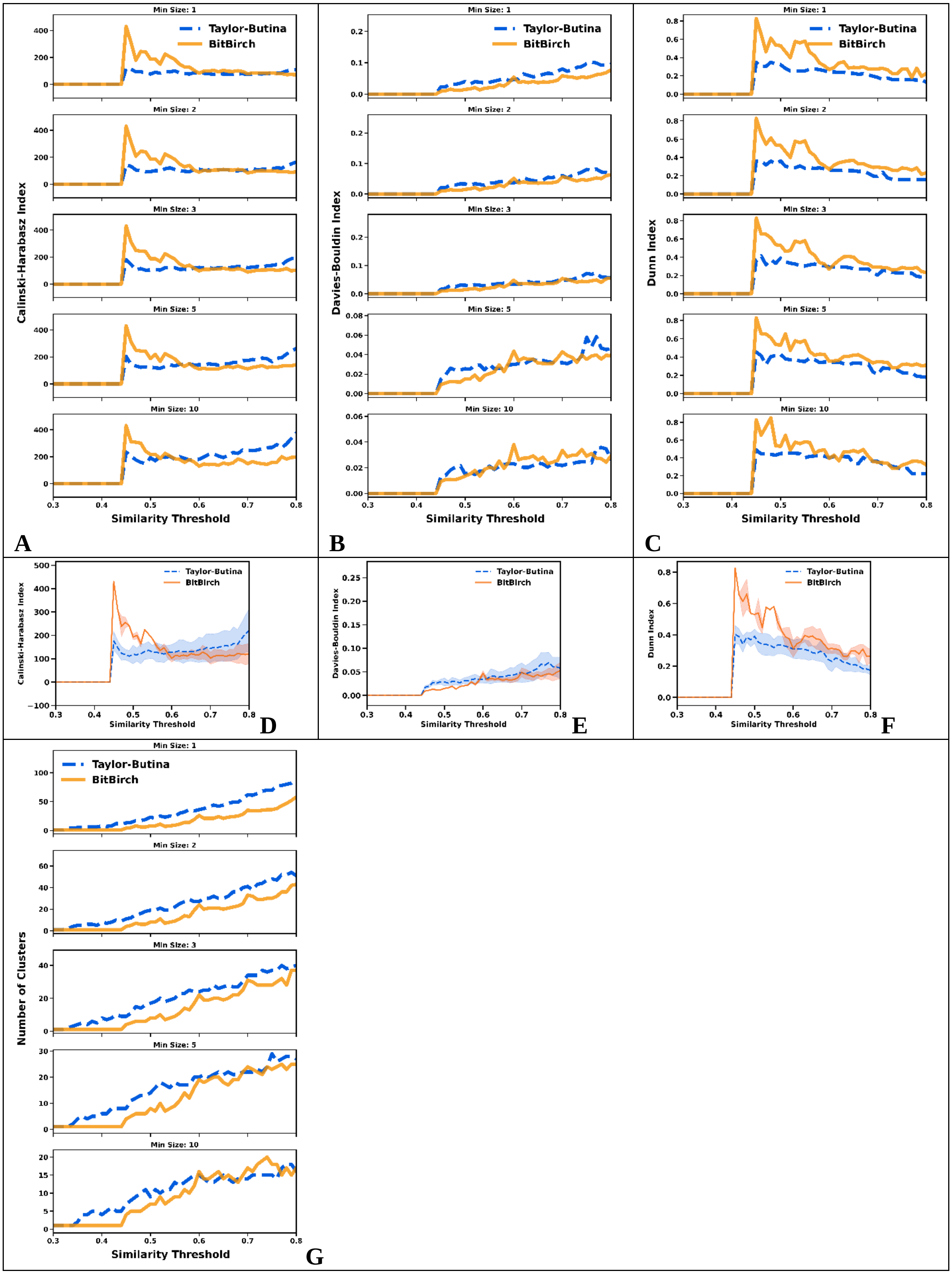

Min Size: 1
Min Size: 2
Min Size: 3
Min Size: 5
Min Size: 10
Taylor-Butina
BitBirch
Calinski-Harabasz Index
Davies-Bouldin Index
Dunn Index
Number of Clusters
Similarity Threshold
A
B
C
D
E
F
G

**Figure S4.16**: **A**: CHI, **B**: DBI, **C**: DI analysis for min_size = 1, 2, 3, 5, 10 for the [16] ChEMBL subset. Average **D**: CHI, **E**: DBI, **F**: DI values over the min_size variable. **G**: Number of clusters. BitBIRCH (orange continuous line), Taylor-Butina (blue dashed line).

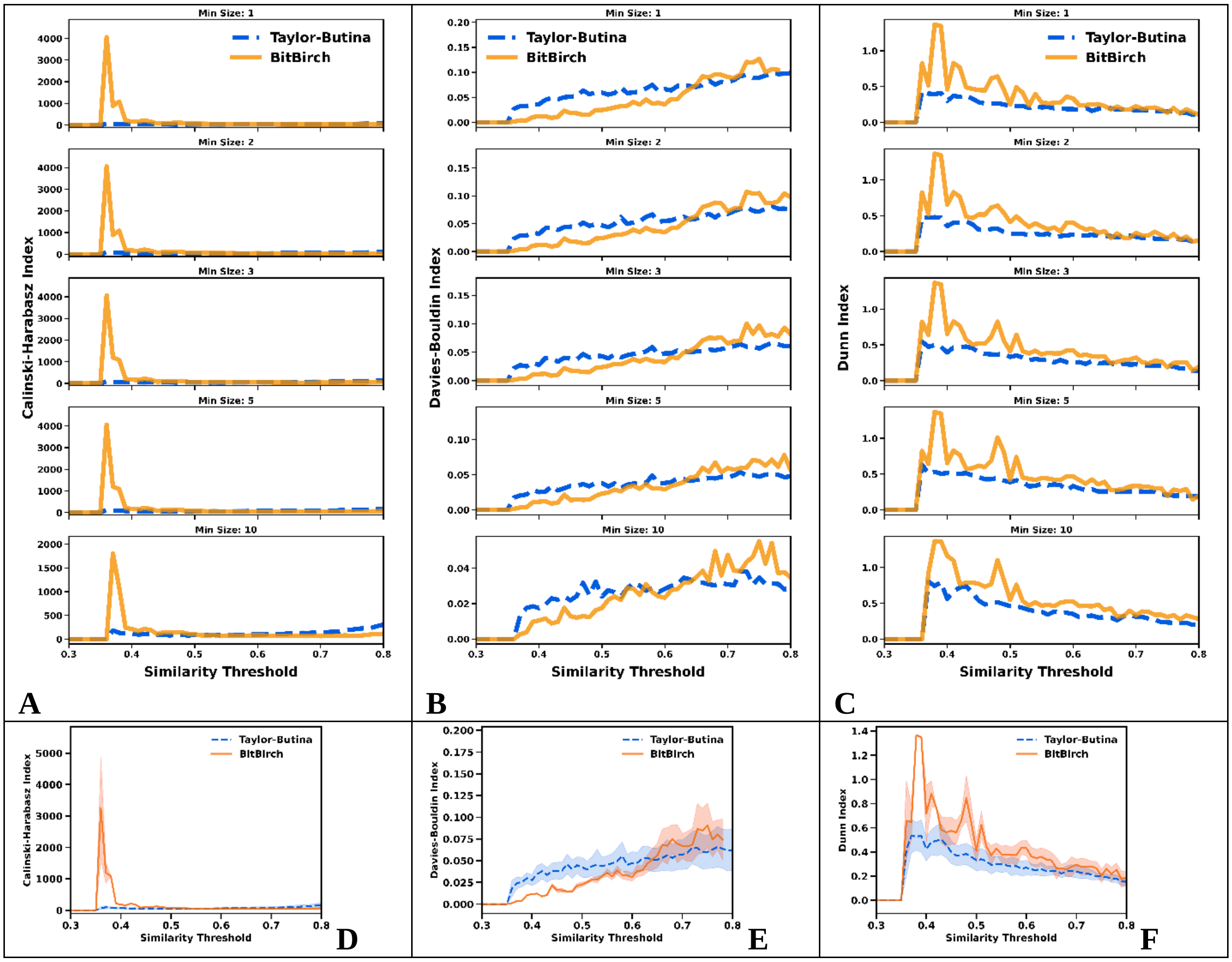

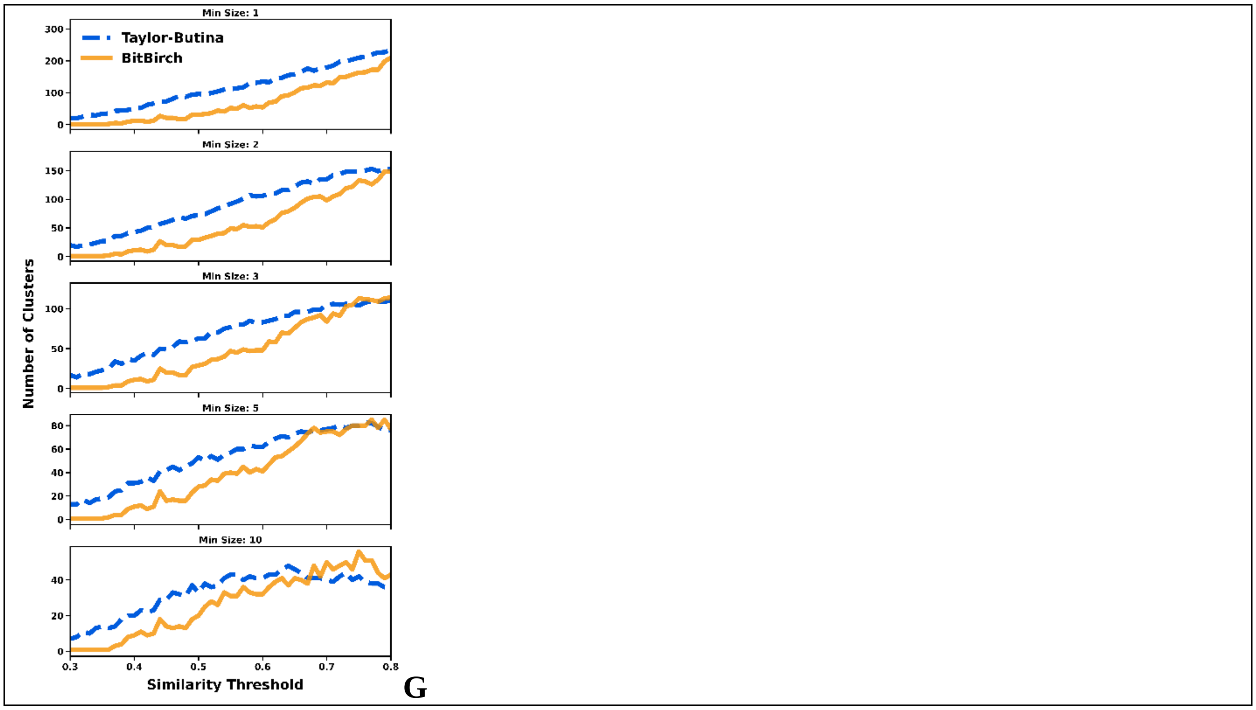


**Figure S4.17**: **A**: CHI, **B**: DBI, **C**: DI analysis for min_size = 1, 2, 3, 5, 10 for the [17] ChEMBL subset. Average **D**: CHI, **E**: DBI, **F**: DI values over the min_size variable. **G**: Number of clusters. BitBIRCH (orange continuous line), Taylor-Butina (blue dashed line).

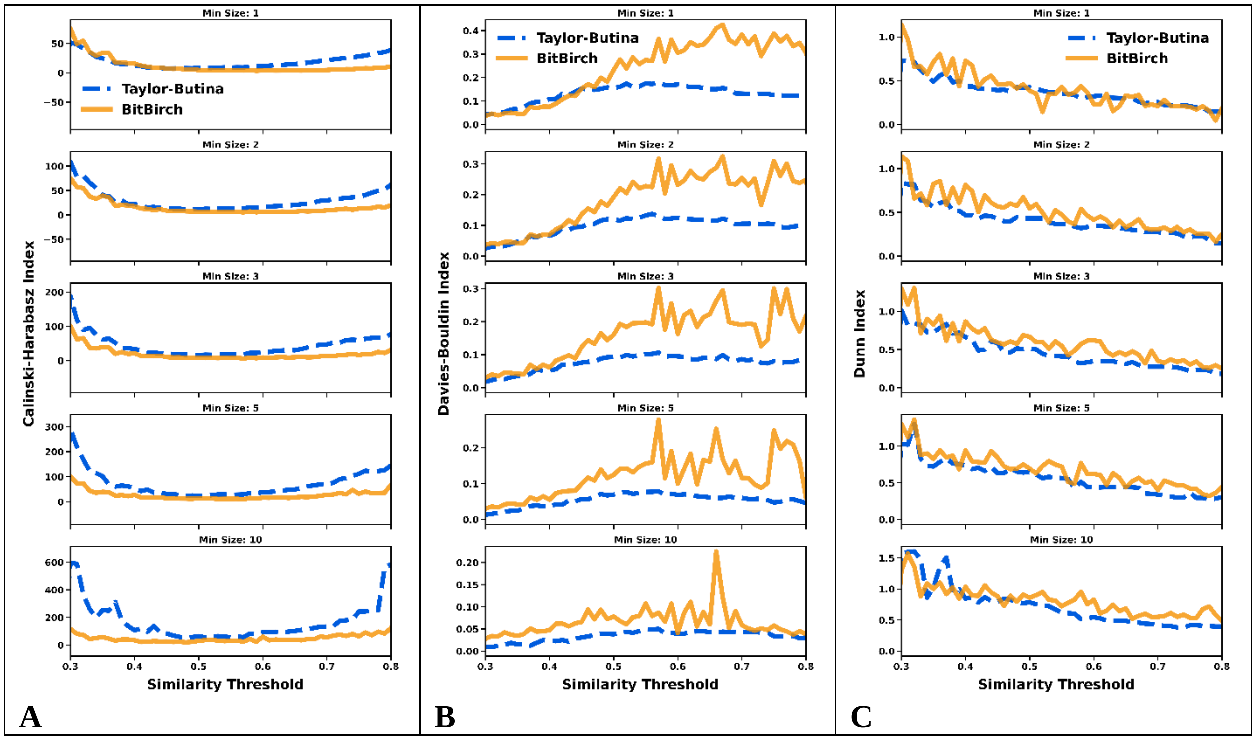

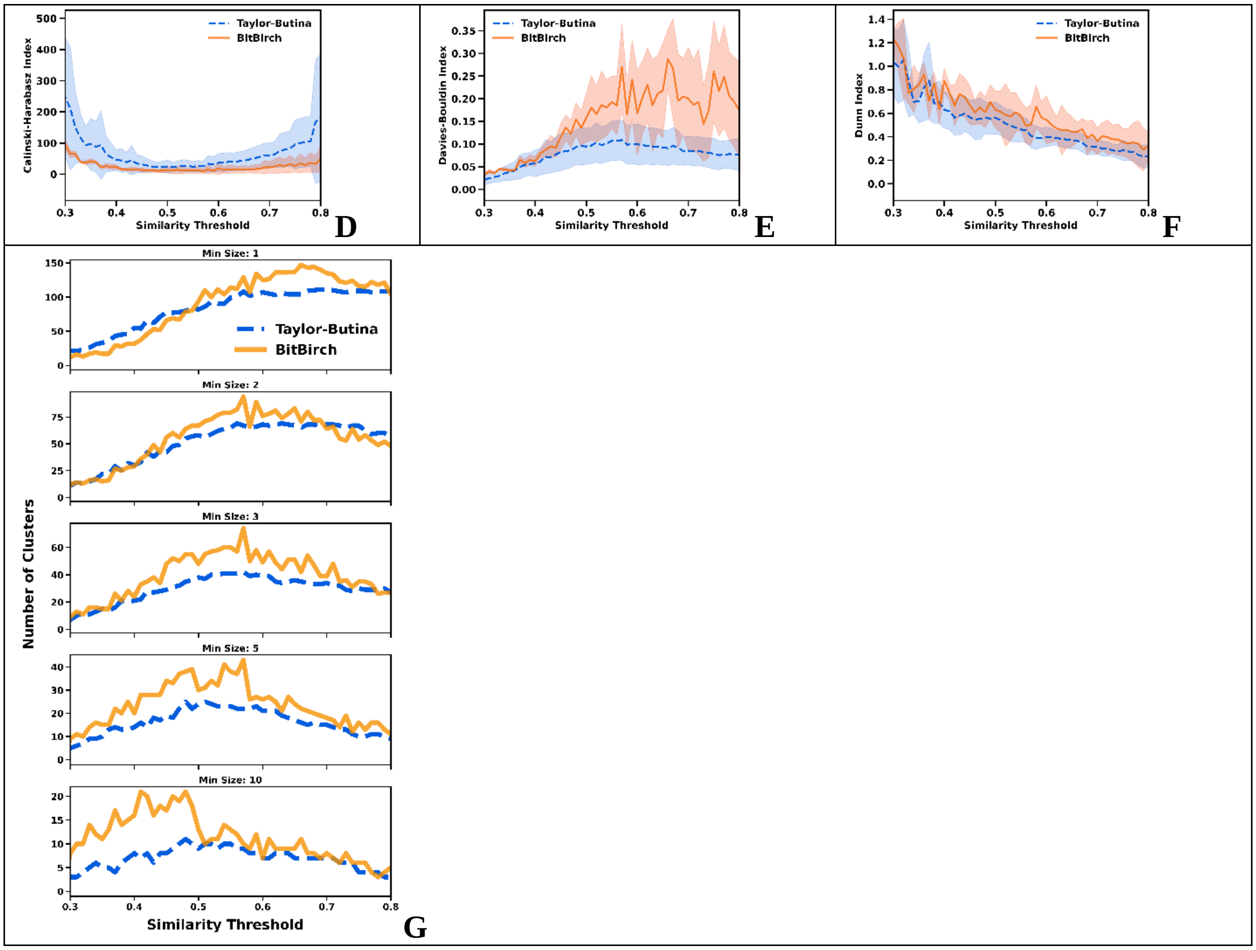


**Figure S4.18**: **A**: CHI, **B**: DBI, **C**: DI analysis for min_size = 1, 2, 3, 5, 10 for the [18] ChEMBL subset. Average **D**: CHI, **E**: DBI, **F**: DI values over the min_size variable. **G**: Number of clusters. BitBIRCH (orange continuous line), Taylor-Butina (blue dashed line).

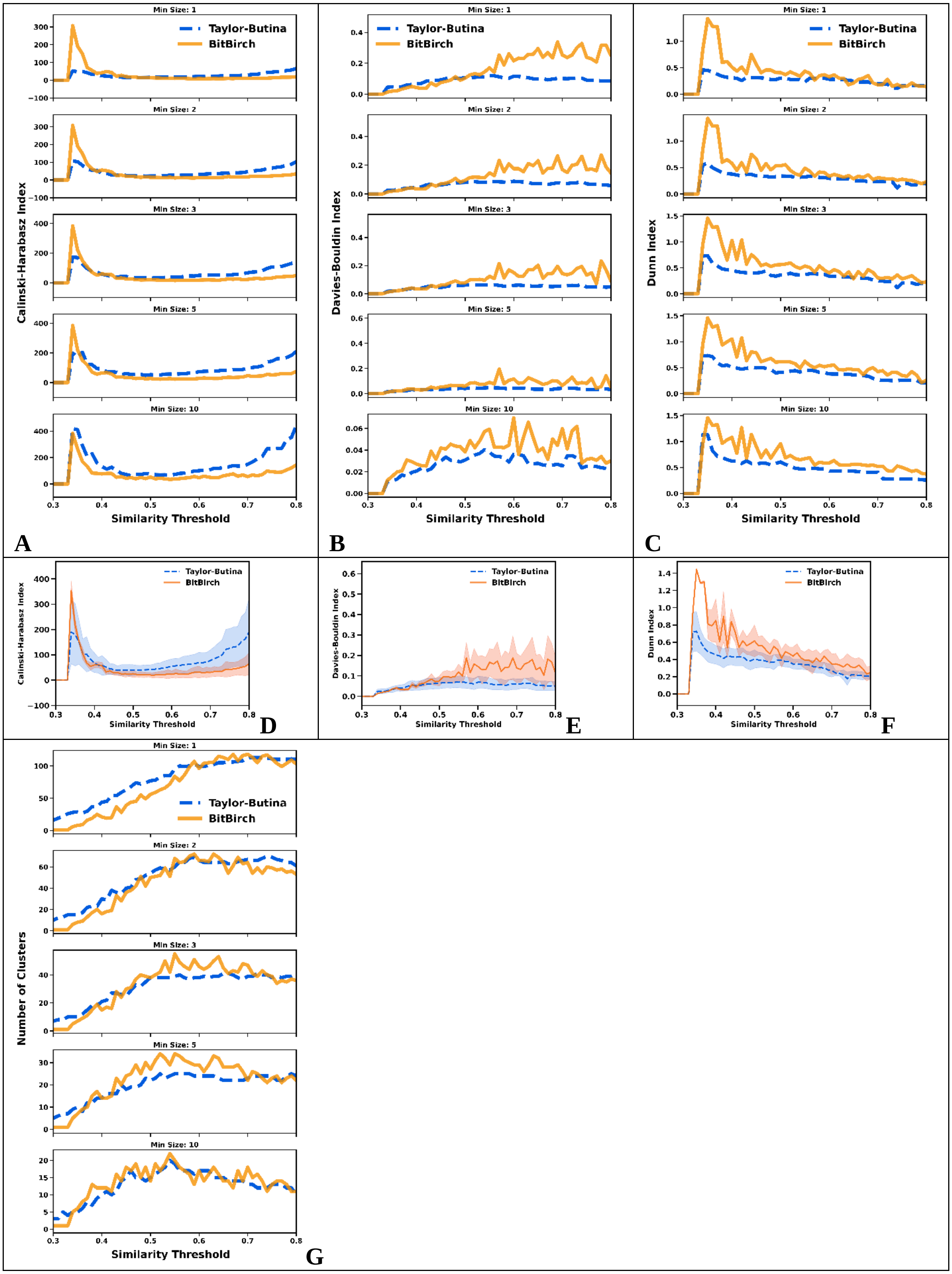

Min Size: 1
Min Size: 2
Min Size: 3
Min Size: 5
Min Size: 10
Taylor-Butina
BitBirch
Calinski-Harabasz Index
Davies-Bouldin Index
Dunn Index
Number of Clusters
Similarity Threshold
A
B
C
D
E
F
G

**Figure S4.19**: **A**: CHI, **B**: DBI, **C**: DI analysis for min_size = 1, 2, 3, 5, 10 for the [19] ChEMBL subset. Average **D**: CHI, **E**: DBI, **F**: DI values over the min_size variable. **G**: Number of clusters. BitBIRCH (orange continuous line), Taylor-Butina (blue dashed line).

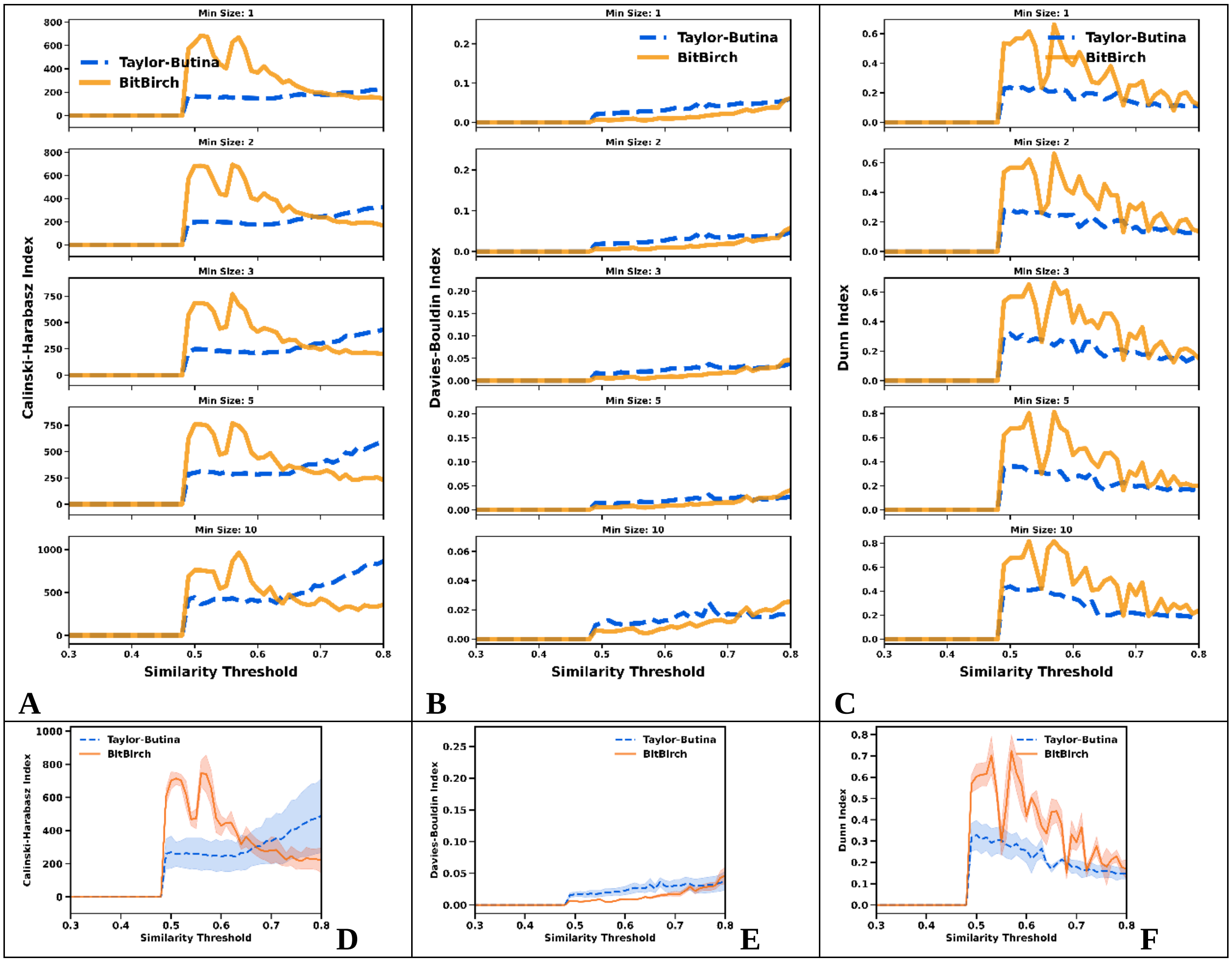

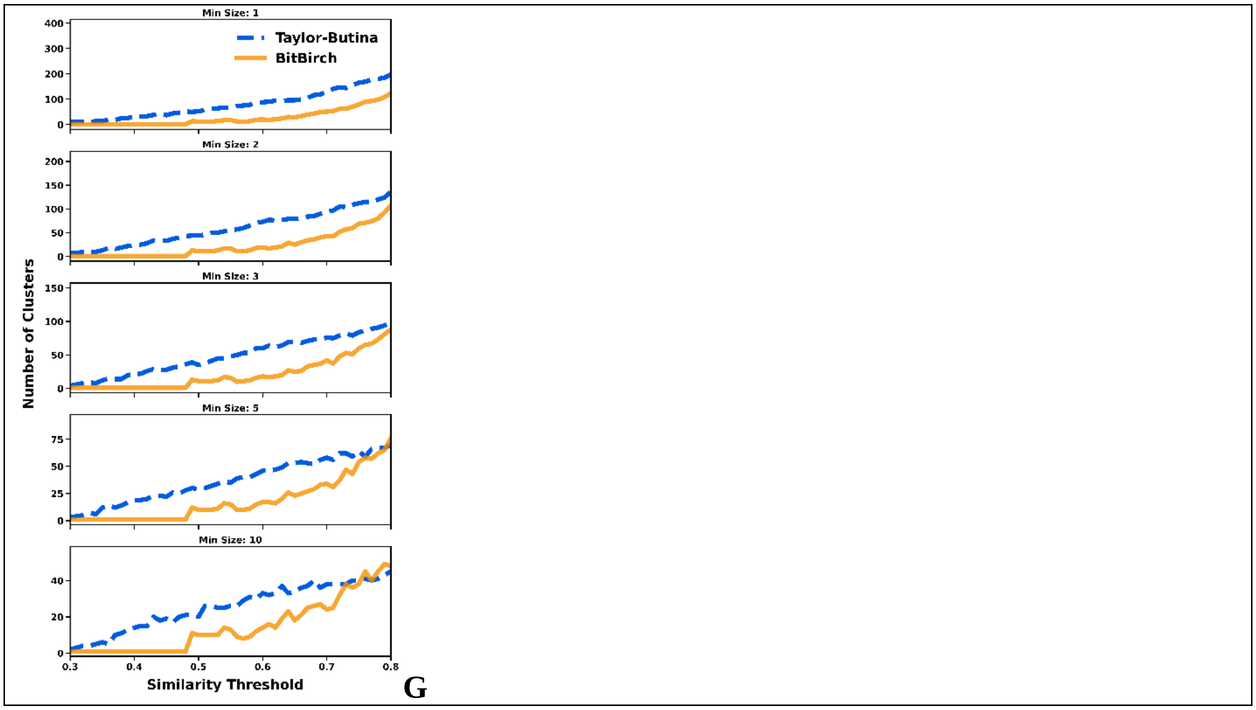


**Figure S4.20**: **A**: CHI, **B**: DBI, **C**: DI analysis for min_size = 1, 2, 3, 5, 10 for the [20] ChEMBL subset. Average **D**: CHI, **E**: DBI, **F**: DI values over the min_size variable. **G**: Number of clusters. BitBIRCH (orange continuous line), Taylor-Butina (blue dashed line).

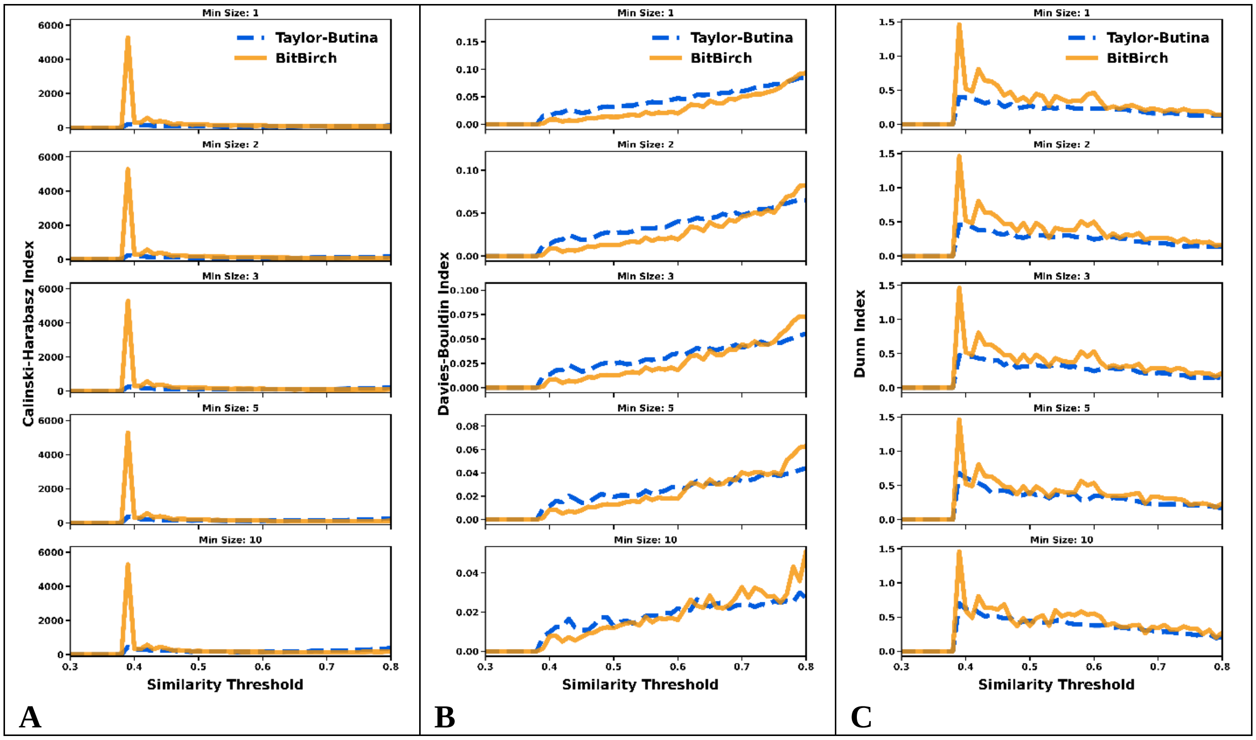

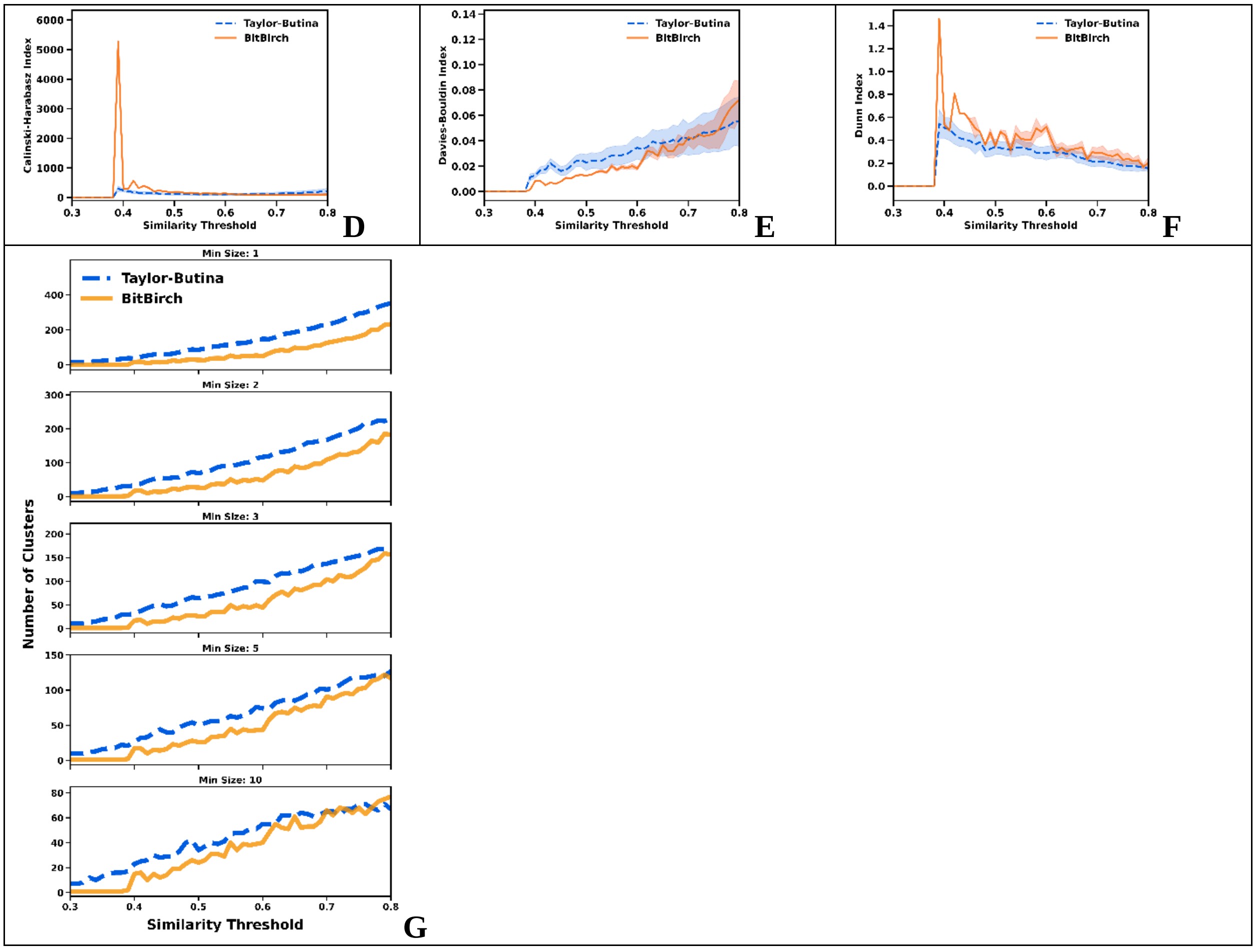


**Figure S4.21**: **A**: CHI, **B**: DBI, **C**: DI analysis for min_size = 1, 2, 3, 5, 10 for the [21] ChEMBL subset. Average **D**: CHI, **E**: DBI, **F**: DI values over the min_size variable. **G**: Number of clusters. BitBIRCH (orange continuous line), Taylor-Butina (blue dashed line).

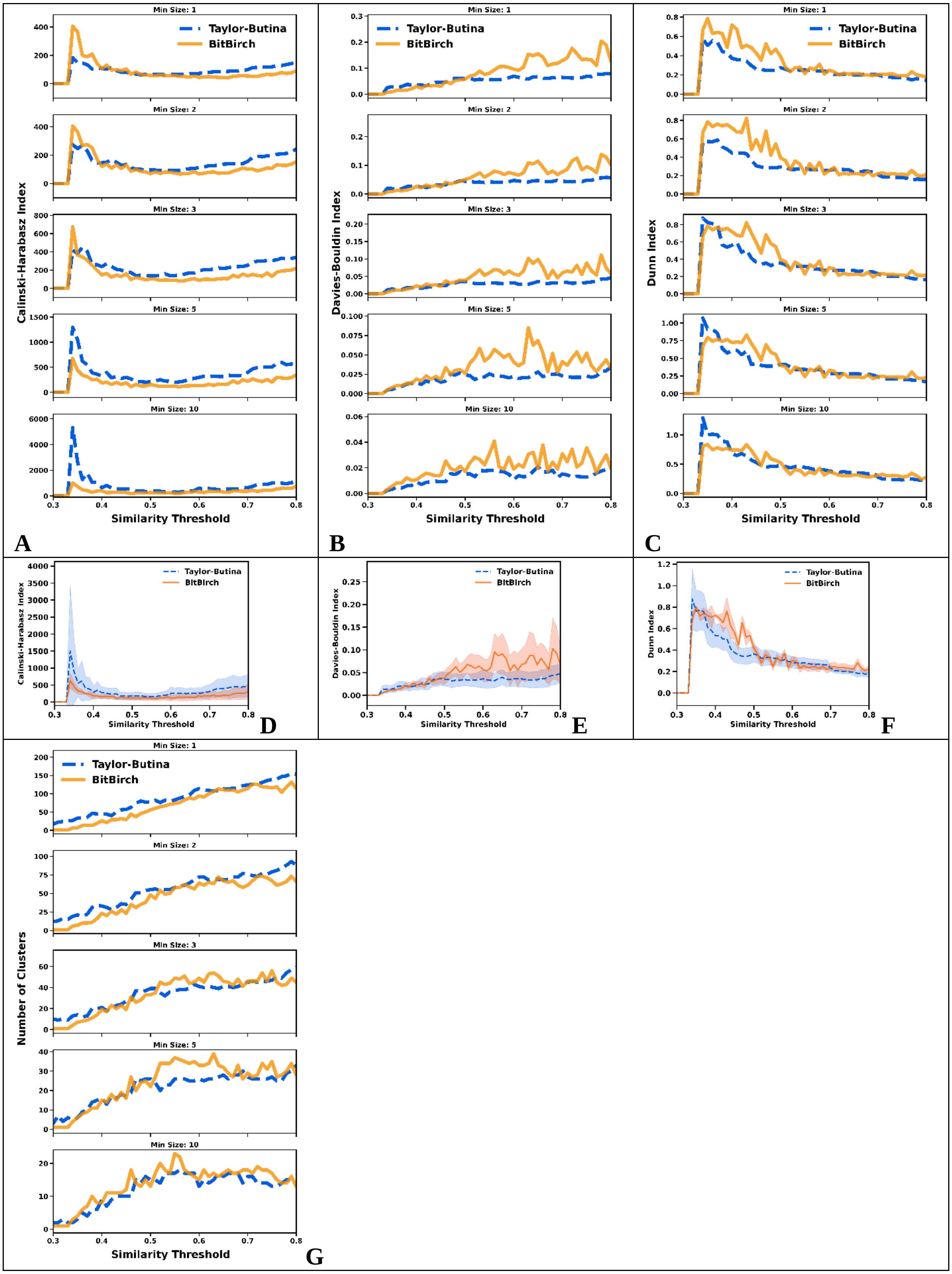

Min Size: 1
Min Size: 2
Min Size: 3
Min Size: 5
Min Size: 10
Taylor-Butina
BitBirch
Calinski-Harabasz Index
Davies-Bouldin Index
Dunn Index
Number of Clusters
Similarity Threshold
A
B
C
D
E
F
G

**Figure S4.22**: **A**: CHI, **B**: DBI, **C**: DI analysis for min_size = 1, 2, 3, 5, 10 for the [22] ChEMBL subset. Average **D**: CHI, **E**: DBI, **F**: DI values over the min_size variable. **G**: Number of clusters. BitBIRCH (orange continuous line), Taylor-Butina (blue dashed line).

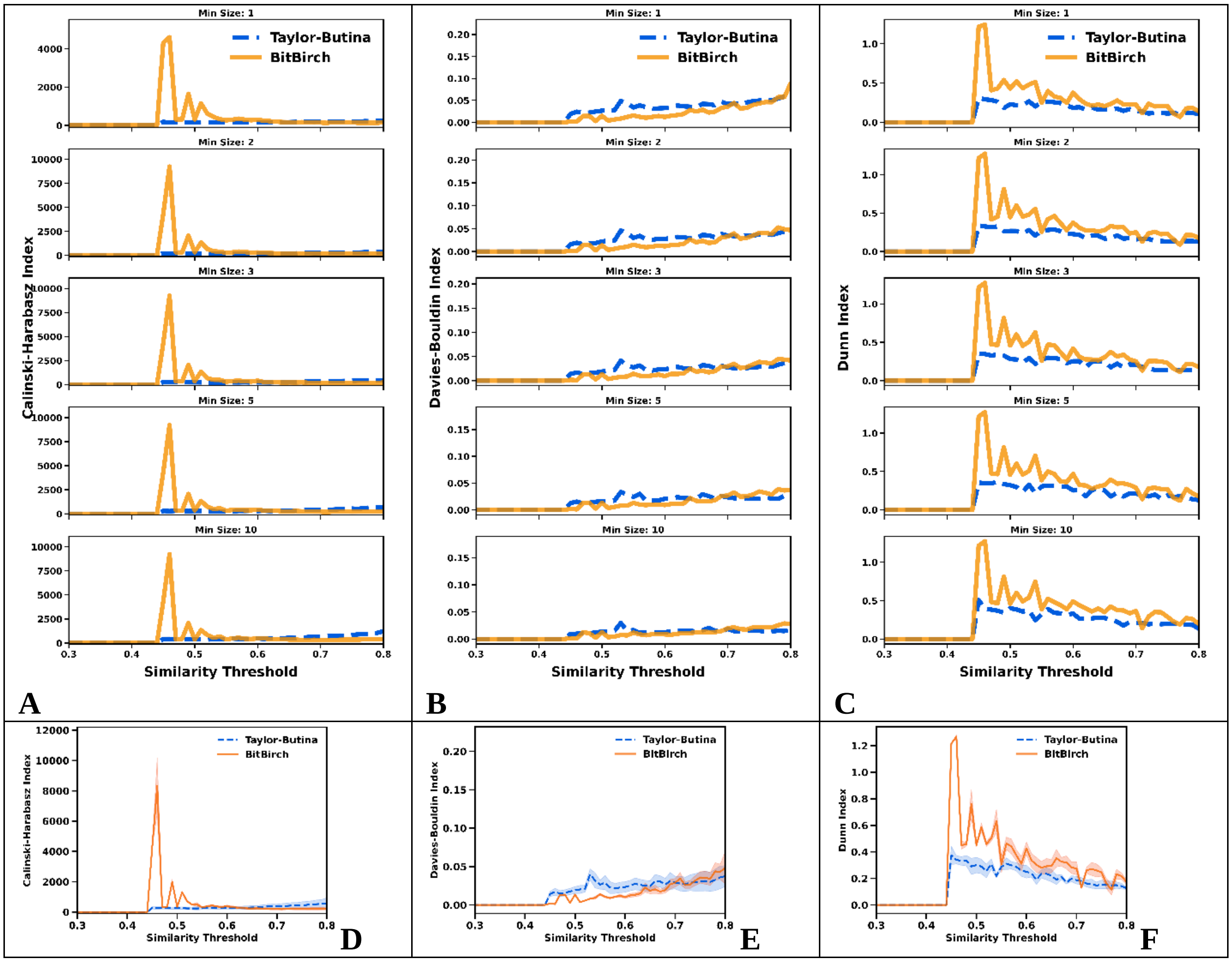

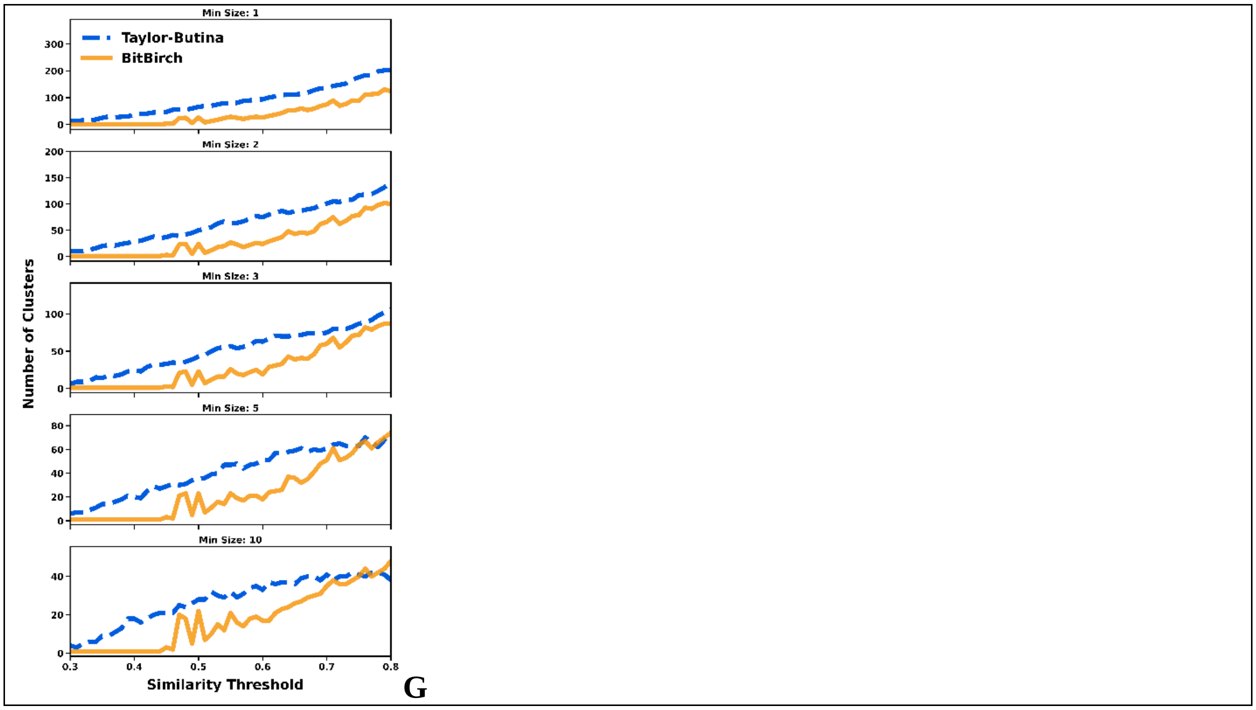


**Figure S4.23**: **A**: CHI, **B**: DBI, **C**: DI analysis for min_size = 1, 2, 3, 5, 10 for the [23] ChEMBL subset. Average **D**: CHI, **E**: DBI, **F**: DI values over the min_size variable. **G**: Number of clusters. BitBIRCH (orange continuous line), Taylor-Butina (blue dashed line).

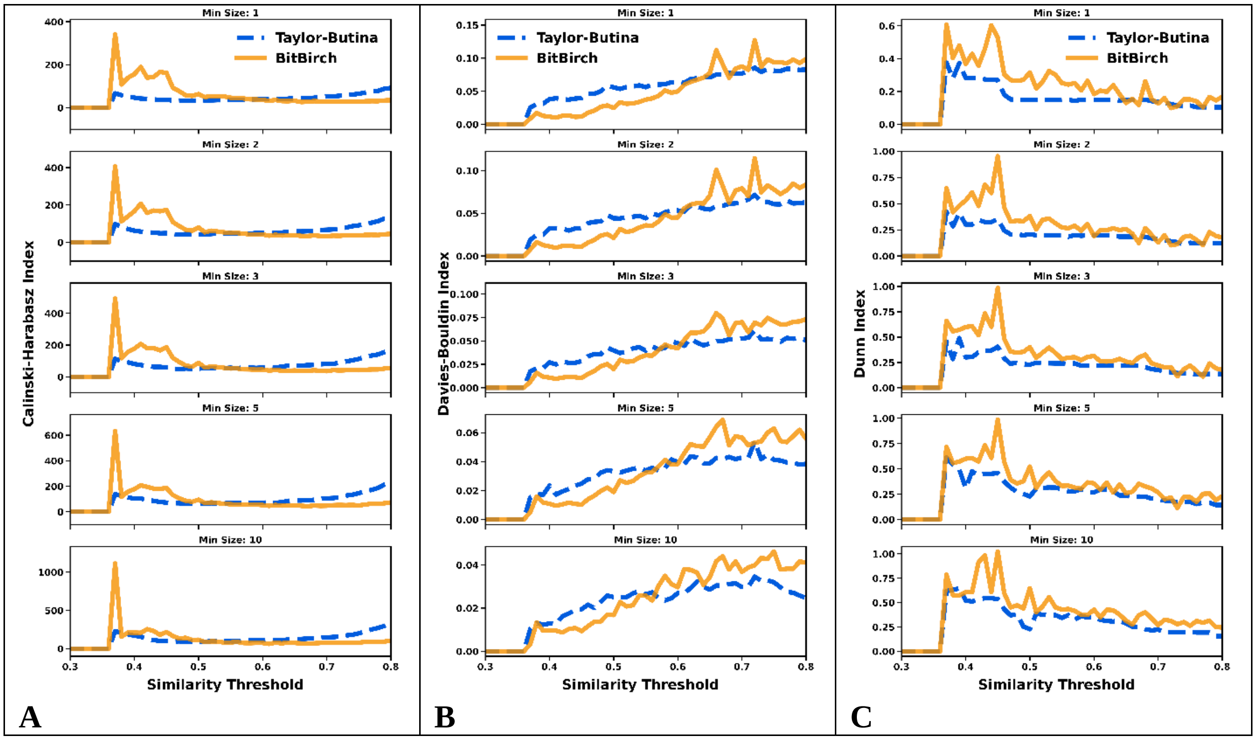

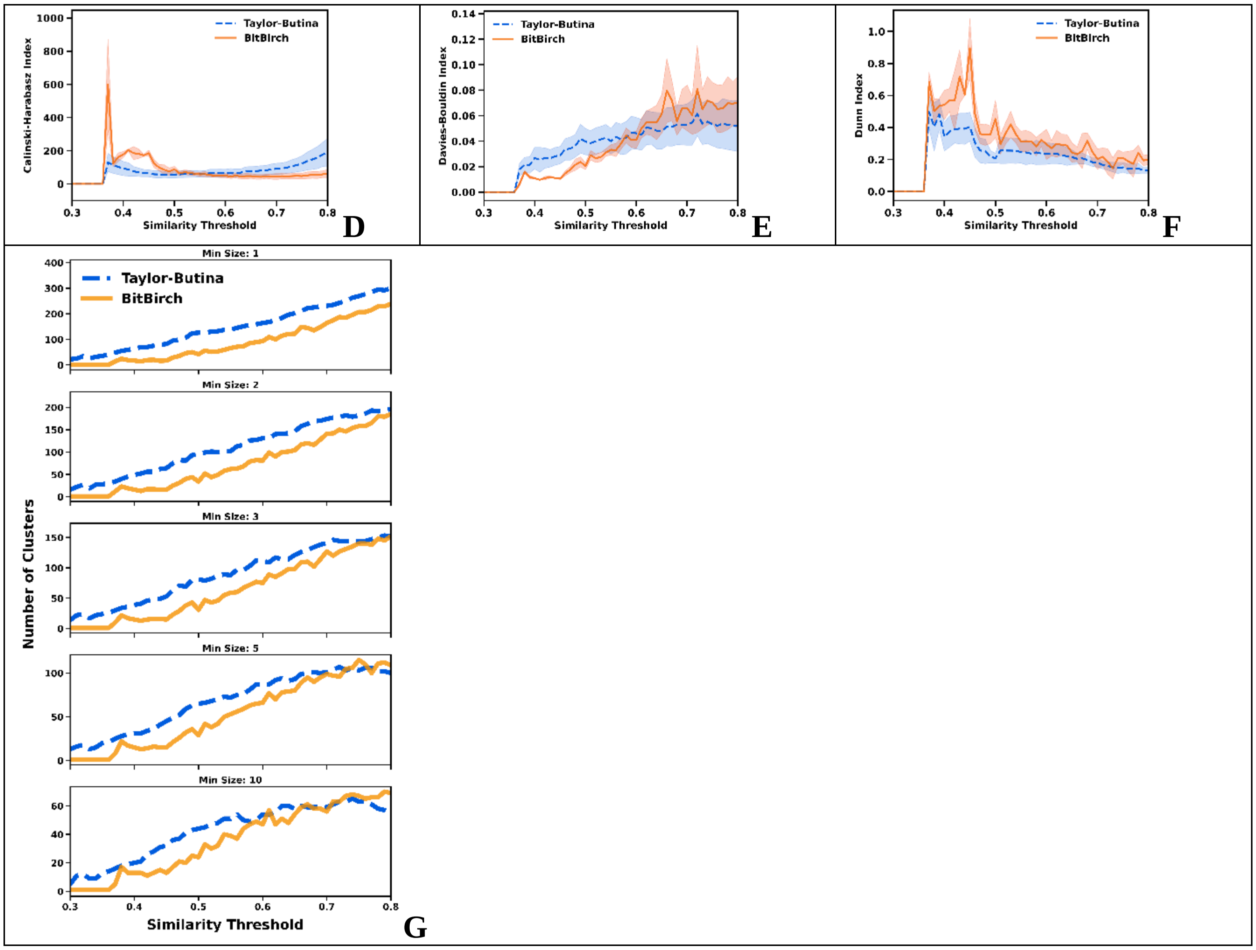


**Figure S4.24**: **A**: CHI, **B**: DBI, **C**: DI analysis for min_size = 1, 2, 3, 5, 10 for the [24] ChEMBL subset. Average **D**: CHI, **E**: DBI, **F**: DI values over the min_size variable. **G**: Number of clusters. BitBIRCH (orange continuous line), Taylor-Butina (blue dashed line).

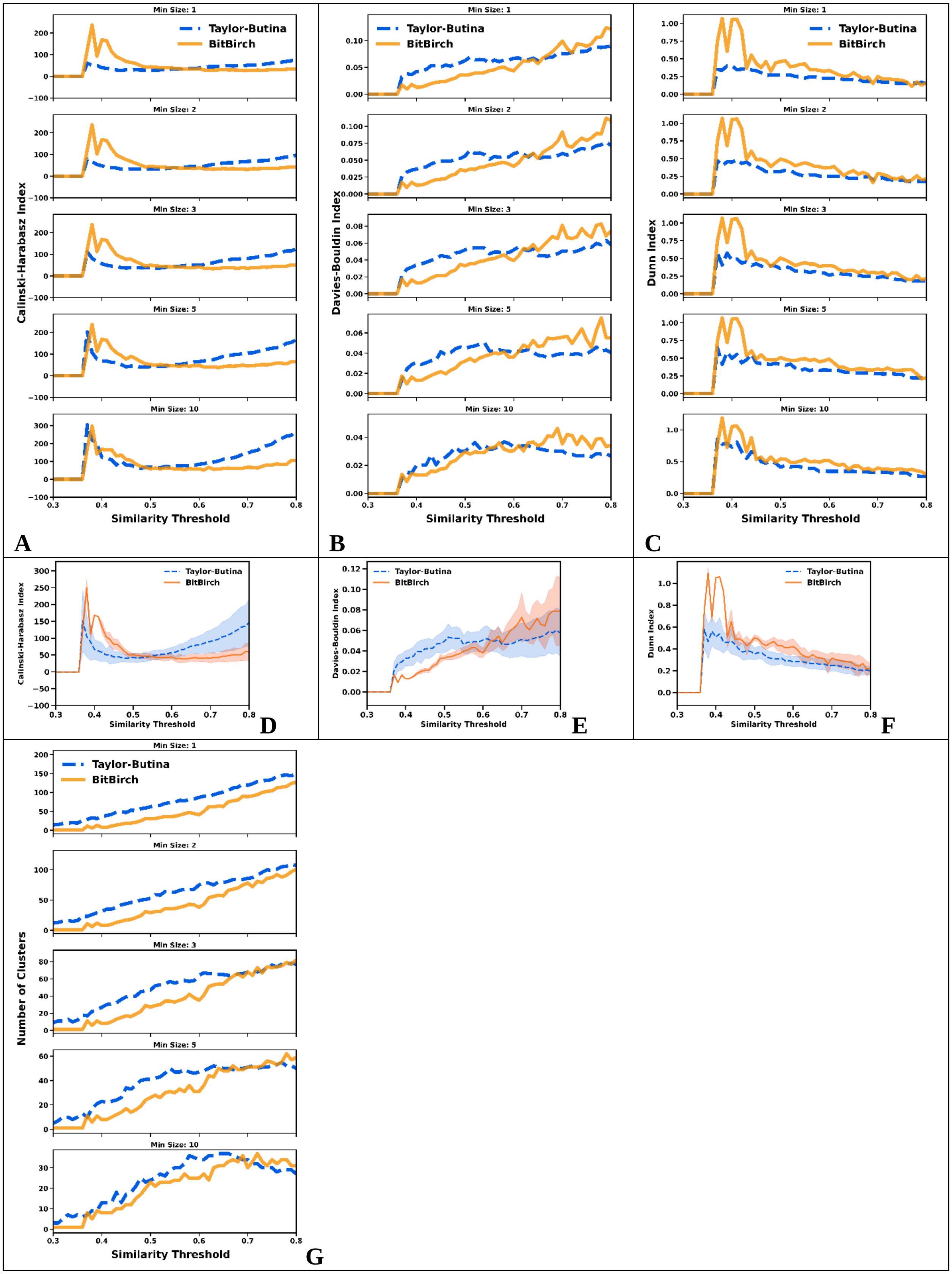
Min Size: 1
Min Size: 2
Min Size: 3
Min Size: 5
Min Size: 10
Taylor-Butina
BitBirch
Calinski-Harabasz Index
Davies-Bouldin Index
Dunn Index
Number of Clusters
Similarity Threshold
A
B
C
D
E
F
G

**Figure S4.25**: **A**: CHI, **B**: DBI, **C**: DI analysis for min_size = 1, 2, 3, 5, 10 for the [25] ChEMBL subset. Average **D**: CHI, **E**: DBI, **F**: DI values over the min_size variable. **G**: Number of clusters. BitBIRCH (orange continuous line), Taylor-Butina (blue dashed line).

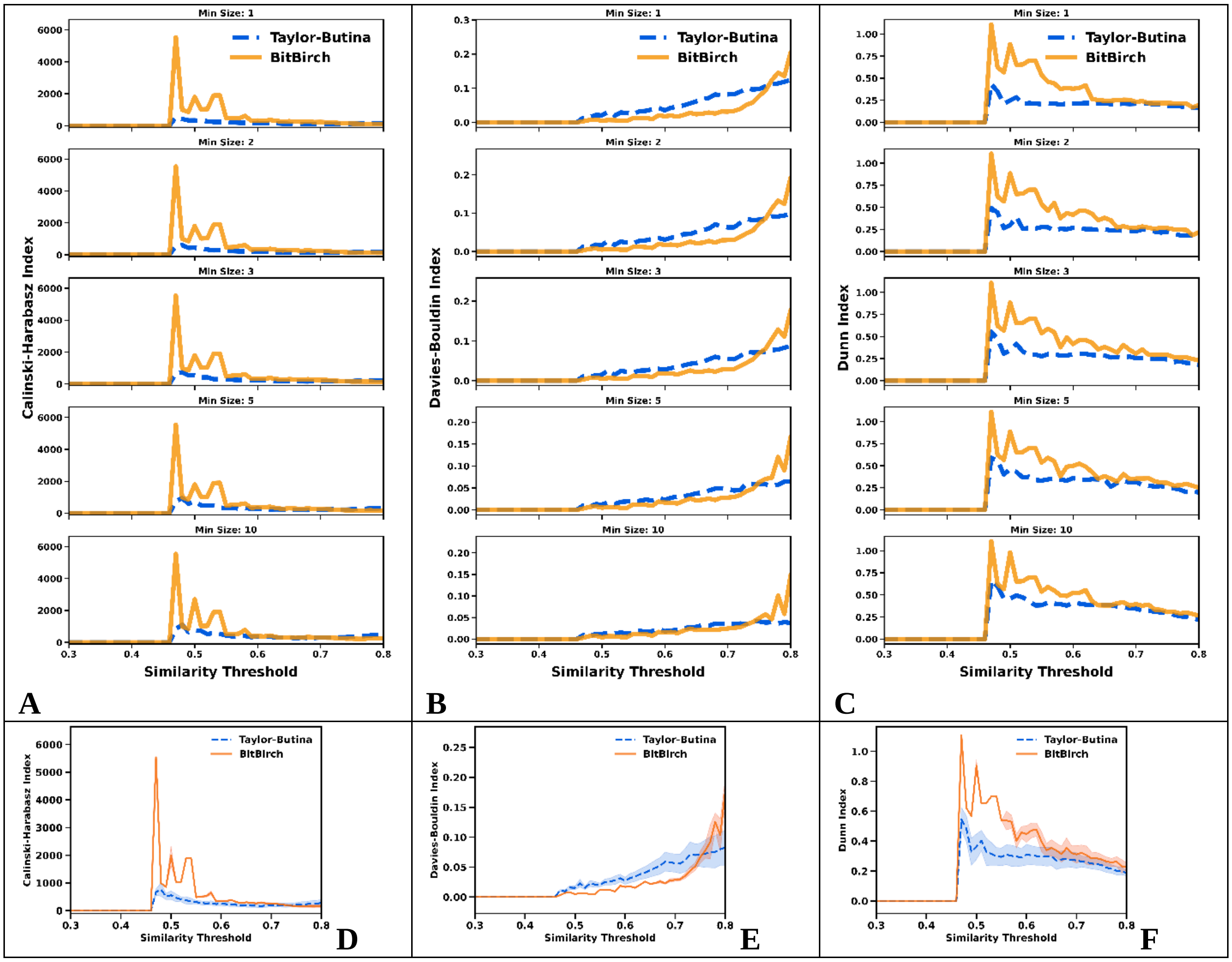

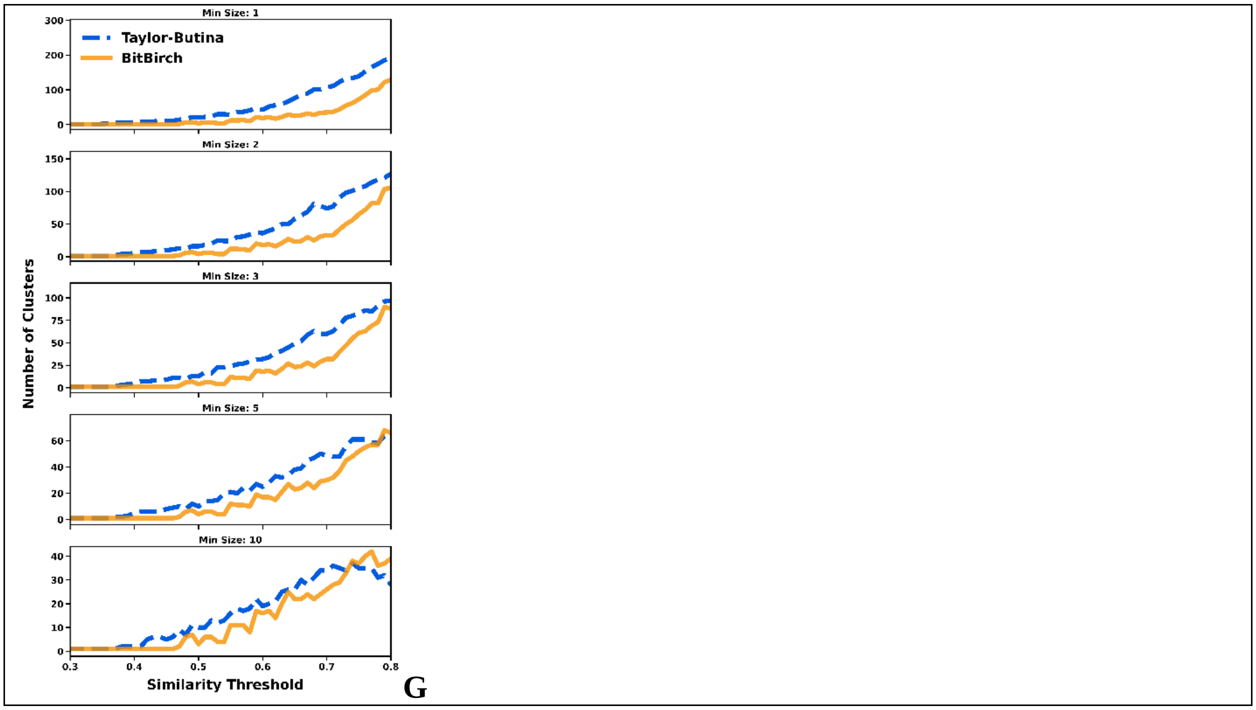


**Figure S4.26**: **A**: CHI, **B**: DBI, **C**: DI analysis for min_size = 1, 2, 3, 5, 10 for the [26] ChEMBL subset. Average **D**: CHI, **E**: DBI, **F**: DI values over the min_size variable. **G**: Number of clusters. BitBIRCH (orange continuous line), Taylor-Butina (blue dashed line).

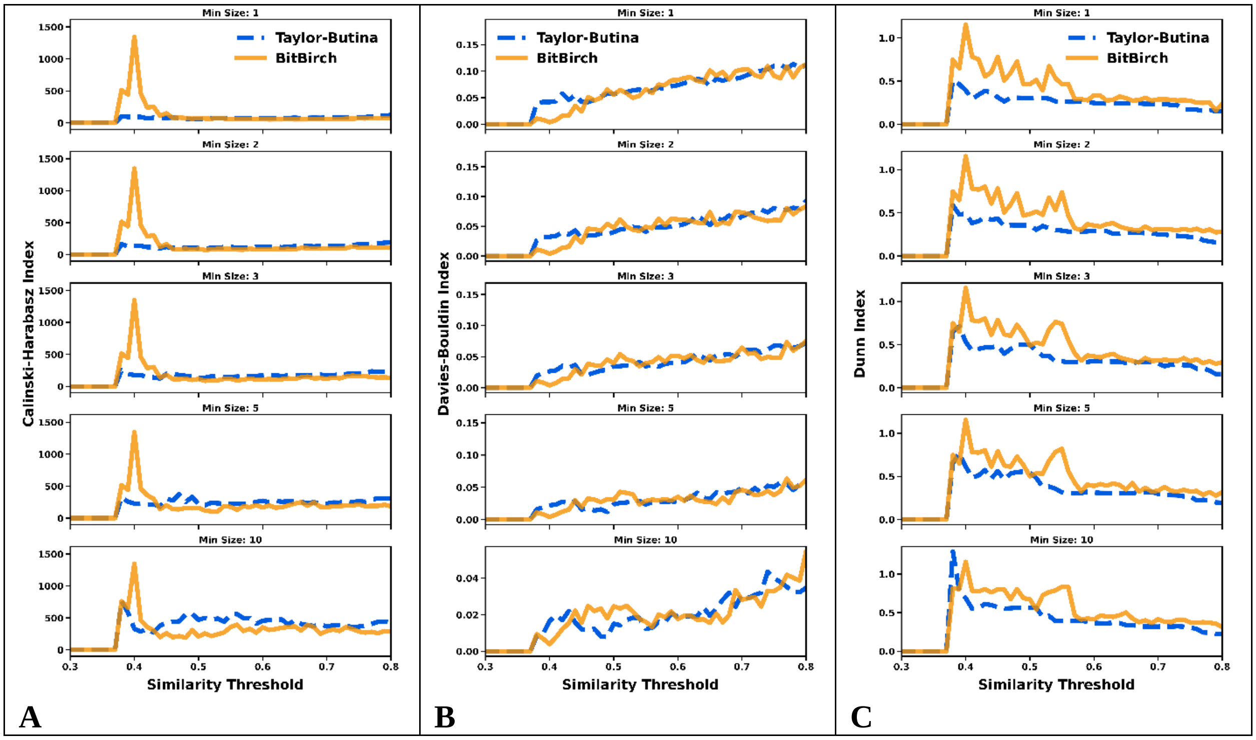

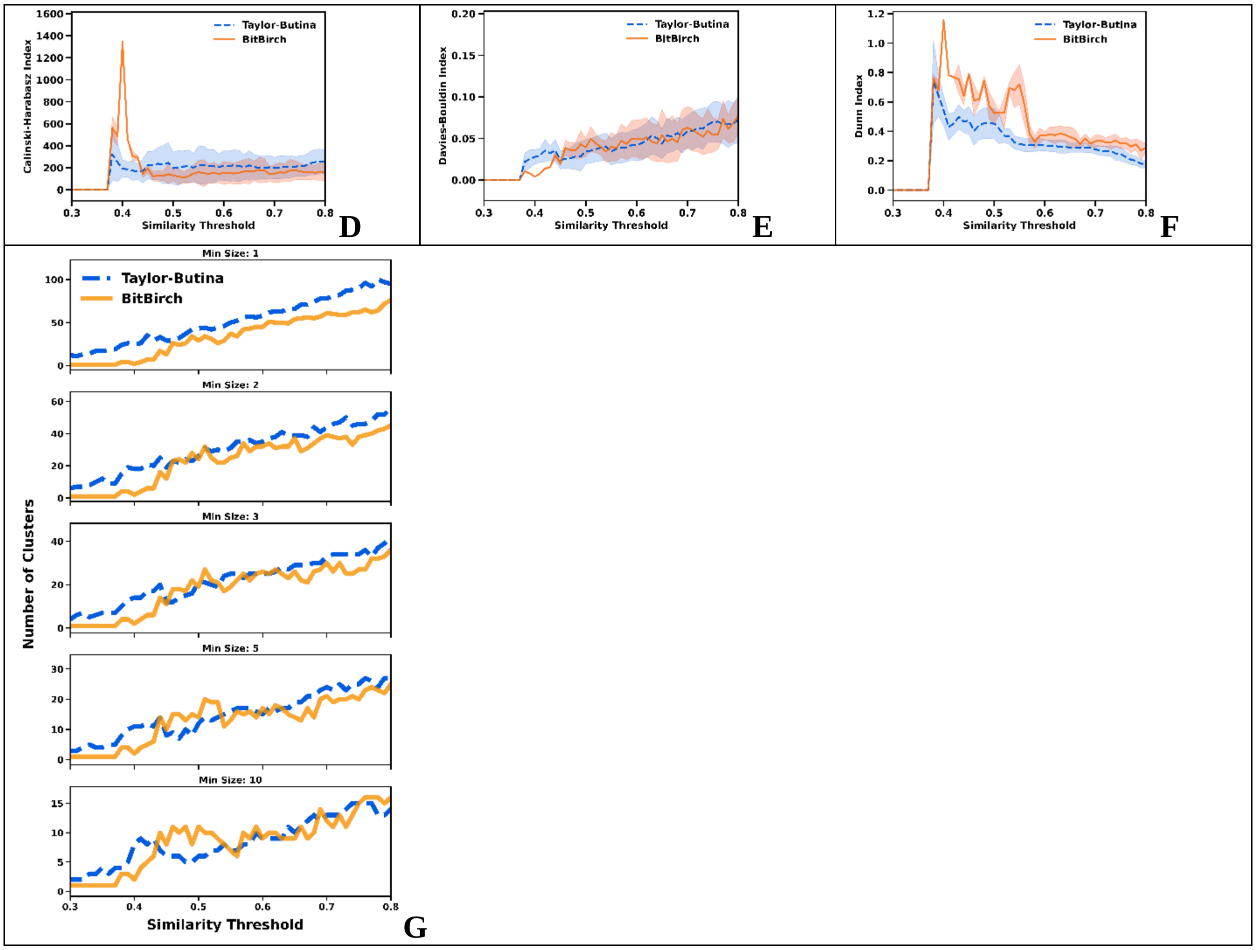


**Figure S4.27**: **A**: CHI, **B**: DBI, **C**: DI analysis for min_size = 1, 2, 3, 5, 10 for the [27] ChEMBL subset. Average **D**: CHI, **E**: DBI, **F**: DI values over the min_size variable. **G**: Number of clusters. BitBIRCH (orange continuous line), Taylor-Butina (blue dashed line).

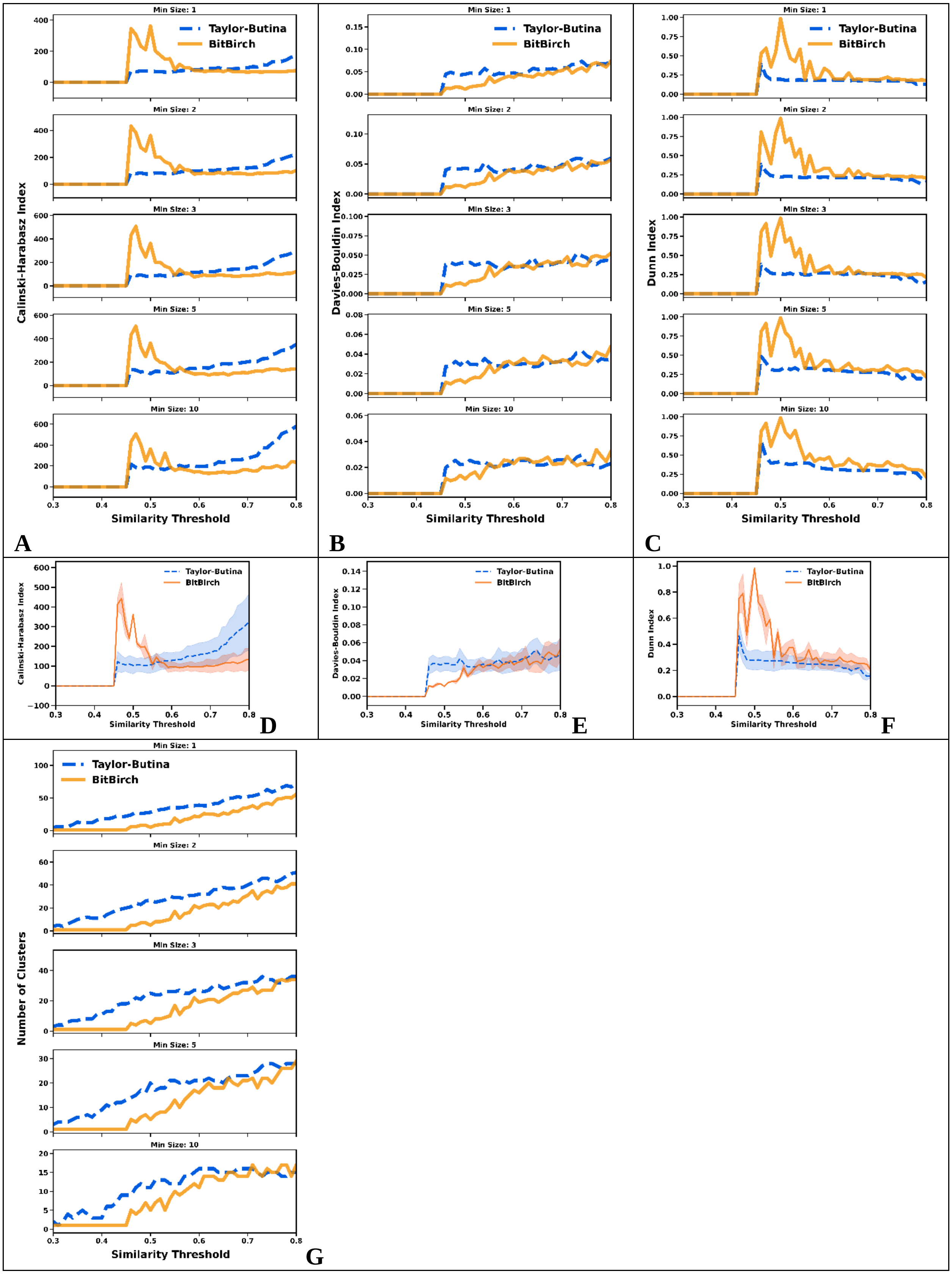
Min Size: 1
Min Size: 2
Min Size: 3
Min Size: 5
Min Size: 10
Taylor-Butina
BitBirch
Calinski-Harabasz Index
Davies-Bouldin Index
Dunn Index
Number of Clusters
Similarity Threshold
A
B
C
D
E
F
G

**Figure S4.28**: **A**: CHI, **B**: DBI, **C**: DI analysis for min_size = 1, 2, 3, 5, 10 for the [28] ChEMBL subset. Average **D**: CHI, **E**: DBI, **F**: DI values over the min_size variable. **G**: Number of clusters. BitBIRCH (orange continuous line), Taylor-Butina (blue dashed line).

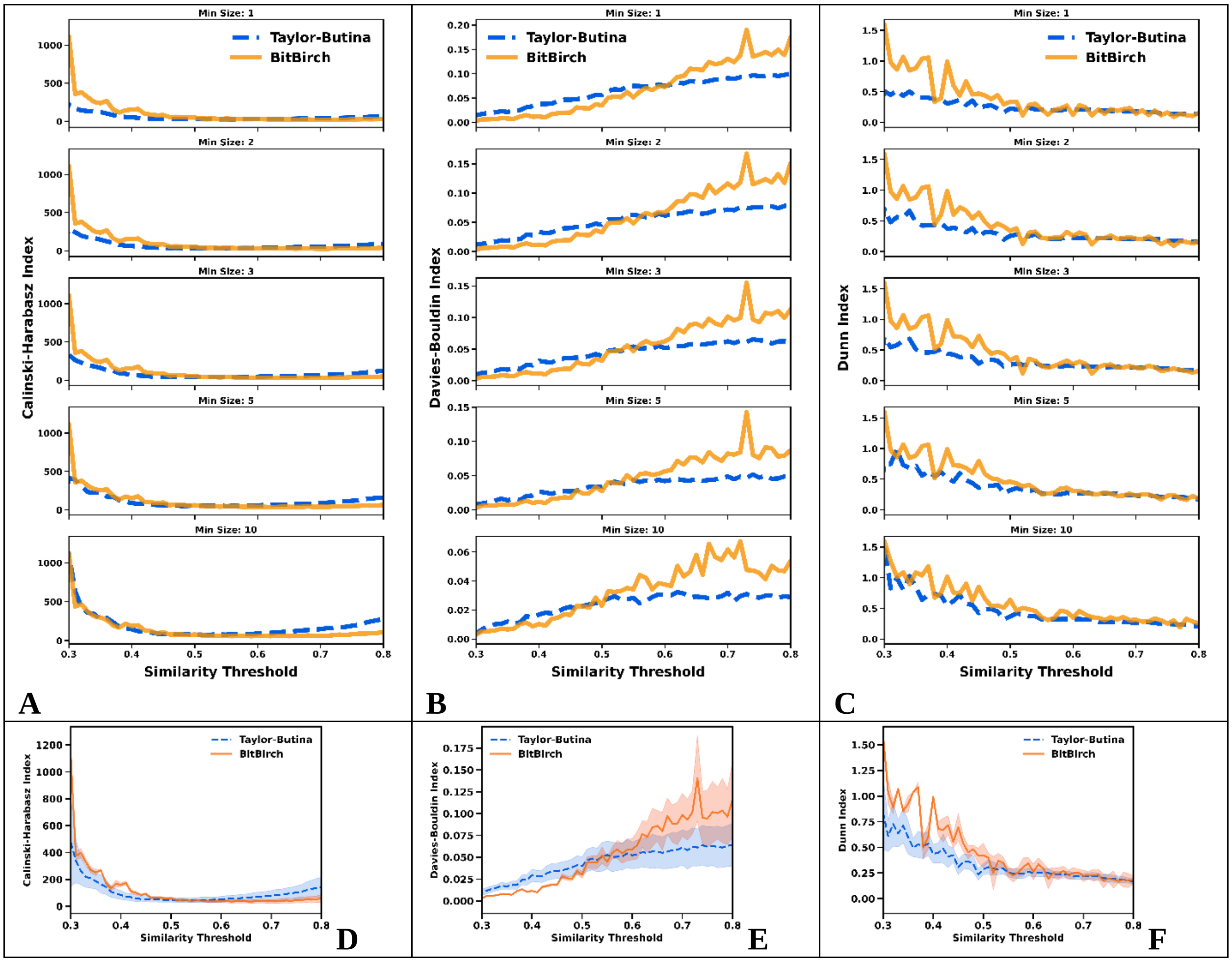

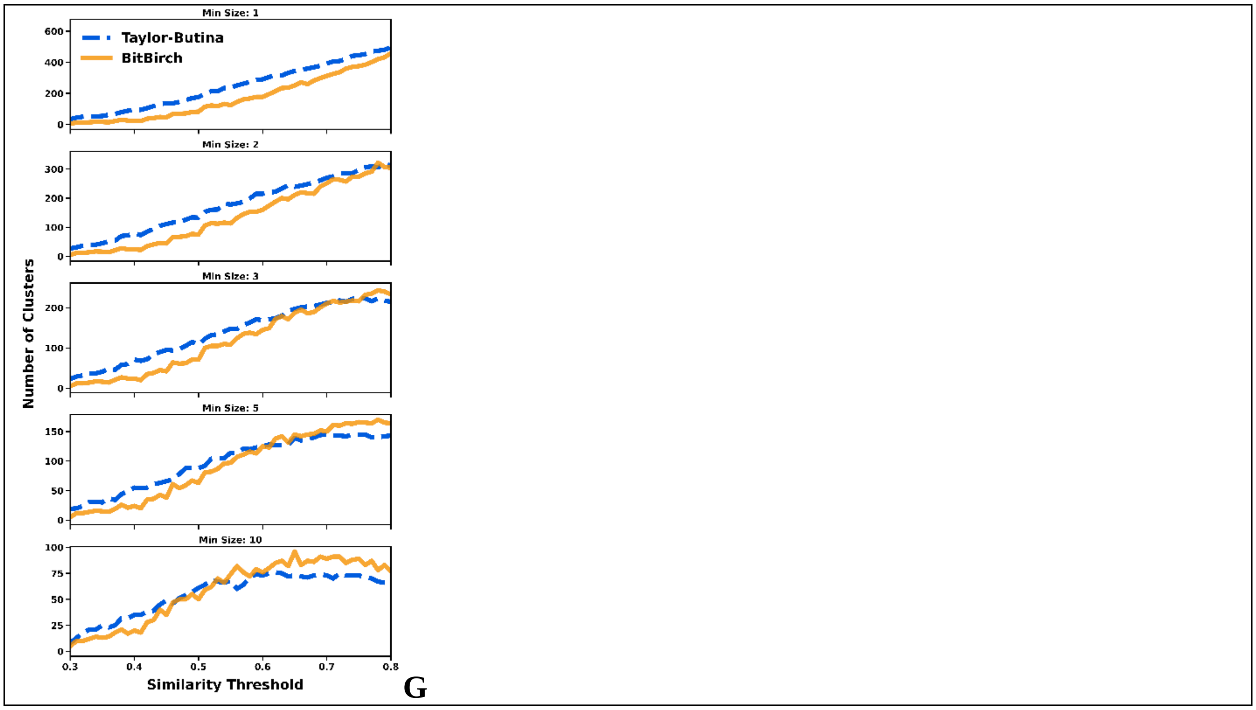


**Figure S4.29**: **A**: CHI, **B**: DBI, **C**: DI analysis for min_size = 1, 2, 3, 5, 10 for the [29] ChEMBL subset. Average **D**: CHI, **E**: DBI, **F**: DI values over the min_size variable. **G**: Number of clusters. BitBIRCH (orange continuous line), Taylor-Butina (blue dashed line).

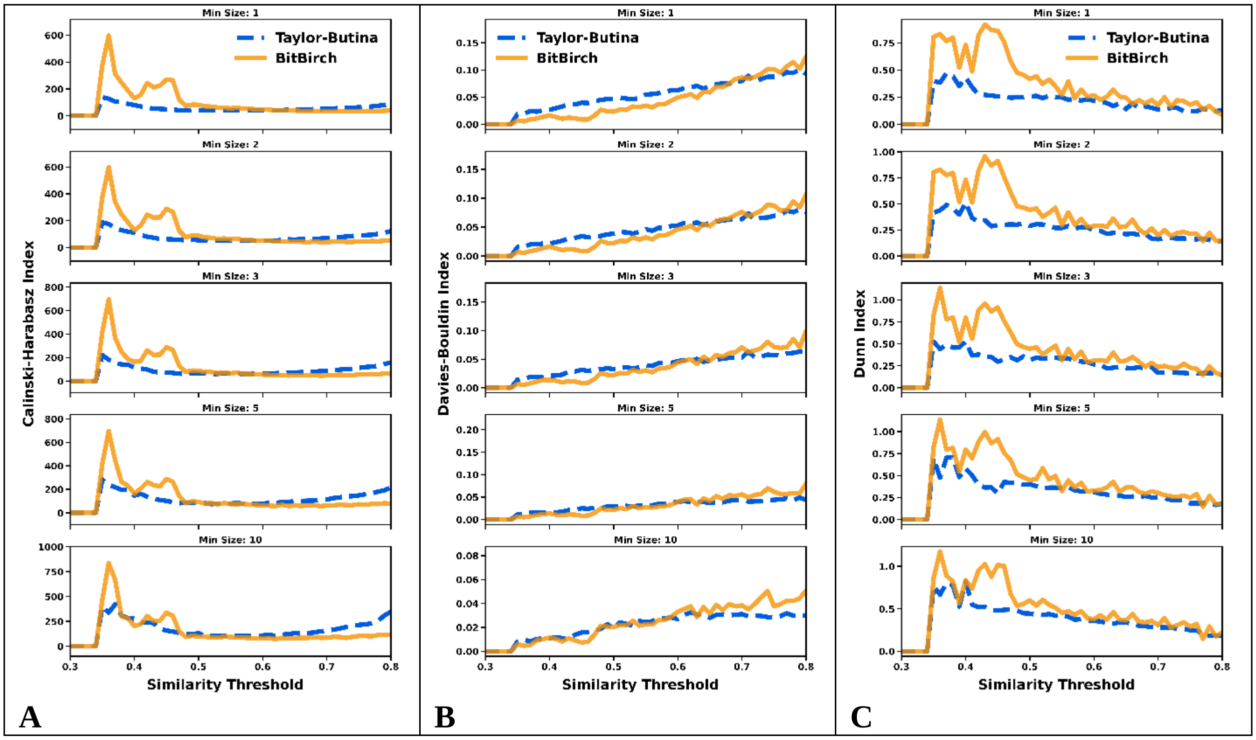

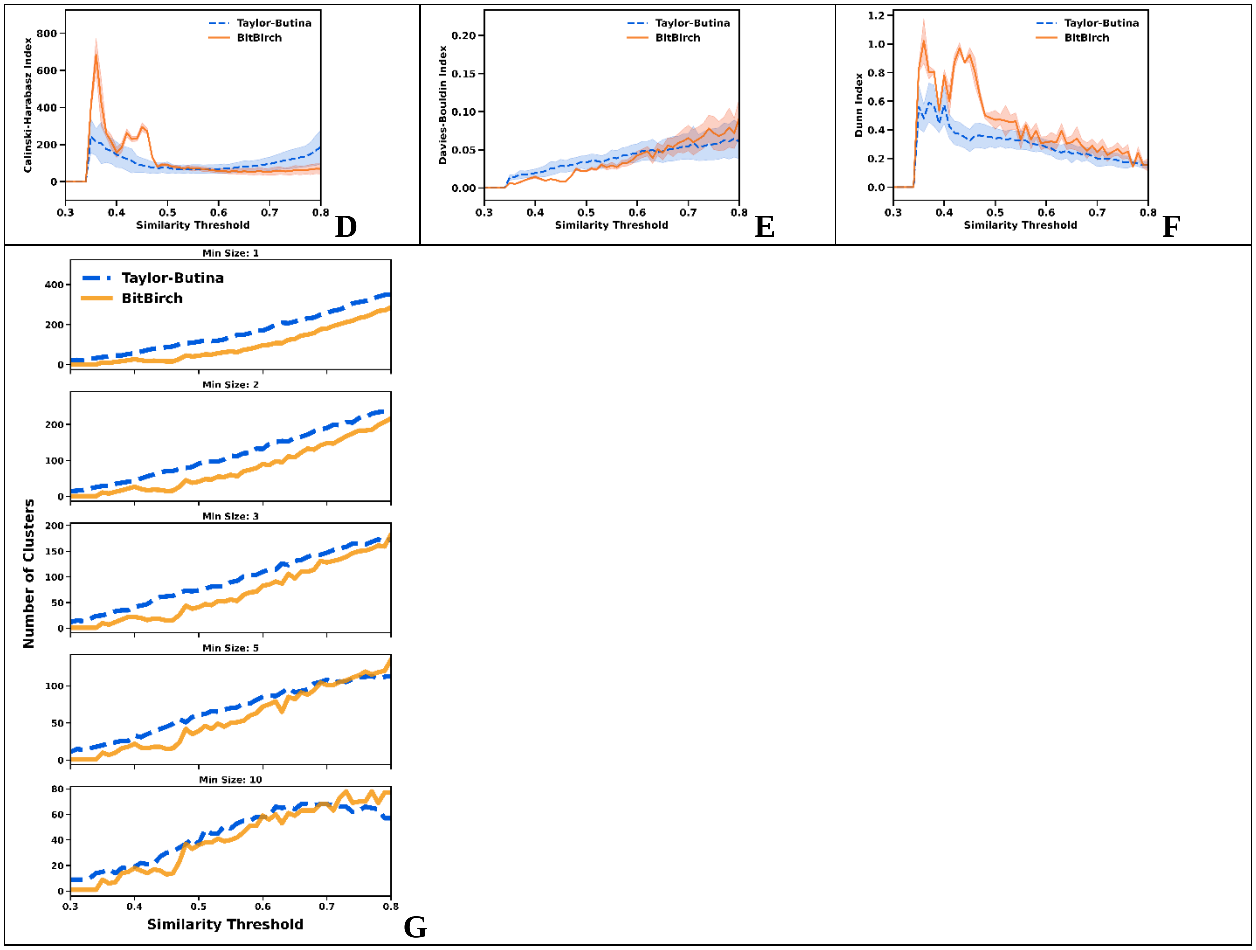


**Figure S4.30**: **A**: CHI, **B**: DBI, **C**: DI analysis for min_size = 1, 2, 3, 5, 10 for the [30] ChEMBL subset. Average **D**: CHI, **E**: DBI, **F**: DI values over the min_size variable. **G**: Number of clusters. BitBIRCH (orange continuous line), Taylor-Butina (blue dashed line).

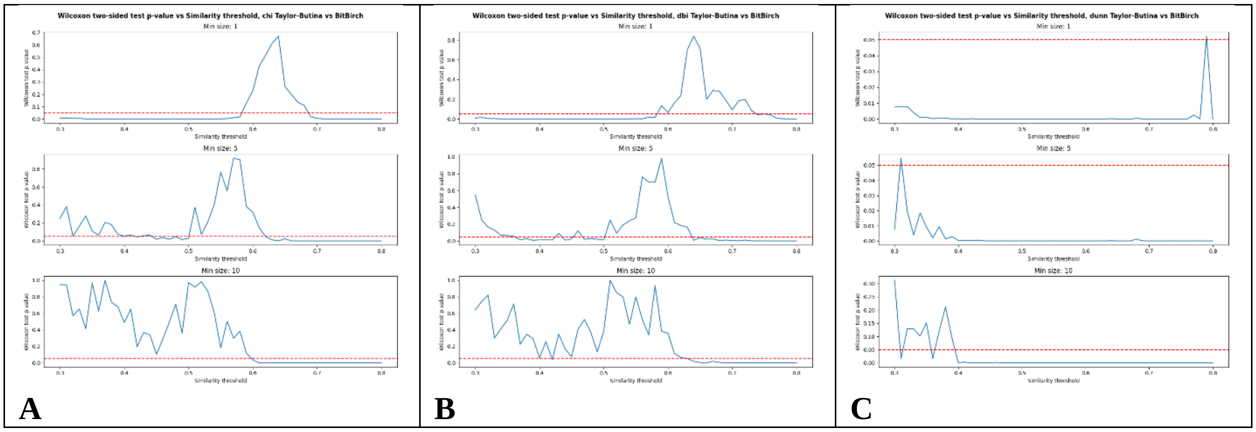

**Figure S4.31**: Wilcoxon two-sided (Ha: TB ≠ BB) test comparing the BitBIRCH and Taylor-Butina **A**: CHI, **B**: DBI, and **C**: DI results for different similarity thresholds and min_size (1, 5, 10) values.

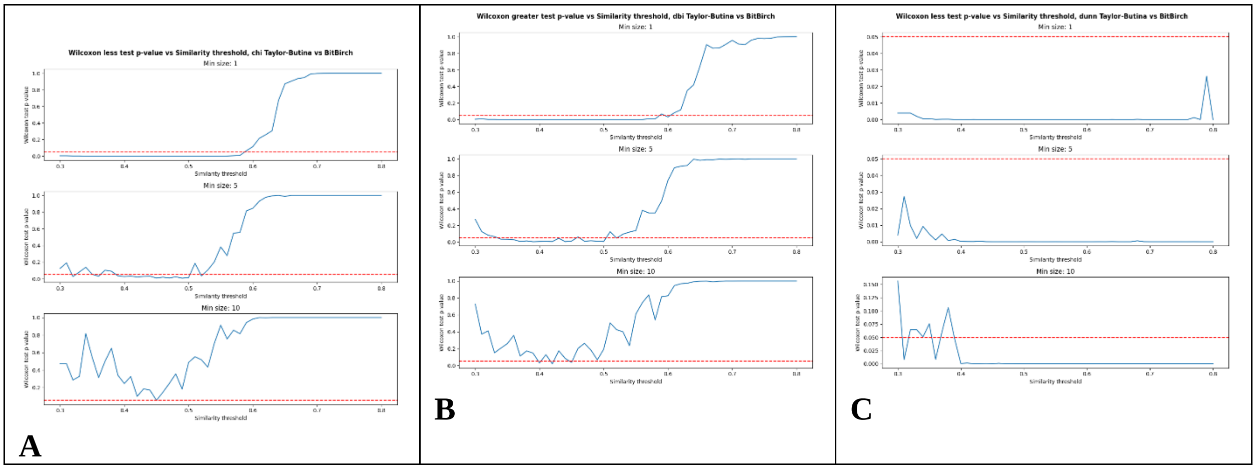


**Figure S4.32**: Wilcoxon one-sided test comparing the BitBIRCH and Taylor-Butina **A**: CHI (Ha: TB < BB), **B**: DBI (Ha: TB > BB), and **C**: DI (Ha: TB < BB), results for different similarity thresholds and min_size (1, 5, 10) values.

### S5: Local Clustering Analysis of BitBIRCH and BitBIRCH-parallel

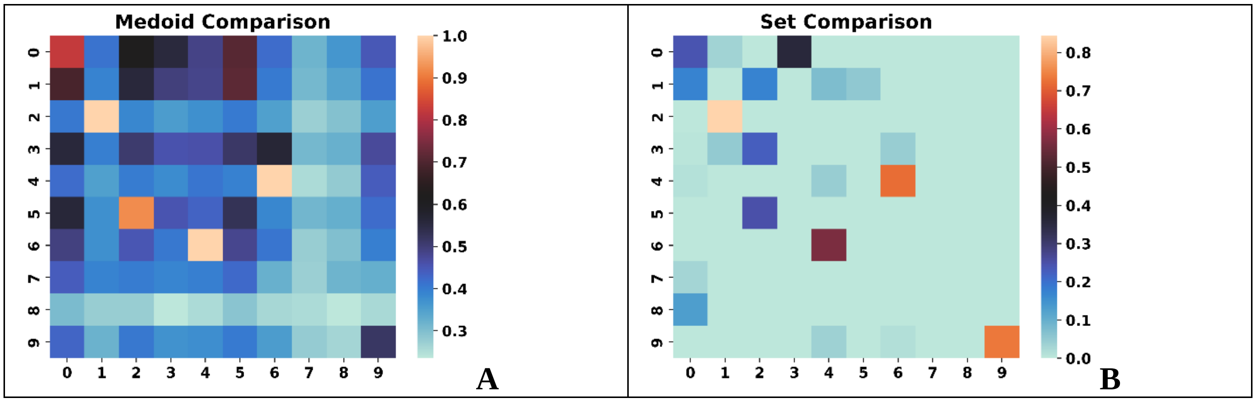


**Figure S5.1**: Comparison of the **A**: medoids, **B**: sets for the top populated clusters of the [1] ChEMBL subset using the BitBIRCH and BitBIRCH-parallel algorithms (similarity threshold = 0.65, min_size = 10).

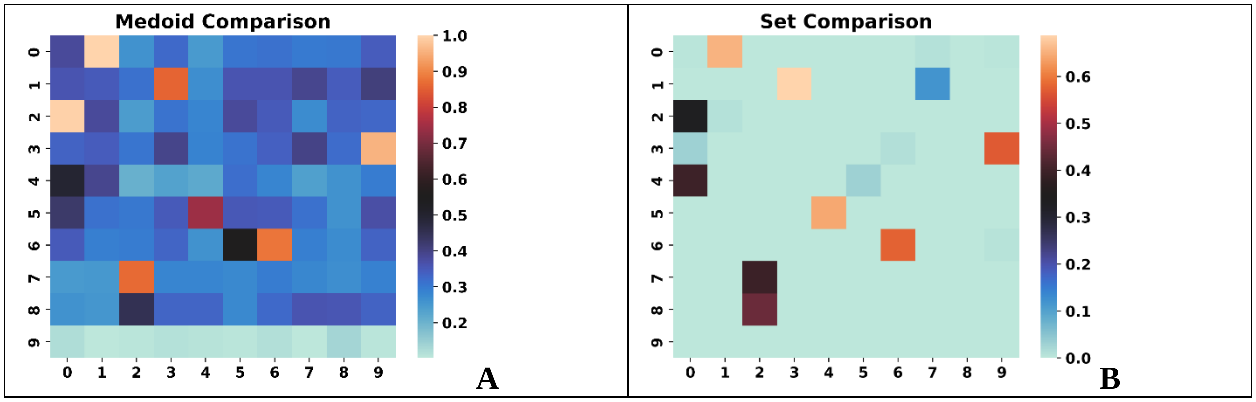


**Figure S5.2**: Comparison of the **A**: medoids, **B**: sets for the top populated clusters of the [2] ChEMBL subset using the BitBIRCH and BitBIRCH-parallel algorithms (similarity threshold = 0.65, min_size = 10).

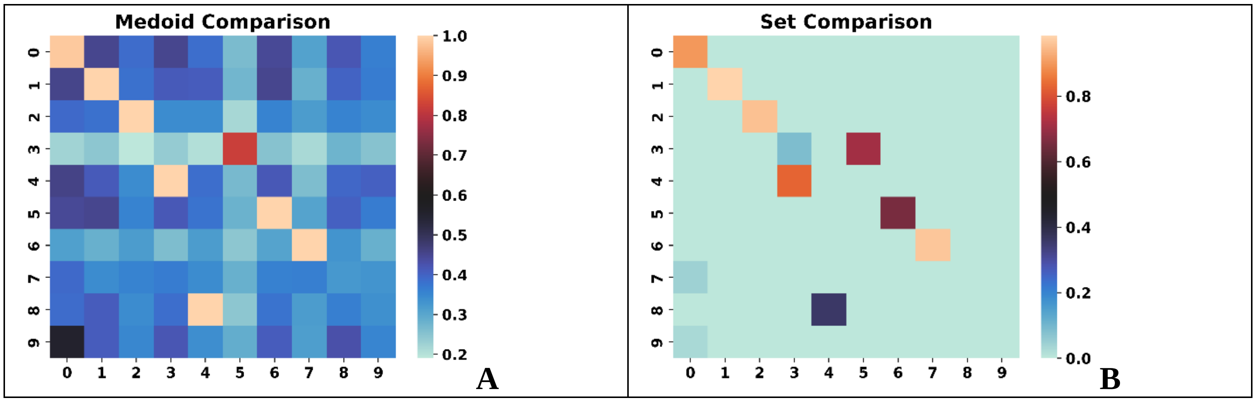


**Figure S5.3**: Comparison of the **A**: medoids, **B**: sets for the top populated clusters of the [3] ChEMBL subset using the BitBIRCH and BitBIRCH-parallel algorithms (similarity threshold = 0.65, min_size = 10).

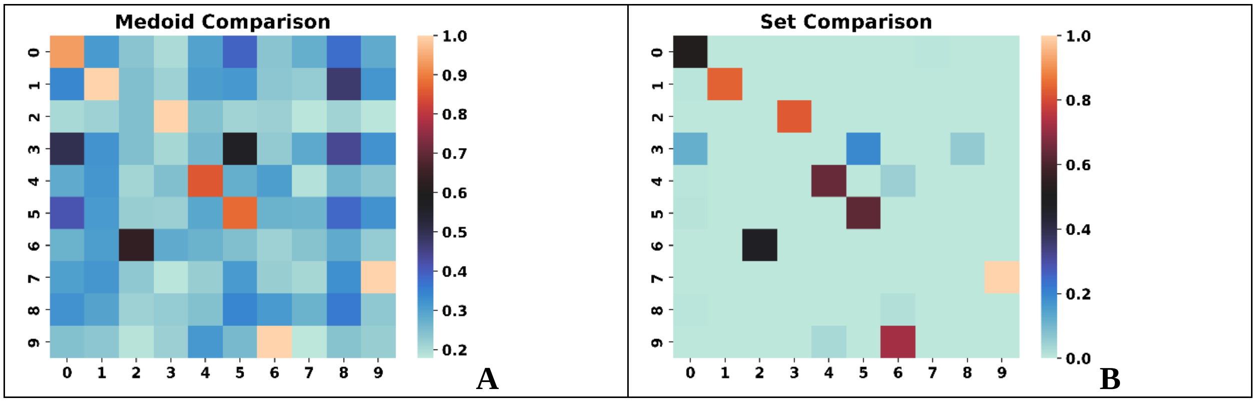

**Figure S5.4**: Comparison of the **A**: medoids, **B**: sets for the top populated clusters of the [4] ChEMBL subset using the BitBIRCH and BitBIRCH-parallel algorithms (similarity threshold = 0.65, min_size = 10).

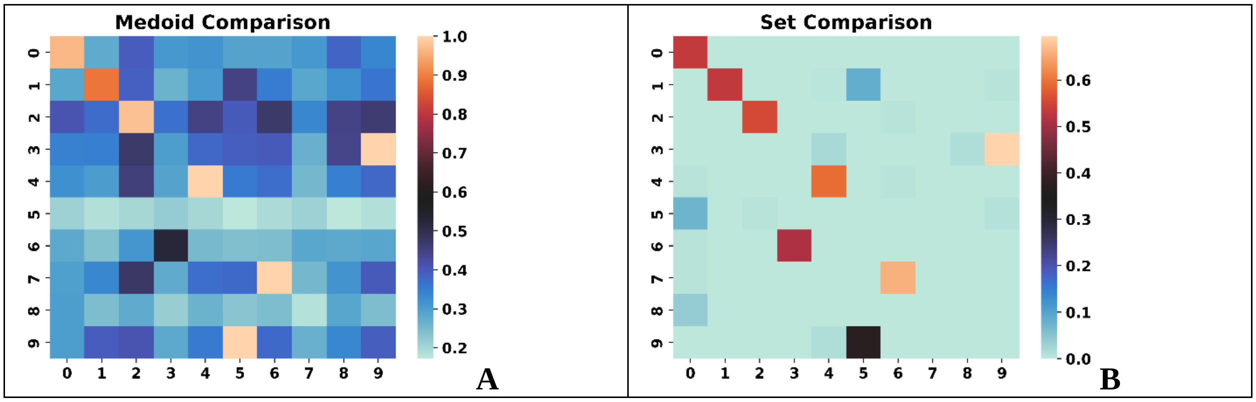


**Figure S5.5**: Comparison of the **A**: medoids, **B**: sets for the top populated clusters of the [5] ChEMBL subset using the BitBIRCH and BitBIRCH-parallel algorithms (similarity threshold = 0.65, min_size = 10).

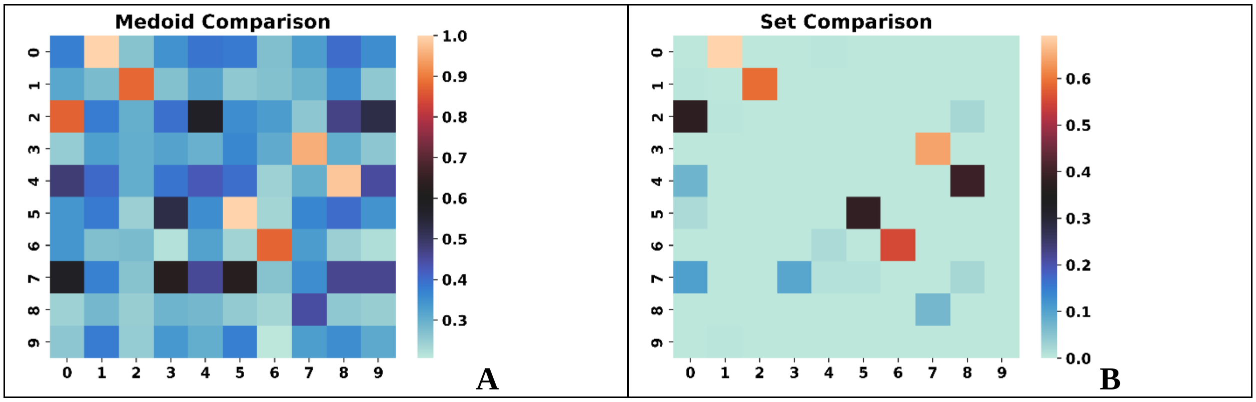


**Figure S5.6**: Comparison of the **A**: medoids, **B**: sets for the top populated clusters of the [6] ChEMBL subset using the BitBIRCH and BitBIRCH-parallel algorithms (similarity threshold = 0.65, min_size = 10).

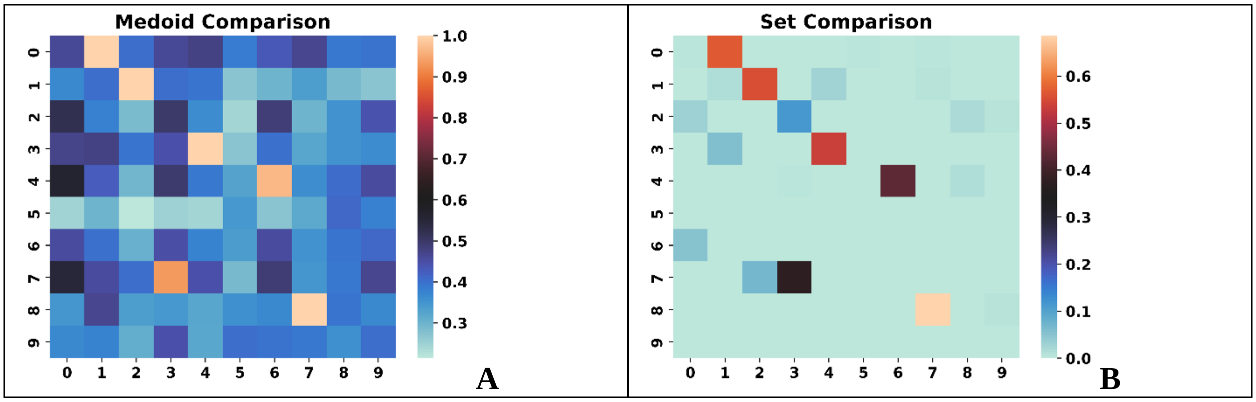


**Figure S5.7**: Comparison of the **A**: medoids, **B**: sets for the top populated clusters of the [7] ChEMBL subset using the BitBIRCH and BitBIRCH-parallel algorithms (similarity threshold = 0.65, min_size = 10).

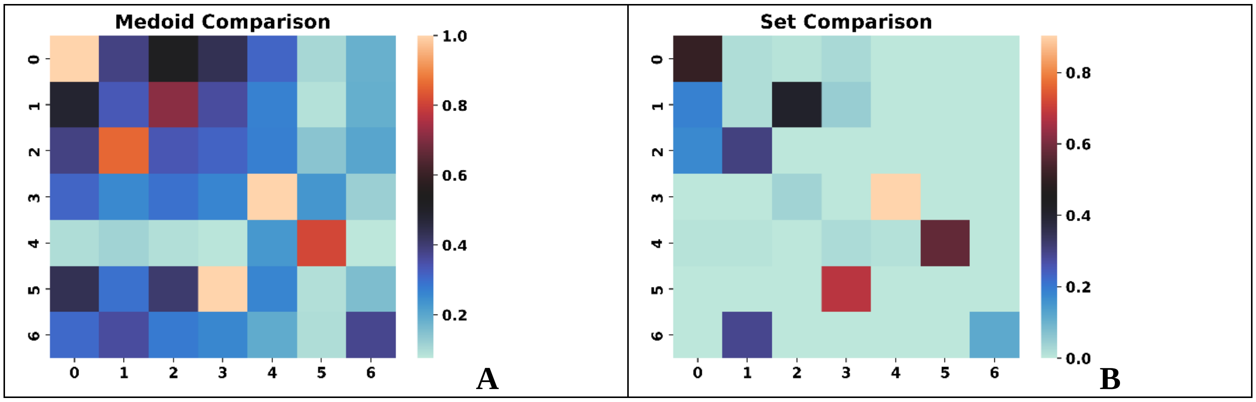


**Figure S5.8**: Comparison of the **A**: medoids, **B**: sets for the top populated clusters of the [8] ChEMBL subset using the BitBIRCH and BitBIRCH-parallel algorithms (similarity threshold = 0.65, min_size = 10).

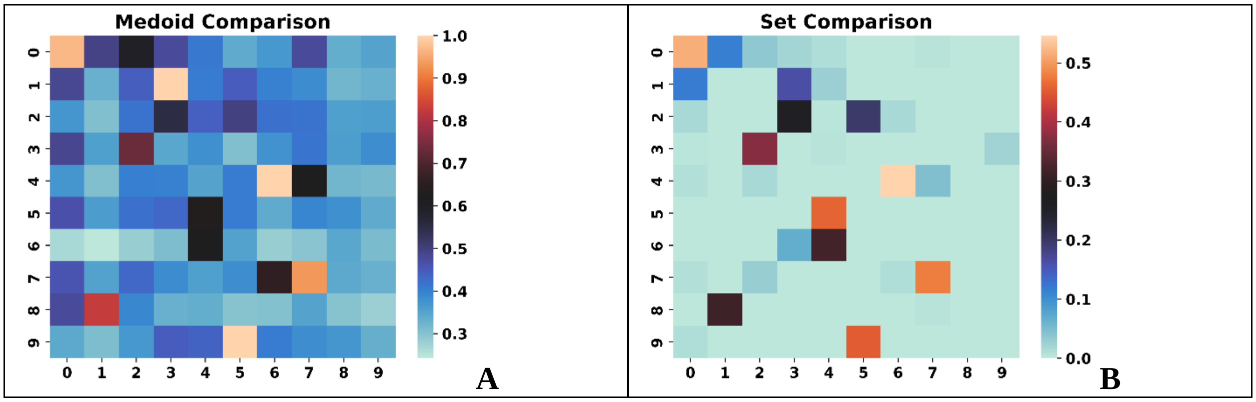

**Figure S5.9**: Comparison of the **A**: medoids, **B**: sets for the top populated clusters of the [9] ChEMBL subset using the BitBIRCH and BitBIRCH-parallel algorithms (similarity threshold = 0.65, min_size = 10).

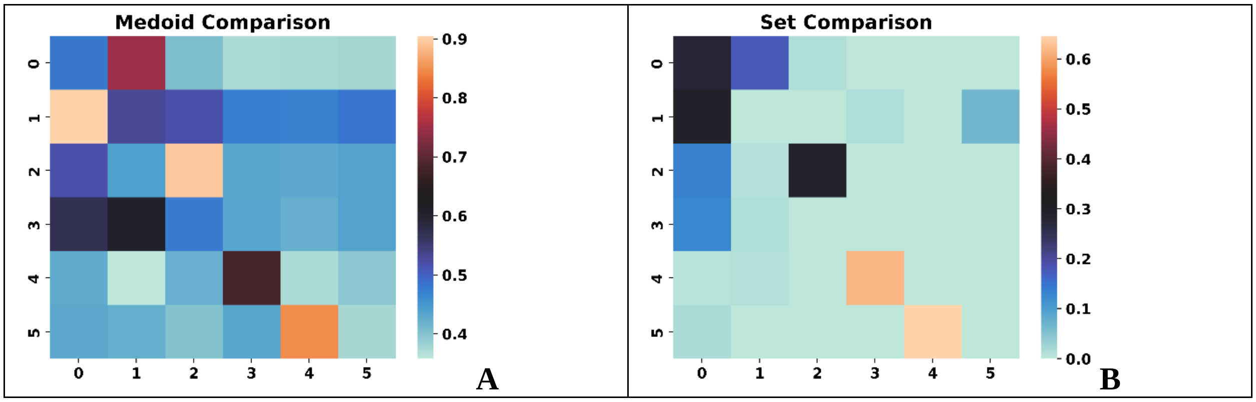


**Figure S5.10**: Comparison of the **A**: medoids, **B**: sets for the top populated clusters of the [10] ChEMBL subset using the BitBIRCH and BitBIRCH-parallel algorithms (similarity threshold = 0.65, min_size = 10).

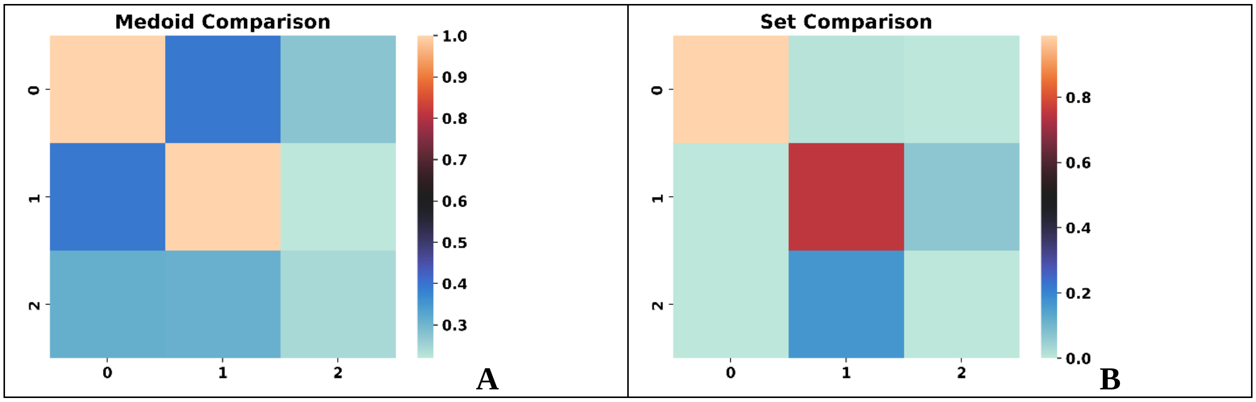


**Figure S5.11**: Comparison of the **A**: medoids, **B**: sets for the top populated clusters of the [11] ChEMBL subset using the BitBIRCH and BitBIRCH-parallel algorithms (similarity threshold = 0.65, min_size = 10).

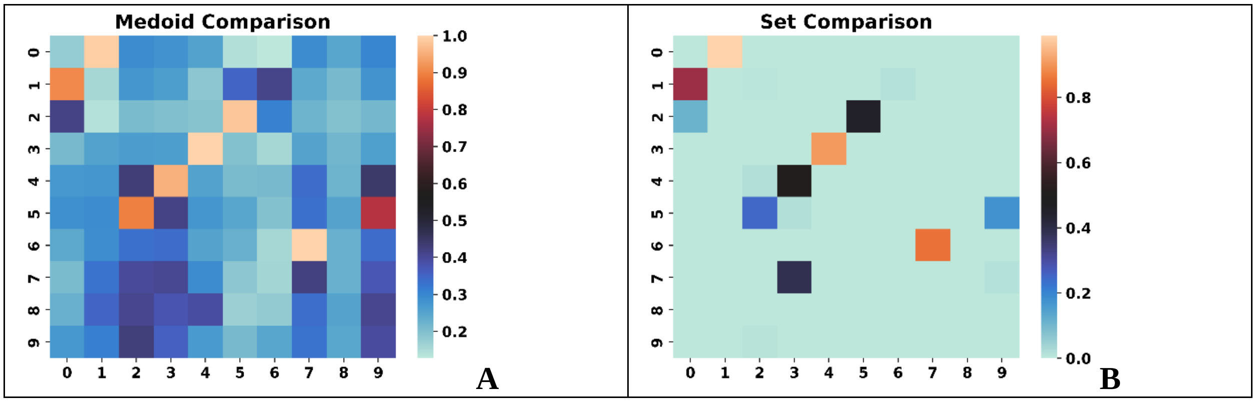


**Figure S5.12**: Comparison of the **A**: medoids, **B**: sets for the top populated clusters of the [12] ChEMBL subset using the BitBIRCH and BitBIRCH-parallel algorithms (similarity threshold = 0.65, min_size = 10).

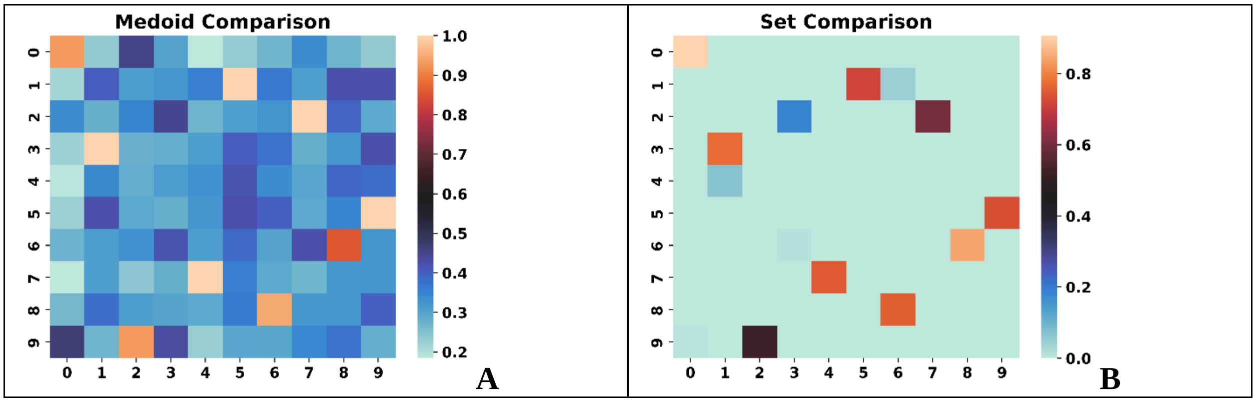


**Figure S5.13**: Comparison of the **A**: medoids, **B**: sets for the top populated clusters of the [13] ChEMBL subset using the BitBIRCH and BitBIRCH-parallel algorithms (similarity threshold = 0.65, min_size = 10).

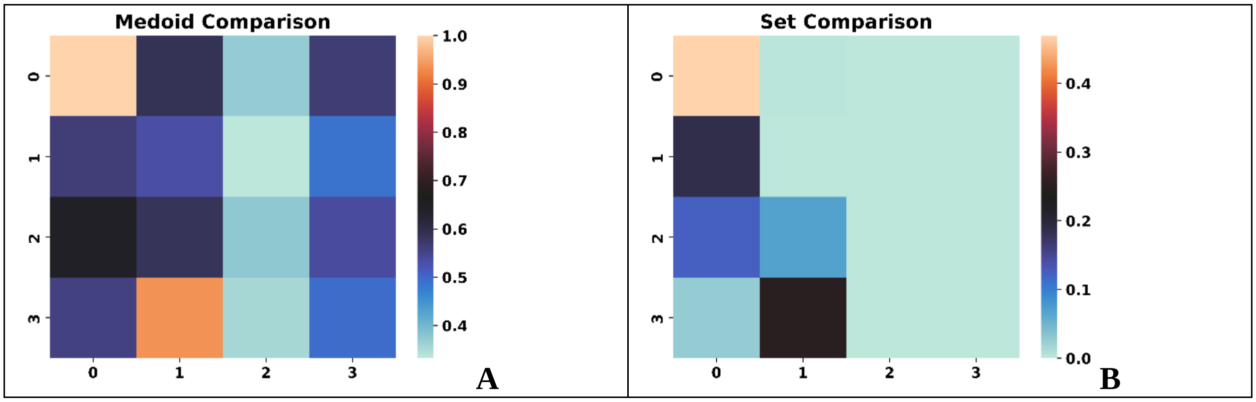

**Figure S5.14**: Comparison of the **A**: medoids, **B**: sets for the top populated clusters of the [14] ChEMBL subset using the BitBIRCH and BitBIRCH-parallel algorithms (similarity threshold = 0.65, min_size = 10).

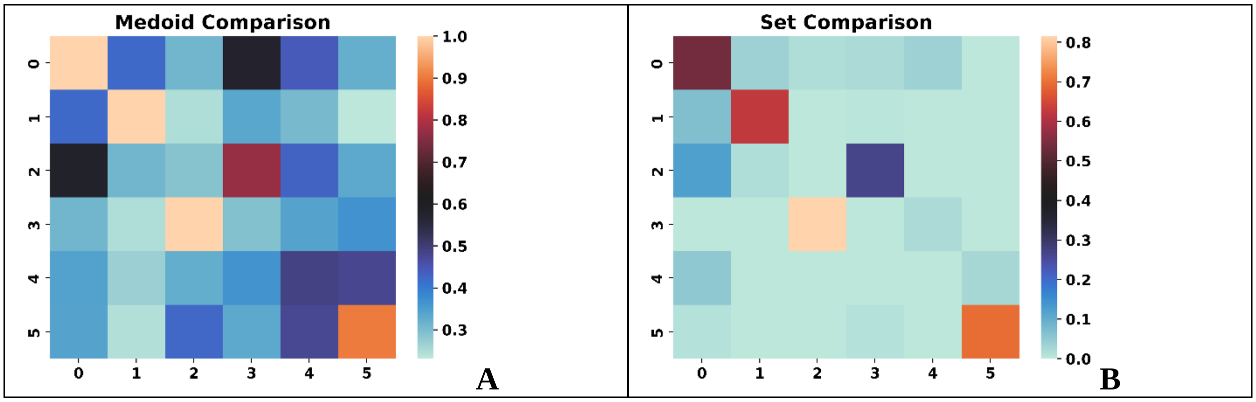


**Figure S5.15**: Comparison of the **A**: medoids, **B**: sets for the top populated clusters of the [15] ChEMBL subset using the BitBIRCH and BitBIRCH-parallel algorithms (similarity threshold = 0.65, min_size = 10).

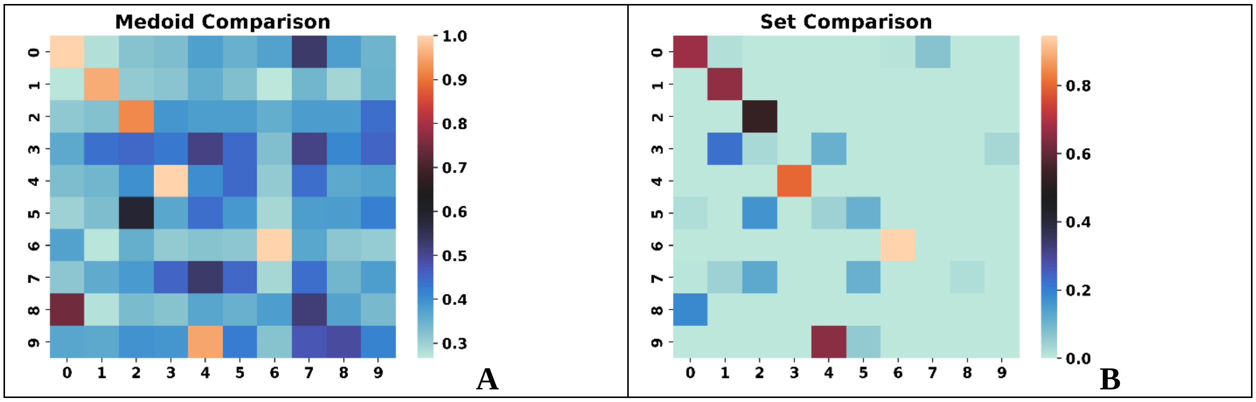


**Figure S5.16**: Comparison of the **A**: medoids, **B**: sets for the top populated clusters of the [16] ChEMBL subset using the BitBIRCH and BitBIRCH-parallel algorithms (similarity threshold = 0.65, min_size = 10).

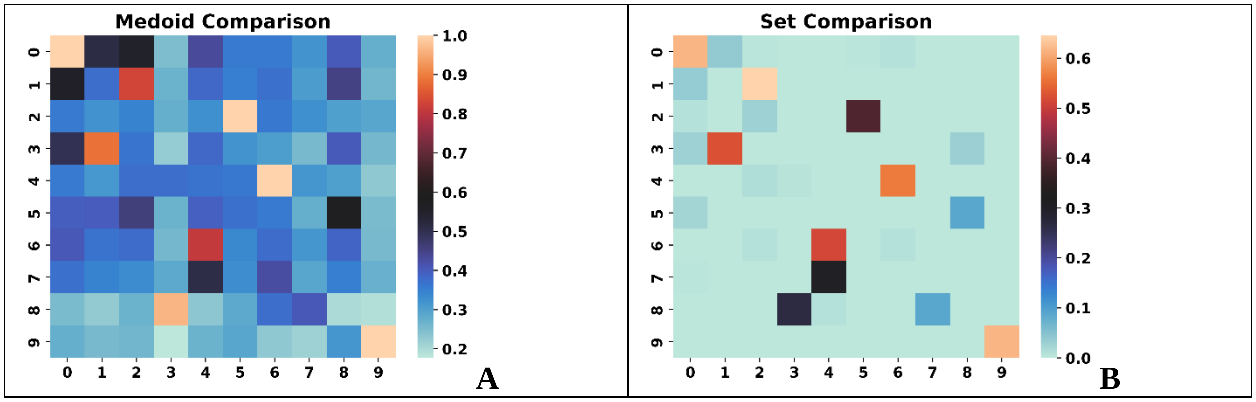


**Figure S5.17**: Comparison of the **A**: medoids, **B**: sets for the top populated clusters of the [17] ChEMBL subset using the BitBIRCH and BitBIRCH-parallel algorithms (similarity threshold = 0.65, min_size = 10).

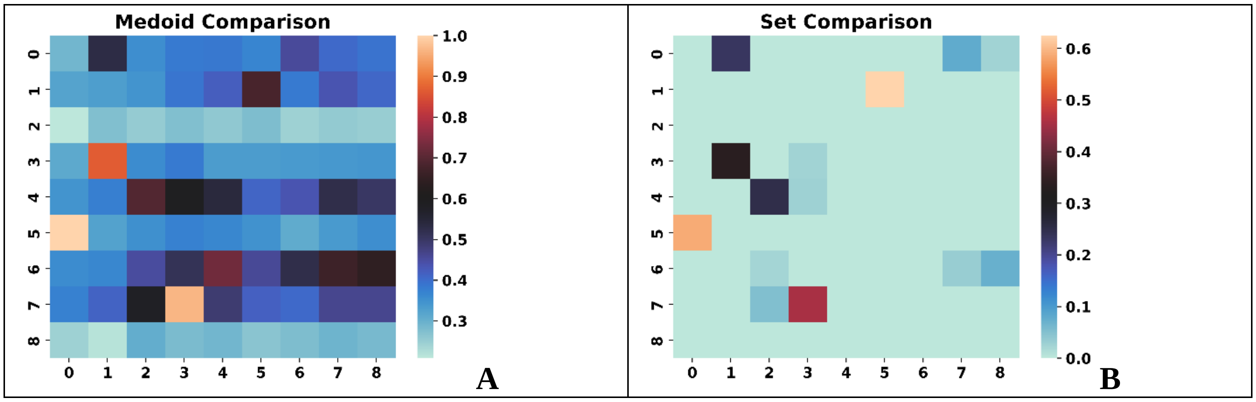


**Figure S5.18**: Comparison of the **A**: medoids, **B**: sets for the top populated clusters of the [18] ChEMBL subset using the BitBIRCH and BitBIRCH-parallel algorithms (similarity threshold = 0.65, min_size = 10).

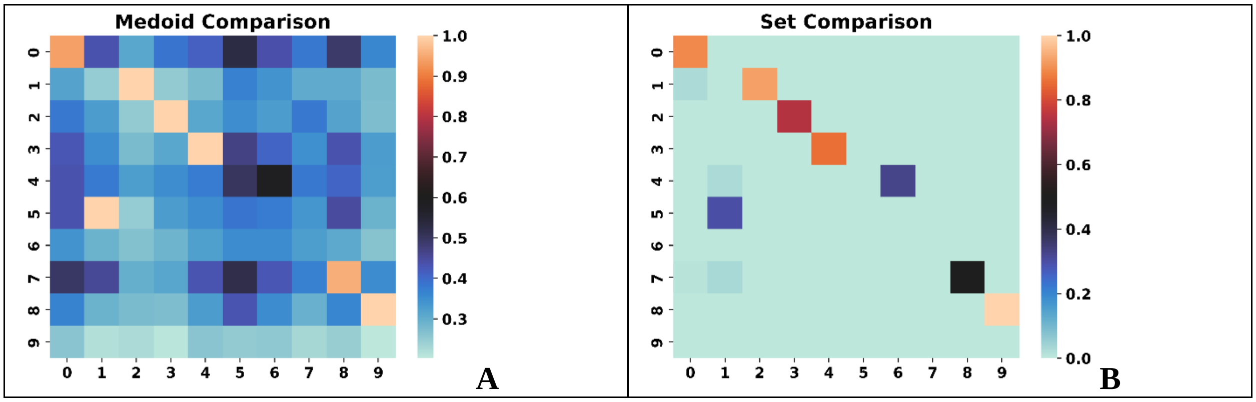

**Figure S5.19**: Comparison of the **A**: medoids, **B**: sets for the top populated clusters of the [19] ChEMBL subset using the BitBIRCH and BitBIRCH-parallel algorithms (similarity threshold = 0.65, min_size = 10).

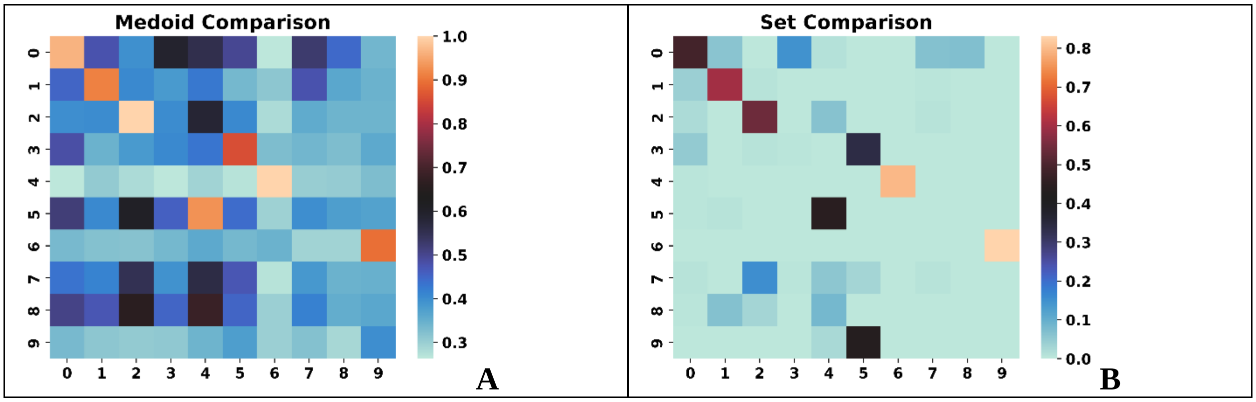


**Figure S5.20**: Comparison of the **A**: medoids, **B**: sets for the top populated clusters of the [20] ChEMBL subset using the BitBIRCH and BitBIRCH-parallel algorithms (similarity threshold = 0.65, min_size = 10).

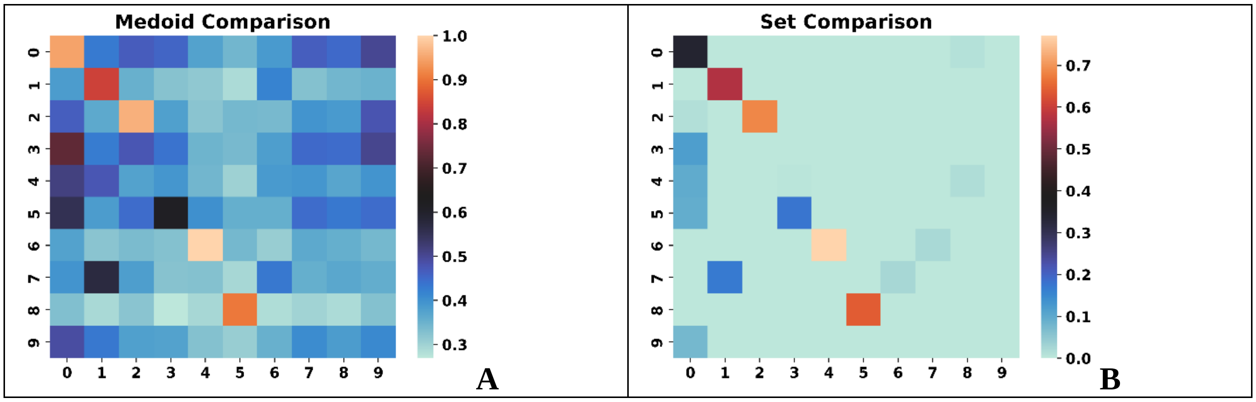


**Figure S5.21**: Comparison of the **A**: medoids, **B**: sets for the top populated clusters of the [21] ChEMBL subset using the BitBIRCH and BitBIRCH-parallel algorithms (similarity threshold = 0.65, min_size = 10).

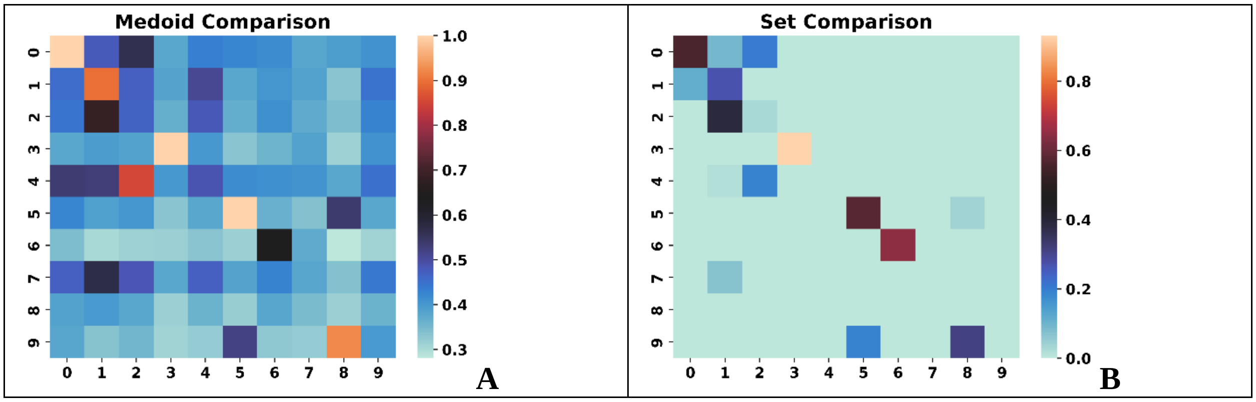


**Figure S5.22**: Comparison of the **A**: medoids, **B**: sets for the top populated clusters of the [22] ChEMBL subset using the BitBIRCH and BitBIRCH-parallel algorithms (similarity threshold = 0.65, min_size = 10).

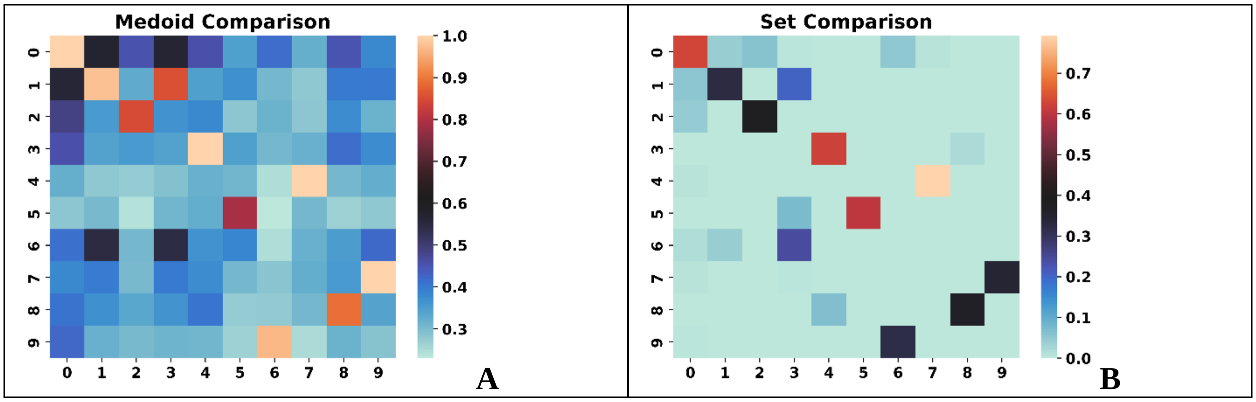


**Figure S5.23**: Comparison of the **A**: medoids, **B**: sets for the top populated clusters of the [23] ChEMBL subset using the BitBIRCH and BitBIRCH-parallel algorithms (similarity threshold = 0.65, min_size = 10).

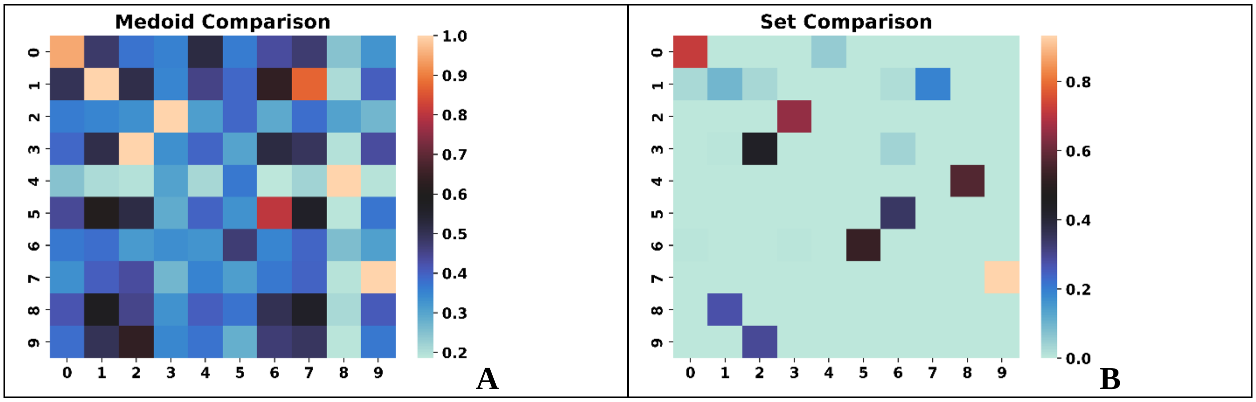

**Figure S5.24**: Comparison of the **A**: medoids, **B**: sets for the top populated clusters of the [24] ChEMBL subset using the BitBIRCH and BitBIRCH-parallel algorithms (similarity threshold = 0.65, min_size = 10).

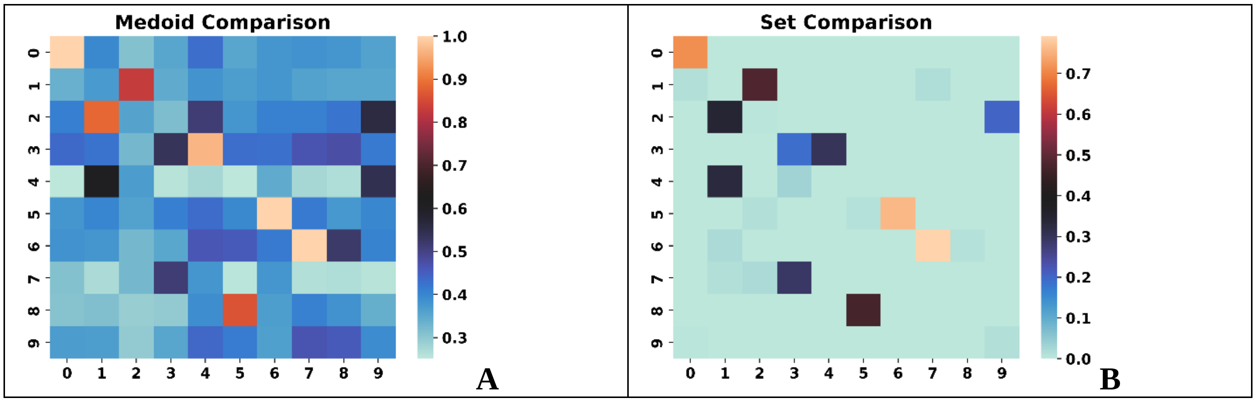


**Figure S5.25**: Comparison of the **A**: medoids, **B**: sets for the top populated clusters of the [25] ChEMBL subset using the BitBIRCH and BitBIRCH-parallel algorithms (similarity threshold = 0.65, min_size = 10).

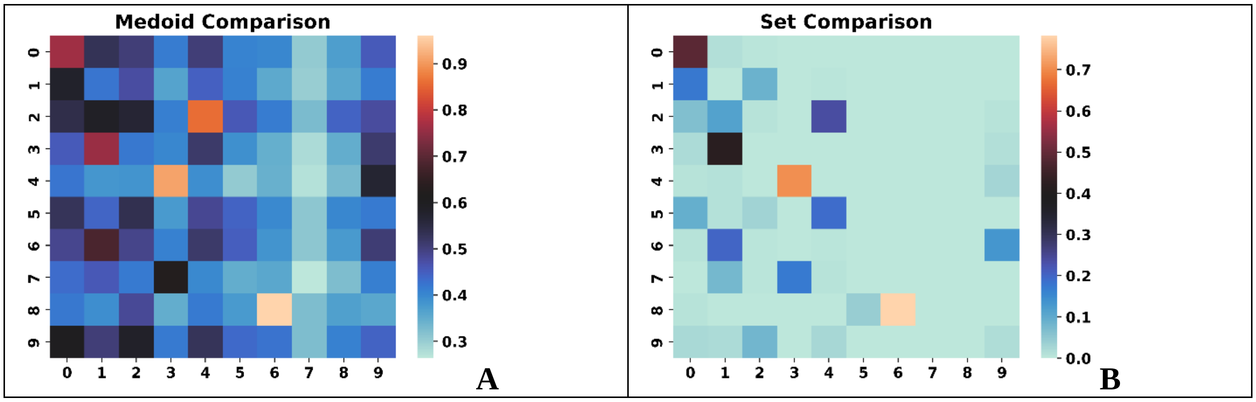


**Figure S5.26**: Comparison of the **A**: medoids, **B**: sets for the top populated clusters of the [26] ChEMBL subset using the BitBIRCH and BitBIRCH-parallel algorithms (similarity threshold = 0.65, min_size = 10).

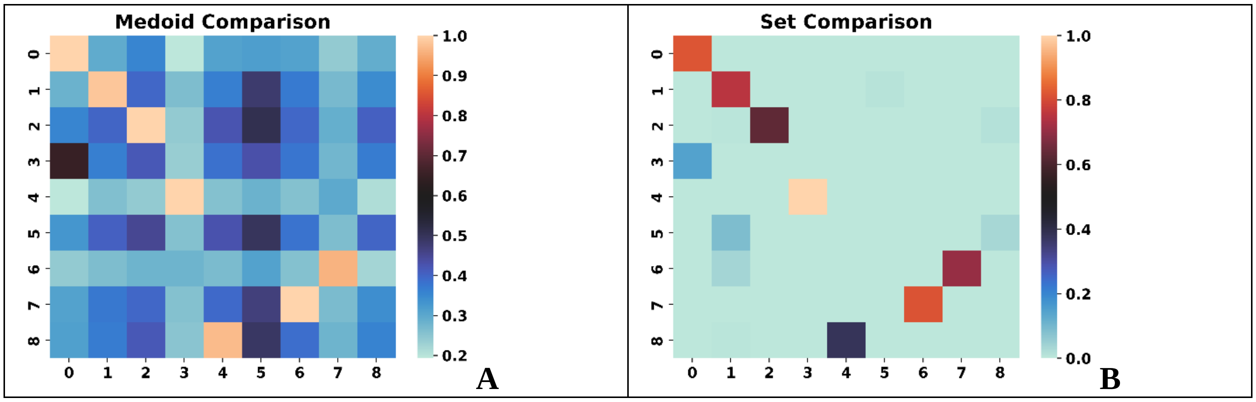


**Figure S5.27**: Comparison of the **A**: medoids, **B**: sets for the top populated clusters of the [27] ChEMBL subset using the BitBIRCH and BitBIRCH-parallel algorithms (similarity threshold = 0.65, min_size = 10).

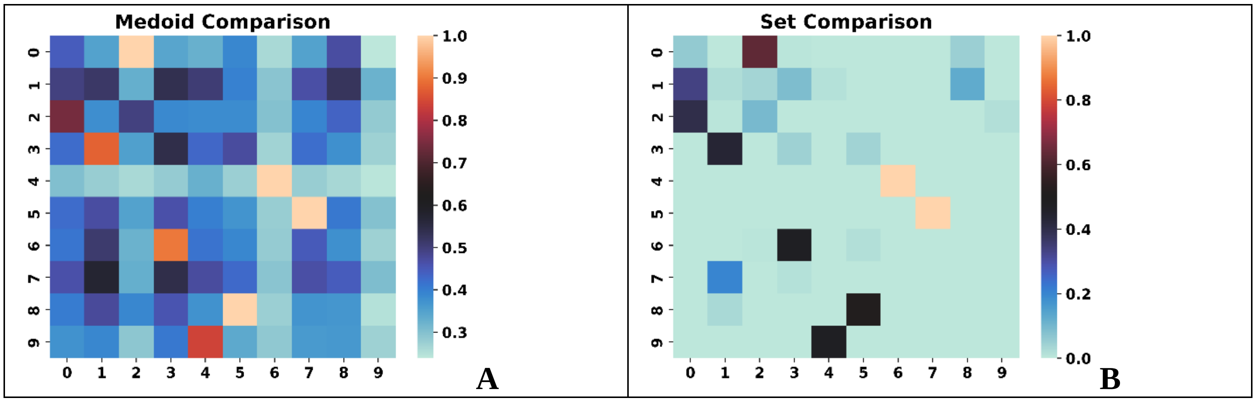


**Figure S5.28**: Comparison of the **A**: medoids, **B**: sets for the top populated clusters of the [28] ChEMBL subset using the BitBIRCH and BitBIRCH-parallel algorithms (similarity threshold = 0.65, min_size = 10).

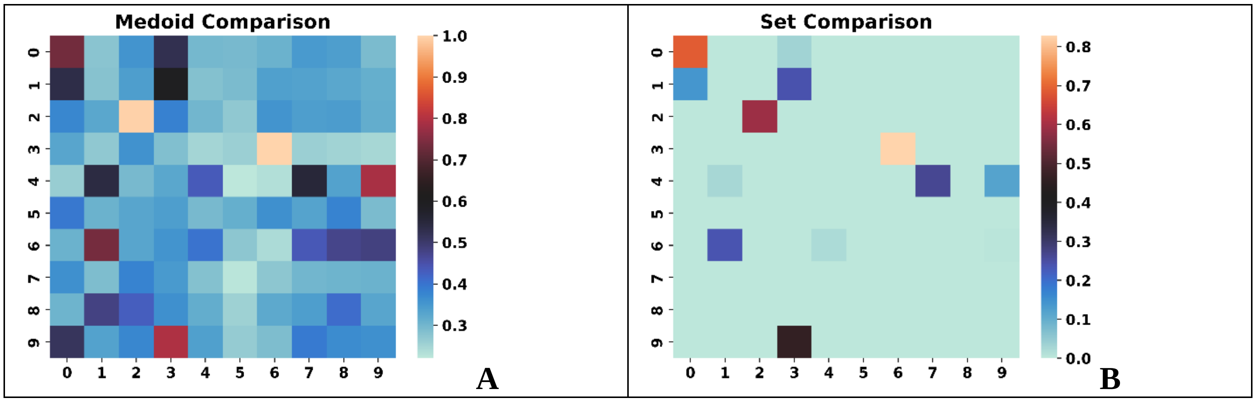

**Figure S5.29**: Comparison of the **A**: medoids, **B**: sets for the top populated clusters of the [29] ChEMBL subset using the BitBIRCH and BitBIRCH-parallel algorithms (similarity threshold = 0.65, min_size = 10).

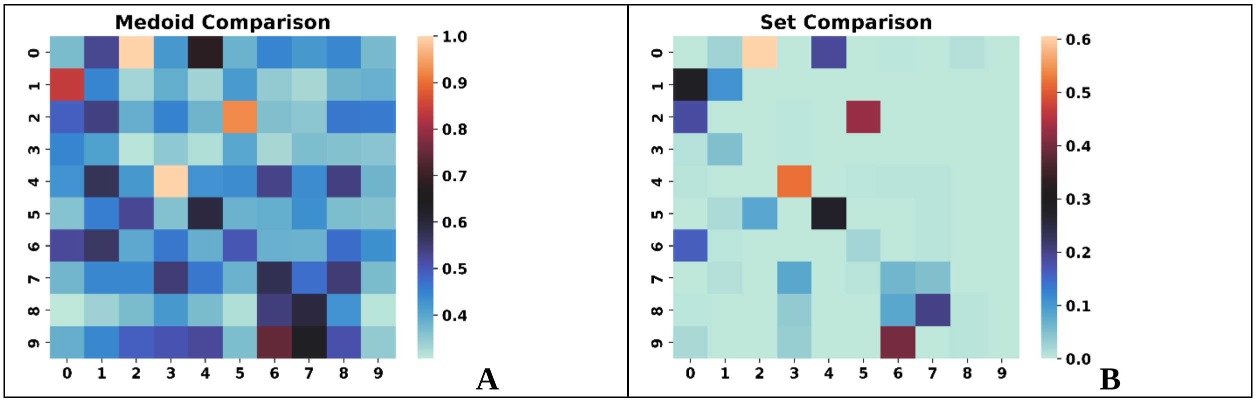


**Figure S5.30**: Comparison of the **A**: medoids, **B**: sets for the top populated clusters of the [30] ChEMBL subset using the BitBIRCH and BitBIRCH-parallel algorithms (similarity threshold = 0.65, min_size = 10).

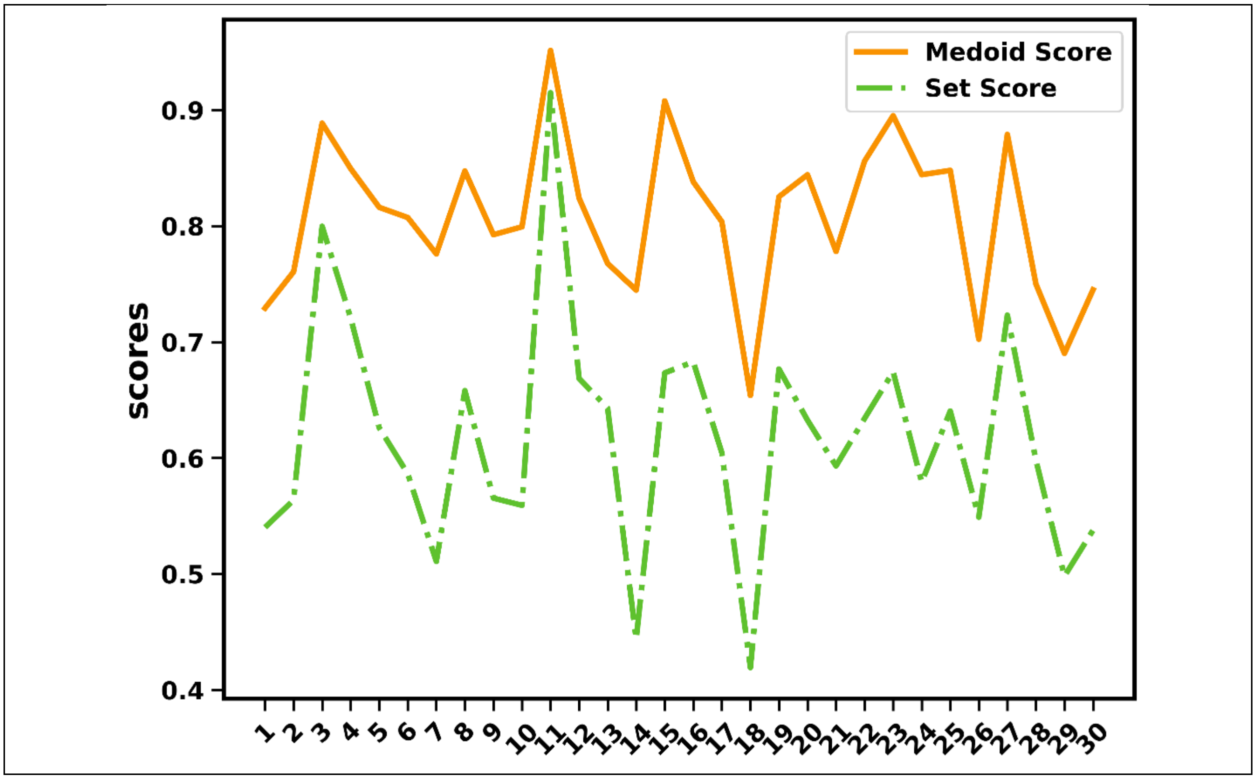


**Figure S5.31**: Summary of the Medoid (continuous orange line) and Set (dashed green line) scores for the 30 ChEMBL subsets for the BitBIRCH and BitBIRCH-parallel algorithms.

| | CHI | DBI | DI |
|---|---|---|---|
| *t*-statistic | 163.0 | 195.0 | 41.0 |
| *p*-value | 0.158 | 0.452 | 1.824e-05 |

**Table S5.1**: *t*-statistic and *p*-value for the two-sided Wilcoxon test comparing the BitBIRCH and BitBIRCH-parallel methods for the 30 ChEMBL subsets at a 0.65 similarity threshold, with min_size = 10.

We also performed a one-sided Dunn test (Ha: BB < BB-parallel), that resulted in a *t*-statistic of 41.0 and a *p*-value of 9.122e-05.

### S6: Local Clustering Analysis of BitBIRCH and BitBIRCH-folded

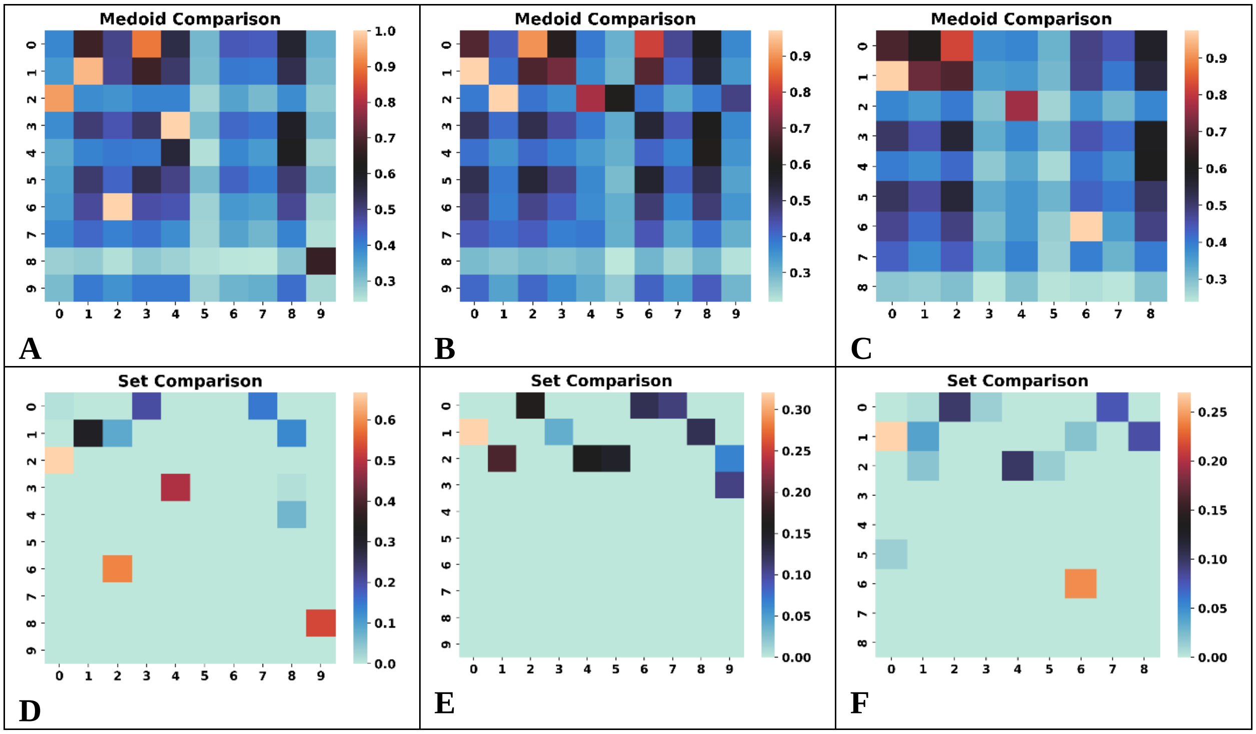


**Figure S6.1**: Comparison of the **A, B, C**: medoids, **D, E, F**: sets for the top populated clusters of the [1] ChEMBL subset using the BitBIRCH and BitBIRCH-folded (**A**, **D**: 1- fold, **B**, **E**: 2-fold, **C**, **F**: 3-fold) algorithms (similarity threshold = 0.65, min_size = 10).

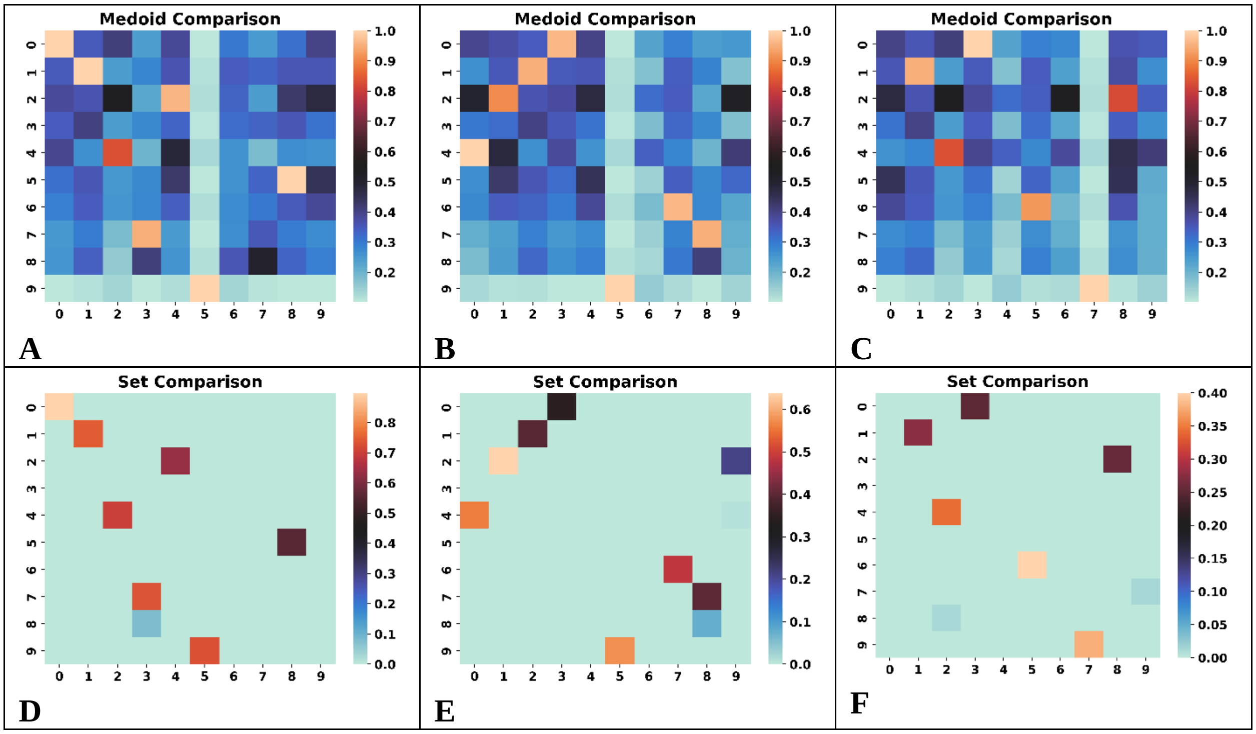


**Figure S6.2**: Comparison of the **A, B, C**: medoids, **D, E, F**: sets for the top populated clusters of the [2] ChEMBL subset using the BitBIRCH and BitBIRCH-folded (**A**, **D**: 1- fold, **B**, **E**: 2-fold, **C**, **F**: 3-fold) algorithms (similarity threshold = 0.65, min_size = 10).

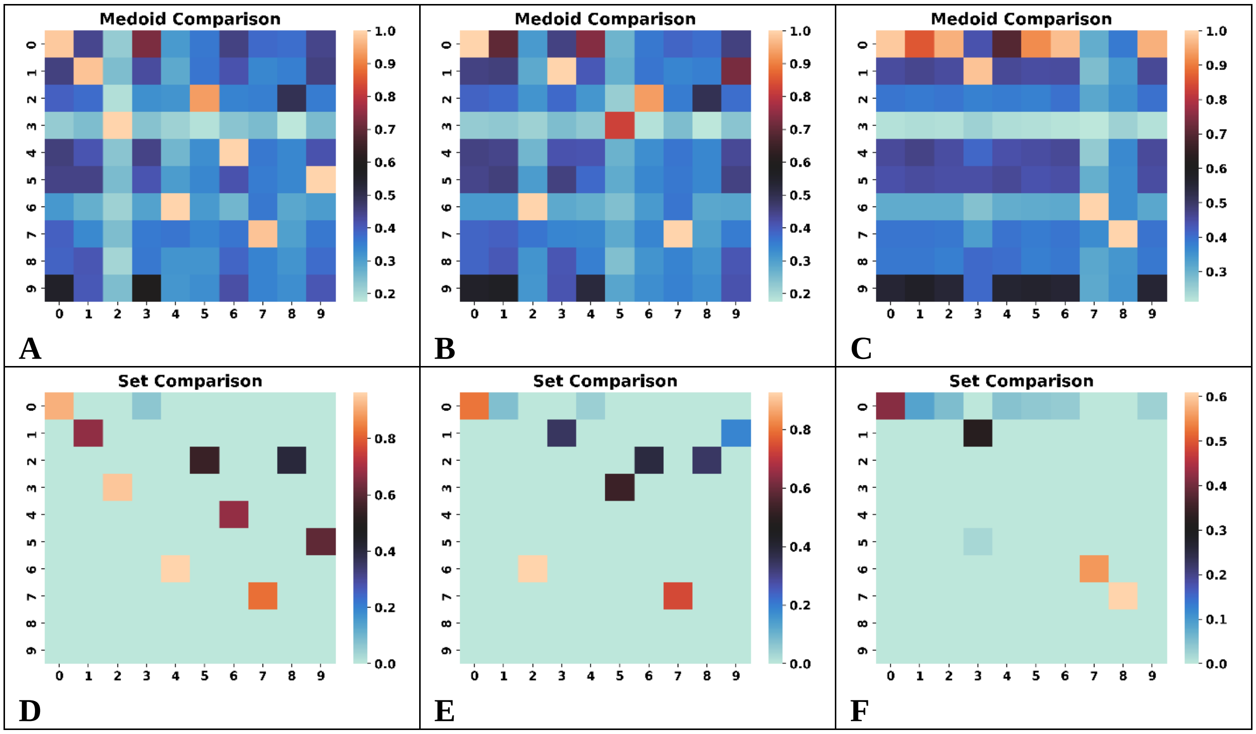

**Figure S6.3**: Comparison of the **A, B, C**: medoids, **D, E, F**: sets for the top populated clusters of the [3] ChEMBL subset using the BitBIRCH and BitBIRCH-folded (**A**, **D**: 1- fold, **B**, **E**: 2-fold, **C**, **F**: 3-fold) algorithms (similarity threshold = 0.65, min_size = 10).

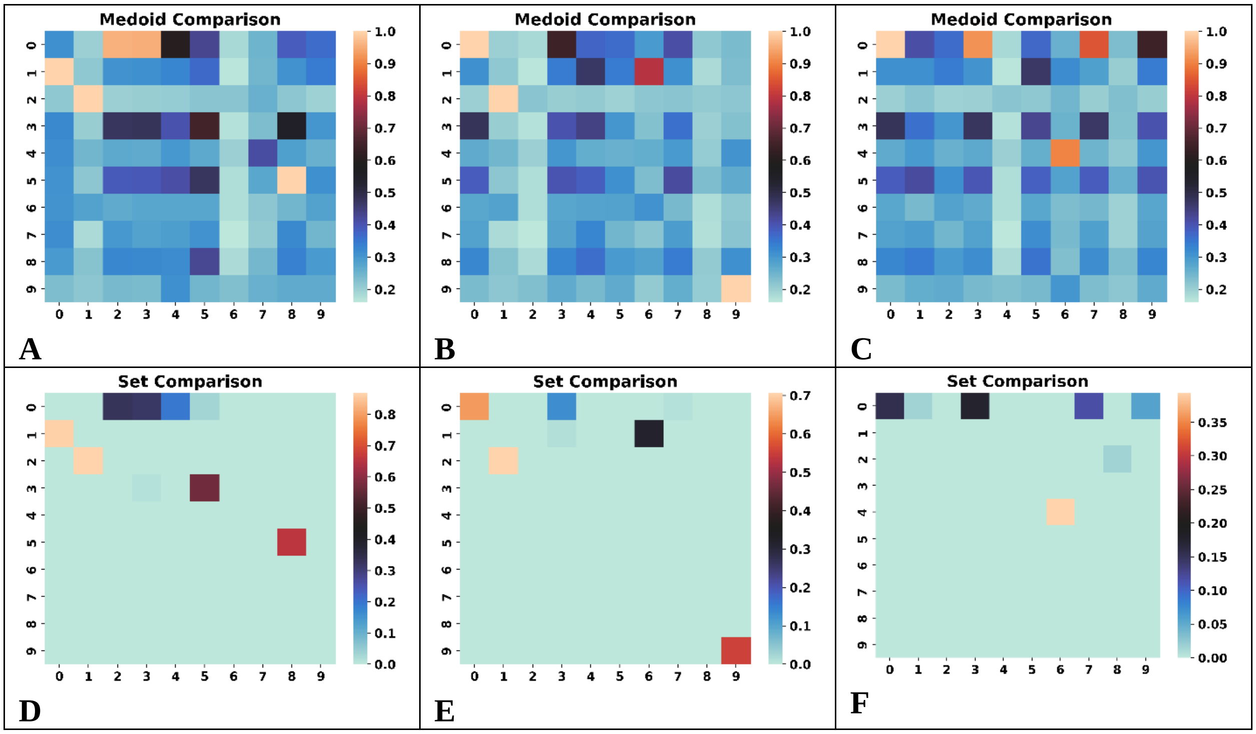


**Figure S6.4**: Comparison of the **A, B, C**: medoids, **D, E, F**: sets for the top populated clusters of the [4] ChEMBL subset using the BitBIRCH and BitBIRCH-folded (**A**, **D**: 1- fold, **B**, **E**: 2-fold, **C**, **F**: 3-fold) algorithms (similarity threshold = 0.65, min_size = 10).

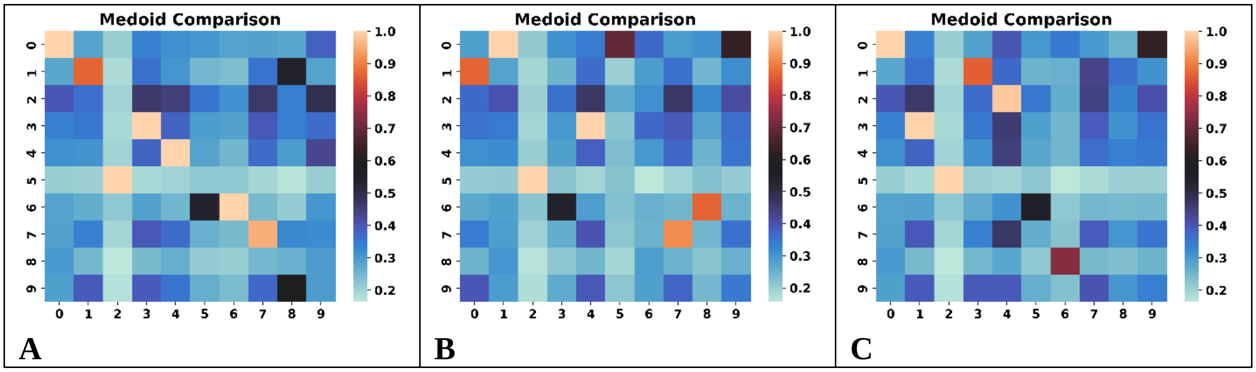

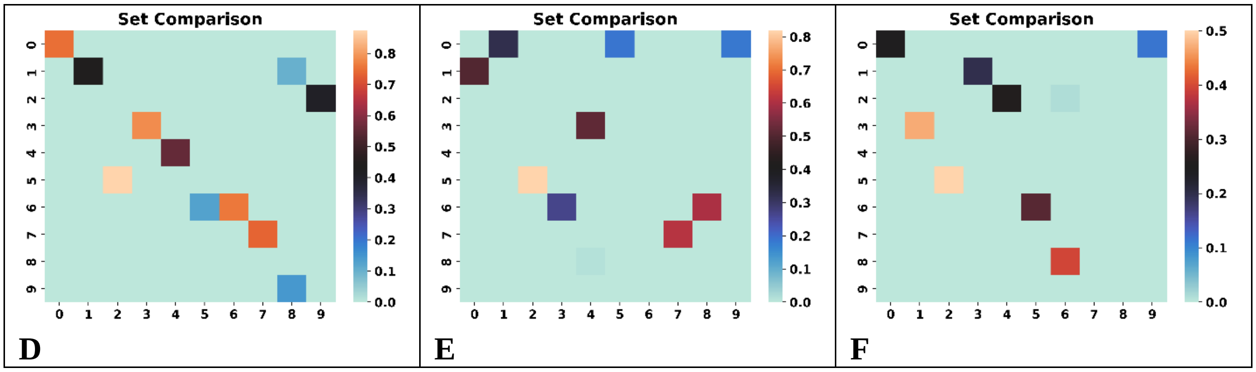


**Figure S6.5**: Comparison of the **A, B, C**: medoids, **D, E, F**: sets for the top populated clusters of the [5] ChEMBL subset using the BitBIRCH and BitBIRCH-folded (**A**, **D**: 1- fold, **B**, **E**: 2-fold, **C**, **F**: 3-fold) algorithms (similarity threshold = 0.65, min_size = 10).

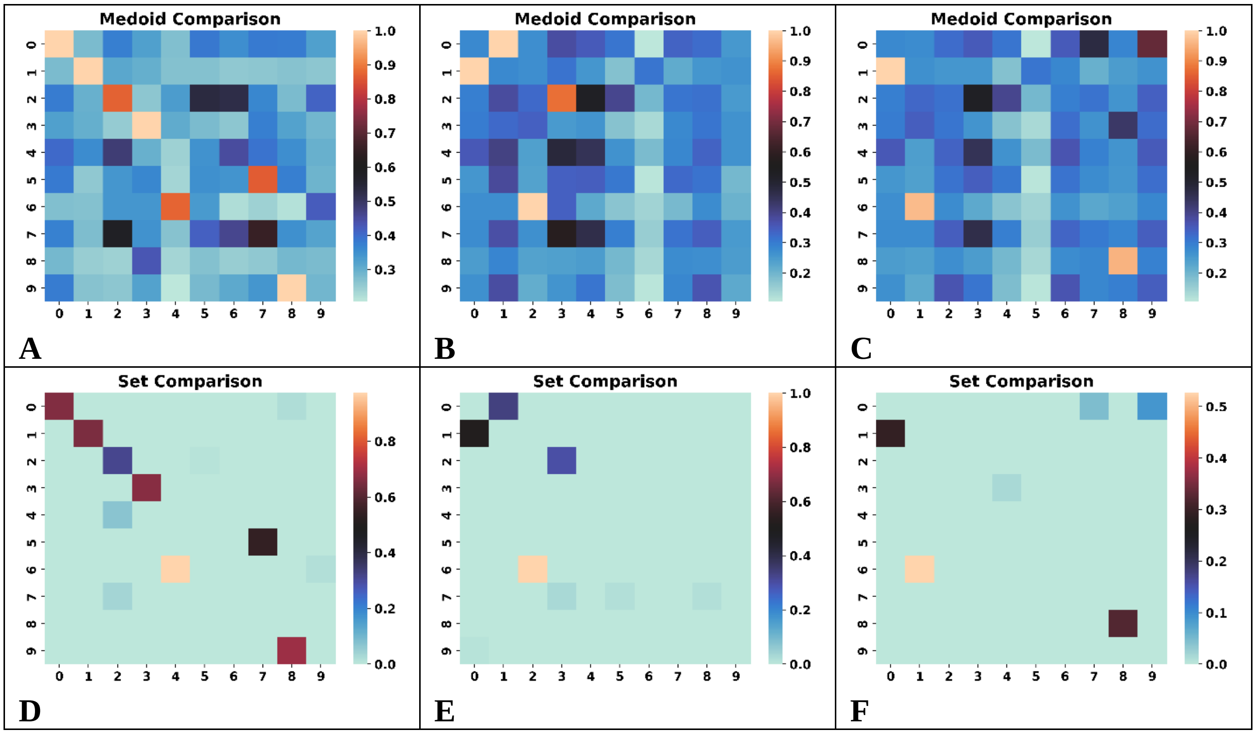


**Figure S6.6**: Comparison of the **A, B, C**: medoids, **D, E, F**: sets for the top populated clusters of the [6] ChEMBL subset using the BitBIRCH and BitBIRCH-folded (**A**, **D**: 1- fold, **B**, **E**: 2-fold, **C**, **F**: 3-fold) algorithms (similarity threshold = 0.65, min_size = 10).

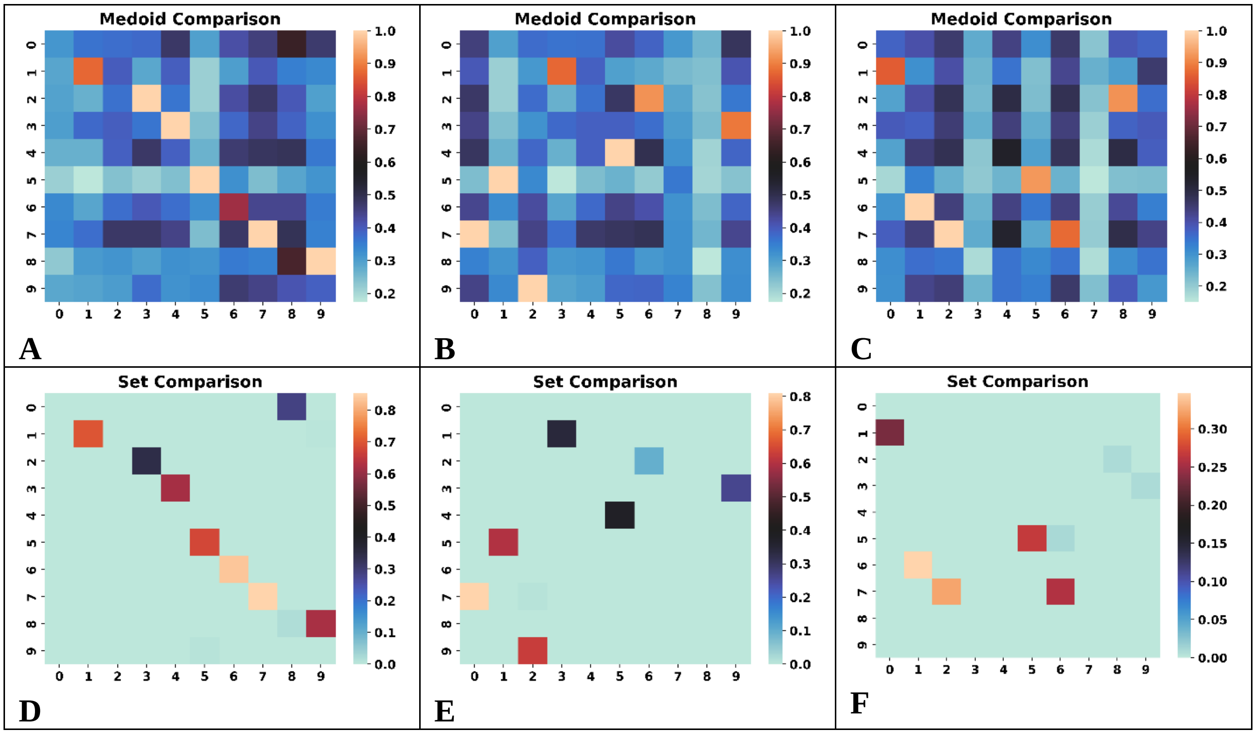


**Figure S6.7**: Comparison of the **A, B, C**: medoids, **D, E, F**: sets for the top populated clusters of the [7] ChEMBL subset using the BitBIRCH and BitBIRCH-folded (**A**, **D**: 1- fold, **B**, **E**: 2-fold, **C**, **F**: 3-fold) algorithms (similarity threshold = 0.65, min_size = 10).

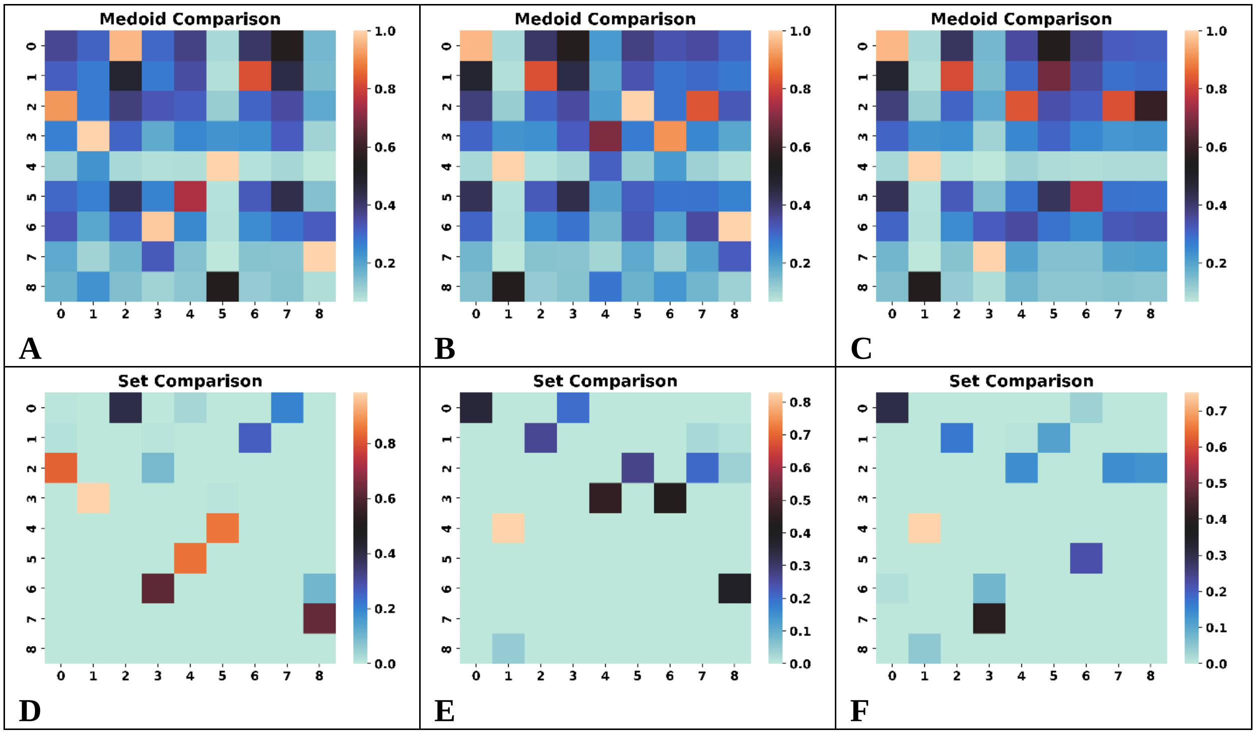

**Figure S6.8**: Comparison of the **A, B, C**: medoids, **D, E, F**: sets for the top populated clusters of the [8] ChEMBL subset using the BitBIRCH and BitBIRCH-folded (**A**, **D**: 1- fold, **B**, **E**: 2-fold, **C**, **F**: 3-fold) algorithms (similarity threshold = 0.65, min_size = 10).

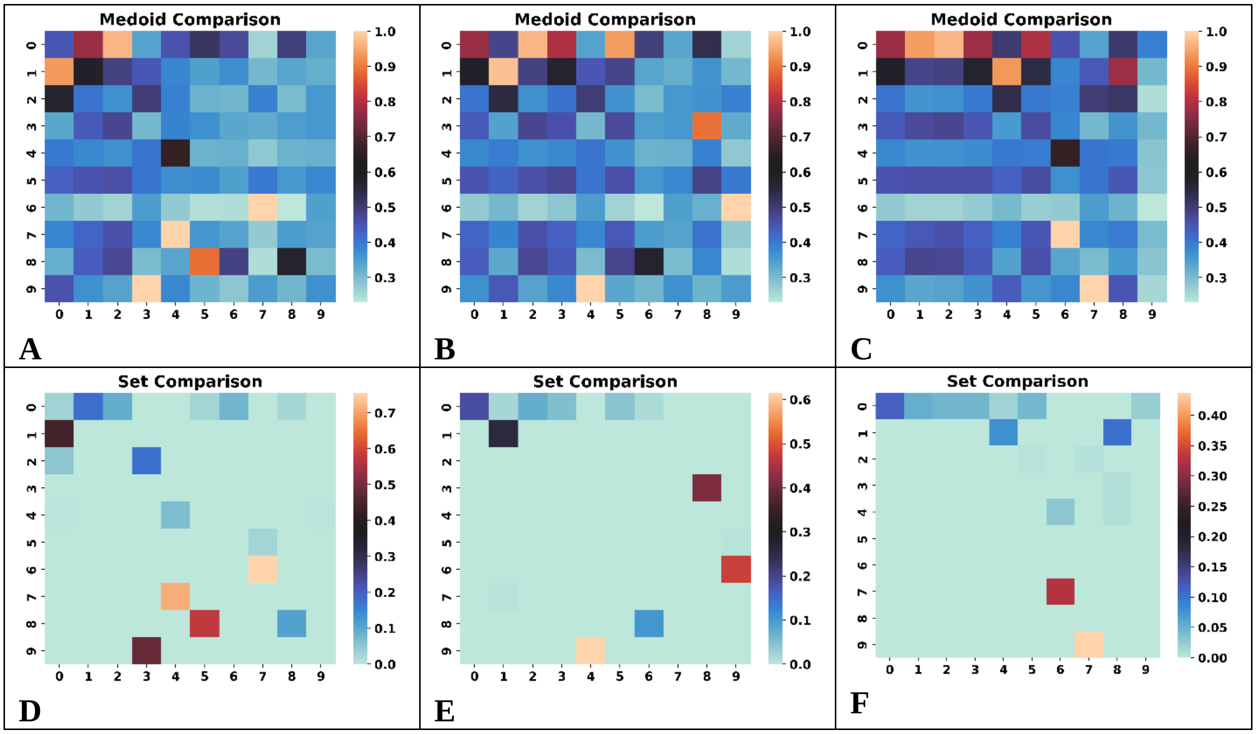


**Figure S6.9**: Comparison of the **A, B, C**: medoids, **D, E, F**: sets for the top populated clusters of the [9] ChEMBL subset using the BitBIRCH and BitBIRCH-folded (**A**, **D**: 1- fold, **B**, **E**: 2-fold, **C**, **F**: 3-fold) algorithms (similarity threshold = 0.65, min_size = 10).

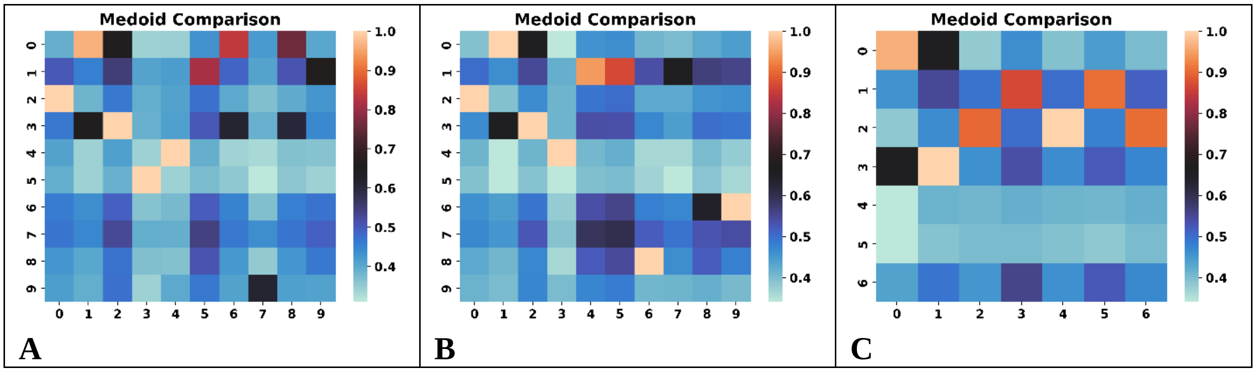

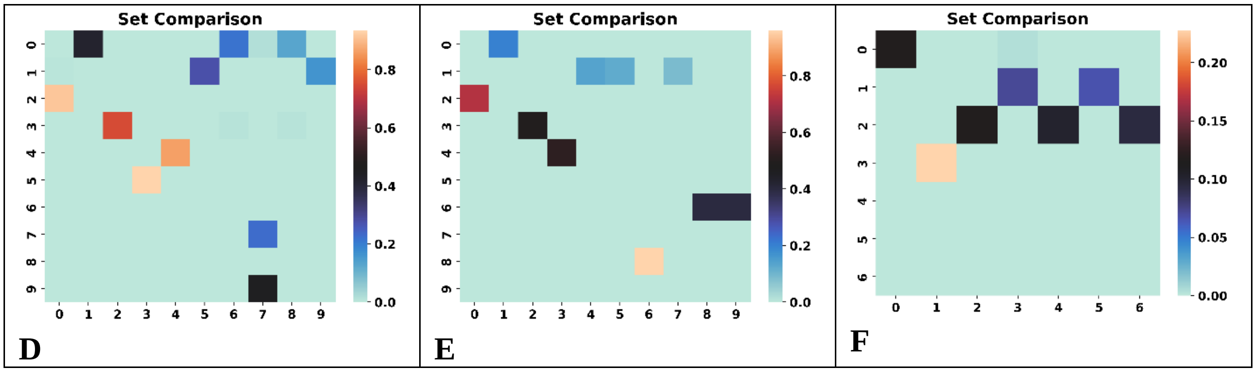


**Figure S6.10**: Comparison of the **A, B, C**: medoids, **D, E, F**: sets for the top populated clusters of the [10] ChEMBL subset using the BitBIRCH and BitBIRCH-folded (**A**, **D**: 1- fold, **B**, **E**: 2-fold, **C**, **F**: 3-fold) algorithms (similarity threshold = 0.65, min_size = 10).

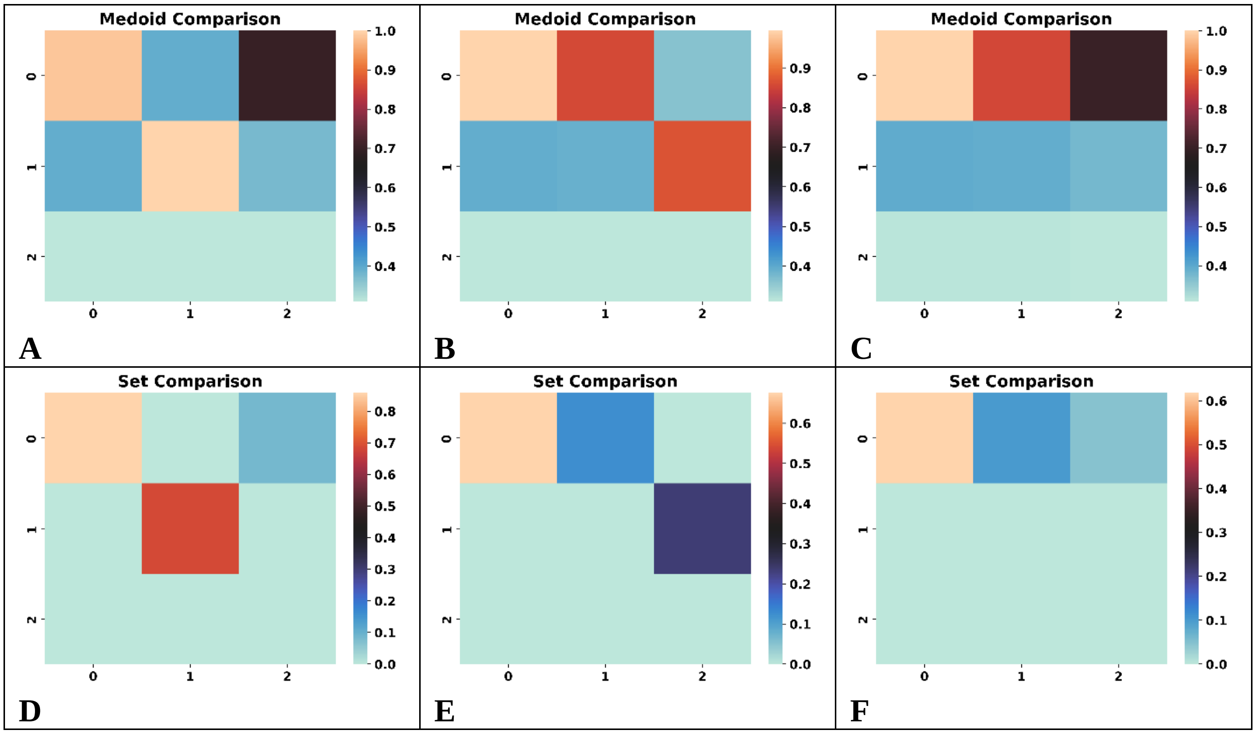


**Figure S6.11**: Comparison of the **A, B, C**: medoids, **D, E, F**: sets for the top populated clusters of the [11] ChEMBL subset using the BitBIRCH and BitBIRCH-folded (**A**, **D**: 1- fold, **B**, **E**: 2-fold, **C**, **F**: 3-fold) algorithms (similarity threshold = 0.65, min_size = 10).

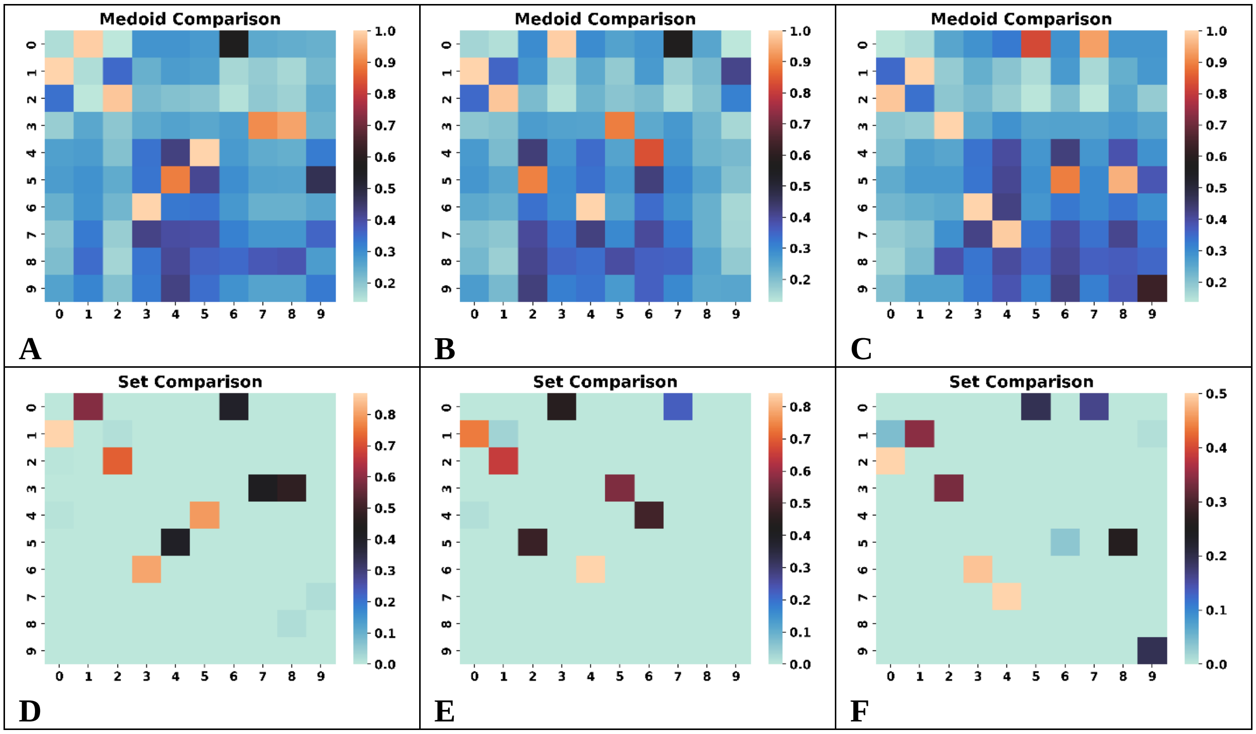


**Figure S6.12**: Comparison of the **A, B, C**: medoids, **D, E, F**: sets for the top populated clusters of the [12] ChEMBL subset using the BitBIRCH and BitBIRCH-folded (**A**, **D**: 1- fold, **B**, **E**: 2-fold, **C**, **F**: 3-fold) algorithms (similarity threshold = 0.65, min_size = 10).

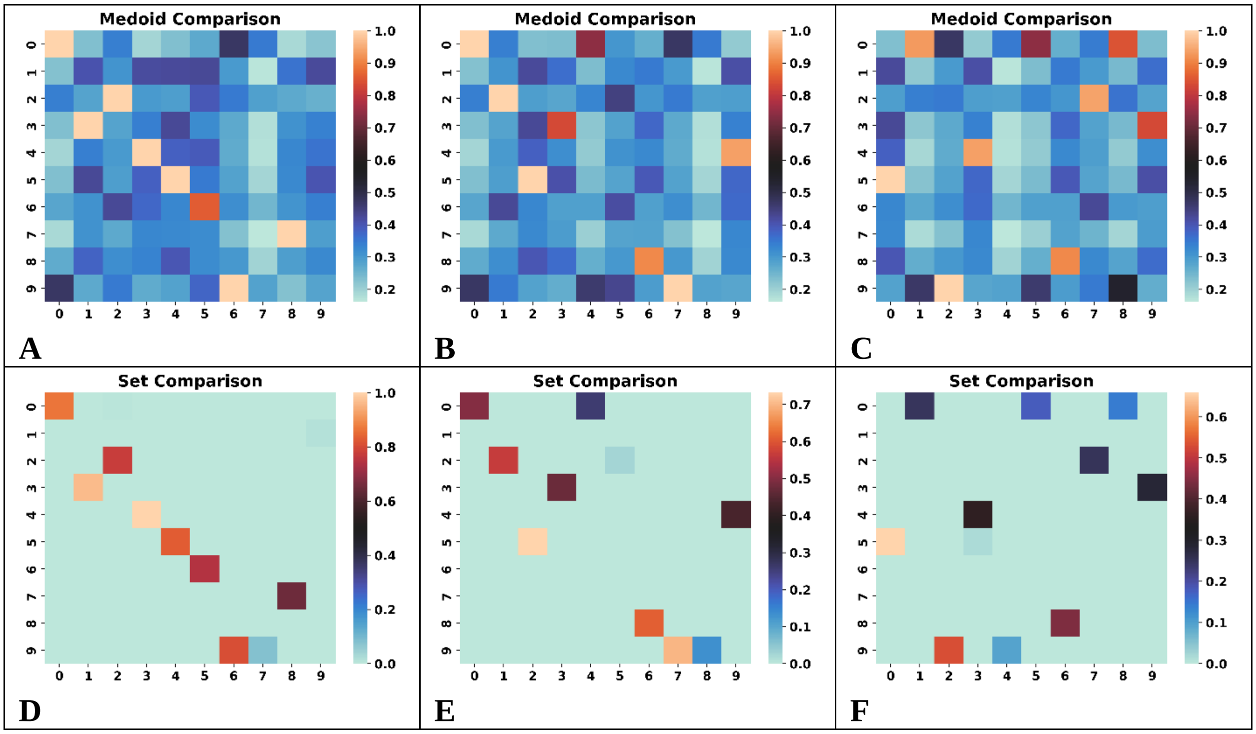

**Figure S6.13**: Comparison of the **A, B, C**: medoids, **D, E, F**: sets for the top populated clusters of the [13] ChEMBL subset using the BitBIRCH and BitBIRCH-folded (**A**, **D**: 1- fold, **B**, **E**: 2-fold, **C**, **F**: 3-fold) algorithms (similarity threshold = 0.65, min_size = 10).

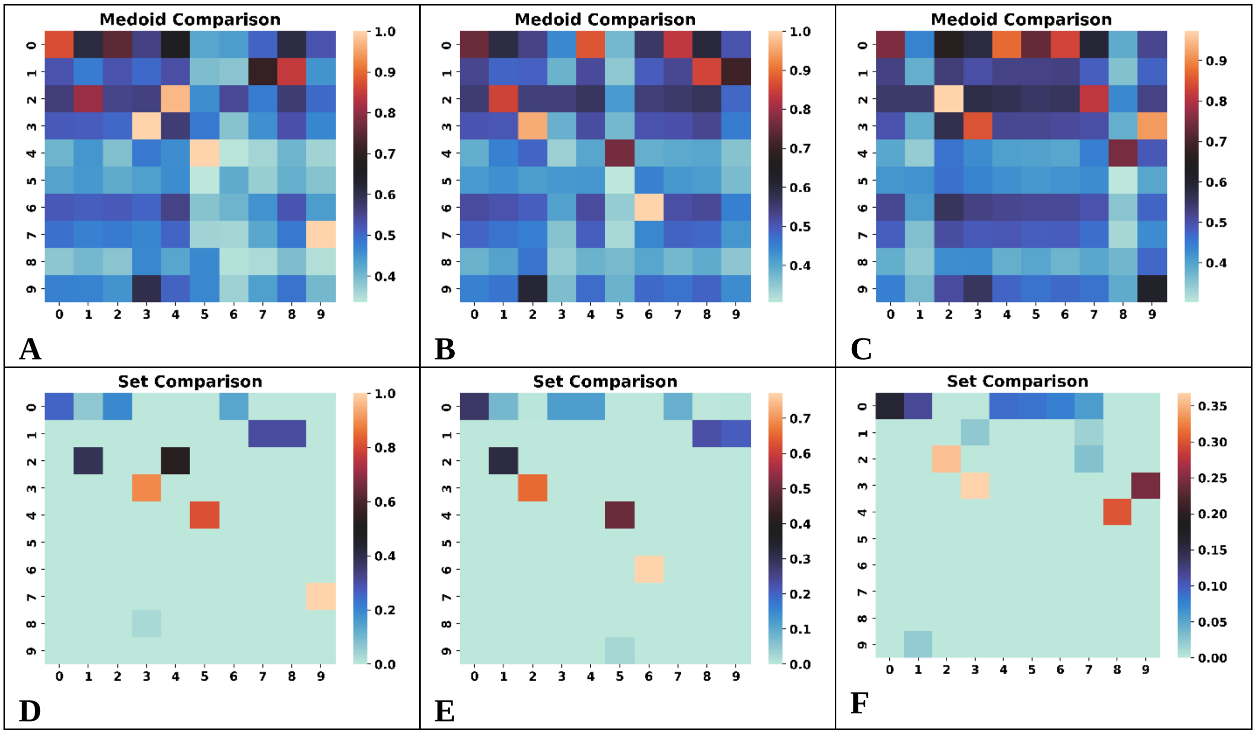


**Figure S6.14**: Comparison of the **A, B, C**: medoids, **D, E, F**: sets for the top populated clusters of the [14] ChEMBL subset using the BitBIRCH and BitBIRCH-folded (**A**, **D**: 1- fold, **B**, **E**: 2-fold, **C**, **F**: 3-fold) algorithms (similarity threshold = 0.65, min_size = 10).

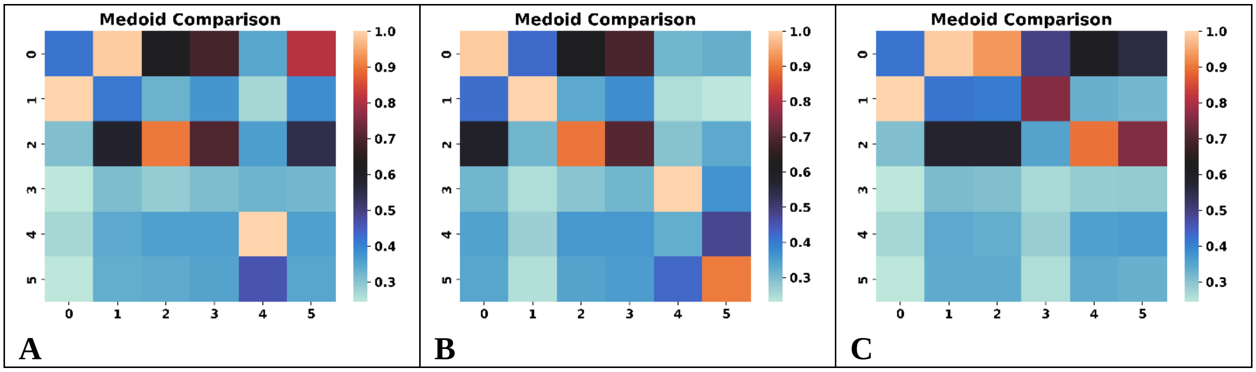

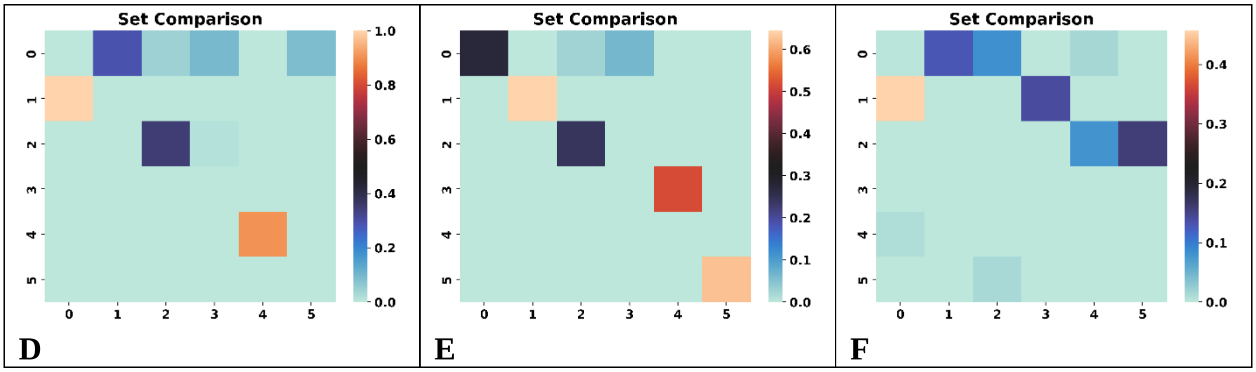


**Figure S6.15**: Comparison of the **A, B, C**: medoids, **D, E, F**: sets for the top populated clusters of the [15] ChEMBL subset using the BitBIRCH and BitBIRCH-folded (**A**, **D**: 1- fold, **B**, **E**: 2-fold, **C**, **F**: 3-fold) algorithms (similarity threshold = 0.65, min_size = 10).

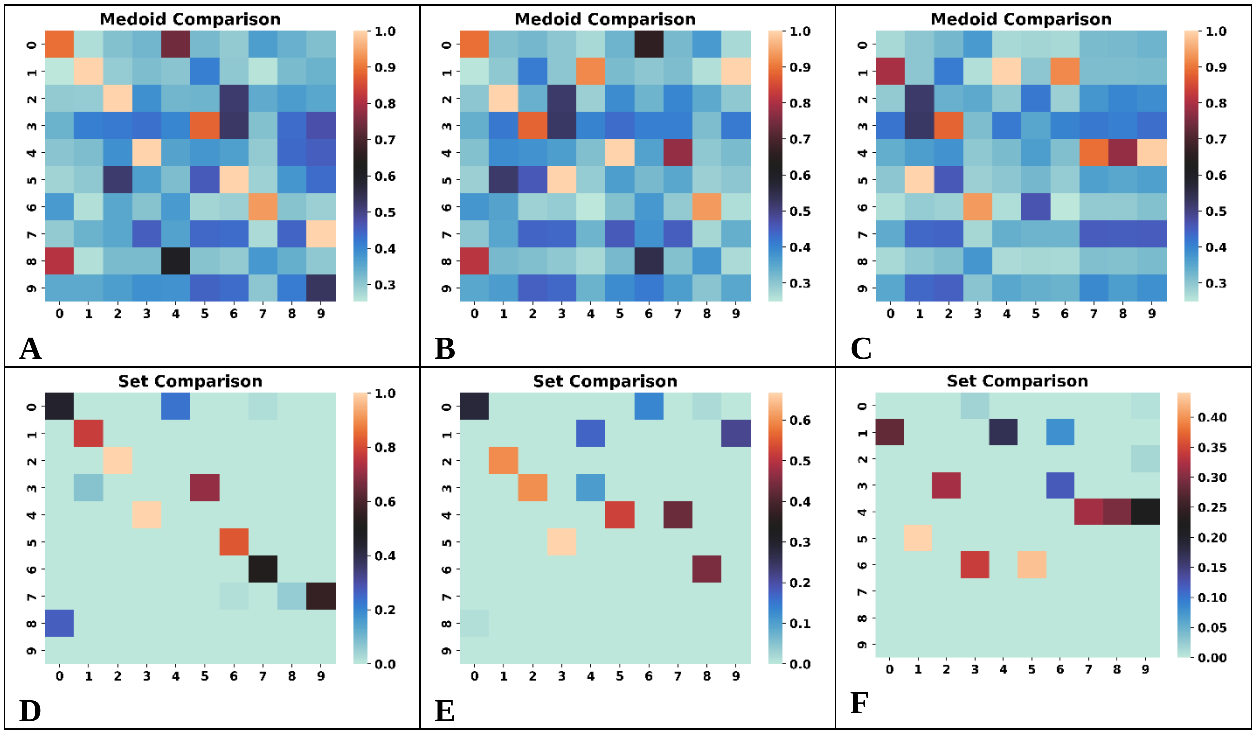


**Figure S6.16**: Comparison of the **A, B, C**: medoids, **D, E, F**: sets for the top populated clusters of the [16] ChEMBL subset using the BitBIRCH and BitBIRCH-folded (**A**, **D**: 1- fold, **B**, **E**: 2-fold, **C**, **F**: 3-fold) algorithms (similarity threshold = 0.65, min_size = 10).

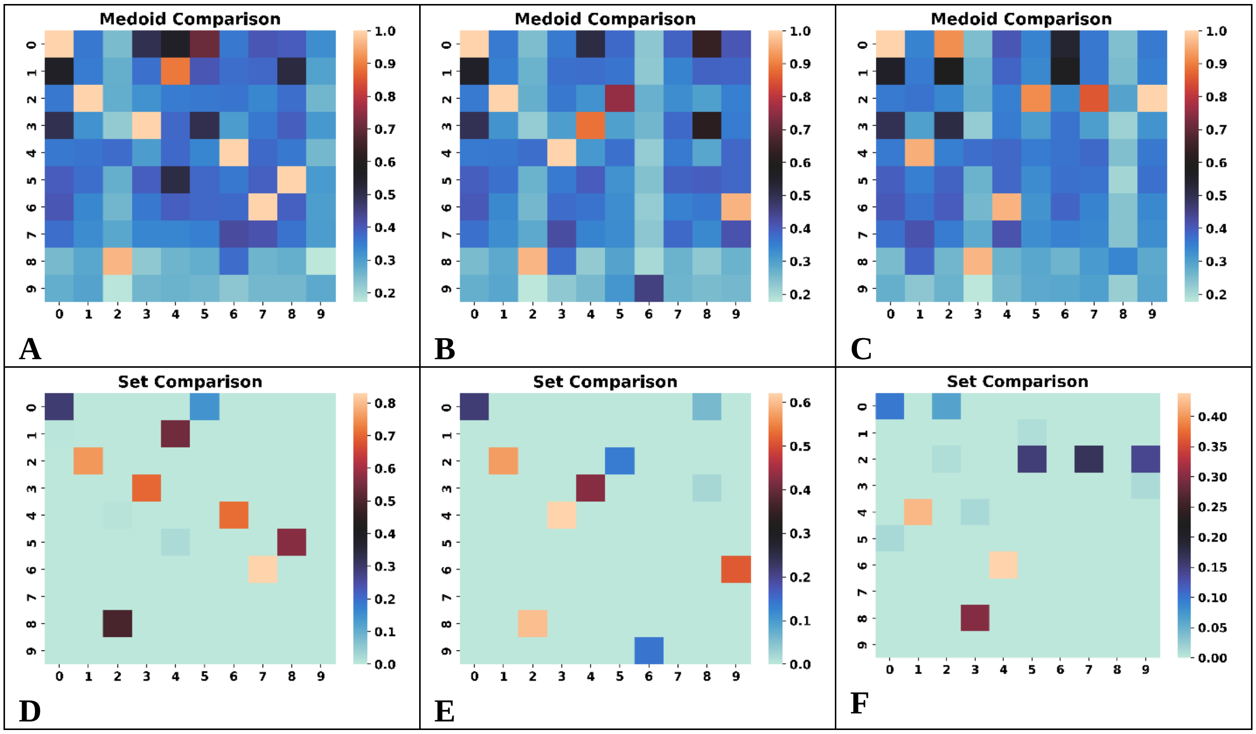


**Figure S6.17**: Comparison of the **A, B, C**: medoids, **D, E, F**: sets for the top populated clusters of the [17] ChEMBL subset using the BitBIRCH and BitBIRCH-folded (**A**, **D**: 1- fold, **B**, **E**: 2-fold, **C**, **F**: 3-fold) algorithms (similarity threshold = 0.65, min_size = 10).

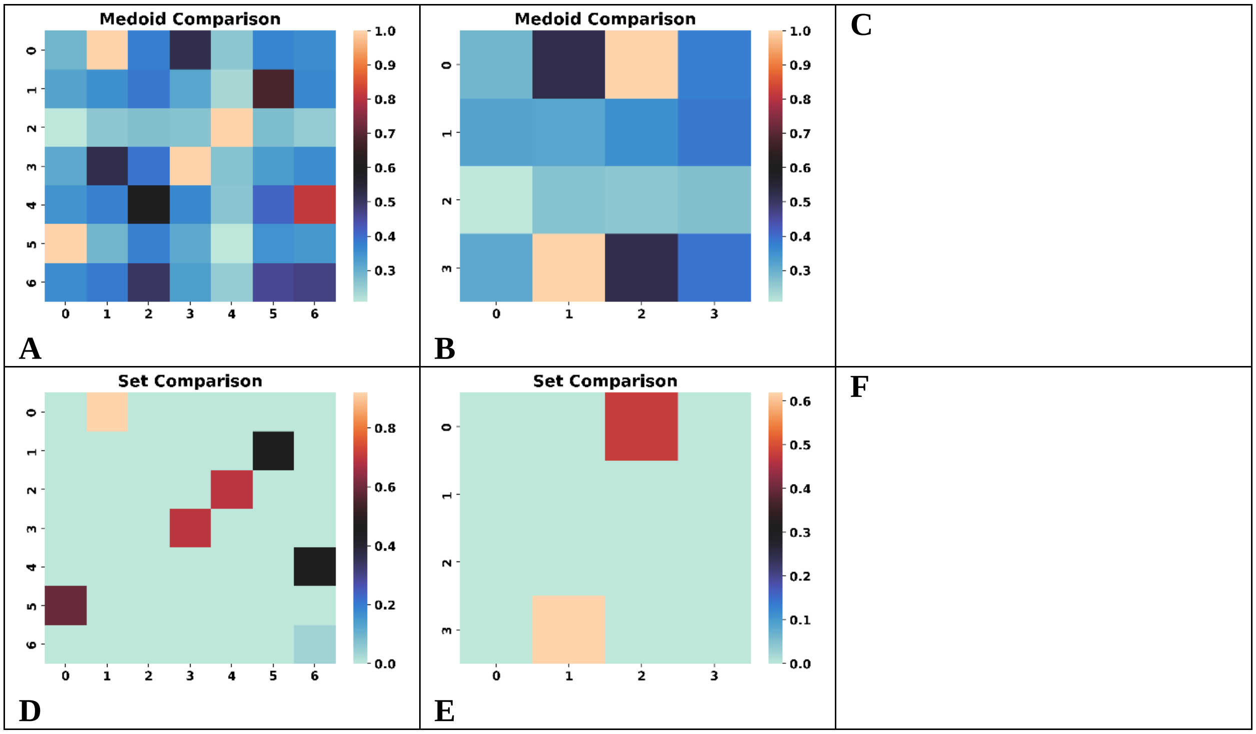

**Figure S6.18**: Comparison of the **A, B, C**: medoids, **D, E, F**: sets for the top populated clusters of the [18] ChEMBL subset using the BitBIRCH and BitBIRCH-folded (**A**, **D**: 1- fold, **B**, **E**: 2-fold, **C**, **F**: 3-fold) algorithms (similarity threshold = 0.65, min_size = 10).

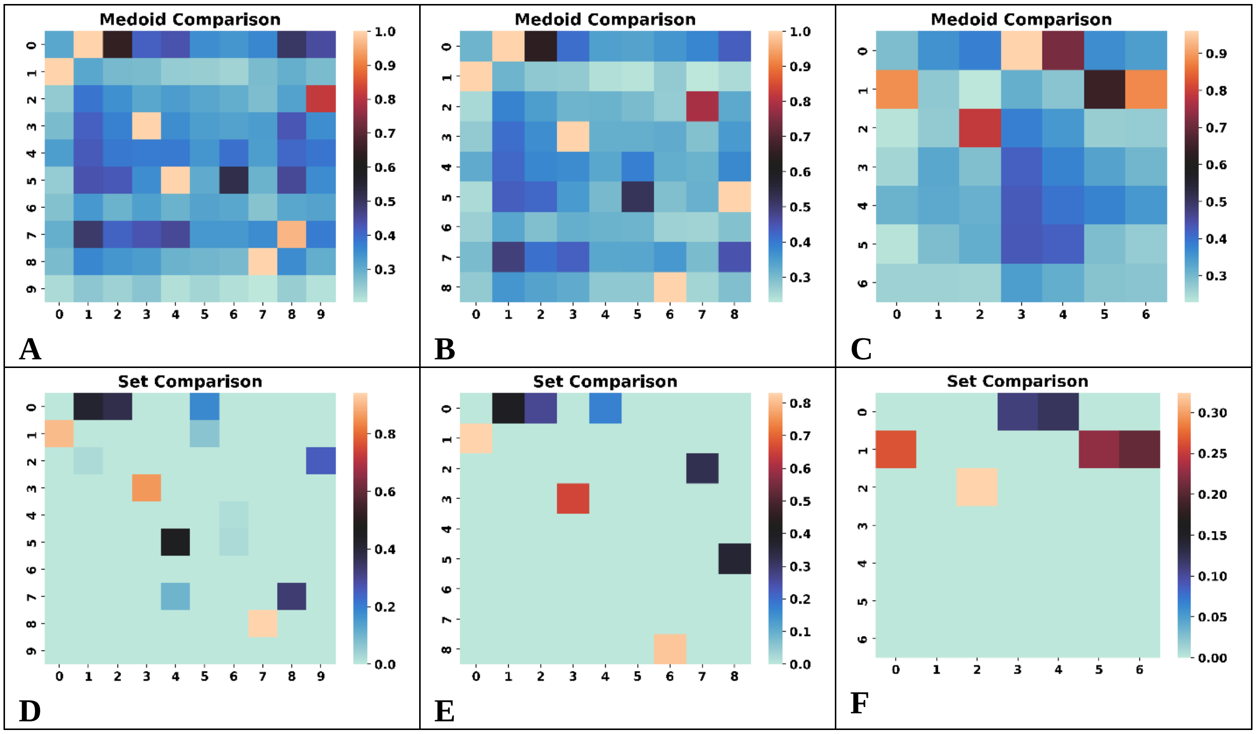


**Figure S6.19**: Comparison of the **A, B, C**: medoids, **D, E, F**: sets for the top populated clusters of the [19] ChEMBL subset using the BitBIRCH and BitBIRCH-folded (**A**, **D**: 1- fold, **B**, **E**: 2-fold, **C**, **F**: 3-fold) algorithms (similarity threshold = 0.65, min_size = 10).

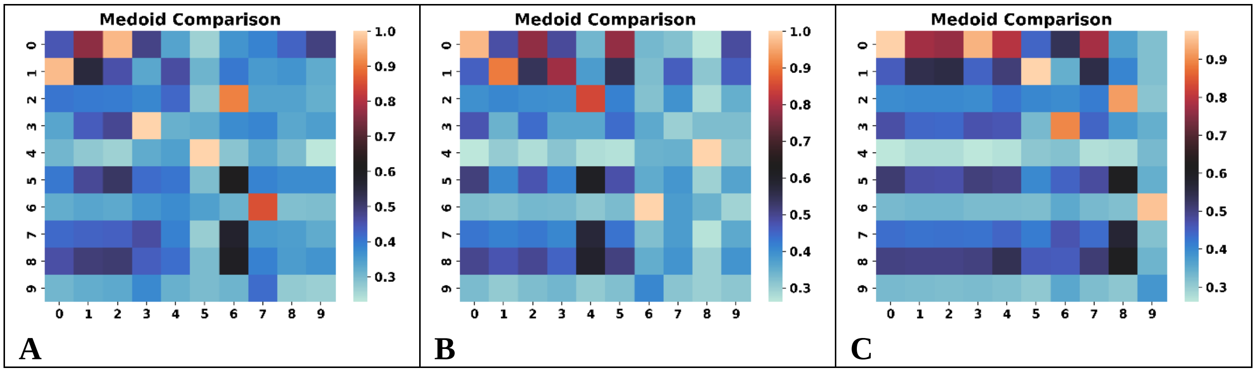

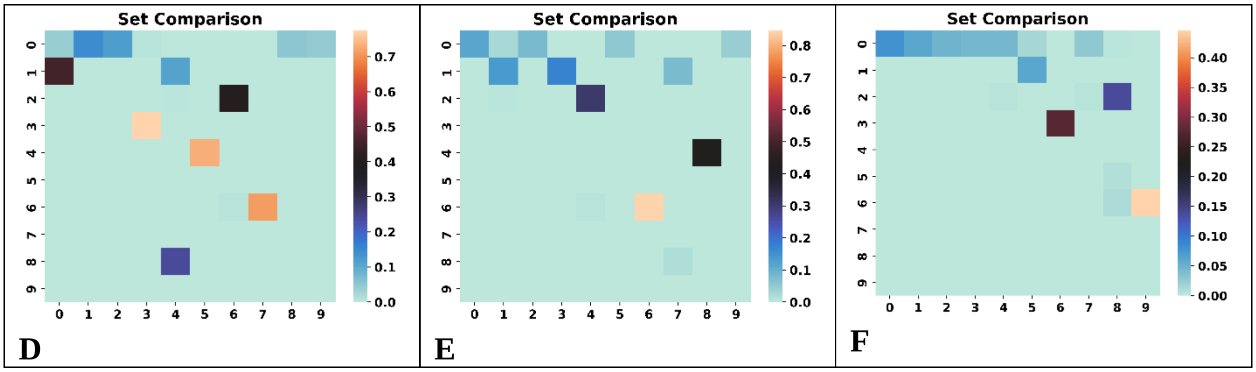


**Figure S6.20**: Comparison of the **A, B, C**: medoids, **D, E, F**: sets for the top populated clusters of the [20] ChEMBL subset using the BitBIRCH and BitBIRCH-folded (**A**, **D**: 1- fold, **B**, **E**: 2-fold, **C**, **F**: 3-fold) algorithms (similarity threshold = 0.65, min_size = 10).

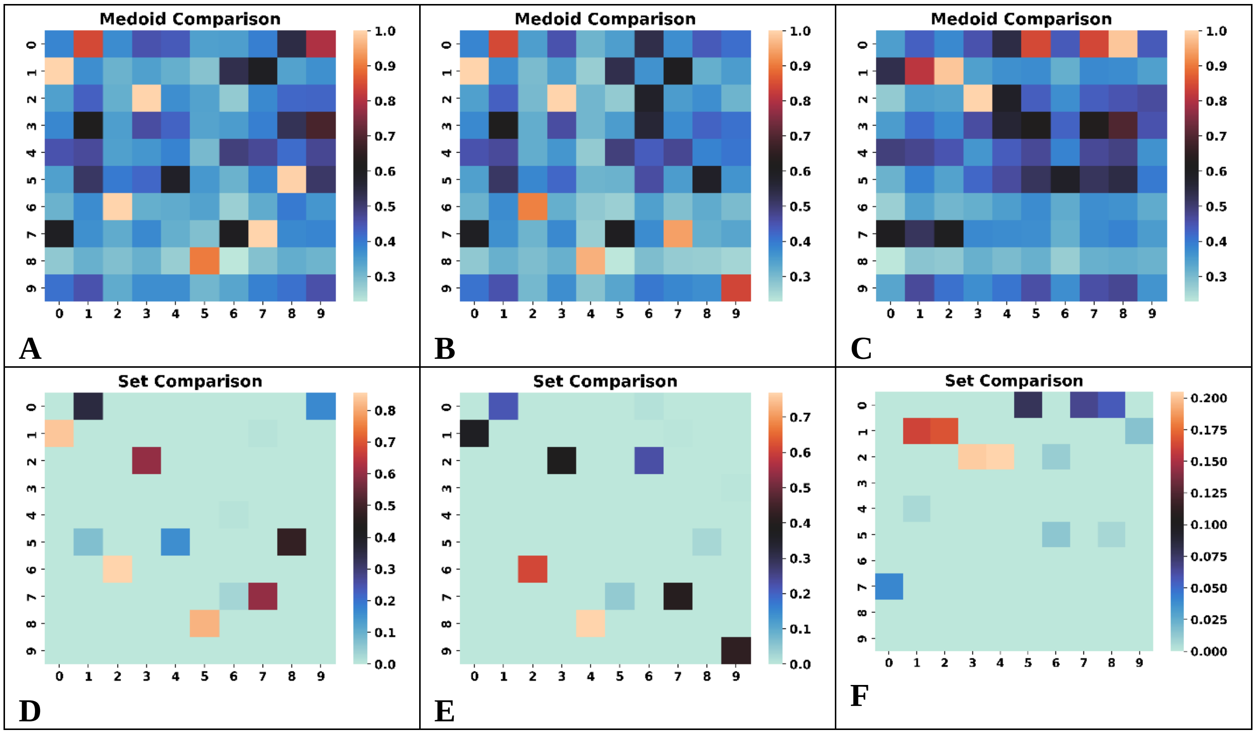


**Figure S6.21**: Comparison of the **A, B, C**: medoids, **D, E, F**: sets for the top populated clusters of the [21] ChEMBL subset using the BitBIRCH and BitBIRCH-folded (**A**, **D**: 1- fold, **B**, **E**: 2-fold, **C**, **F**: 3-fold) algorithms (similarity threshold = 0.65, min_size = 10).

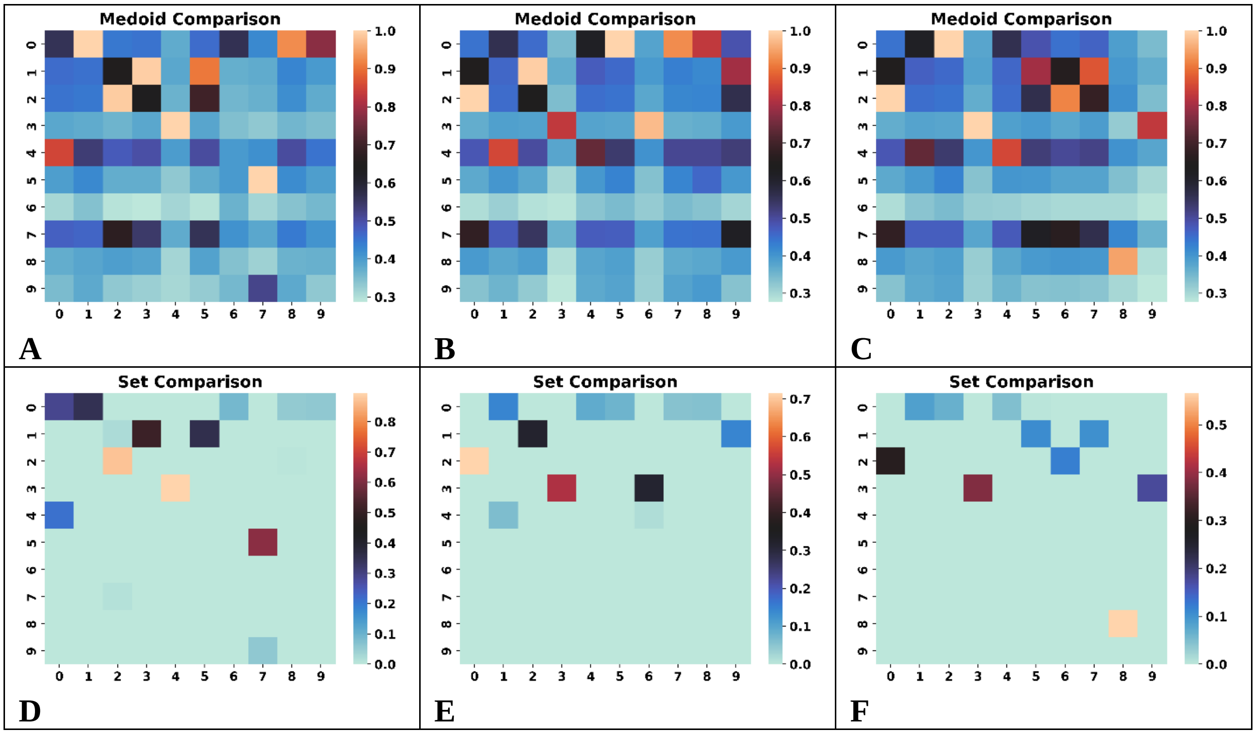


**Figure S6.22**: Comparison of the **A, B, C**: medoids, **D, E, F**: sets for the top populated clusters of the [22] ChEMBL subset using the BitBIRCH and BitBIRCH-folded (**A**, **D**: 1- fold, **B**, **E**: 2-fold, **C**, **F**: 3-fold) algorithms (similarity threshold = 0.65, min_size = 10).

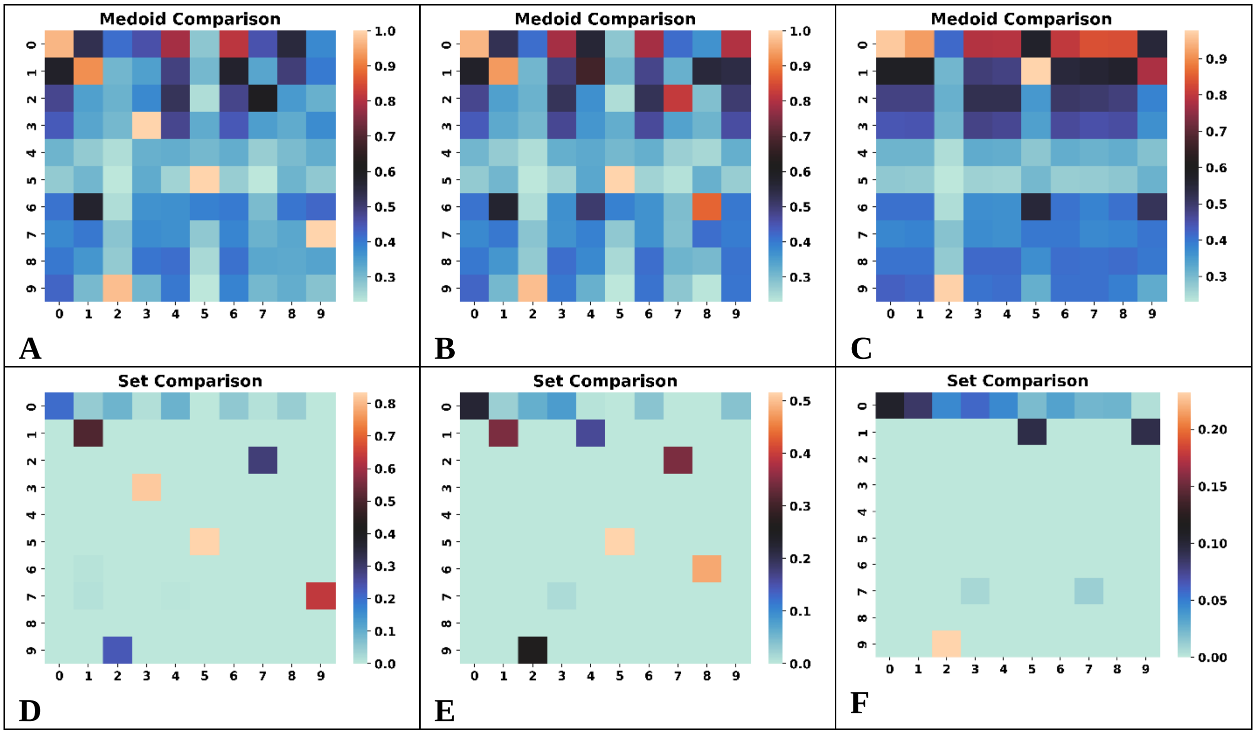

**Figure S6.23**: Comparison of the **A, B, C**: medoids, **D, E, F**: sets for the top populated clusters of the [23] ChEMBL subset using the BitBIRCH and BitBIRCH-folded (**A**, **D**: 1- fold, **B**, **E**: 2-fold, **C**, **F**: 3-fold) algorithms (similarity threshold = 0.65, min_size = 10).

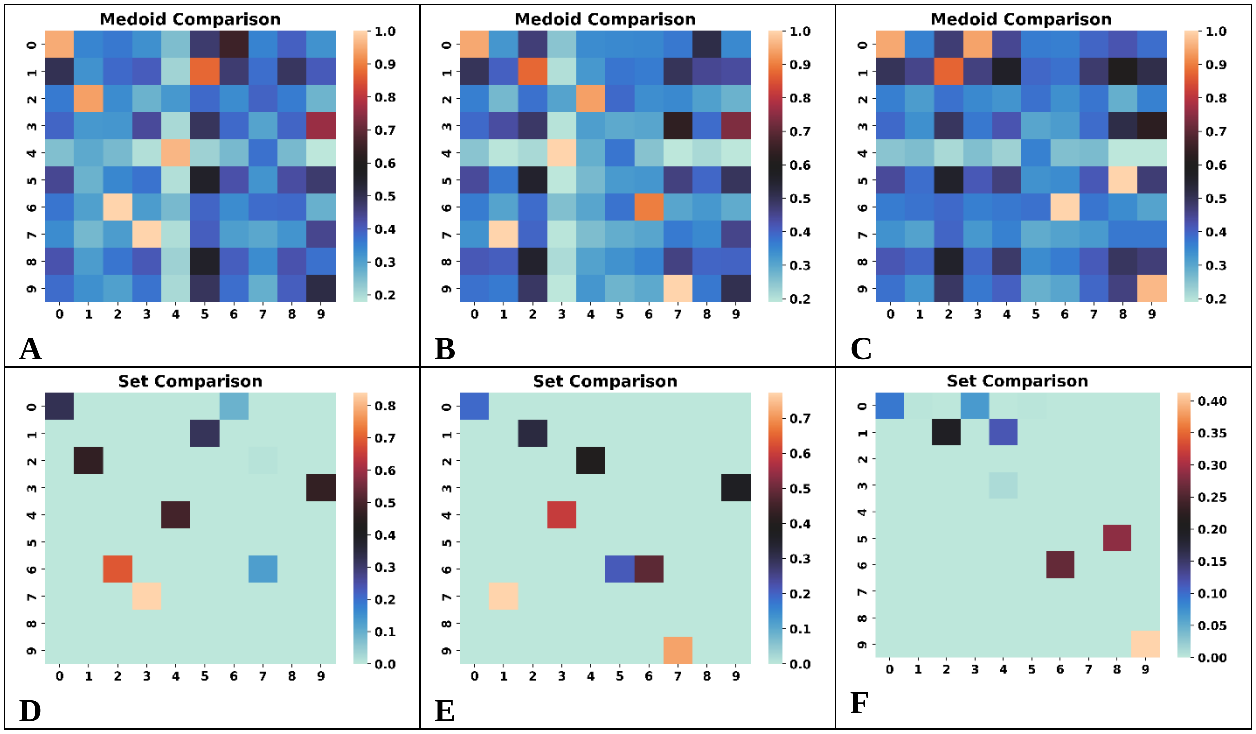


**Figure S6.24**: Comparison of the **A, B, C**: medoids, **D, E, F**: sets for the top populated clusters of the [24] ChEMBL subset using the BitBIRCH and BitBIRCH-folded (**A**, **D**: 1- fold, **B**, **E**: 2-fold, **C**, **F**: 3-fold) algorithms (similarity threshold = 0.65, min_size = 10).

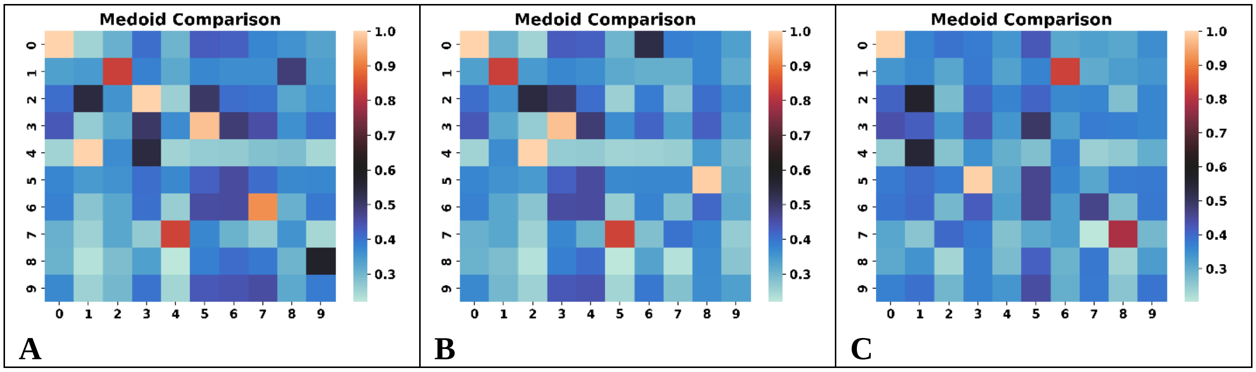

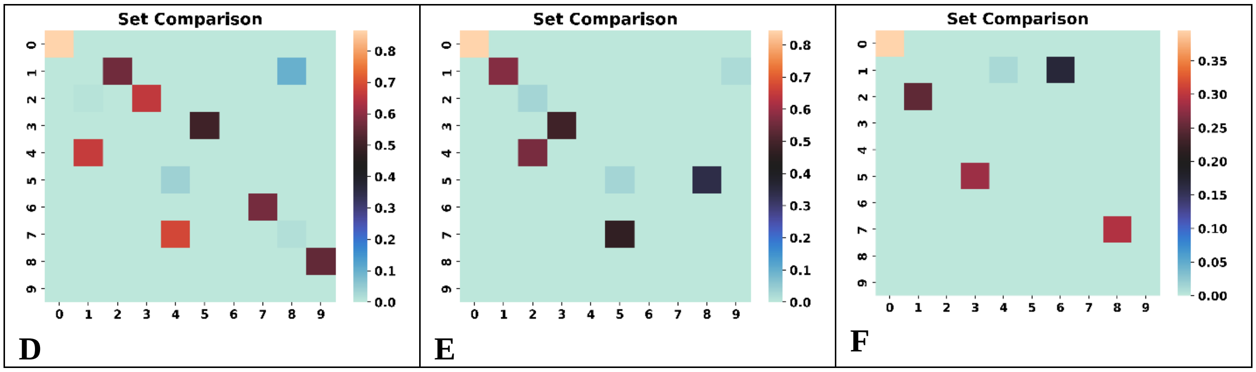


**Figure S6.25**: Comparison of the **A, B, C**: medoids, **D, E, F**: sets for the top populated clusters of the [25] ChEMBL subset using the BitBIRCH and BitBIRCH-folded (**A**, **D**: 1- fold, **B**, **E**: 2-fold, **C**, **F**: 3-fold) algorithms (similarity threshold = 0.65, min_size = 10).

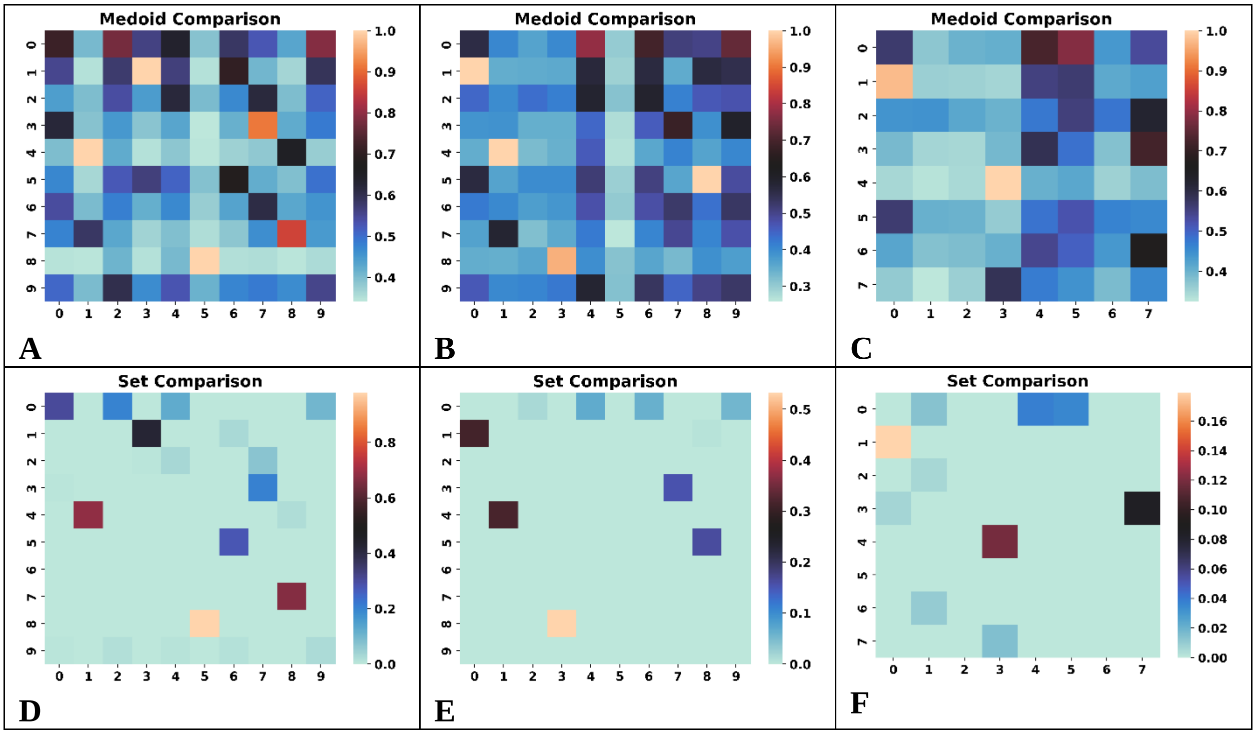


**Figure S6.26**: Comparison of the **A, B, C**: medoids, **D, E, F**: sets for the top populated clusters of the [26] ChEMBL subset using the BitBIRCH and BitBIRCH-folded (**A**, **D**: 1- fold, **B**, **E**: 2-fold, **C**, **F**: 3-fold) algorithms (similarity threshold = 0.65, min_size = 10).

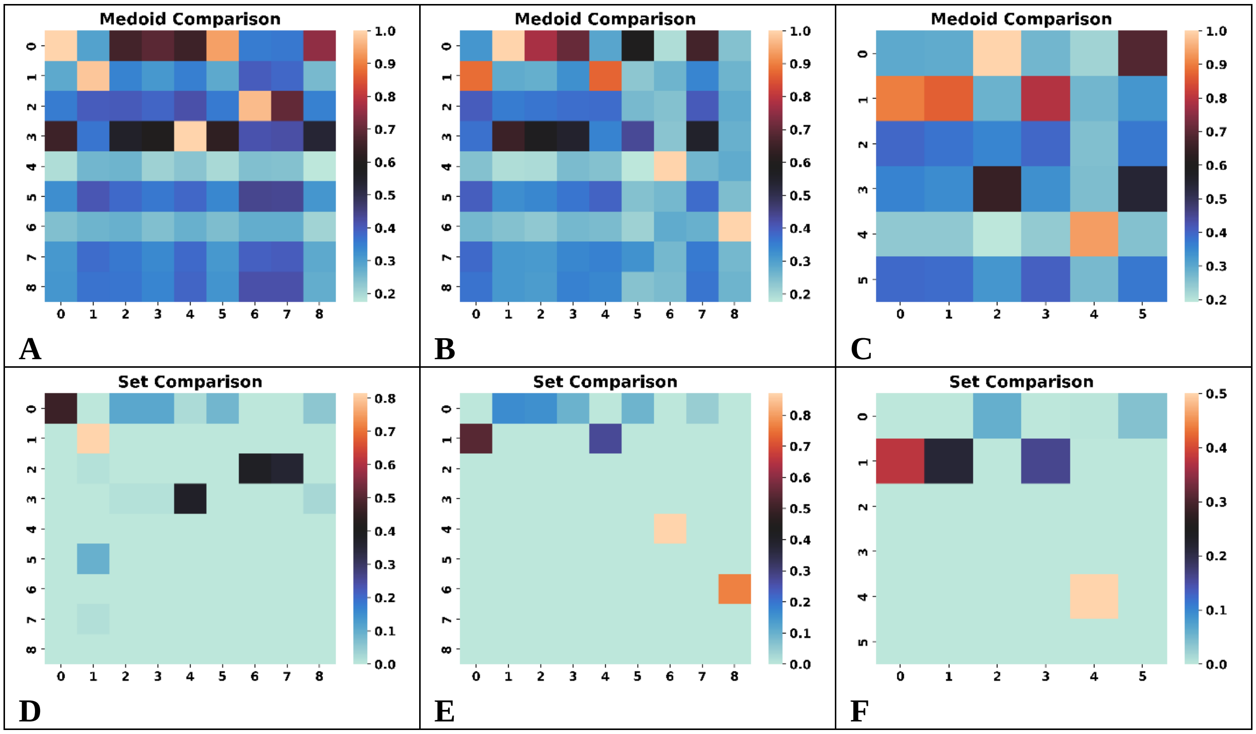


**Figure S6.27**: Comparison of the **A, B, C**: medoids, **D, E, F**: sets for the top populated clusters of the [27] ChEMBL subset using the BitBIRCH and BitBIRCH-folded (**A**, **D**: 1- fold, **B**, **E**: 2-fold, **C**, **F**: 3-fold) algorithms (similarity threshold = 0.65, min_size = 10).

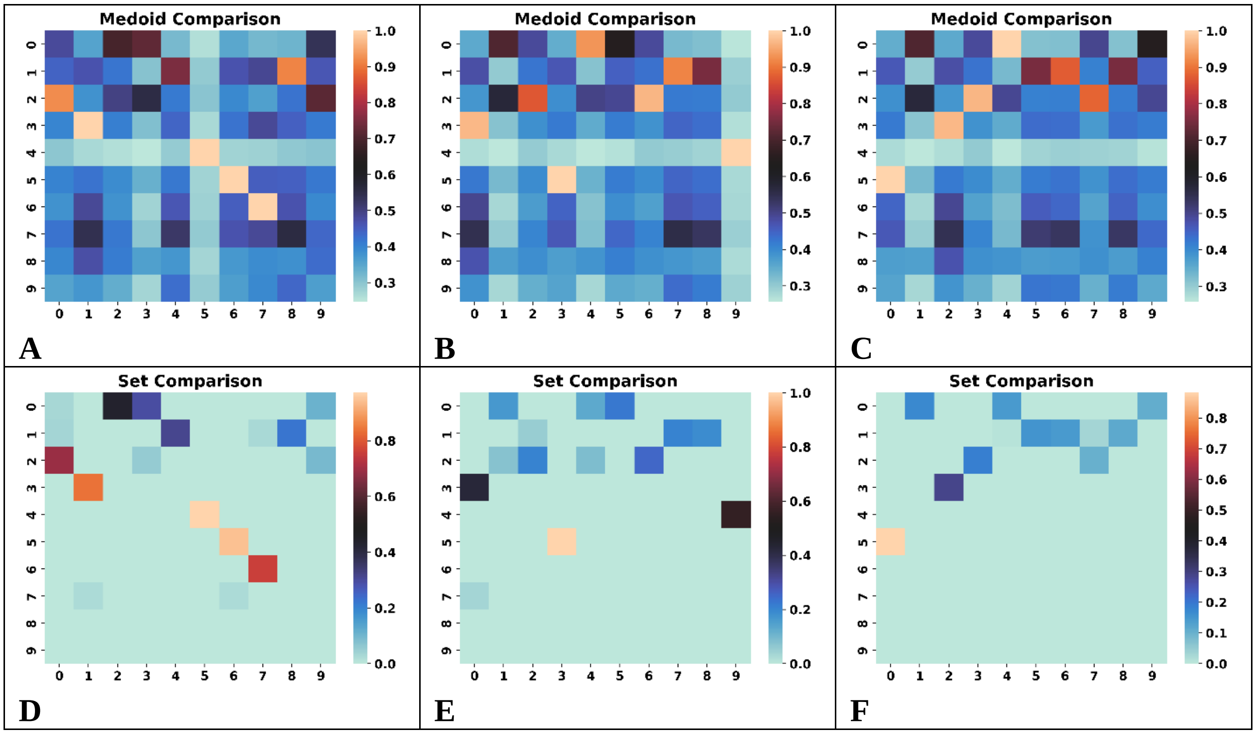

**Figure S6.28**: Comparison of the **A, B, C**: medoids, **D, E, F**: sets for the top populated clusters of the [28] ChEMBL subset using the BitBIRCH and BitBIRCH-folded (**A**, **D**: 1- fold, **B**, **E**: 2-fold, **C**, **F**: 3-fold) algorithms (similarity threshold = 0.65, min_size = 10).

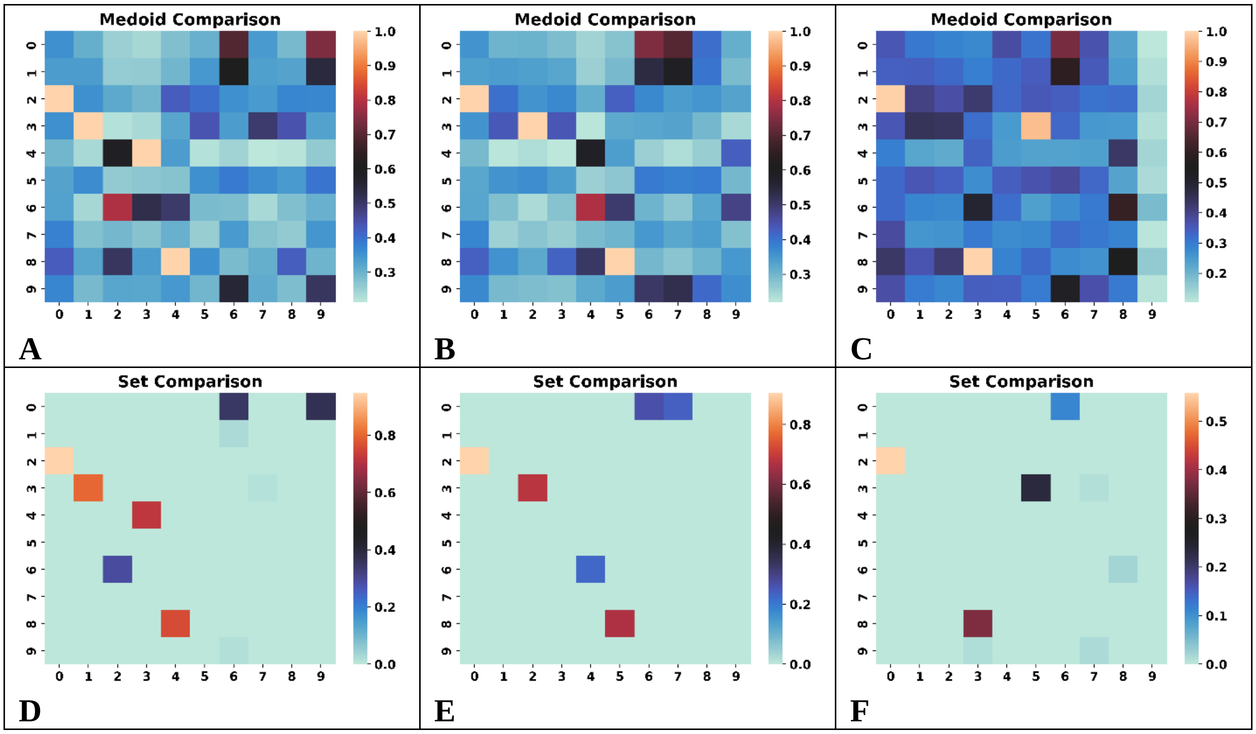


**Figure S6.29**: Comparison of the **A, B, C**: medoids, **D, E, F**: sets for the top populated clusters of the [29] ChEMBL subset using the BitBIRCH and BitBIRCH-folded (**A**, **D**: 1- fold, **B**, **E**: 2-fold, **C**, **F**: 3-fold) algorithms (similarity threshold = 0.65, min_size = 10).

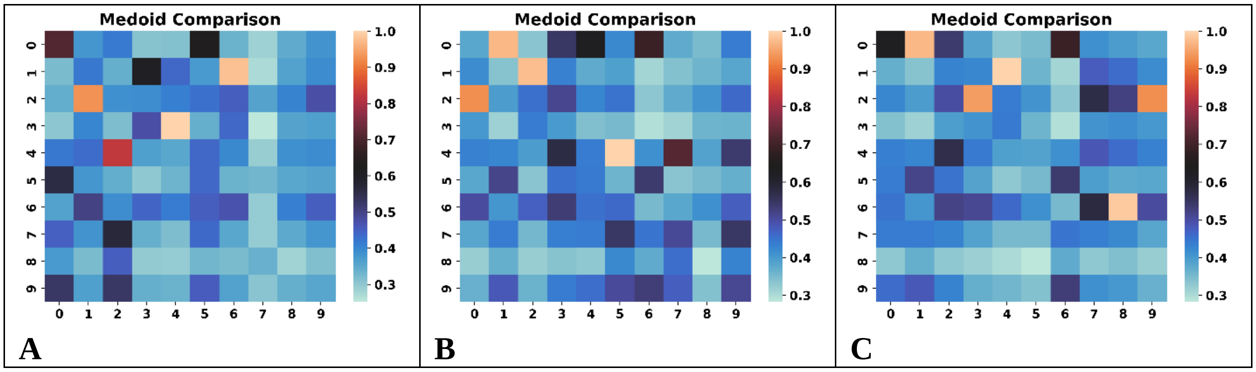

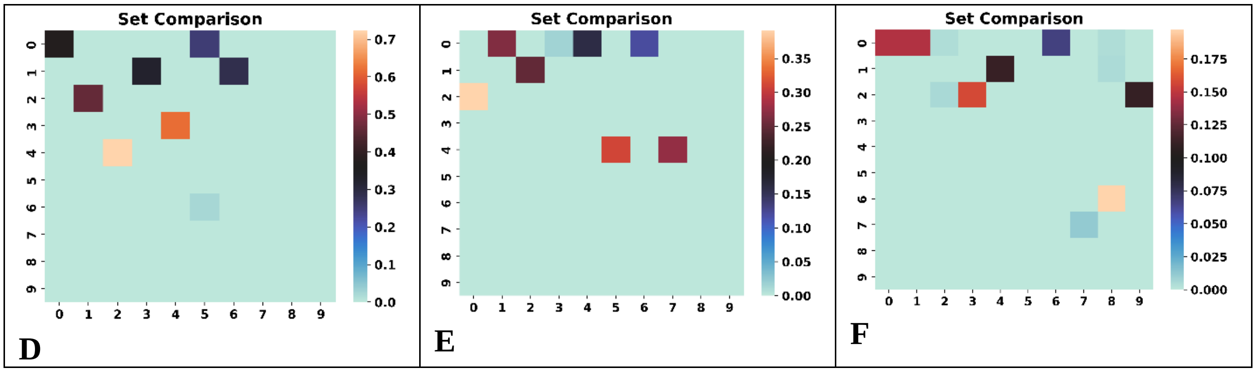


**Figure S6.30**: Comparison of the **A, B, C**: medoids, **D, E, F**: sets for the top populated clusters of the [30] ChEMBL subset using the BitBIRCH and BitBIRCH-folded (**A**, **D**: 1- fold, **B**, **E**: 2-fold, **C**, **F**: 3-fold) algorithms (similarity threshold = 0.65, min_size = 10).

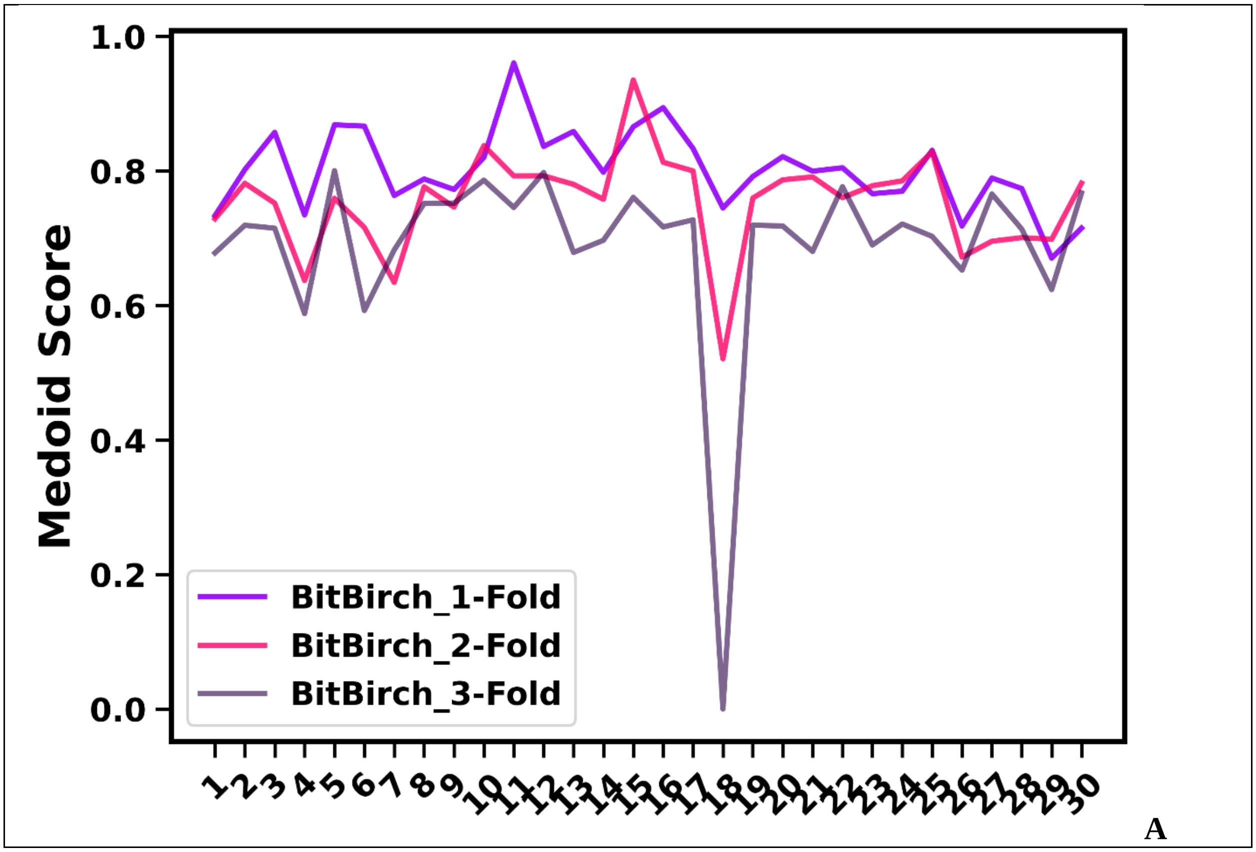

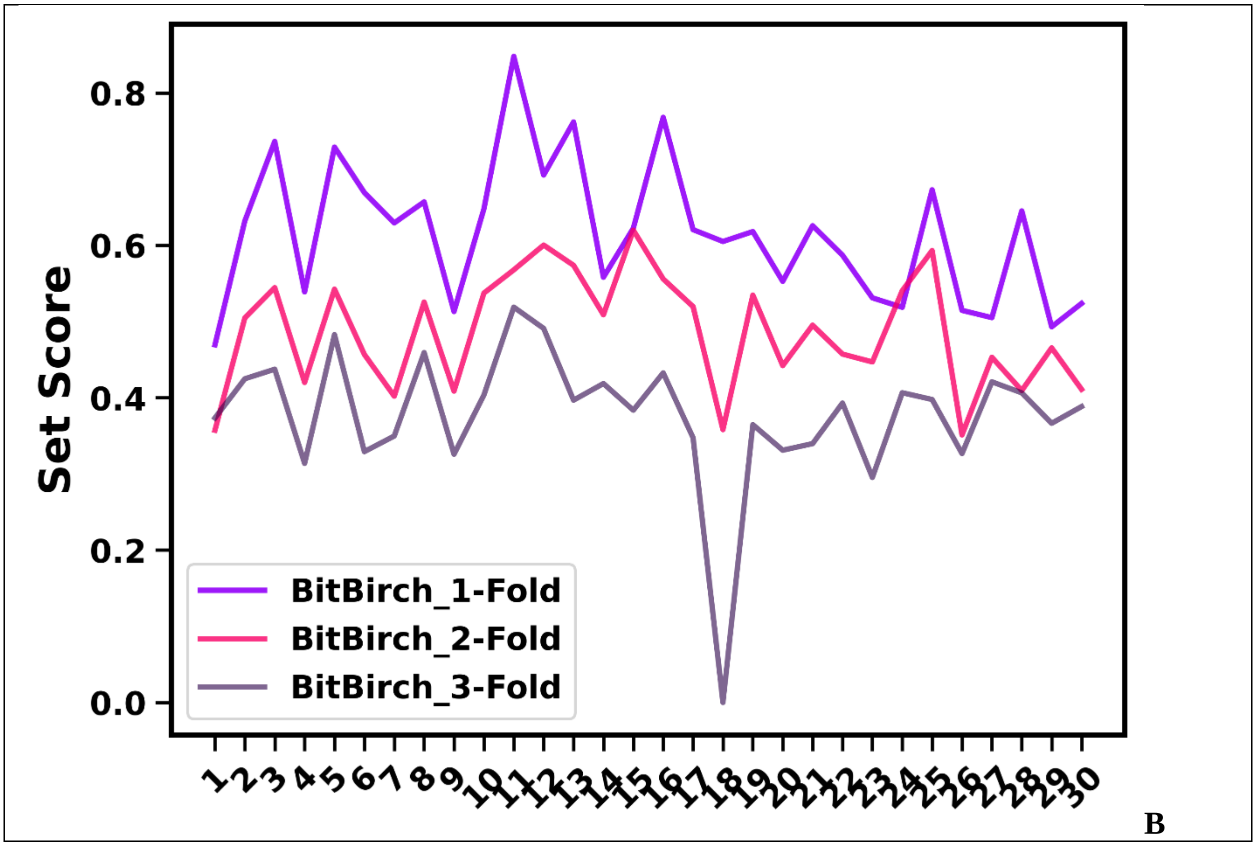


**Figure S6.31**: Summary of the Medoid (A) and Set (B) scores for the 30 ChEMBL subsets for the BitBIRCH and BitBIRCH-folded algorithms.

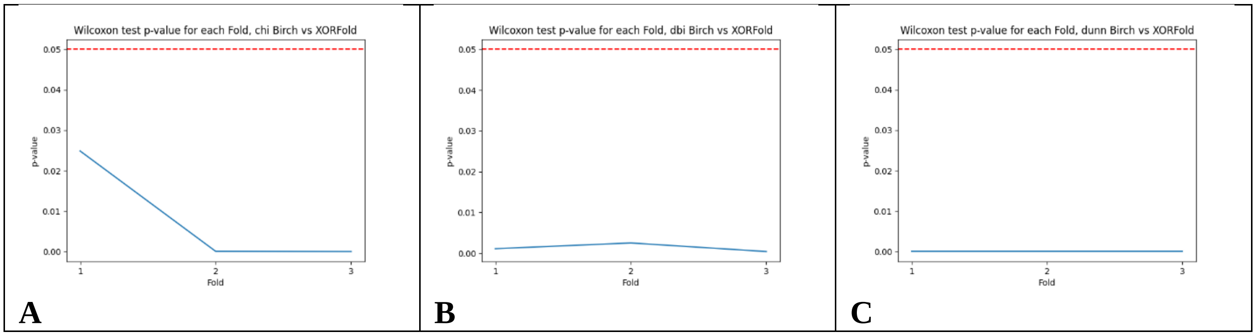


**Figure S6.32**: Wilcoxon two-sided tests results comparing the BitBIRCH and BitBIRCH-folded methods for the **A**: CHI (Ha: BB < BB-fold), **B**: DBI (Ha: BB > BB-fold) , and **C**: DI (Ha: BB > BB-fold).

### S7: Ultra large libraries

The billion molecules were obtained from several randomly selected tranches with more than 1 million molecules from the ZINC22-2D database (https://cartblanche.docking.org/tranches/2d).

From each selected tranche, $\lfloor n\_molecules \ / \ 1{,}000{,}000 \rfloor$ subsets were taken in order of apparition. The remaining molecules from each tranche were not included in our study. All subsets were included, except for H27, H28 and H29, only the subsets to complete the billion were used.

**Table S7.1**: Used tranches from the ZINC22 database codes, number of molecules in each tranche and number of 1 million subsets used.

| ZINC22 Tranche | Number of molecules | Number of subsets used (1 mill. molecules each) |
|---|---|---|
| H15M000 | 1,487,831 | 1 |
| H17M100 | 1,094,594 | 1 |
| H18P090 | 2,524,328 | 2 |
| H20P000 | 2,925,212 | 2 |
| H20P140 | 7,022,735 | 7 |
| H21P260 | 10,716,148 | 10 |
| H22P470 | 2,835,583 | 2 |
| H22P190 | 25,318,774 | 25 |
| H22P100 | 19,790,141 | 19 |
| H23P000 | 13,076,376 | 13 |
| H23P280 | 27,504,053 | 27 |
| H24P130 | 53,244,587 | 53 |
| H25P410 | 26,304,860 | 26 |
| H26P470 | 17,859,872 | 17 |
| H26M200 | 12,760,298 | 12 |
| H26P320 | 125,434,222 | 125 |
| H27P210 | 275,242,750 | 267* |
| H28P280 | 373,968,228 | 340* |
| H29P600 | 70,902,324 | 51* |
| TOTAL | --- | 1000 |
